# Alcohol, Opioid, and Combined Alcohol and Opioid Use Disorders Affect Shared and Unique Pathways: A Proteomic Analysis of Postmortem Brains

**DOI:** 10.1101/2022.05.30.494031

**Authors:** Edison Leung, Laura Stertz, Antonio L. Teixeira, Thomas D. Meyer, Sudhakar Selvaraj, Consuelo Walss-Bass

**Affiliations:** Louis. A. Faillace, MD, Department of Psychiatry and Behavioral Sciences, McGovern Medical School, The University of Texas Health Science Center at Houston, 1941 East Rd, Houston, TX 77054

**Keywords:** Alcohol Use Disorder, Opioid Use Disorder, Polysubstance Use Disorder, Proteomics, Substance use

## Abstract

Substance misuse is a major burden for not only patients but the healthcare system. Understanding how substance misuse alters a patient’s genome, proteome, and neurocircuitry is an important step toward identifying targets for treatments. In this exploratory study, we performed a proteomic analysis of postmortem brain from individuals with substance misuse disorder (n=29), including alcohol (n=11) and opioid misuse disorder (n=12), or comorbidity with both disorders (n=6), compared to controls (n=12). The results demonstrate that these substances affect both common and different biological pathways compared to controls. Alcohol misuse affects primarily protein translation, rRNA processing, and energy metabolism pathways, while opioid influences G-protein signaling, protein translation, rRNA processing, energy metabolism, and angiogenesis pathways. Alcohol and opioid misuse combined appears to have a synergistic effect on protein expression of the TCA cycle, respiratory transport, mitochondrial function, LGI-ADAM interactions, interleukin signaling, neuronal system changes, and actin polymerization. These findings offer new insights into how substance misuse leads to protein alterations and how combined substance misuse leads to different protein changes compared to single substance misuse.

## Introduction

Substance use disorders (SUD) are a major public health problem with an annual medical cost exceeding $13 billion in 2017^1^. In addition to the financial burden, the US drug overdose death rate has also tripled in the last two decades^2^. This has created an urgent need to understand the neurotoxic consequences of substance use to develop targeted treatments to minimize and/or reverse the damage induced by these drugs. While the mechanisms leading to substance use and addiction are multifactorial^3^ and likely involve environmental^3, 4^, genetic^5, 6^, and socioeconomic factors^7, 8^, examining how different substances lead to protein changes in the brain can provide some insight into the underlying biological mechanisms of neurotoxicity.

There are a variety of substances that can cause problems for individuals, the most common being alcohol, opioids, amphetamines, hallucinogens, cocaine, cannabis, and benzodiazepines. Each substance affects the brain differently, with some activating, others being sedatives, among other effects. How each substance affects the brain has been investigated using genomic^9–11^ and epigenetic^12–14^ approaches with both in vitro and in vivo models^15–17^, but studies examining the proteome at a systemic level are scarce. Further, few studies have explored the proteomics landscape in the human postmortem brain. Even fewer studies have examined how polysubstance use alters proteins in the brain despite the high prevalence of polysubstance use ^18, 19^. It has been reported that 5.6% of adults in the U.S. have used both alcohol and another illicit drug and 1.1% met diagnostic criteria for alcohol use disorder and another SUD^20^. Also, having one SUD is known to increase the susceptibility to a different SUD^19^. Alcohol use, in particular, leads to two fold increase in heroin dependence^19^. Despite these concerning statistics, how comorbid substance use affects the brain proteome is under-investigated. To help address this gap in understanding the protein changes after co-morbid alcohol and opioid use disorder, this study performed a comprehensive evaluation of protein alterations in postmortem brain from individuals with alcohol use disorder, opioid use disorder, combined alcohol and opioid use disorders, and non-substance users (controls). Understanding the protein changes occurring in these samples may shed light into how the brain adapts to chronic use of alcohol, opioids, or a combination of these substances.

## Methods

### Sample Collection

Sample collection was performed as described previously^21^. Briefly, with Institutional Review Board approval, postmortem brain tissue was obtained from the University of Texas Health Science Center at Houston Brain Collection in collaboration with the Harris County Institute of Forensic Science. Demographic data, autopsy and toxicology reports, and medical and psychiatric notes were obtained for each subject. The UT Health Psychological Autopsy Interview Schedule (UTH-PAIS) was used for each individual interviewing the next of kin^22^, where psychiatric clinical syndromes (depression/mania/psychosis), age of onset of drug use, types of drugs used, alcohol and smoking history, and other co-morbidities were obtained. A main diagnosis of opioid use disorder, alcohol use disorder (with no other substance use), comorbid alcohol and opioid use disorder or No Axis 1 diagnosis (non-psychiatric control) was reached by a panel of three trained clinicians after review of all information using DSM-5 criteria.

For this study, samples from 12 non-psychiatric controls, 11 alcohol use disorder (AUD), 12 opioid use disorder (OUD), and 6 combined alcohol and opioid use disorder (OUD + AUD) were used to analyze proteomics data. Postmortem interval (PMI) was calculated as the interval from time of death until tissue preservation. Within the dorsolateral prefrontal cortex, Brodmann Area 9 (BA9) was defined between the cingulate sulcus and the superior frontal gyrus. Dissections were obtained using a 4 mm punch through the cortex, yielding approximately 100 mg of tissue. Cerebellar pH was measured.

### Proteomics Analysis

Proteomics data was generated by nanoflow liquid chromatography-tandem mass spectrometry (NanoLC MS/MS), with sample processing, data acquisition, and peptide identification performed in the same manner as described previously^21^. The raw data for the mass spectrometry proteomics have been deposited to the ProteomeXchange Consortium via the PRIDE partner repository with the dataset identifier PXD025269.

### Statistical Analysis

Statistical analysis on the normalized and imputed proteomics data was performed using Jamovi software^23^. Non-parametric ANOVA analysis was performed comparing all SUD to controls and any protein that showed a statistical difference of p < 0.05 was selected. Post hoc analysis was performed using Dwass-Steel-Critchlow-Fligner pairwise comparisons comparing individual SUDs or combined SUDs to controls. Proteins showing p < 0.05 between individual substance use or co-morbid SUDs to controls were selected for linear regression analysis, which was performed on each protein including covariates gender, age bracket, PMI, ethnicity, and pH. Proteins showing a difference between individual substance use or co-morbid SUDs with a p < 0.05 after linear regression analysis and adjusting for covariates were included in the final pathway analysis.

### Pathway Analysis

Pathway analysis was performed using Reactome^24, 25^ with human species as criteria. The data was displayed as Reacfoam graphs showing enriched protein pathways. Subsequently, pathway analysis was performed using Cytoscape with String Nodes to generate pathway figures.

## Results

Postmortem brain samples from 41 individuals were used for analysis. The demographic data for the control and each of the SUD groups are summarized in Table 1. Detailed demographic information including age, gender, cause of death, drugs found in toxicology screen, as well as other diagnosis or comorbidities associated with the samples are included as Supplemental Table 1. In general, the samples were predominantly from white males, with an average age of 48 for controls and 45 for SUD individuals.

**Table 1.** Demographic data. In total, 41 post-mortem samples were collected, and the demographic data of the samples are listed. In general, the samples were predominately from white males in the age bracket of 30-49 years old.

| Diagnosis Group | Race |  |  |  |  |
| --- | --- | --- | --- | --- | --- |
|  | White | Black | Hispanic | Asian | white |
| control | 6 | 3 | 1 | 1 | 1 |
| alcohol use disorder | 9 | 1 | 1 | 0 | 0 |
| opioid use disorder | 8 | 4 | 0 | 0 | 0 |
| opioid + alcohol use disorder | 6 | 0 | 0 | 0 | 0 |

| Diagnosis Group | Gender |  |
| --- | --- | --- |
|  | Male | Female |
| control | 11 | 1 |
| alcohol use disorder | 10 | 1 |
| opioid use disorder | 7 | 5 |
| opioid + alcohol use disorder | 3 | 3 |

| Diagnosis Group | Age Bracket |  |  |  |
| --- | --- | --- | --- | --- |
|  | <30 | 30-49 | 50-64 | 65+ |
| control | 2 | 2 | 7 | 1 |
| alcohol use disorder | 0 | 5 | 4 | 2 |
| opioid use disorder | 3 | 7 | 2 | 0 |
| opioid + alcohol use disorder | 1 | 1 | 3 | 1 |

A total of 4,612 unique proteins were analyzed in all samples (Supplemental Table 2). As a first step, and to reduce computing power, non-parametric ANOVA analysis was performed comparing all individuals with SUD to controls. This analysis yielded 1,296 proteins with p < 0.05, (Supplemental Table 3) that were used for post-hoc analysis using the Dwass-Steel-Critchlow-Fligner method. Using this methodology, 18 proteins were found to be of significance for AUD (Table 2), 97 for OUD (Table 3), and 103 for OUD+AUD (Table 4). Covariance analysis was performed for each protein in each arm using linear regression including age bracket (less than 30, 30-49, 50-64, and 65+ years of age), ethnicity (Asian, White, Black, Hispanic), gender (male, female), PMI and pH (Supplemental Table 4). Covariates found to be statistically significant are summarized in Table 5. Proteins found to be of significance after linear regression analysis were used for pathway analysis.

**Table 2.** Significant Proteins in AUD. In this table are the proteins that were found to be statistically significant compared to controls in AUD samples after ANOVA, Dwass-Steel-Critchlow-Fligner, and linear regression analysis.

| Protein | Percent Change Compared to Control | Mean Protein Level AUD Only | Mean Protein Level of Control | P Value from Linear Regression |
| --- | --- | --- | --- | --- |
| GGPS1 | -3.39% | 22.8 | 23.6 | <0.01 |
| EIF3F | -2.07% | 23.7 | 24.2 | 0.001 |
| PDPK1 | -5.33% | 23.1 | 24.4 | 0.002 |
| RAB5B | 0.72% | 27.8 | 27.6 | 0.003 |
| SLC16A1 | 5.00% | 25.2 | 24 | 0.004 |
| RPS11 | -7.31% | 24.1 | 26 | 0.012 |
| RPS27 | 6.41% | 24.9 | 23.4 | 0.012 |
| UCHL5 | -3.32% | 23.3 | 24.1 | 0.014 |
| IDH3A | -0.64% | 31 | 31.2 | 0.021 |
| SORD | -1.47% | 26.8 | 27.2 | 0.021 |
| ADAM11 | 5.37% | 25.5 | 24.2 | 0.023 |
| RBP4 | -2.13% | 23 | 23.5 | 0.025 |
| SSBP1 | -0.35% | 28.2 | 28.3 | 0.033 |
| NDUFC2 | 4.85% | 28.1 | 26.8 | 0.036 |
| AUH | 0.70% | 28.8 | 28.6 | 0.036 |
| EEF1G | -0.67% | 29.6 | 29.8 | 0.042 |
| PPP1R1B | 1.53% | 26.6 | 26.2 | 0.042 |
| CCDC124 | -2.95% | 23 | 23.7 | 0.045 |

**Table 3.** Significant Proteins in OUD. In this table are the proteins that were found to be statistically significant compared to controls in OUD samples after ANOVA, Dwass-Steel-Critchlow-Fligner, and linear regression analysis.

| Protein | Percent change | Mean Protein Level- Control | Mean Protein Level OUD Only | P Value from Linear Regression |
| --- | --- | --- | --- | --- |
| NCS1 | -4.84% | 24.8 | 23.6 | < .001 |
| WARS | 2.11% | 28.4 | 29 | < .001 |
| HIST1H2BO | -7.45% | 25.5 | 23.6 | 0.002 |
| CPPED1 | -6.23% | 25.7 | 24.1 | 0.002 |
| PITRM1 | 4.89% | 26.6 | 27.9 | 0.002 |
| SIRPA | 1.23% | 32.5 | 32.9 | 0.002 |
| GNAI3 | -1.92% | 26 | 25.5 | 0.003 |
| EXOC8 | -3.63% | 24.8 | 23.9 | 0.003 |
| PGK1 | 0.88% | 33.9 | 34.2 | 0.003 |
| RAB5B | 0.72% | 27.6 | 27.8 | 0.003 |
| HNRNPK | -0.66% | 30.2 | 30 | 0.004 |
| BSG | 1.36% | 29.5 | 29.9 | 0.005 |
| GNB1 | 0.58% | 34.3 | 34.5 | 0.005 |
| SHMT2 | -5.95% | 25.2 | 23.7 | 0.006 |
| TCEB1 | -2.97% | 26.9 | 26.1 | 0.006 |
| HECTD4 | -3.25% | 24.6 | 23.8 | 0.006 |
| SH3BGRL2 | -9.29% | 26.9 | 24.4 | 0.008 |
| LRRC57 | 1.85% | 27.1 | 27.6 | 0.008 |
| RPL32 | 3.81% | 23.6 | 24.5 | 0.008 |
| ATP6V1G2 | 1.92% | 31.3 | 31.9 | 0.009 |
| EIF1 | -2.48% | 24.2 | 23.6 | 0.01 |
| NLGN2 | 2.42% | 24.8 | 25.4 | 0.01 |
| ATP2B4 | 0.63% | 31.6 | 31.8 | 0.011 |
| DPYSL3 | 0.96% | 31.4 | 31.7 | 0.012 |
| CCBL2 | -4.03% | 27.3 | 26.2 | 0.013 |
| VPS13C | -7.39% | 25.7 | 23.8 | 0.013 |
| EML1 | 2.59% | 23.2 | 23.8 | 0.013 |
| PRPSAP1 | 3.59% | 25.1 | 26 | 0.013 |
| NEGR1 | 1.01% | 29.6 | 29.9 | 0.013 |
| HSPA12A | -1.66% | 30.1 | 29.6 | 0.014 |
| ACTR3B | 1.41% | 28.4 | 28.8 | 0.014 |
| UBE2V2 | 1.74% | 28.8 | 29.3 | 0.015 |
| ICAM5 | 1.97% | 30.5 | 31.1 | 0.015 |
| RPS29 | 4.27% | 23.4 | 24.4 | 0.016 |
| NRCAM | 0.95% | 31.6 | 31.9 | 0.016 |
| SYNE1 | -3.67% | 24.5 | 23.6 | 0.017 |
| GNAO1 | 0.29% | 34.5 | 34.6 | 0.017 |
| RDH11 | -3.98% | 25.1 | 24.1 | 0.019 |
| PPP1R12B | 3.45% | 23.2 | 24 | 0.019 |
| SLC25A22 | 0.97% | 30.8 | 31.1 | 0.019 |
| PDHA1 | -0.96% | 31.2 | 30.9 | 0.02 |
| OAT | -2.57% | 27.2 | 26.5 | 0.021 |
| EIF2S3 | 0.75% | 26.8 | 27 | 0.021 |
| LAMTOR3 | 8.55% | 23.4 | 25.4 | 0.021 |
| IDH3A | -0.96% | 31.2 | 30.9 | 0.022 |
| SSBP1 | -0.71% | 28.3 | 28.1 | 0.023 |
| ACADS | 3.66% | 24.6 | 25.5 | 0.023 |
| EIF6 | 1.98% | 25.3 | 25.8 | 0.023 |
| STX12 | 1.81% | 27.7 | 28.2 | 0.023 |
| MT3 | 5.36% | 28 | 29.5 | 0.024 |
| FAM49B | 0.68% | 29.2 | 29.4 | 0.024 |
| CADM1 | 0.68% | 29.5 | 29.7 | 0.024 |
| TUFM | -2.64% | 30.3 | 29.5 | 0.025 |
| EIF4A2 | -0.67% | 29.7 | 29.5 | 0.026 |
| DENR | 1.88% | 26.6 | 27.1 | 0.026 |
| USP5 | -1.74% | 28.7 | 28.2 | 0.027 |
| ACAT2 | 1.02% | 29.3 | 29.6 | 0.027 |
| CDV3 | 1.84% | 27.2 | 27.7 | 0.027 |
| PCBP2 | -0.71% | 28 | 27.8 | 0.028 |
| RAB1B | 2.07% | 29 | 29.6 | 0.028 |
| COA6 | 1.12% | 26.7 | 27 | 0.028 |
| NSF | -1.25% | 32.1 | 31.7 | 0.029 |
| TMX4 | -5.53% | 25.3 | 23.9 | 0.029 |
| CRMP1 | 0.61% | 32.9 | 33.1 | 0.029 |
| ATP2B1 | 0.32% | 31.1 | 31.2 | 0.031 |
| VCP | 0.95% | 31.7 | 32 | 0.031 |
| RPS12 | 2.93% | 27.3 | 28.1 | 0.032 |
| RASAL1 | 5.67% | 24.7 | 26.1 | 0.033 |
| NLGN3 | 3.40% | 23.5 | 24.3 | 0.033 |
| CCDC132 | -3.64% | 24.7 | 23.8 | 0.034 |
| G3BP2 | -3.35% | 26.9 | 26 | 0.034 |
| RPL7 | 0.69% | 29 | 29.2 | 0.034 |
| PRDX6 | -1.90% | 31.5 | 30.9 | 0.035 |
| NPTXR | 3.56% | 25.3 | 26.2 | 0.035 |
| SH3GLB2 | 1.00% | 30 | 30.3 | 0.036 |
| DNM2 | -8.68% | 26.5 | 24.2 | 0.037 |
| WDR44 | 2.69% | 26 | 26.7 | 0.037 |
| MAL2 | 7.36% | 25.8 | 27.7 | 0.037 |
| PACSIN1 | 0.61% | 32.8 | 33 | 0.037 |
| PPP2R1A | -0.95% | 31.6 | 31.3 | 0.038 |
| AARS | -1.35% | 29.7 | 29.3 | 0.038 |
| VAT1 | -1.39% | 28.7 | 28.3 | 0.039 |
| PFKP | -1.63% | 30.7 | 30.2 | 0.04 |
| SYNPR | 2.53% | 27.7 | 28.4 | 0.04 |
| TBC1D24 | 1.07% | 28 | 28.3 | 0.04 |
| GLTP | -4.69% | 25.6 | 24.4 | 0.041 |
| TNR | 0.61% | 32.6 | 32.8 | 0.041 |
| PPP2R5D | -4.91% | 26.5 | 25.2 | 0.042 |
| ATP2A2 | 0.64% | 31.4 | 31.6 | 0.042 |
| SET | 2.15% | 23.3 | 23.8 | 0.043 |
| PDXP | 1.02% | 29.4 | 29.7 | 0.043 |
| UQCC2 | 2.58% | 23.3 | 23.9 | 0.043 |
| CPNE5 | 1.42% | 28.2 | 28.6 | 0.043 |
| CDH13 | 1.42% | 28.1 | 28.5 | 0.044 |
| PPP5C | 0.71% | 28.1 | 28.3 | 0.046 |
| SORD | -1.10% | 27.2 | 26.9 | 0.047 |
| CDS2 | -1.81% | 27.7 | 27.2 | 0.048 |

**Table 4.**
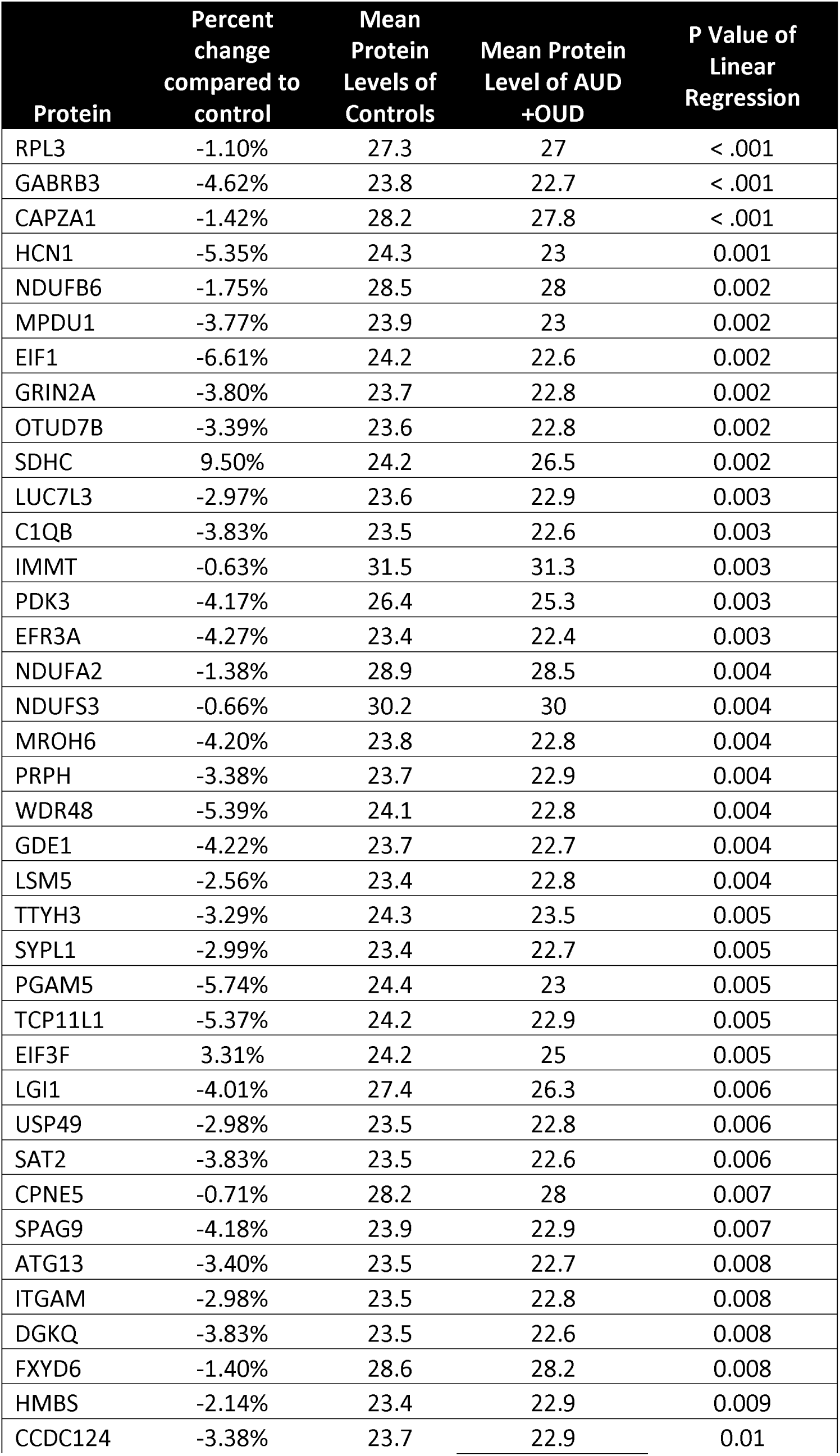
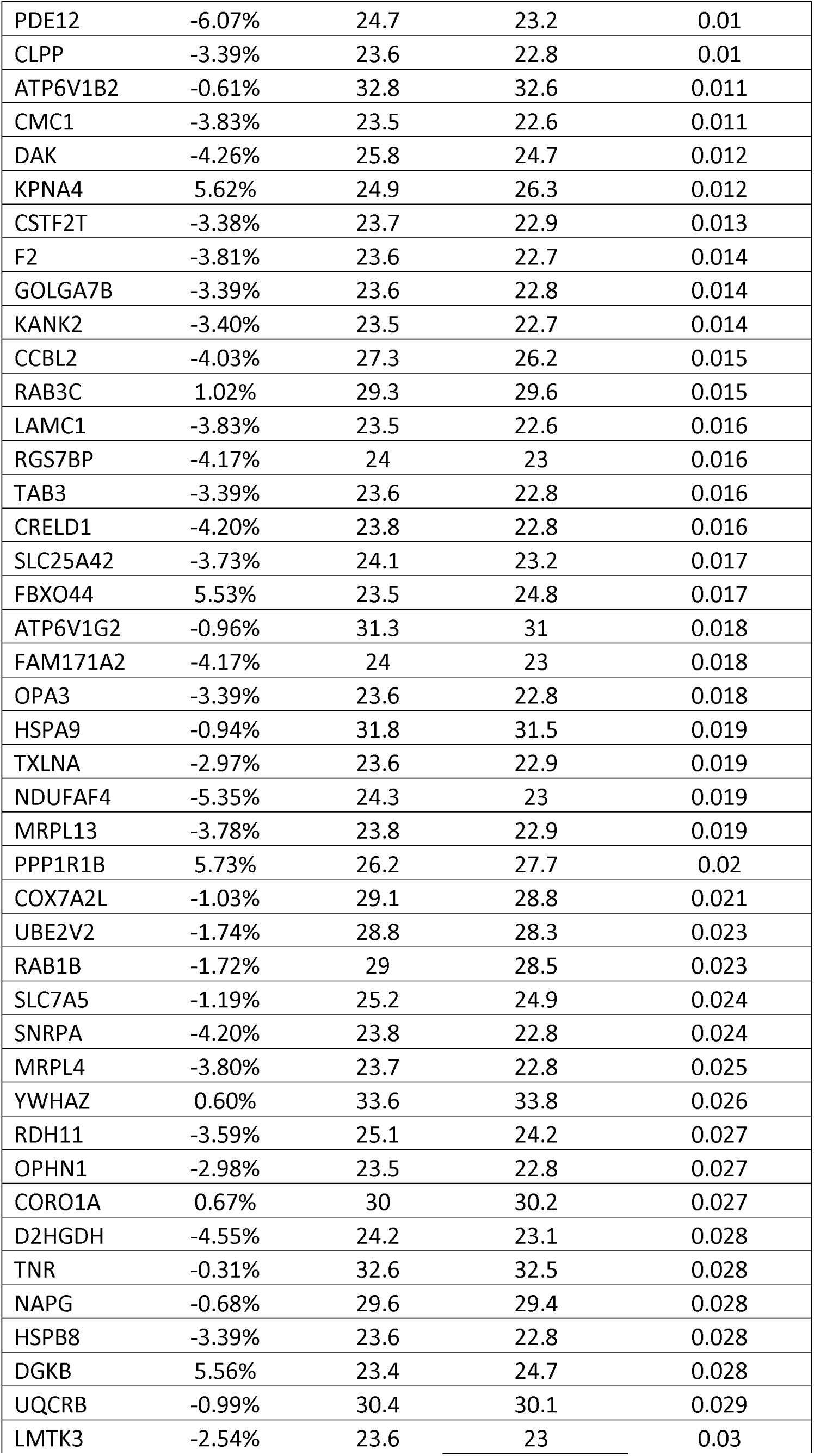
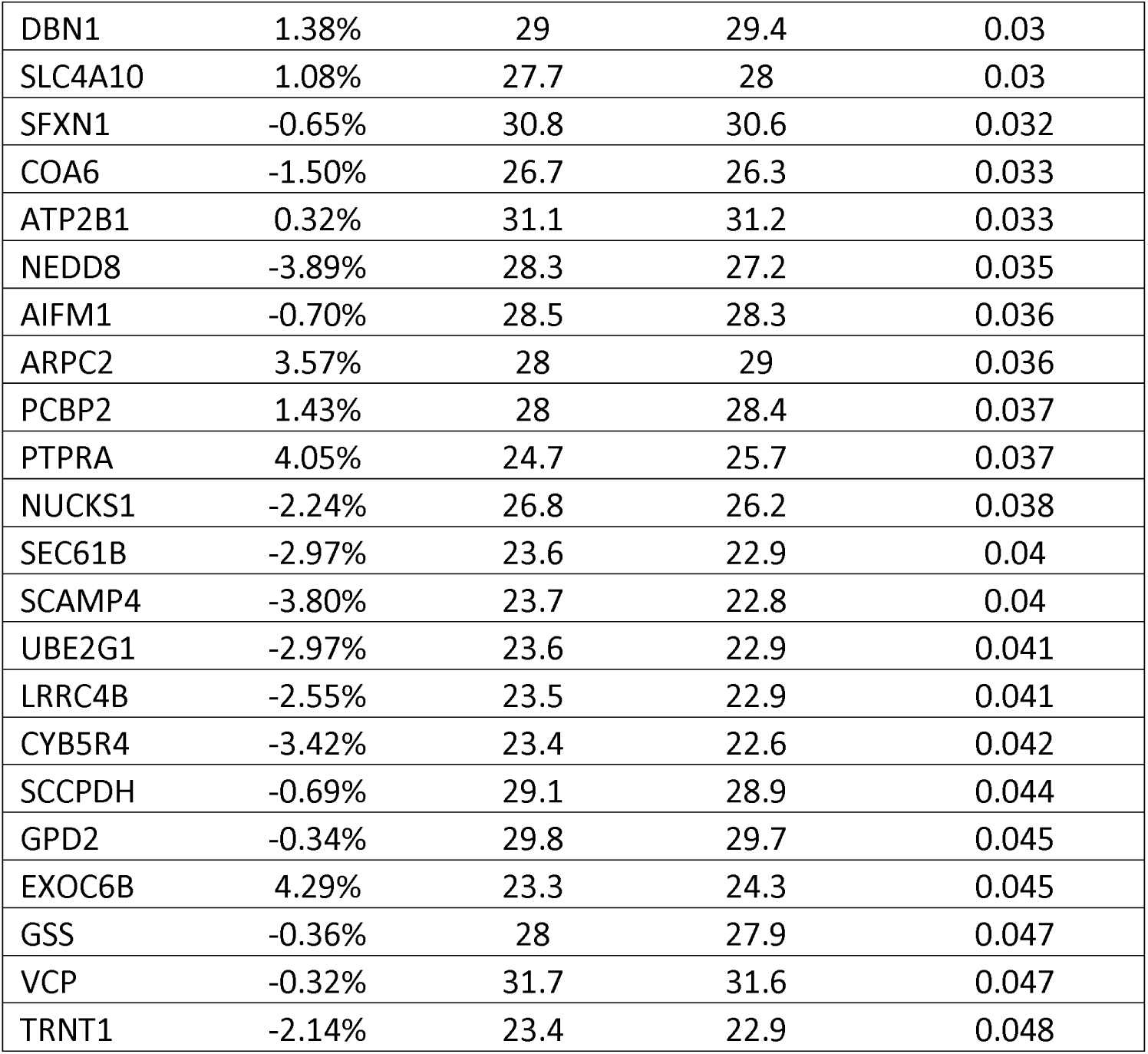
Significant Proteins in AUD + OUD. In this table are the proteins that were found to be statistically significant compared to controls in AUD + OUD samples after ANOVA, Dwass-Steel-Critchlow-Fligner, and linear regression analysis.

**Table 5.** Significant Covariates. The covariates included demographic data such as age bracket (less than 30, 30-49, 50-64, and 65+ years of age), ethnicity (Asian, White, Black, Hispanic), and gender (male, female) as well as factors for post-mortem sample collection such as post-mortem interval (PMI) and pH. The covariates that were statistically significant in the linear regression analysis are listed for each protein above in the table. For further information on non-significant covariates, please see Supplemental Table 4.

| Covariate analysis |  |  |
| --- | --- | --- |
| Alcohol Use Disorder Alone | Opioid Use Disorder Alone | Alcohol and Opioid Use Disorder Combined |
| Age (years) | Age (years) | Age (years) |
| <u><30</u><br>EIF3F | <u><30</u><br>EIF3F<br>DNM2<br>PRPSAP1<br>CCDC132<br>STX12<br>SLC25A22<br>RPL32 | <u><30</u><br>PCBP2<br>EIF3F<br>CAPZA1 |
| <u>50-64</u><br>EIF3F<br>PPP1R1B | <u>50-64</u><br>SORD<br>EIF3F<br>PPP1R1B<br>USP5<br>TUFM<br>PFKP<br>SIRPA<br>UBE2V2<br>SLC25A22<br>NLGN3 | <u>50-64</u><br>PCBP2<br>ATP2B1<br>GPD2<br>UBE2V2<br>TTYH3<br>EIF3F<br>MPDU1<br>PPP1R1B |
| <u>65+</u><br>EIF3F | <u>65+</u><br>EIF3F<br>AARS<br>GNB1<br>EIF4A2<br>SLC25A22 | <u>65+</u><br>EIF3F<br>KANK2 |
| PMI | PMI | PMI |
| PDPK1<br>EIF3F | PDPK1<br>EIF3F<br>DNM2<br>DPYSL3<br>PRPSAP1<br>CCBL2<br>CDH13<br>TCEB1<br>UBE2V2 | ATP6V1G2<br>CCBL2<br>UBE2V2<br>TTYH3<br>EIF3F<br>NAPG |

|  | NEGR1<br>STX12<br>ACAT2<br>NLGN3<br>TMX4<br>CADM1 |  |
| --- | --- | --- |
| pH | pH | pH |
| RPS11 | RPS11<br>CDH13<br>EIF1<br>SYNE1 | KPNA4<br>EIF1 |
| Ethnicity | Ethnicity | Ethnicity |
| <u>**Asian**</u><br>NDUFC2 | <u>**Asian**</u><br>NDUFC2<br>EIF4A2<br>UBE2V2<br>LRRC57 | <u>**Asian**</u><br>HMBS<br>SLC7A5<br>UBE2V2<br>SFXN1<br>NDUFB6<br>SFXN1<br>HCN1 |
| <u>**Black**</u> | <u>**Black**</u><br>SYNE1 | <u>**Black**</u><br>IMMT<br>FBXO44 |
| <u>**Hispanic**</u> | <u>**Hispanic**</u><br>PRPSAP1<br>WDR44<br>SET<br>NCS1<br>TCEB1<br>VAT1<br>HECTD4 | <u>**Hispanic**</u><br>GPD2<br>RPL3<br>SCCPDH<br>NDUFAF4<br>TRNT1 |
| Gender | Gender | Gender |
| <u>**Male**</u> | <u>**Male**</u><br>EXOC8<br>HECTD4 | <u>**Male**</u><br>PCBP2<br>RDH11<br>RPL3 |

The Reactome database was used to visualize protein pathways that were altered in each of the SUD compared to controls (Supplemental Figures 1-3). To visualize the significant pathways more clearly, barplots of each pathway that showed enrichment for each SUD were generated (Figures 1-3). Tables were also generated of the pathways that were enriched (Tables 5-7). In general, AUD showed changes in protein translation, translational silencing of ceruloplasmin expression, rRNA processing, nonsense mediated decay, and selenoamino acid metabolism (Figure 1 and Table 6). OUD showed major alterations in PP2A mediated dephosphorylation, platelet homeostasis and platelet sensitization, calcium levels regulation, rRNA processing, non-sense mediated decay, translation of proteins, glucose metabolism, and G protein activation (Figure 2 and Table 7). In OUD+AUD, the TCA cycle and respiratory electron transport synthesis were mainly affected (Figure 3 and Table 8).

**Figure 1.**
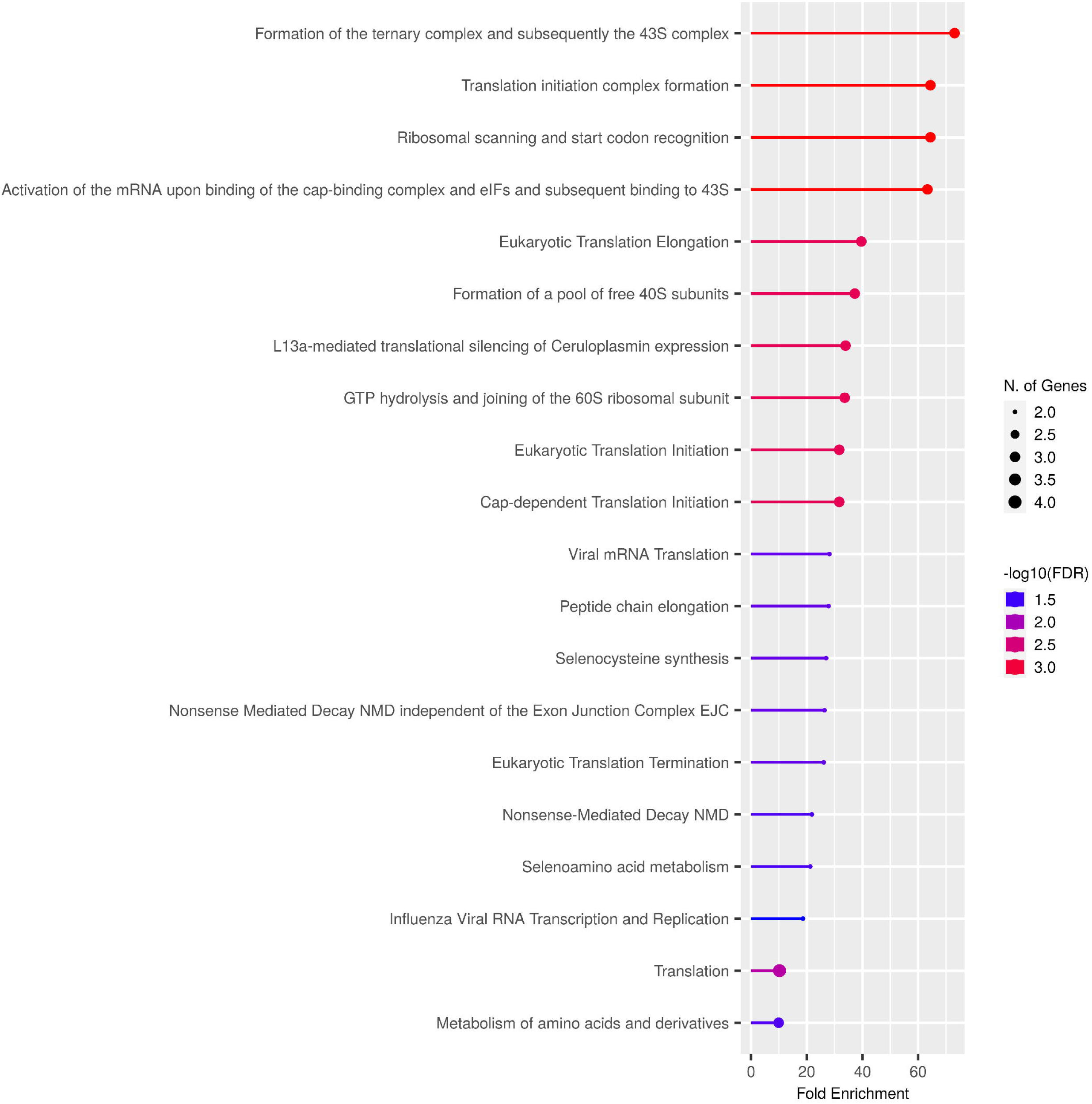
Bar plot of AUD. The bar plot demonstrates that there is enrichment in pathways for protein translation, selenocysteine synthesis, and nonsense mediated decay in addition to other pathways in AUD.

**Figure 2.**
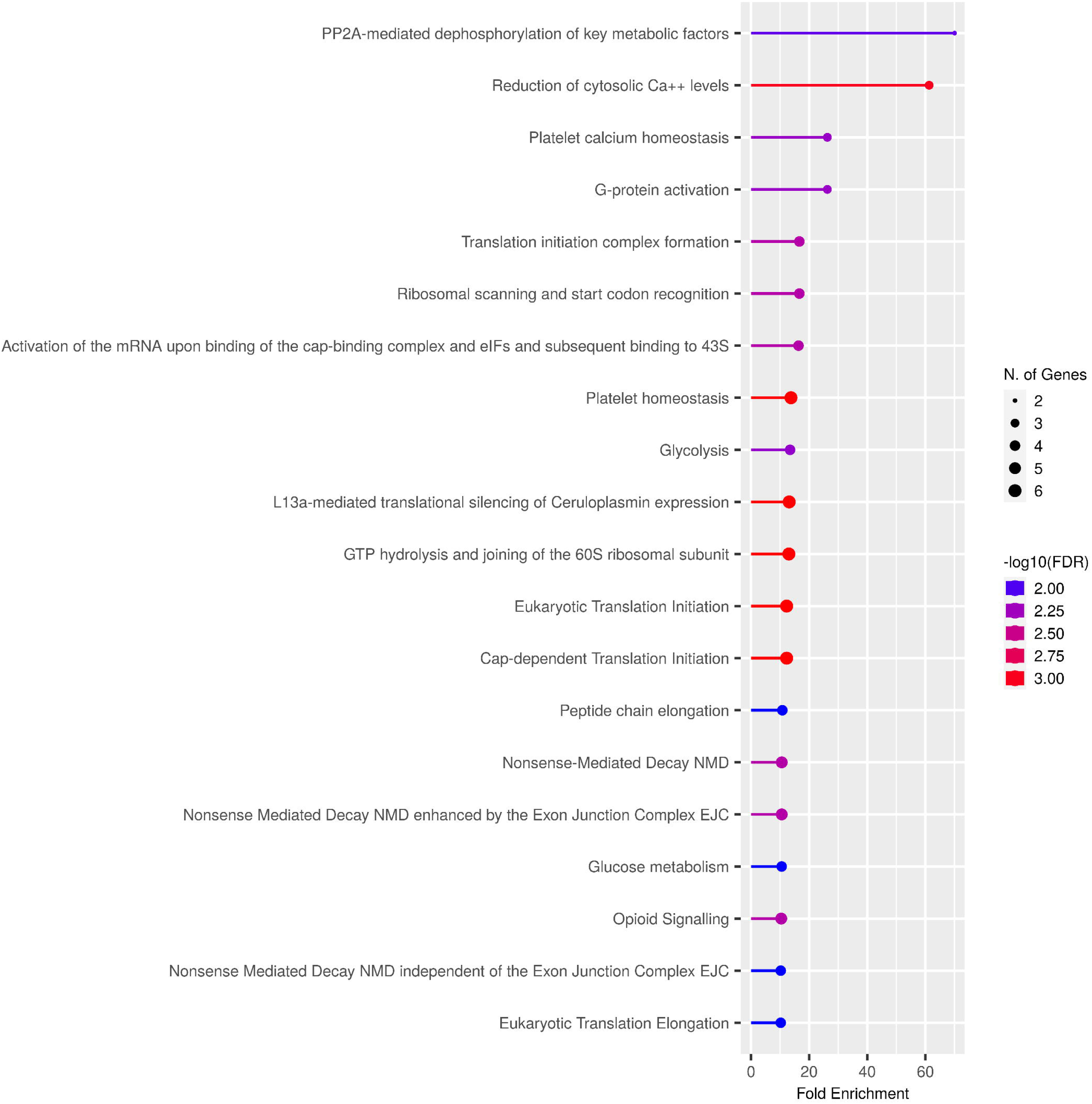
Bar plot of OUD. The bar plot demonstrates that there is enrichment in pathways for calcium levels, PP2A mediated dephosphorylation, platelet calcium homeostasis, and G protein activation as well as others in OUD.

**Figure 3.**
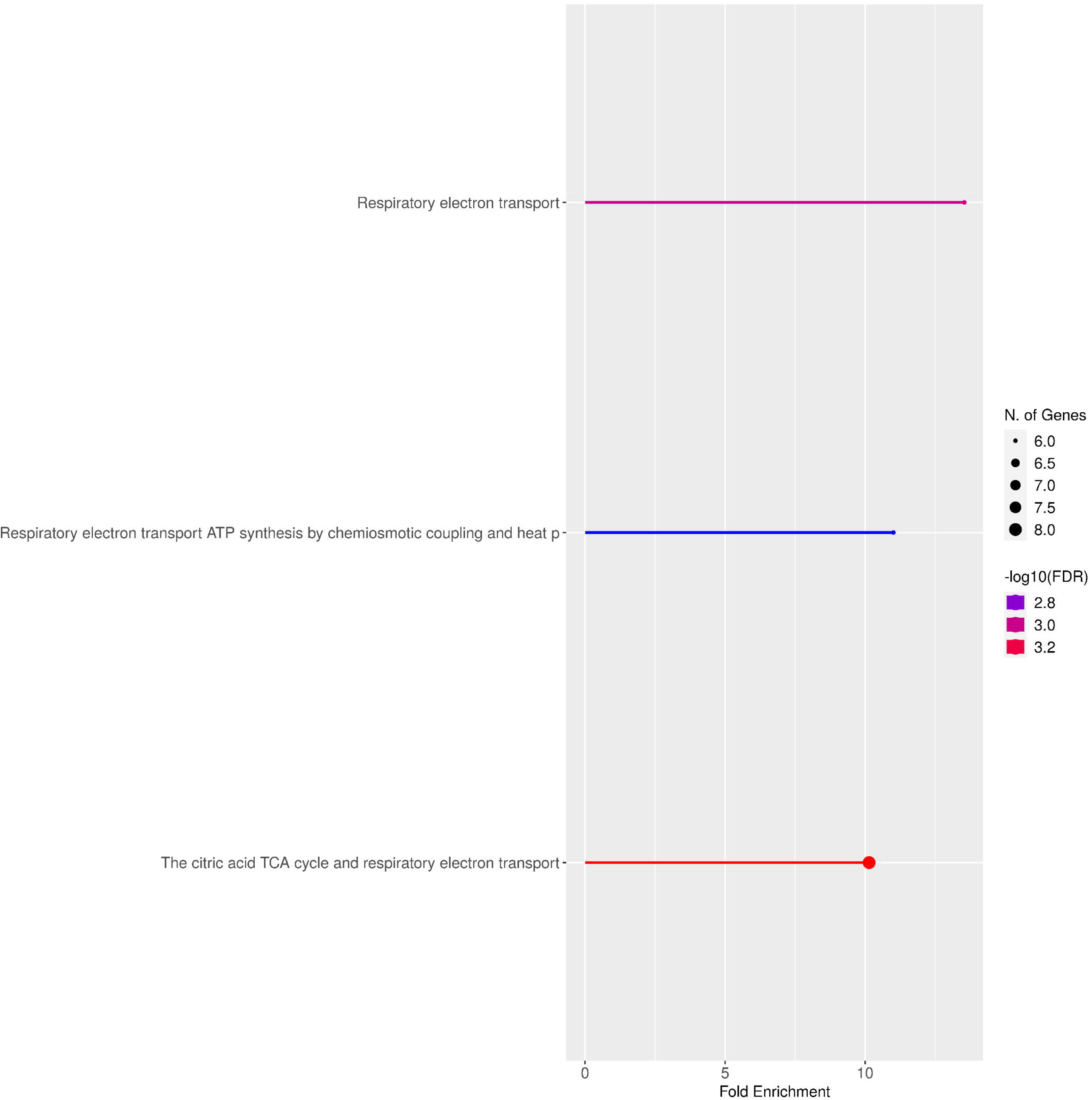
Bar plot of AUD + OUD. The bar plot demonstrates that the respiratory electron transport and TCA cycle pathways are enriched in AUD + OUD.

**Table 6.**
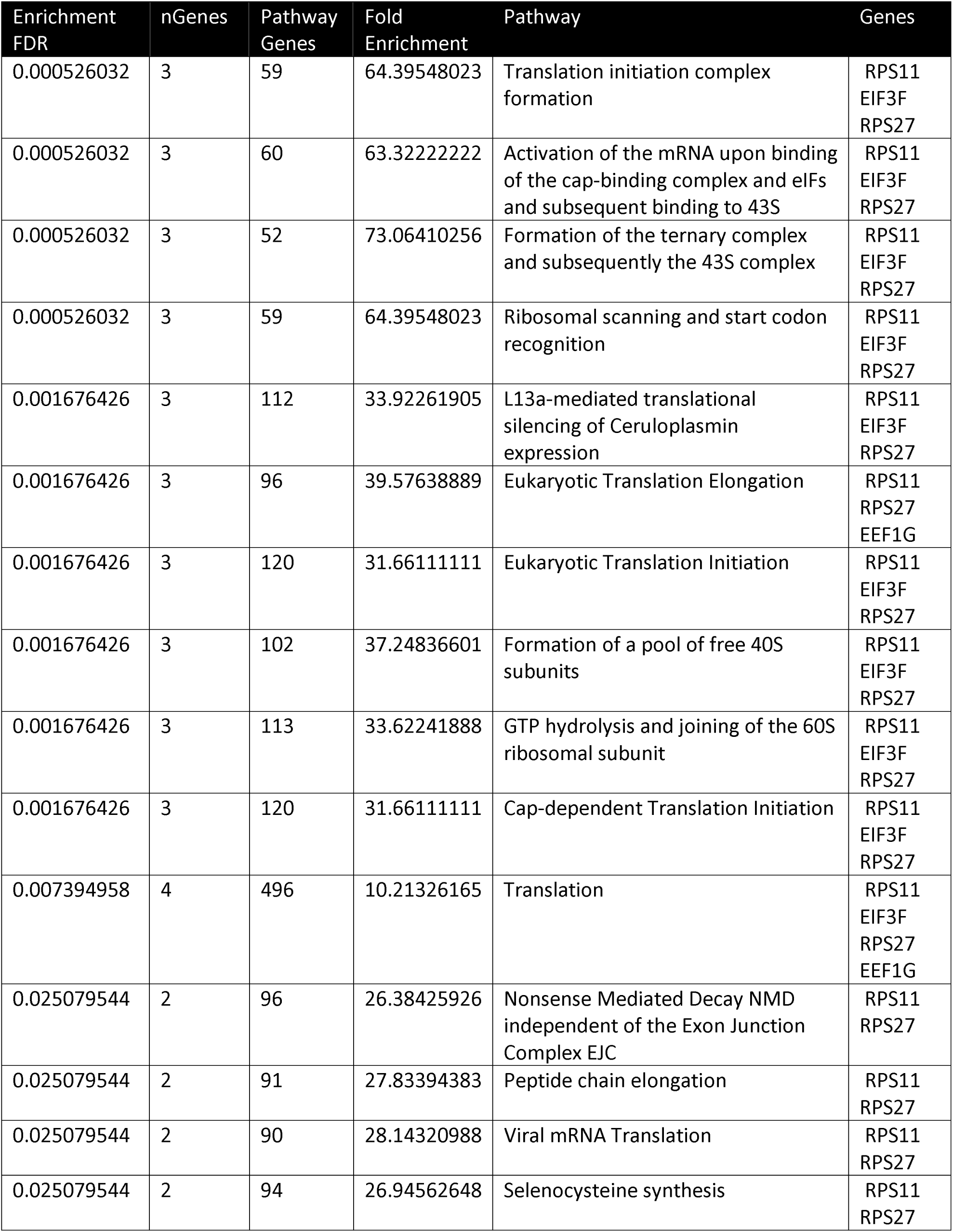
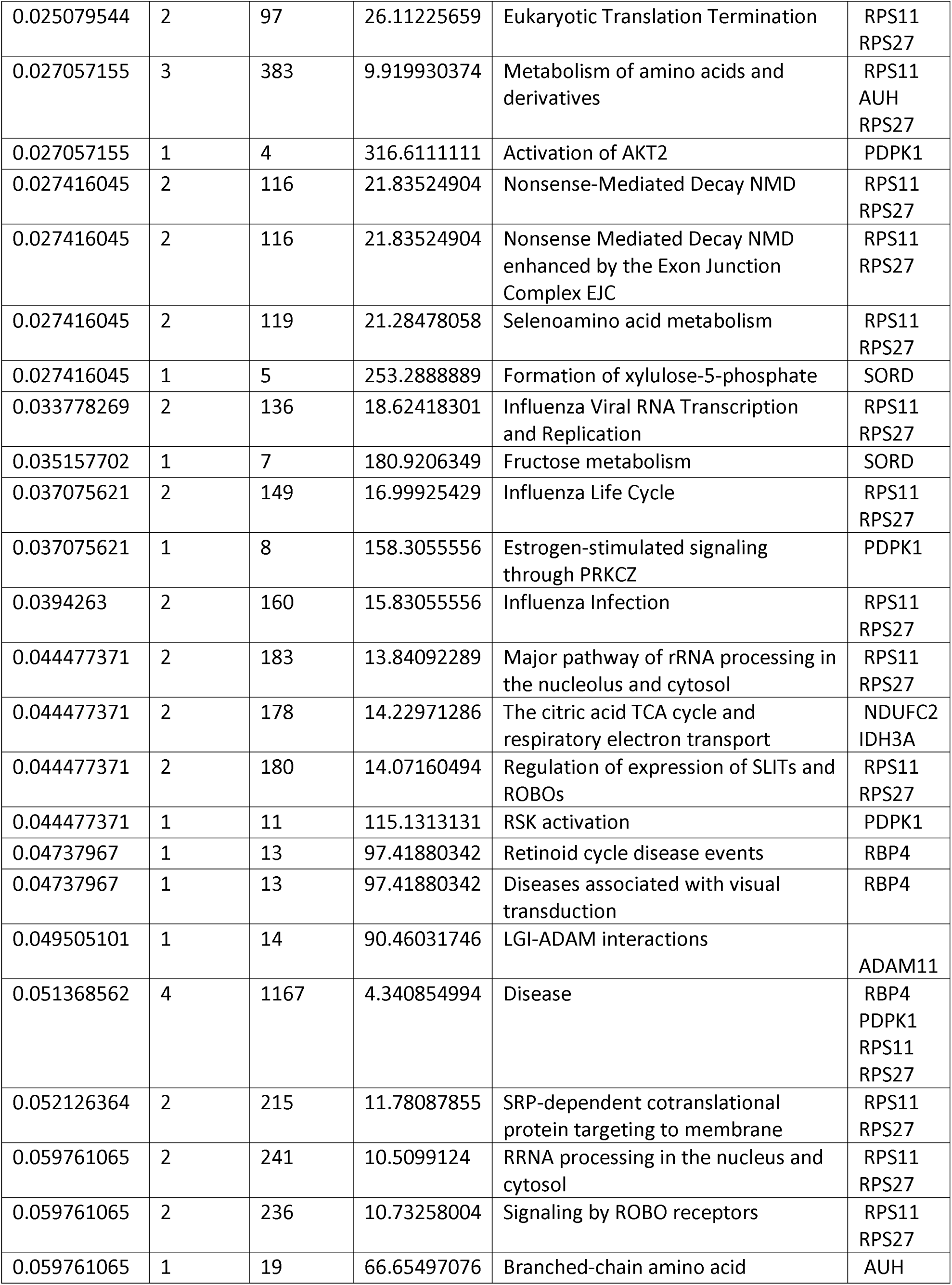
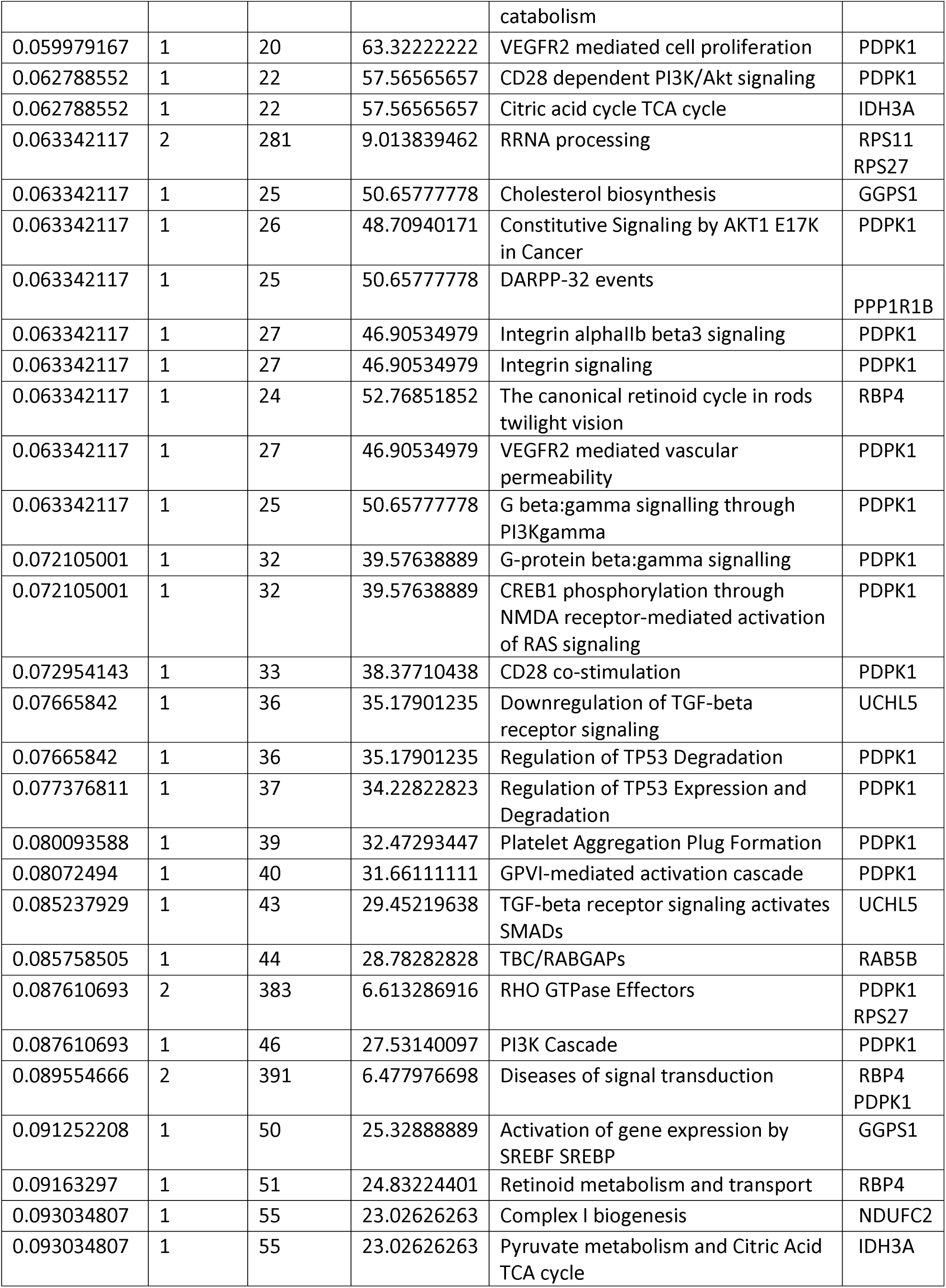
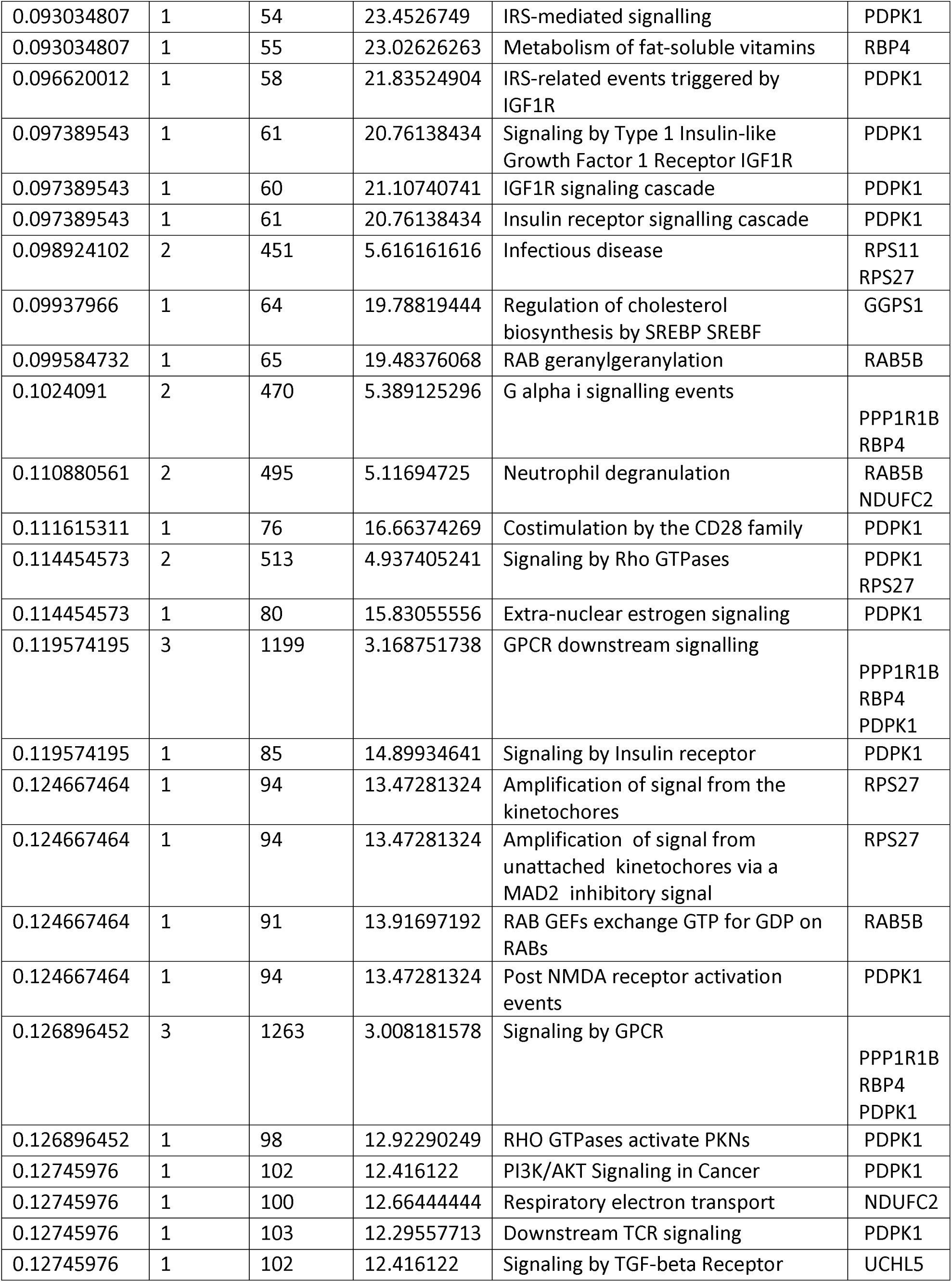
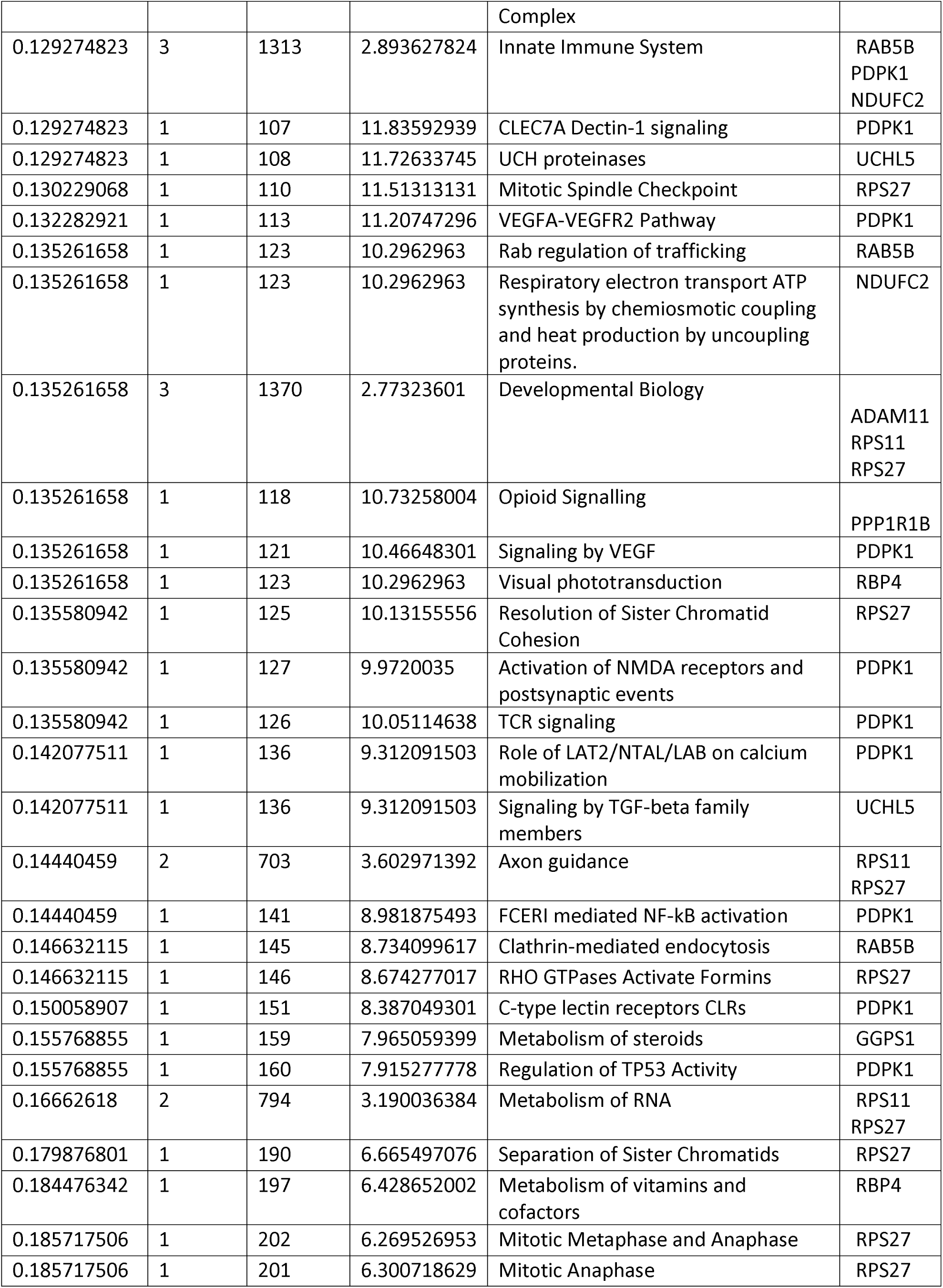
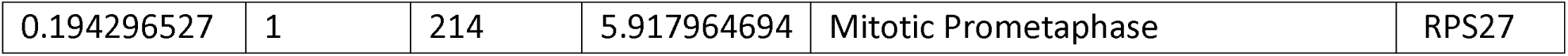
Enriched Pathways in AUD. The table shows the enrichment FDR, the number of proteins/genes enriched, the total number of genes in the pathway, the fold enrichment, the pathway that is enriched, and the genes/proteins that were found to be enriched in the pathway for AUD samples. A simplified view of the data via a bar plot is included in Figure 1.

**Table 7.**
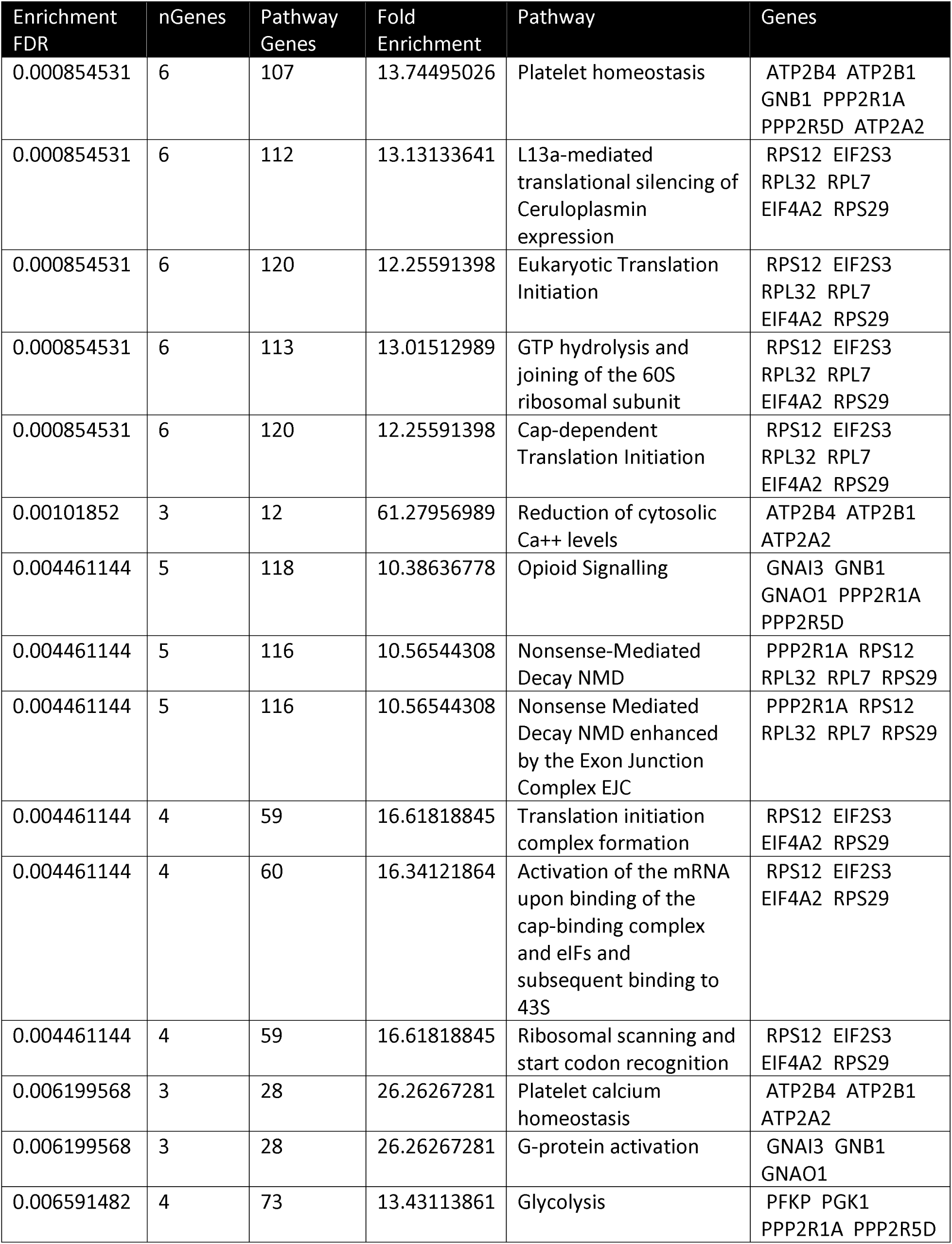
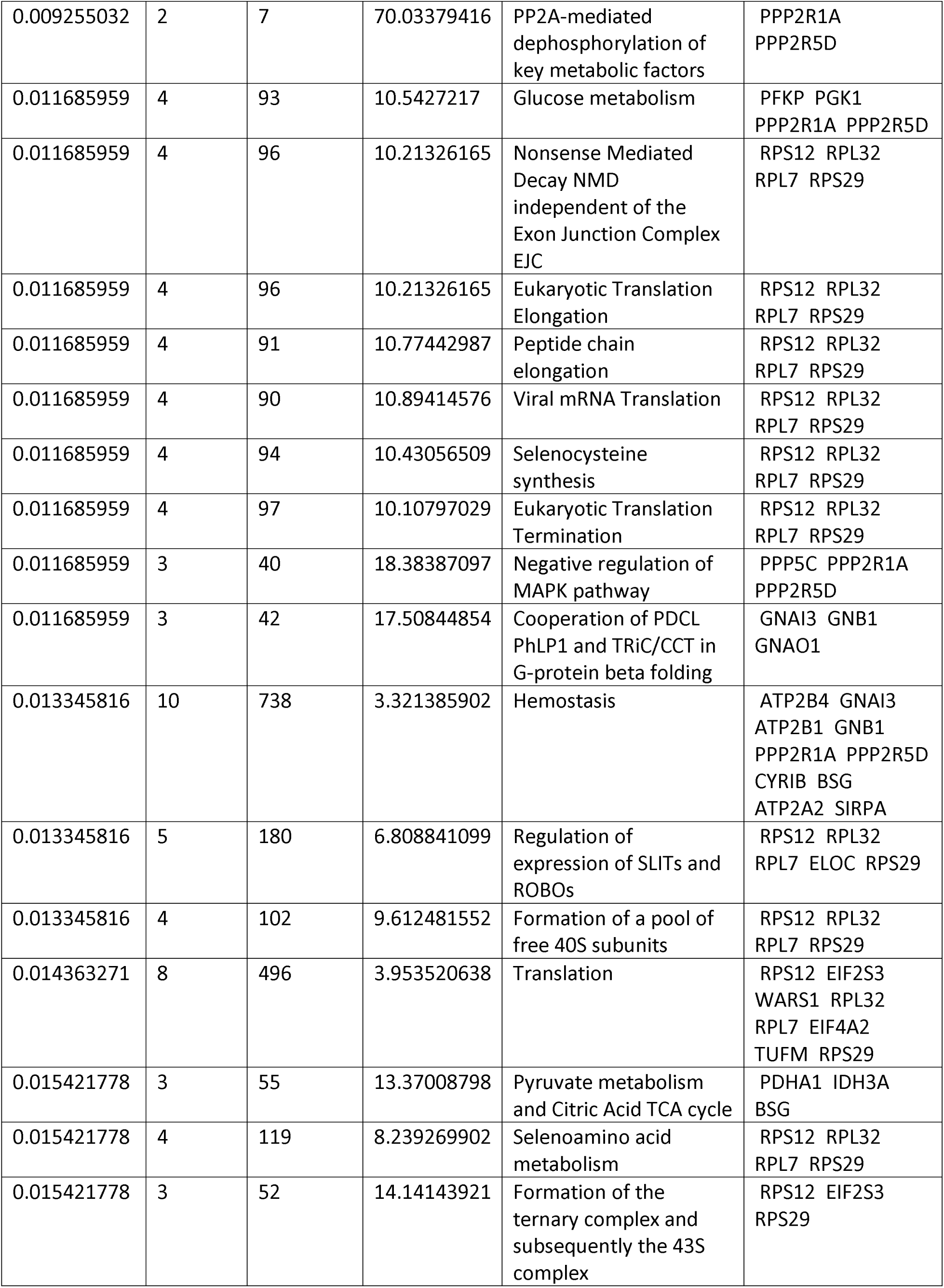
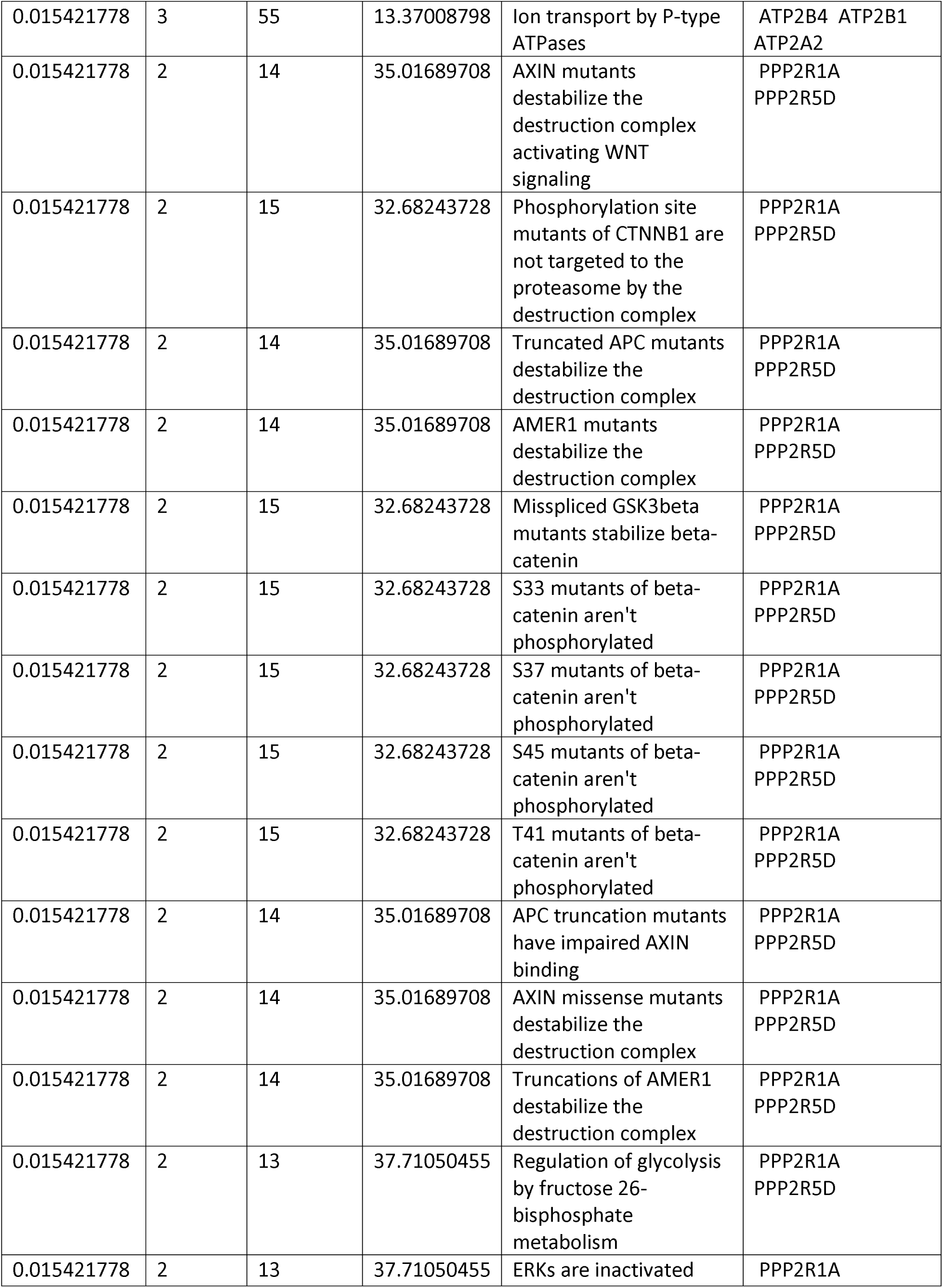
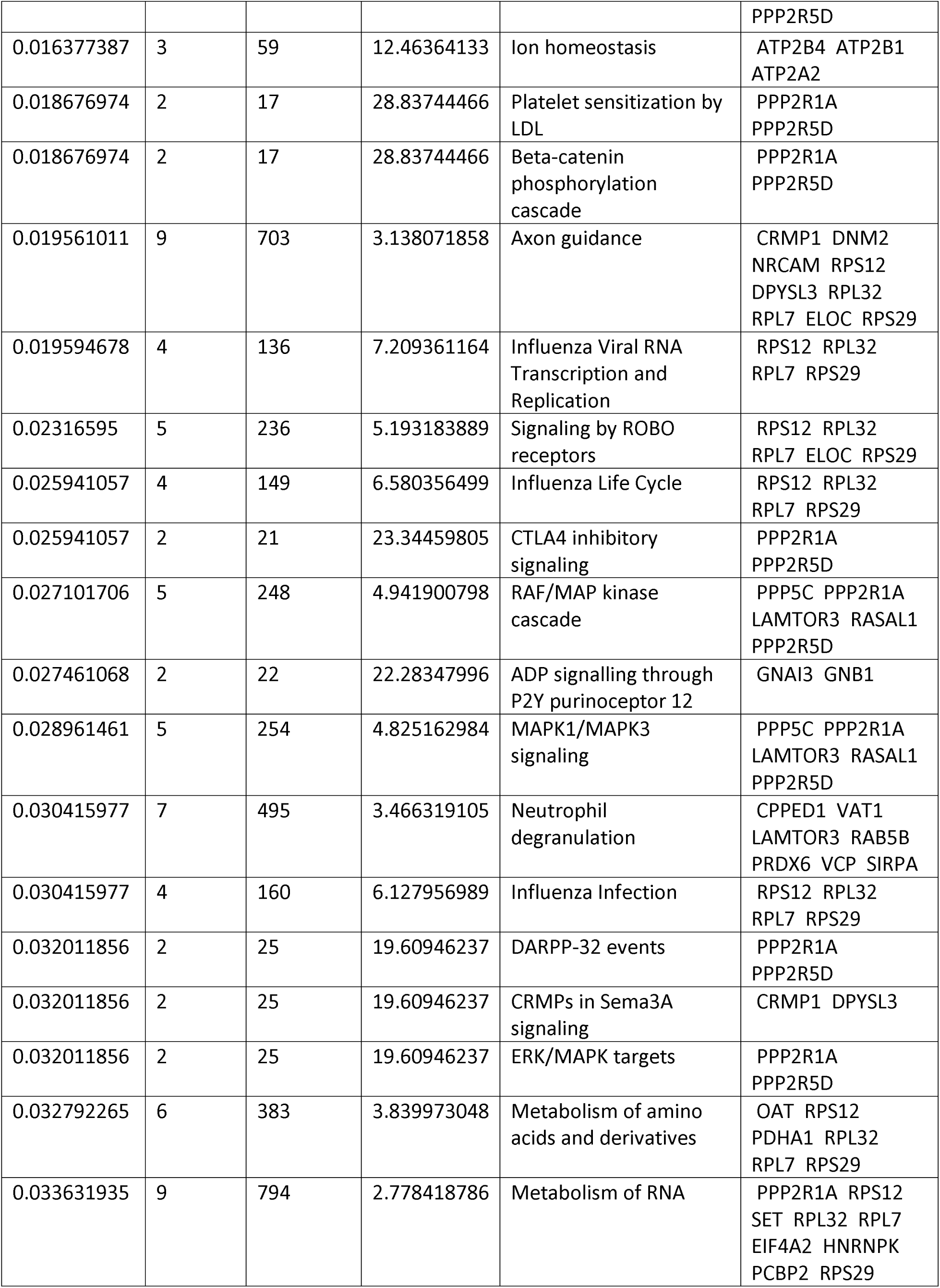
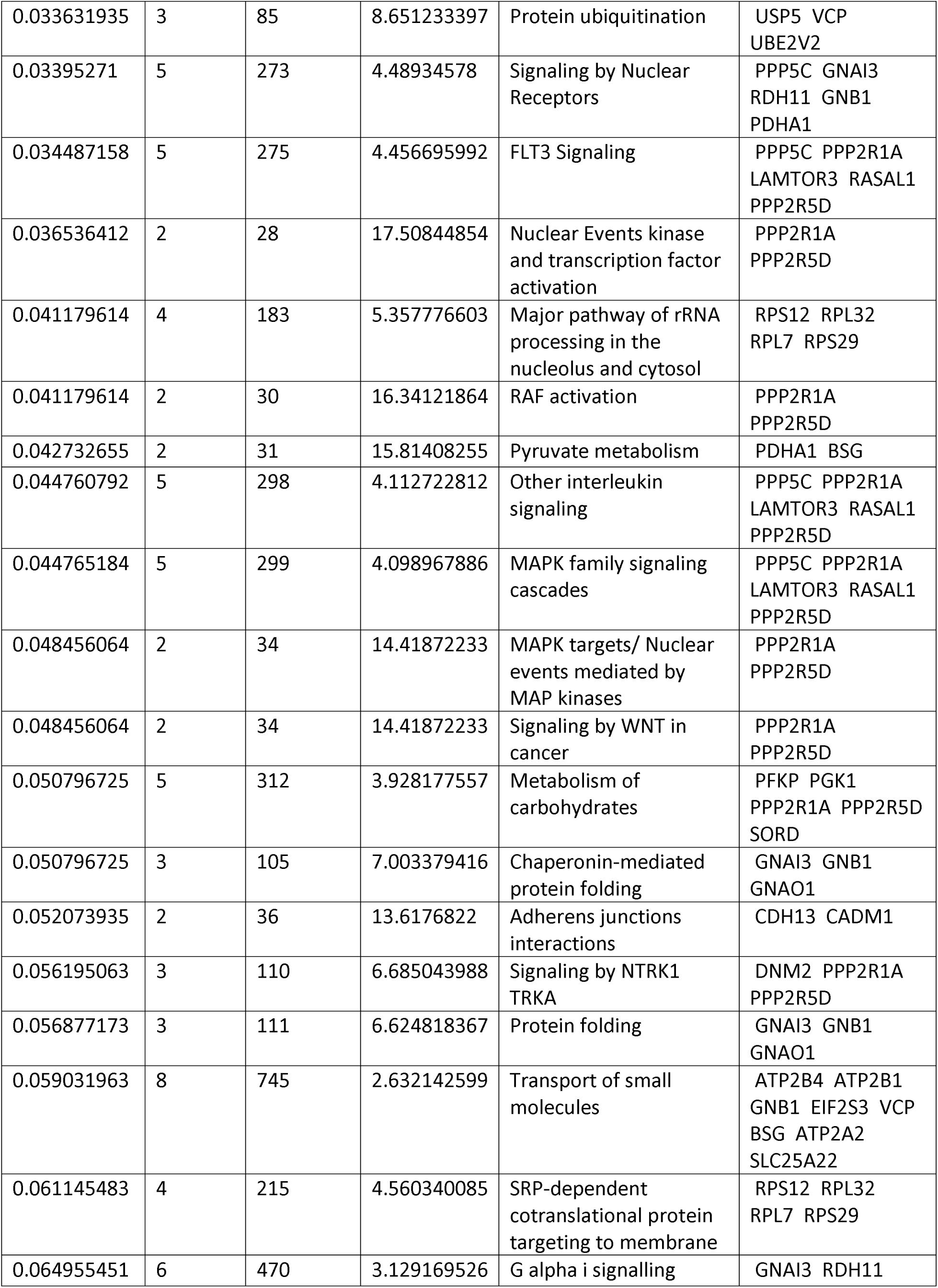
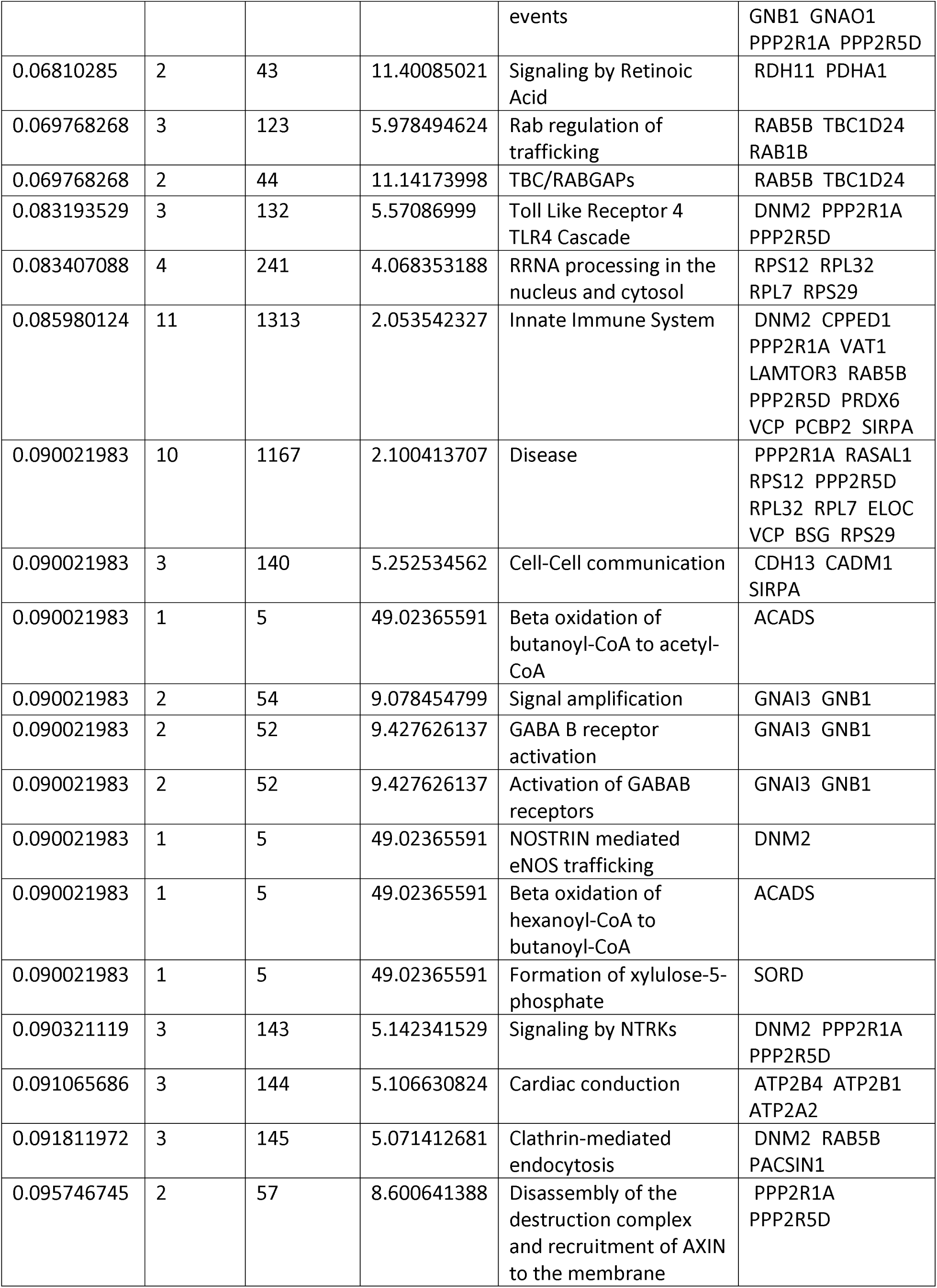
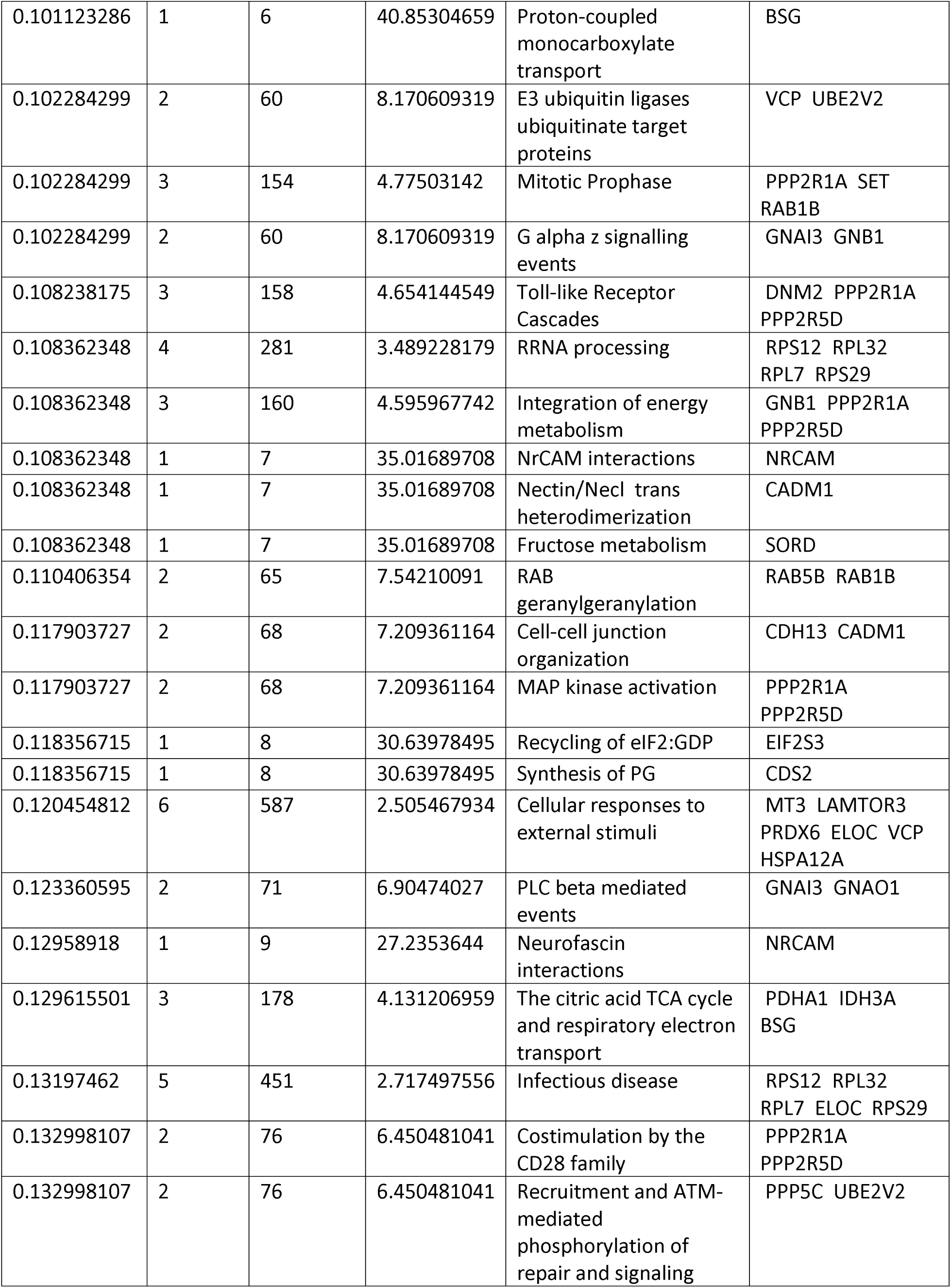
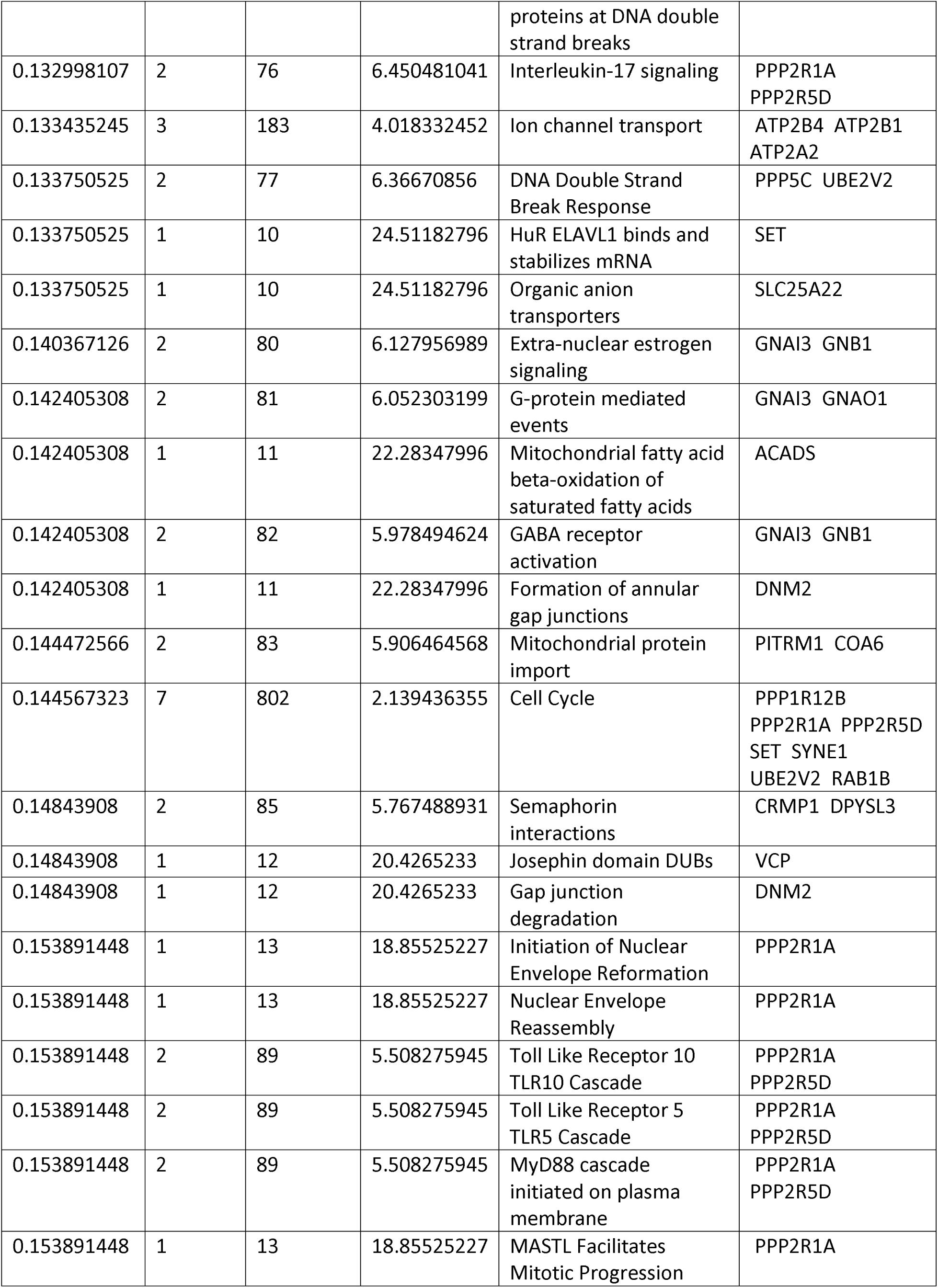
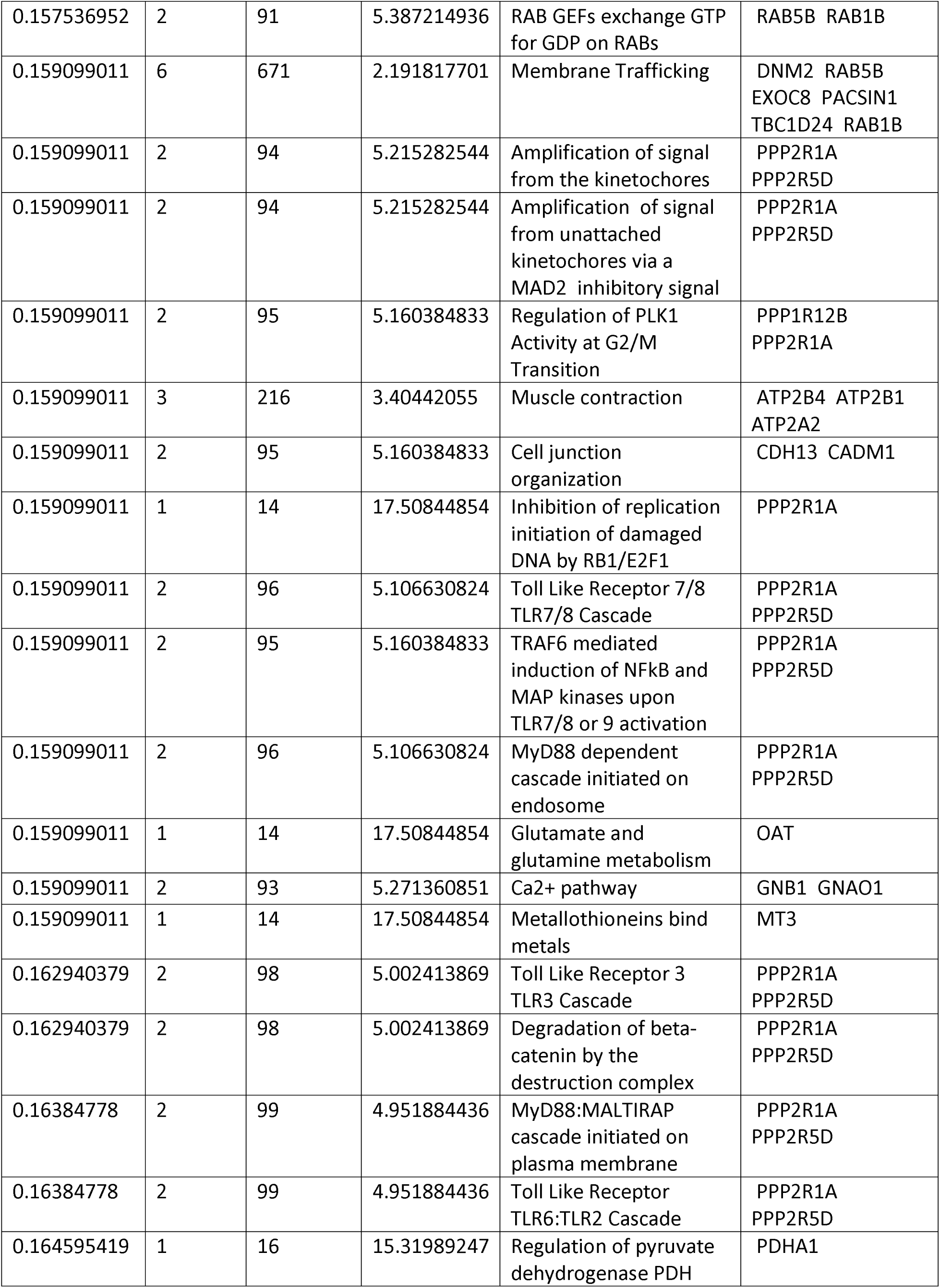
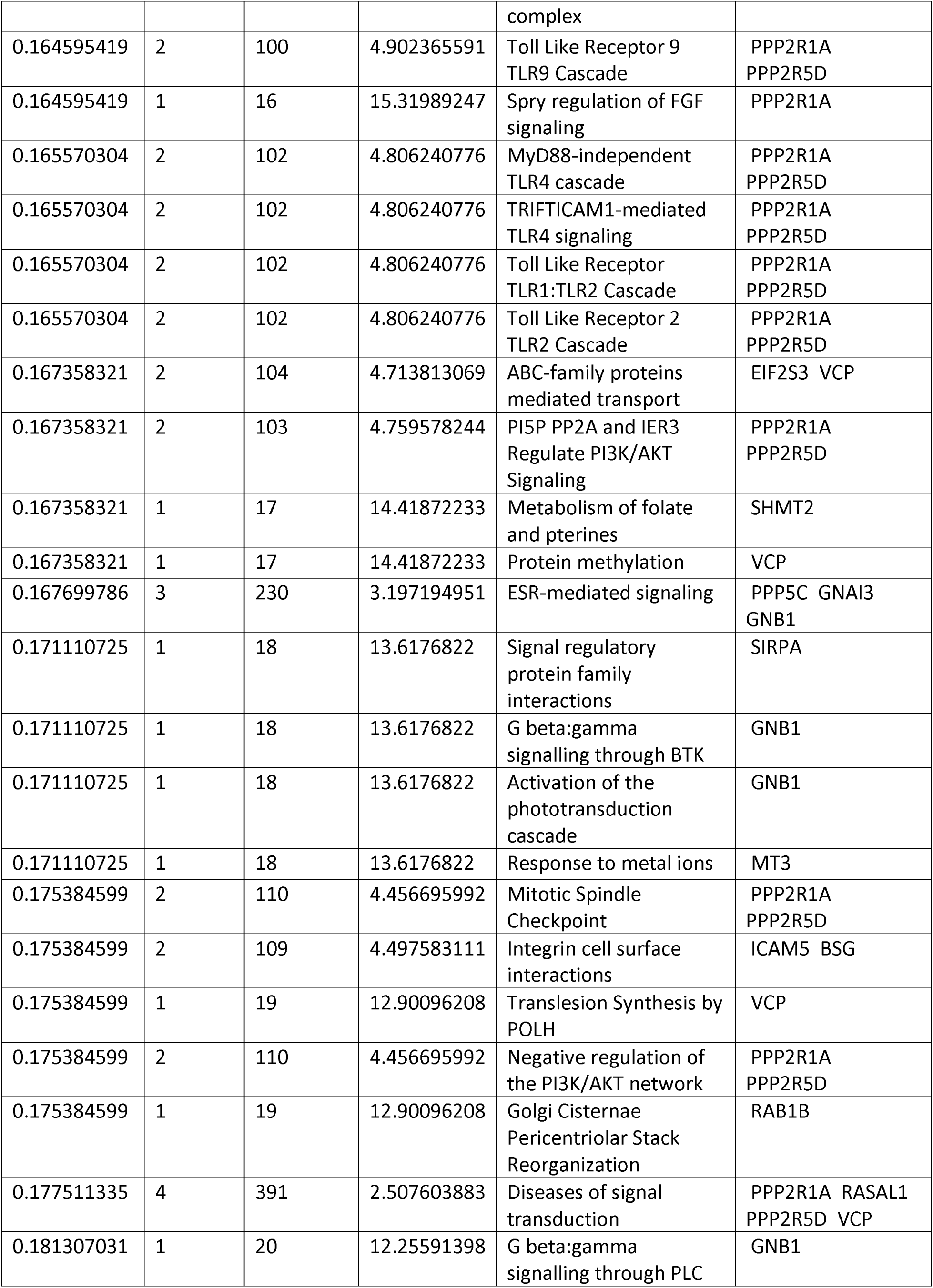
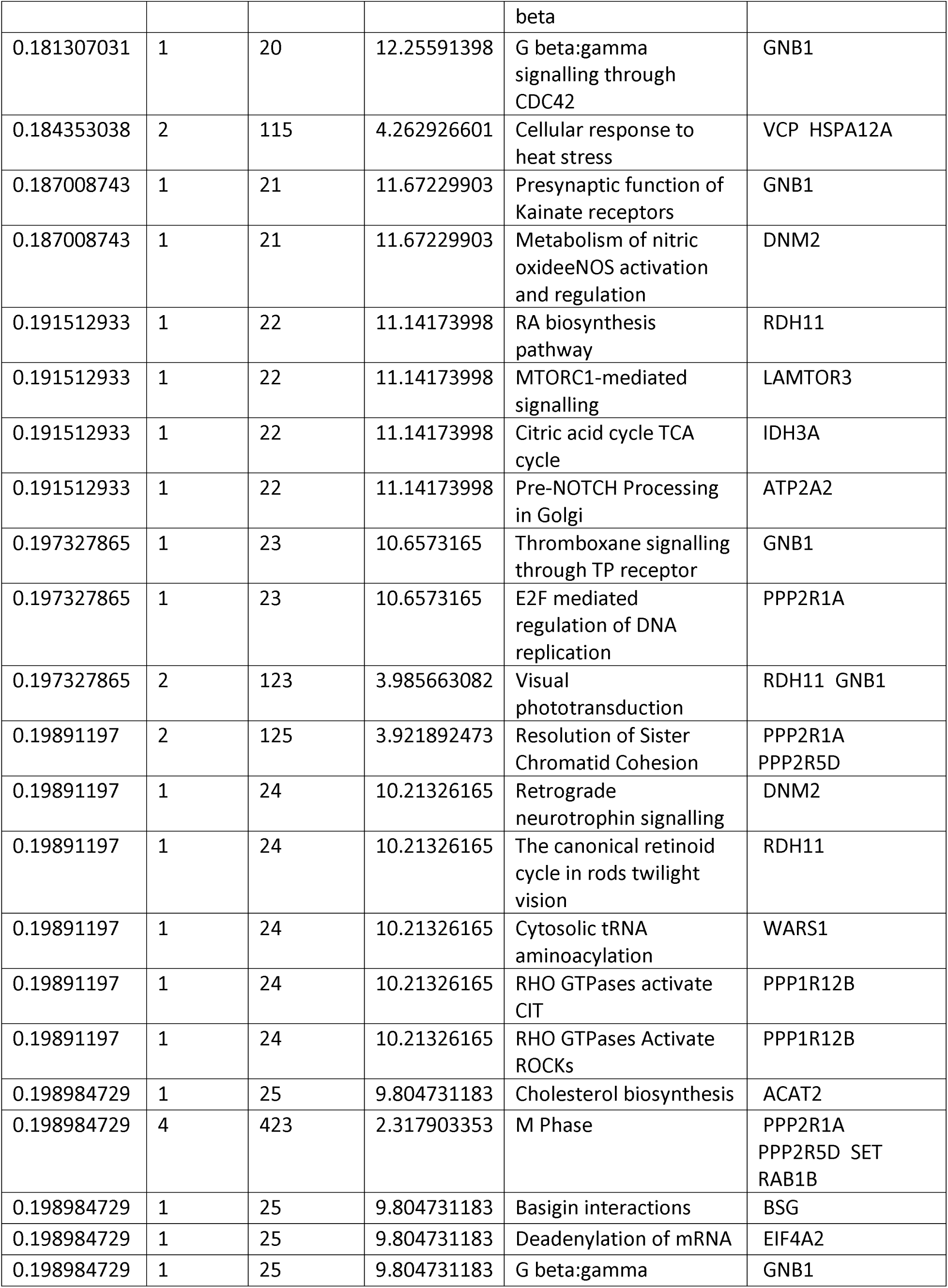
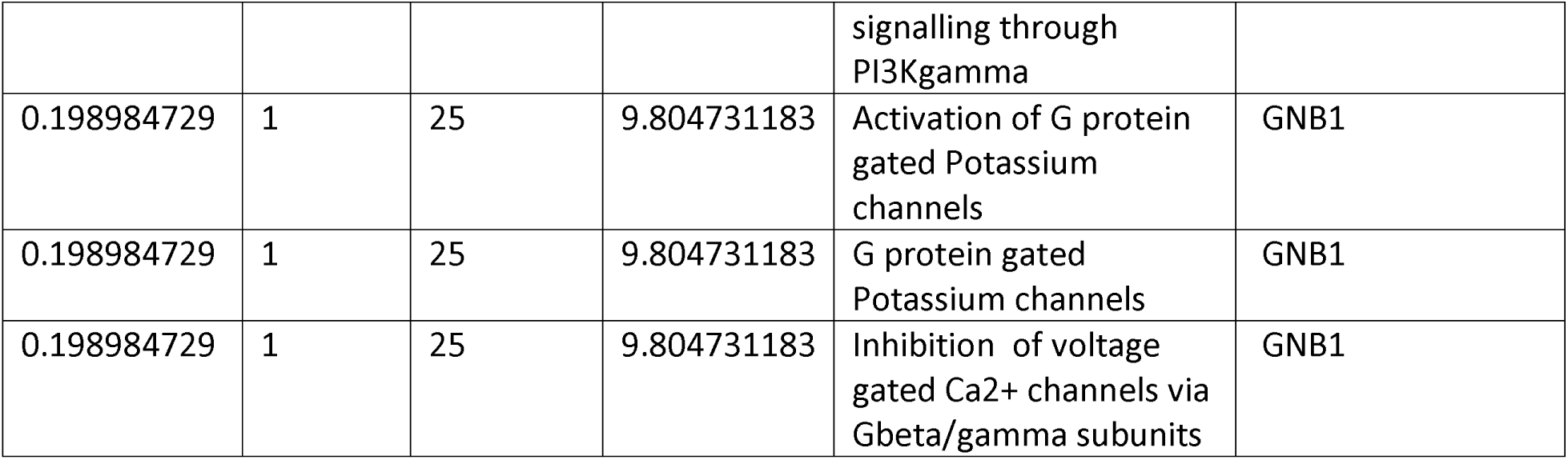
Enriched Pathways in OUD. The table shows the enrichment FDR, the number of proteins/genes enriched, the total number of genes in the pathway, the fold enrichment, the pathway that is enriched, and the genes/proteins that were found to be enriched in the pathway for OUD samples. A simplified view of the data via a bar plot is included in Figure 2.

**Table 8.**
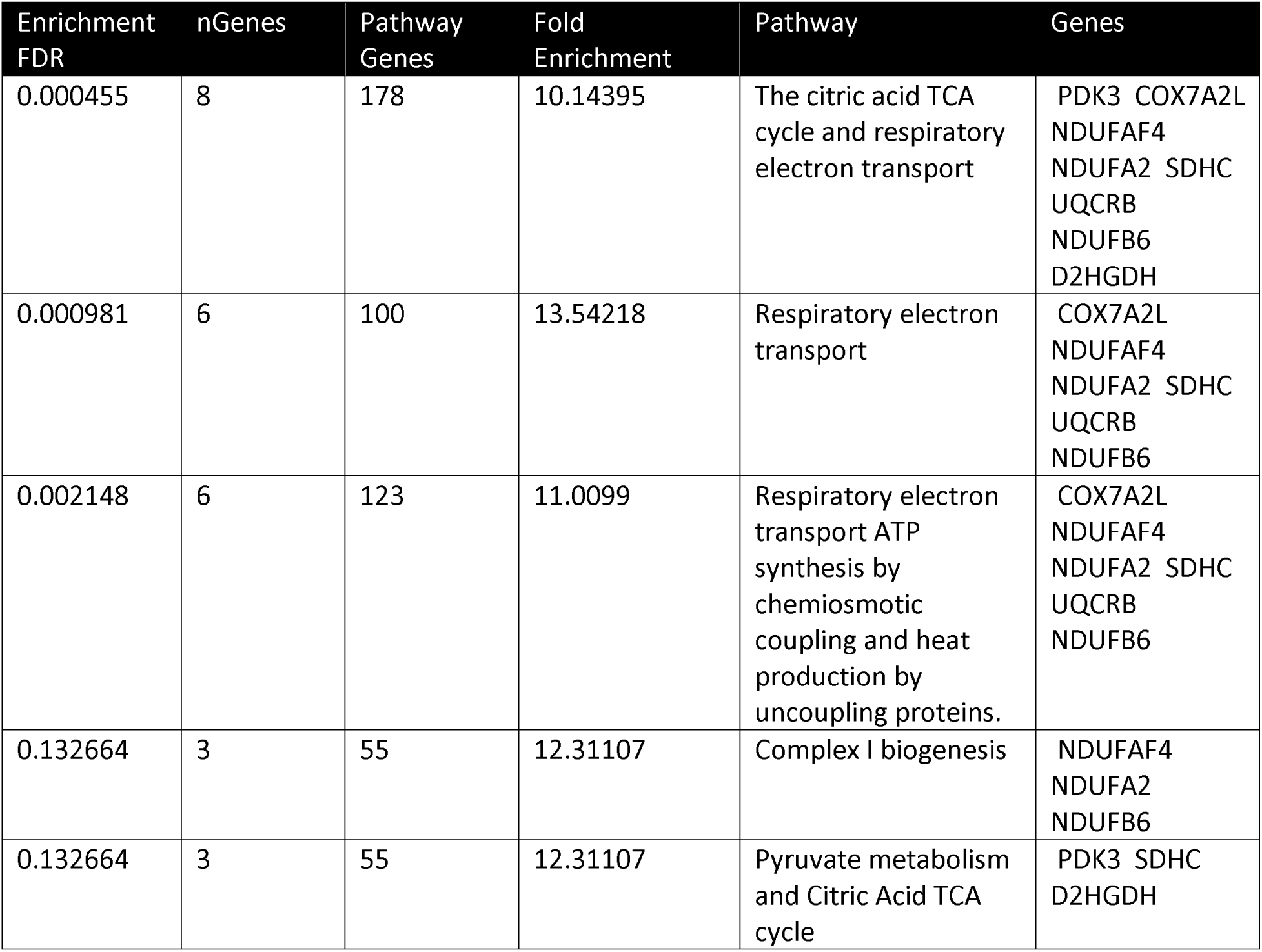
Enriched Pathways in AUD+ OUD. The table shows the enrichment FDR, the number of proteins/genes enriched, the total number of genes in the pathway, the fold enrichment, the pathway that is enriched, and the genes/proteins that were found to be enriched in the pathway for AUD + OUD samples. A simplified view of the data via a bar plot is included in Figure 3.

Four proteins (IDH3A, RAB5B, SORD, and SSBP1) overlapped between AUD and OUD, three proteins (CCDC124, EIF3F, and PPPR1R1B) overlapped between AUD and AUD + OUD, and 13 proteins (ATP2B1, ATP6V1G2, CCBL2, COA6, CPNE5, EIF1, RAB1B, RDH11, SSBP1, TNR, UBE2V2, VCP, and PCBP2) overlapped between OUD and AUD + OUD. A Venn Diagram demonstrating the pattern of protein overlap is shown in Figure 4.

**Figure 4.**
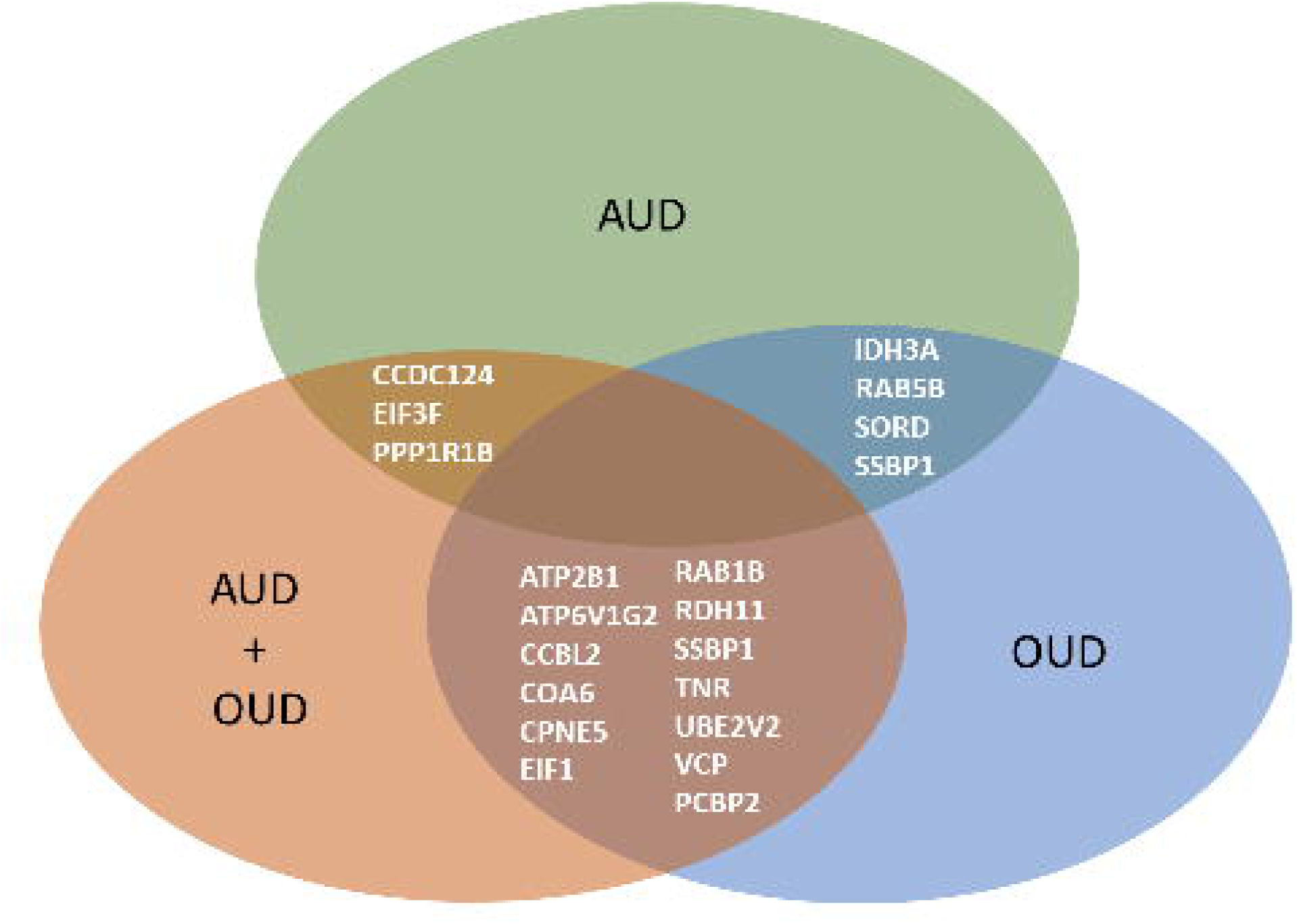
Venn Diagram of Overlapping Proteins in Each SUD. In green is alcohol use disorder (AUD), blue is opioid use disorder (OUD), and orange is combined AUD and OUD.

## Discussion

To our knowledge, this study represents the first comprehensive proteomic analysis in postmortem brain of individuals with OUD and AUD. We identified alterations in proteins not previously reported in SUD. In addition, we found that chronic exposure to opioids and alcohol alters both unique and common protein pathways, and when combined, certain pathways are enriched.

### Alcohol Use Disorder

In general, AUD affects proteins involved in metabolic pathways, including NDUFC2, IDH3A, AUH, GGPS1, SORD, and SLC16A1. These proteins are involved in the respiratory electron transport chain (NDUFC2)^26, 27^, TCA cycle (IDH3A)^28, 29^, amino acid catabolism (AUH)^30, 31^, cholesterol biosynthesis (GGPS1)^32^, and ketose-polyol conversions (SORD)^33, 34^. This is in line with clinical data showing that thiamine is often deficient in AUD due to rapid metabolism of carbohydrates^35^. Alcohol interferes with the conversion of thiamine to the active form thiamine pyrophosphate, which affects ATP production from glucose metabolism^36^. This may explain the upregulation of proteins involved in energy metabolism such as SLC16A1, AUH, and NDUFC2. SLC16A1 is a proton monocarboxylate transporter that moves molecules such as lactate, pyruvate, and ketones between mitochondria and cytoplasm^37^. AUH is involved in catabolism of leucine^30, 31^, and NDUFC2 is associated with the mitochondrial respiratory chain enzyme NADH dehydrogenase^26, 27^.

AUD is also known to affect GABAergic, glutamatergic, glycine, adenosine, serotonin, dopamine, opioid, nicotinic, and endocannabinoid signaling^38–41^. We found alterations in proteins in these signaling pathways, including PDPK1, GNAO1, RAB5B, and PPP1R1B. PDPK1 plays a role in calcium mediated signaling and protein kinase B activity, among other signaling pathways^42–44^. RAB5B is involved in vesicular trafficking and is part of the GTPase family^45, 46^. Upregulation of Rab5b after glutamate receptor stimulation was associated with reduced neuronal excitotoxic injury^47^, suggesting a possible protective mechanism for chronic AUD. PPP1R1B is known as a dopamine and cAMP regulated neuronal phosphoprotein that is important in many signaling pathways such as calcium signaling, cAMP signaling, and AKT signaling^48–50^.

We identified a few proteins not previously implicated in AUD such as RPS27 and CCDC124. RPS27 is a ribosomal protein subunit^51^ involved in DNA repair, cell division, and ribosomal biogenesis^52, 53^ with known localization to glioblastoma tumor cells and neurons, but not healthy astrocytes^54^. It is possible that RPS27 serves as a stress response to the neuronal damage caused by alcohol use. CCDC124 is a coiled-coil domain containing protein that is involved in RNA binding, transcription coactivation, and regulation of cell cycle/division^55^. The protein interacts with Ras-Guanine Nucleotide Exchange Factor1B (RasGEF1B), which is involved in RAP2 signaling^55^. RAP2 signaling in turn mediates mechanoresponses^56, 57^. Further research is necessary to determine if alcohol is responsible for impaired mechanoresponses in neuronal cells, as there is evidence of alcohol damaging the epithelial cell barriers by disrupting RAP2 activity in the intestine^58^.

### Opioid Use Disorder

The opioid pathway is highly dependent on G protein signaling, which can modulate other signaling pathways including calcium, potassium, MAPK, and βγ signaling pathways^59, 60^. Our proteomic data show alterations in these signaling pathways in OUD, particularly calcium signaling and opioid signaling through G protein signaling.

We also found alterations in proteins involved in the processes of translation and respiratory electron transport metabolism. The abundance of proteins upregulated in these pathways, including RPL32, RPS29, RPL7, RPS12, and LAMTOR3 suggests increased energy requirements in the brain of these individuals. LAMTOR3 is a scaffold protein found in late endosomes that is involved in amino acid sensing and activation of mTORC1^61, 62^. It is part of the regulator complex that activates Rag in response to energy levels, amino acids, and growth factors^62^.

We have previously reported dysregulation of angiogenesis and endothelial cell function in OUD^21^. In the present work, we found alterations in proteins involved in platelet homeostasis including PPP2R5D, GNB1, PPP2R1A, ATP2B1, ATP2A2, and ATP2B4. The ATPase proteins are important for coagulation, endothelial cell signaling, and neuro-immunomodulatory regulation^63–65^. Further, PPP2R5D is implicated in angiogenesis^66^ and GNB1 has been found to be related to VEGF signaling in renal carcinoma^67^. Further investigation on how opioids modulate vascularization would be important in understanding the pathophysiology of OUD.

### Opioid and Alcohol Use Disorder comorbidity

Surprisingly, when examining proteins that are altered in individuals with comorbid opioid and alcohol use disorder compared to controls, we found only a few proteins that overlapped with the alcohol or opioid only substance users. In general, the energy metabolism pathways overlapped between the comorbidity and each of the individual disorder groups. However, no specific protein overlapped in all three groups as seen in Figure 4. This suggests that the protein changes may not be just additive, but that the two SUDs combined produce synergistic effects on the brain leading to greater enrichment of specific pathways.

The overlapping proteins between co-morbid substance use are mainly involved in the TCA cycle, respiratory transport, and mitochondria function, suggesting that a higher level of metabolic and respiratory stress is produced in the brain with chronic exposure to both alcohol and opioids. Indeed, several TCA cycle proteins including D2HGDH, UQCRB, COX7A2L, NDUFS3, NDUFAF4, NDUFB6, NDUFA2, SDHC, and PDK3 are upregulated when both SUDs are present. In addition, interleukin signaling is strongly enriched, with proteins such as TAB3, ITGAM, HSPA9, CAPZA1, and TXLNA being more pronounced in the combined SUDs. Further research examining the mechanisms by which these biological pathways are altered in the presence of AUD+OUD may lead to identification of therapeutic targets that could help reduce the cellular stress and related damage caused by these substances.

### Comparison to Other Studies

Our findings are in line with several other GWAS studies demonstrating metabolism, trafficking, and immune signaling are altered in AUD and OUD^21, 68–72^. However, proteomics of both SUDs combined is limited^68, 73^. Our study is the first to report data suggesting a synergistic effect of alcohol and opioids on metabolism and cellular stress pathways.

### Limitations

The analysis was performed in a conservative manner where only the proteins found to be different from controls were analyzed. Relaxing this criterion could have yielded more proteins of significance when analyzing between groups. This criterion was kept, however, to ensure a more rigid analysis with higher confidence in the data. By performing an ANOVA analysis followed by a sub-analysis and then linear regression on the proteins found to be of significance, this could have eliminated any significant proteins that were affected by covariance first, leading to false negatives.

Most of the samples in our study are from middle age white males. Future studies with a larger and more diverse cohort are needed to help generalize the current results. Further, larger sample sizes are needed to determine the effect of covariates included in this study, such as age, gender and pH, as well as the effects of medical comorbidities and other substances such as benzodiazepines used by some individuals, which were not investigated in this study.

## Conclusions

The present study demonstrates that alcohol and opioid chronic exposure leads to specific protein alterations in the brain, with both common and unique pathways affected by each of the disorders. The combination of both disorders appears to have a synergistic effect instead of just an additive effect given the limited number of overlapping protein changes. In addition, we identified proteins that were not previously reported to be involved in substance use disorders such as RPS27 and CCDC124. Further investigation of these proteins is warranted for a better understanding the pathophysiology of alcohol and opioid use disorders.

## Acknowledgments

We are grateful for the invaluable donations and participation from families, as well as for the generous collaboration of the medical examiners at the Harris County Institute of Forensic Sciences. This study was supported by R01DA044859 to CWB. The University of Texas System provided funding for the Neuropsychiatric Proteome Database, for which proteomics data from brain tissue was generated by the Mass Spectrometry Core at the University of Texas Medical Branch.

## Author Contributions

EL, CWB, and SS were responsible for the study concept and design. LS performed the proteomics analysis. EL, LS, ALT, TDM, SS, and and CWB assisted with data analysis and interpretation of findings. EL drafted the manuscript. EL, LS, ALT, TDM, SS, and CWB provided critical revision of the manuscript for important intellectual content. All authors critically reviewed content and approved final version for publication.

---

Supplemental Figure 1 – Reacfoam graph of AUD samples. The more intense the yellow, the higher the enrichment of that pathway.

Supplemental Figure 2 – Reacfoam graph of OUD samples. The more intense the yellow, the higher the enrichment of that pathway.

Supplemental Figure 3 – Reacfoam graph of AUD+OUD samples. The more intense the yellow, the higher the enrichment of that pathway.

Supplemental Table 1 – Demographics of Samples. The demographics of the post-mortem samples including diagnosis, race, gender, urine drug screen (UDS) results, age, and diagnosis are included.

Supplemental Table 2 – Protein Levels. The protein levels of each protein from each sample are listed.

Supplemental Table 3 – ANOVA Results of Significant Proteins. The ANOVA results of each protein that was found to have some difference between controls and substance use was included for further analysis through linear regression analysis if the p value of that protein was <0.05.

Supplemental Table 4 – Linear Regression Results of Significant Proteins. Proteins found to be of significance from the initial ANOVA screen were selected for further analysis in linear regression to incorporate covariates including age, ethnicity, gender, pH, and PMI. Each protein’s name is listed next to the model coefficients table title. The p value of each appropriate covariate or variable is listed in the table. The R and R2 values are included above the table to demonstrate the model fit measures.

