## Supplementary figures and images for "Alcohol, Opioid, and Combined Alcohol and Opioid Use Disorders Affect Shared and Unique Pathways: A Proteomic Analysis of Postmortem Brains"

### Supplemental Figure 1

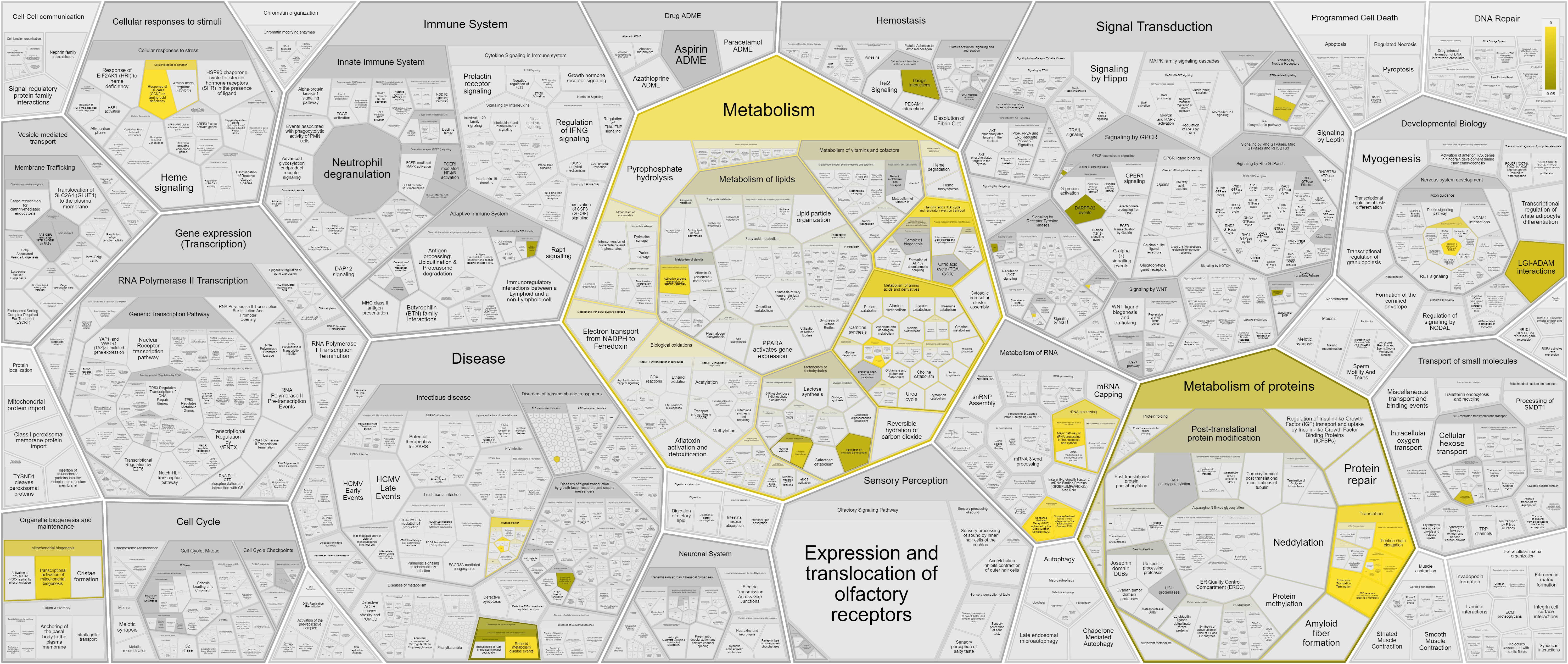

### Supplemental Figure 2

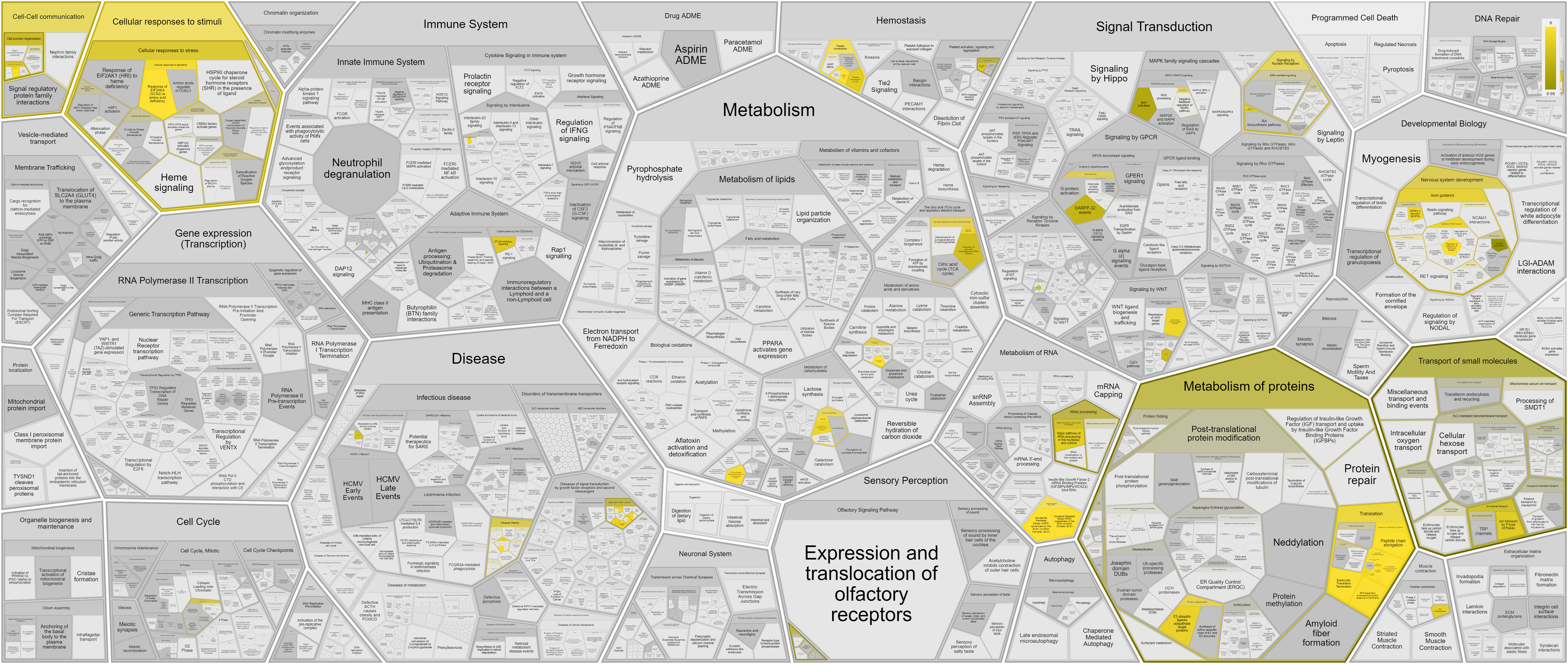

### Supplemental Figure 3

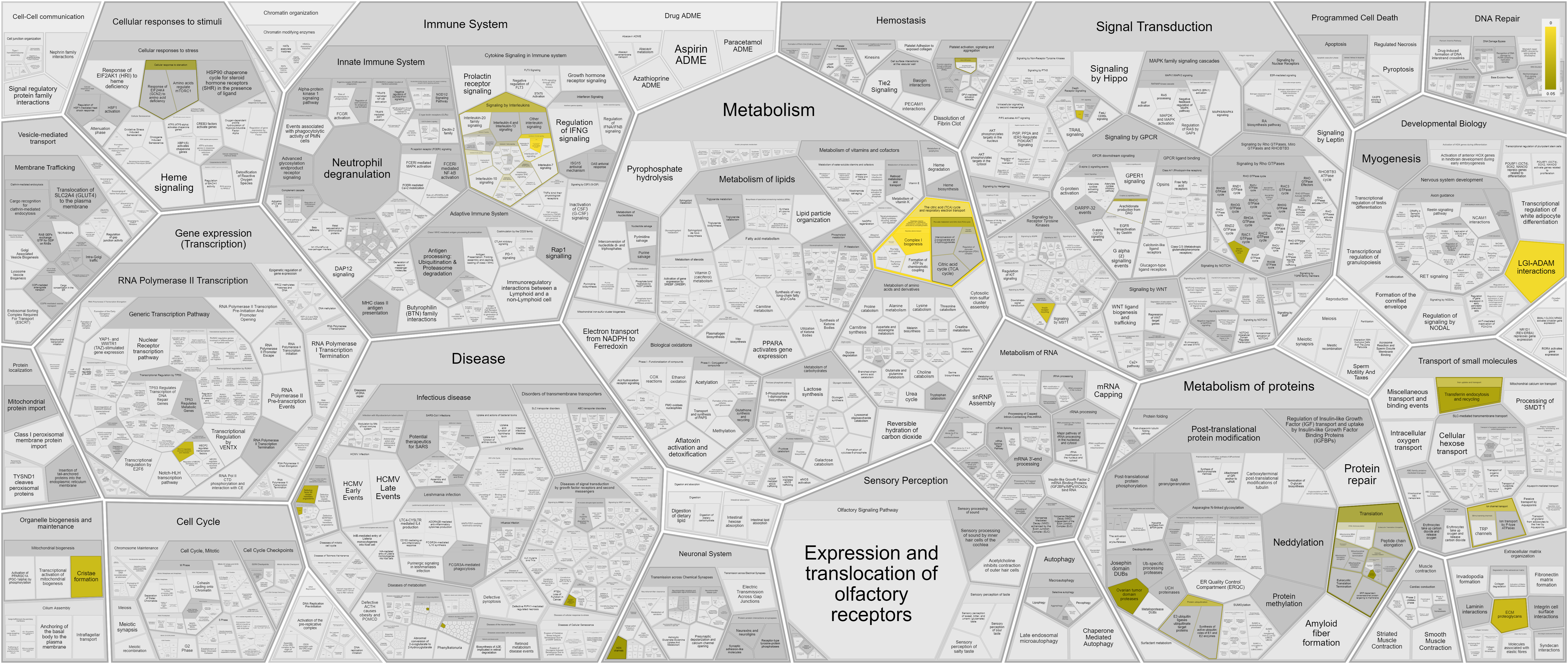
