## Supplemental Table 4 for "Alcohol, Opioid, and Combined Alcohol and Opioid Use Disorders Affect Shared and Unique Pathways: A Proteomic Analysis of Postmortem Brains"

### Results

#### Descriptives

Descriptives

| | Arm | GGPS1 | WFS1 | NDUFC2 | DDX3X | PDPK1 | RBP4 | ADAM11 | HSPA12A | PRKCG | ITPKA | ACADS | NEFM | EEF1G | RPS27 | GGT5 | ALDH4A1 | [illegible] |
|---|---|---|---|---|---|---|---|---|---|---|---|---|---|---|---|---|---|---|
| Mean | alcohol use disorder | 22.8 | 25.0 | 28.1 | 24.0 | 23.1 | 23.0 | 25.5 | 29.7 | 28.0 | 23.6 | 25.8 | 30.6 | 29.6 | 24.9 | 24.2 | 30.2 | |
| | control | 23.6 | 24.1 | 26.8 | 24.7 | 24.4 | 23.5 | 24.2 | 30.1 | 28.4 | 25.1 | 24.6 | 31.3 | 29.8 | 23.4 | 23.3 | 29.8 | |
| | opioid use disorder | 23.7 | 24.7 | 27.8 | 23.9 | 23.6 | 23.7 | 24.1 | 29.6 | 28.3 | 24.2 | 25.5 | 30.8 | 29.6 | 24.2 | 23.6 | 29.9 | |
| | opioid+alcohol use disorder | 23.0 | 23.7 | 26.2 | 24.8 | 24.1 | 23.6 | 23.8 | 30.2 | 28.8 | 25.6 | 23.9 | 31.1 | 29.9 | 23.1 | 23.3 | 29.9 | |

#### Linear Regression

Model Fit Measures

| Model | R | $R^2$ |
|---|---|---|
| 1 | 0.791 | 0.626 |

Model Coefficients - GGPS1

| Predictor | Estimate | SE | t | p |
|---|---|---|---|---|
| Intercept [a] | 20.46204 | 2.5177 | 8.127 | < .001 |
| PMI converted to hours | 0.00645 | 0.0106 | 0.607 | 0.550 |
| pH | 0.41454 | 0.3676 | 1.128 | 0.271 |
| Arm: | | | | |
| alcohol use disorder – control | -1.08318 | 0.2649 | -4.089 | < .001 |
| opioid use disorder – control | 0.11530 | 0.2478 | 0.465 | 0.646 |
| opioid+alcohol use disorder – control | -0.55423 | 0.2931 | -1.891 | 0.071 |
| Age Cateogry: | | | | |
| 50-64 – 30-49 | -0.04929 | 0.2640 | -0.187 | 0.853 |
| 50-65 – 30-49 | 0.58857 | 0.4057 | 1.451 | 0.160 |
| 65+ – 30-49 | 0.54711 | 0.3059 | 1.788 | 0.086 |
| <30 – 30-49 | -0.45824 | 0.3198 | -1.433 | 0.165 |
| Ethnicity: | | | | |
| Asian – White | -0.49856 | 0.5326 | -0.936 | 0.359 |
| Black – White | 0.26803 | 0.2168 | 1.236 | 0.228 |
| Hispanic – White | -0.53757 | 0.4147 | -1.296 | 0.207 |
| Gender: | | | | |
| Male – Female | 0.42076 | 0.2563 | 1.642 | 0.114 |

[a] Represents reference level

#### Linear Regression

Model Fit Measures

| Model | R | $R^2$ |
|---|---|---|
| 1 | 0.660 | 0.436 |

Model Coefficients - WFS1

| Predictor | Estimate | SE | t | p |
|---|---|---|---|---|
| Intercept [a] | 20.8573 | 4.3090 | 4.8404 | < .001 |
| Arm: | | | | |
| alcohol use disorder – control | 0.8070 | 0.4534 | 1.7799 | 0.088 |
| opioid use disorder – control | 0.6726 | 0.4241 | 1.5857 | 0.126 |
| opioid+alcohol use disorder – control | -0.0838 | 0.5016 | -0.1670 | 0.869 |
| Age Cateogry: | | | | |
| 50-64 – 30-49 | -0.4580 | 0.4519 | -1.0135 | 0.321 |
| 50-65 – 30-49 | -0.2668 | 0.6943 | -0.3843 | 0.704 |
| 65+ – 30-49 | -0.2139 | 0.5236 | -0.4085 | 0.687 |
| <30 – 30-49 | -0.0953 | 0.5474 | -0.1741 | 0.863 |
| Ethnicity: | | | | |
| Asian – White | -0.3655 | 0.9115 | -0.4010 | 0.692 |
| Black – White | -0.0209 | 0.3711 | -0.0564 | 0.955 |
| Hispanic – White | 0.4706 | 0.7097 | 0.6631 | 0.514 |
| Gender: | | | | |
| Male – Female | 0.4019 | 0.4386 | 0.9164 | 0.369 |
| PMI converted to hours | 0.0196 | 0.0182 | 1.0786 | 0.291 |
| pH | 0.3809 | 0.6292 | 0.6054 | 0.551 |

[a] Represents reference level

### Linear Regression

Model Fit Measures

| Model | R | $R^2$ |
|---|---|---|
| 1 | 0.791 | 0.625 |

Model Coefficients - NDUFC2

| Predictor | Estimate | SE | t | p |
|---|---|---|---|---|
| Intercept [a] | 21.5622 | 6.3692 | 3.385 | 0.002 |
| Arm: | | | | |
| alcohol use disorder – control | 1.4855 | 0.6702 | 2.216 | 0.036 |
| opioid use disorder – control | 1.2001 | 0.6269 | 1.914 | 0.068 |
| opioid+alcohol use disorder – control | -0.3350 | 0.7414 | -0.452 | 0.655 |
| Age Cateogry: | | | | |
| 50-64 – 30-49 | 0.7629 | 0.6680 | 1.142 | 0.265 |
| 50-65 – 30-49 | 1.3183 | 1.0262 | 1.285 | 0.211 |
| 65+ – 30-49 | -1.3826 | 0.7739 | -1.787 | 0.087 |
| <30 – 30-49 | -0.4737 | 0.8091 | -0.585 | 0.564 |
| Ethnicity: | | | | |
| Asian – White | -3.0692 | 1.3473 | -2.278 | 0.032 |
| Black – White | 0.5413 | 0.5485 | 0.987 | 0.334 |
| Hispanic – White | -0.3397 | 1.0490 | -0.324 | 0.749 |
| Gender: | | | | |
| Male – Female | 0.2888 | 0.6483 | 0.445 | 0.660 |
| PMI converted to hours | 0.0391 | 0.0269 | 1.454 | 0.159 |
| pH | 0.5566 | 0.9300 | 0.598 | 0.555 |

[a] Represents reference level

### Linear Regression

Model Fit Measures

| Model | R | $R^2$ |
|---|---|---|
| 1 | 0.686 | 0.470 |

Model Coefficients - DDX3X

| Predictor | Estimate | SE | t | p |
|---|---|---|---|---|
| Intercept [a] | 25.8596 | 3.8402 | 6.734 | < .001 |
| Arm: | | | | |
| alcohol use disorder – control | -0.2936 | 0.4041 | -0.727 | 0.474 |
| opioid use disorder – control | -0.6210 | 0.3780 | -1.643 | 0.113 |
| opioid+alcohol use disorder – control | 0.0750 | 0.4470 | 0.168 | 0.868 |
| Age Cateogry: | | | | |
| 50-64 – 30-49 | 0.6367 | 0.4028 | 1.581 | 0.127 |
| 50-65 – 30-49 | -0.2111 | 0.6187 | -0.341 | 0.736 |
| 65+ – 30-49 | 0.7610 | 0.4666 | 1.631 | 0.116 |
| <30 – 30-49 | 0.3272 | 0.4878 | 0.671 | 0.509 |
| Ethnicity: | | | | |
| Asian – White | 1.2202 | 0.8123 | 1.502 | 0.146 |
| Black – White | 0.2651 | 0.3307 | 0.802 | 0.431 |
| Hispanic – White | 0.7548 | 0.6325 | 1.193 | 0.244 |
| Gender: | | | | |
| Male – Female | -0.3433 | 0.3909 | -0.878 | 0.388 |
| PMI converted to hours | -0.0158 | 0.0162 | -0.972 | 0.341 |
| pH | -0.1522 | 0.5607 | -0.271 | 0.788 |

[a] Represents reference level

### Linear Regression

Model Fit Measures

| Model | R | $R^2$ |
|---|---|---|
| 1 | 0.784 | 0.615 |

Model Coefficients - PDPK1

| Predictor | Estimate | SE | t | p |
|---|---|---|---|---|
| Intercept [a] | 22.2695 | 3.6496 | 6.1019 | < .001 |
| Arm: | | | | |
| alcohol use disorder – control | -1.3112 | 0.3840 | -3.4142 | 0.002 |
| opioid use disorder – control | -0.4430 | 0.3592 | -1.2332 | 0.229 |
| opioid+alcohol use disorder – control | -0.1415 | 0.4248 | -0.3330 | 0.742 |
| Age Cateogry: | | | | |
| 50-64 – 30-49 | 0.4286 | 0.3828 | 1.1197 | 0.274 |
| 50-65 – 30-49 | 0.0499 | 0.5880 | 0.0848 | 0.933 |
| 65+ – 30-49 | 0.5689 | 0.4434 | 1.2830 | 0.212 |
| <30 – 30-49 | -0.8910 | 0.4636 | -1.9218 | 0.067 |
| Ethnicity: | | | | |
| Asian – White | -0.7279 | 0.7720 | -0.9428 | 0.355 |
| Black – White | 0.3420 | 0.3143 | 1.0881 | 0.287 |
| Hispanic – White | -0.6480 | 0.6011 | -1.0780 | 0.292 |
| Gender: | | | | |
| Male – Female | 0.3028 | 0.3715 | 0.8150 | 0.423 |
| PMI converted to hours | 0.0377 | 0.0154 | 2.4461 | 0.022 |
| pH | 0.0971 | 0.5329 | 0.1822 | 0.857 |

[a] Represents reference level

### Linear Regression

Model Fit Measures

| Model | R | $R^2$ |
|---|---|---|
| 1 | 0.570 | 0.325 |

Model Coefficients - RBP4

| Predictor | Estimate | SE | t | p |
|---|---|---|---|---|
| Intercept [a] | 22.8670 | 3.5348 | 6.469 | < .001 |
| Arm: | | | | |
| alcohol use disorder – control | -0.8897 | 0.3719 | -2.392 | 0.025 |
| opioid use disorder – control | -0.2048 | 0.3479 | -0.589 | 0.562 |
| opioid+alcohol use disorder – control | -0.1079 | 0.4115 | -0.262 | 0.795 |
| Age Cateogry: | | | | |
| 50-64 – 30-49 | -0.5948 | 0.3707 | -1.604 | 0.122 |
| 50-65 – 30-49 | -0.3910 | 0.5695 | -0.686 | 0.499 |
| 65+ – 30-49 | -0.3141 | 0.4295 | -0.731 | 0.472 |
| <30 – 30-49 | -0.6938 | 0.4490 | -1.545 | 0.135 |
| Ethnicity: | | | | |
| Asian – White | -0.6732 | 0.7477 | -0.900 | 0.377 |
| Black – White | -0.1062 | 0.3044 | -0.349 | 0.730 |
| Hispanic – White | 0.0796 | 0.5822 | 0.137 | 0.892 |
| Gender: | | | | |
| Male – Female | 0.0568 | 0.3598 | 0.158 | 0.876 |
| PMI converted to hours | -0.0123 | 0.0149 | -0.821 | 0.420 |
| pH | 0.2437 | 0.5161 | 0.472 | 0.641 |

[a] Represents reference level

### Linear Regression

Model Fit Measures

| Model | R | $R^2$ |
|---|---|---|
| 1 | 0.645 | 0.417 |

Model Coefficients - ADAM11

| Predictor | Estimate | SE | t | p |
|---|---|---|---|---|
| Intercept [a] | 27.63090 | 6.1175 | 4.517 | < .001 |
| Arm: | | | | |
| alcohol use disorder – control | 1.56558 | 0.6437 | 2.432 | 0.023 |
| opioid use disorder – control | -0.38748 | 0.6022 | -0.643 | 0.526 |
| opioid+alcohol use disorder – control | -0.88127 | 0.7121 | -1.238 | 0.228 |
| Age Cateogry: | | | | |
| 50-64 – 30-49 | 0.23363 | 0.6416 | 0.364 | 0.719 |
| 50-65 – 30-49 | -1.10262 | 0.9857 | -1.119 | 0.274 |
| 65+ – 30-49 | -0.46979 | 0.7433 | -0.632 | 0.533 |
| <30 – 30-49 | 1.20555 | 0.7771 | 1.551 | 0.134 |
| Ethnicity: | | | | |
| Asian – White | -1.16436 | 1.2940 | -0.900 | 0.377 |
| Black – White | -0.77067 | 0.5268 | -1.463 | 0.156 |
| Hispanic – White | -0.59854 | 1.0076 | -0.594 | 0.558 |
| Gender: | | | | |
| Male – Female | -0.29452 | 0.6227 | -0.473 | 0.641 |
| PMI converted to hours | -0.00721 | 0.0258 | -0.279 | 0.782 |
| pH | -0.43224 | 0.8932 | -0.484 | 0.633 |

[a] Represents reference level

### Linear Regression

Model Fit Measures

| Model | R | $R^2$ |
|---|---|---|
| 1 | 0.767 | 0.589 |

Model Coefficients - HSPA12A

| Predictor | Estimate | SE | t | p |
|---|---|---|---|---|
| Intercept [a] | 29.96345 | 1.56526 | 19.1428 | < .001 |
| Arm: | | | | |
| alcohol use disorder – control | -0.20744 | 0.16470 | -1.2594 | 0.220 |
| opioid use disorder – control | -0.40841 | 0.15407 | -2.6508 | 0.014 |
| opioid+alcohol use disorder – control | 0.09654 | 0.18220 | 0.5299 | 0.601 |
| Age Cateogry: | | | | |
| 50-64 – 30-49 | 0.31293 | 0.16416 | 1.9062 | 0.069 |
| 50-65 – 30-49 | 0.04877 | 0.25220 | 0.1934 | 0.848 |
| 65+ – 30-49 | 0.15284 | 0.19019 | 0.8036 | 0.429 |
| <30 – 30-49 | 0.16547 | 0.19884 | 0.8321 | 0.414 |
| Ethnicity: | | | | |
| Asian – White | 0.21610 | 0.33110 | 0.6527 | 0.520 |
| Black – White | -0.00703 | 0.13479 | -0.0522 | 0.959 |
| Hispanic – White | 0.29992 | 0.25780 | 1.1634 | 0.256 |
| Gender: | | | | |
| Male – Female | -0.18154 | 0.15933 | -1.1394 | 0.266 |
| PMI converted to hours | -0.00811 | 0.00661 | -1.2274 | 0.232 |
| pH | 0.04630 | 0.22855 | 0.2026 | 0.841 |

[a] Represents reference level

#### Linear Regression

Model Fit Measures

| Model | R | $R^2$ |
|---|---|---|
| 1 | 0.769 | 0.591 |

Model Coefficients - PRKCG

| Predictor | Estimate | SE | t | p |
|---|---|---|---|---|
| Intercept [a] | 30.72391 | 1.34268 | 22.882 | < .001 |
| Arm: | | | | |
| alcohol use disorder – control | -0.16468 | 0.14128 | -1.166 | 0.255 |
| opioid use disorder – control | -0.08404 | 0.13216 | -0.636 | 0.531 |
| opioid+alcohol use disorder – control | 0.24645 | 0.15629 | 1.577 | 0.128 |
| Age Cateogry: | | | | |
| 50-64 – 30-49 | 0.20642 | 0.14082 | 1.466 | 0.156 |
| 50-65 – 30-49 | -0.17942 | 0.21633 | -0.829 | 0.415 |
| 65+ – 30-49 | 0.10709 | 0.16314 | 0.656 | 0.518 |
| <30 – 30-49 | 0.27206 | 0.17057 | 1.595 | 0.124 |
| Ethnicity: | | | | |
| Asian – White | 0.19233 | 0.28402 | 0.677 | 0.505 |
| Black – White | 0.06061 | 0.11563 | 0.524 | 0.605 |
| Hispanic – White | -0.09067 | 0.22114 | -0.410 | 0.685 |
| Gender: | | | | |
| Male – Female | -0.23532 | 0.13667 | -1.722 | 0.098 |
| PMI converted to hours | -0.00654 | 0.00567 | -1.154 | 0.260 |
| pH | -0.31928 | 0.19605 | -1.629 | 0.116 |

[a] Represents reference level

#### Linear Regression

Model Fit Measures

| Model | R | $R^2$ |
|---|---|---|
| 1 | 0.688 | 0.474 |

Model Coefficients - ITPKA

| Predictor | Estimate | SE | t | p |
|---|---|---|---|---|
| Intercept [a] | 29.95292 | 5.8707 | 5.102 | < .001 |
| Arm: | | | | |
| alcohol use disorder – control | -0.80526 | 0.6177 | -1.304 | 0.205 |
| opioid use disorder – control | -0.26172 | 0.5779 | -0.453 | 0.655 |
| opioid+alcohol use disorder – control | 0.71209 | 0.6834 | 1.042 | 0.308 |
| Age Cateogry: | | | | |
| 50-64 – 30-49 | 0.82698 | 0.6157 | 1.343 | 0.192 |
| 50-65 – 30-49 | -0.26849 | 0.9459 | -0.284 | 0.779 |
| 65+ – 30-49 | 0.96731 | 0.7133 | 1.356 | 0.188 |
| <30 – 30-49 | 0.17054 | 0.7458 | 0.229 | 0.821 |
| Ethnicity: | | | | |
| Asian – White | 1.84410 | 1.2418 | 1.485 | 0.151 |
| Black – White | 0.37101 | 0.5056 | 0.734 | 0.470 |
| Hispanic – White | 1.10013 | 0.9669 | 1.138 | 0.266 |
| Gender: | | | | |
| Male – Female | -0.09546 | 0.5976 | -0.160 | 0.874 |
| PMI converted to hours | 0.00587 | 0.0248 | 0.237 | 0.815 |
| pH | -0.89443 | 0.8572 | -1.043 | 0.307 |

[a] Represents reference level

#### Linear Regression

Model Fit Measures

| Model | R | $R^2$ |
|---|---|---|
| 1 | 0.776 | 0.601 |

Model Coefficients - ACADS

| Predictor | Estimate | SE | t | p |
|---|---|---|---|---|
| Intercept [a] | 17.3718 | 4.9660 | 3.4981 | 0.002 |
| Arm: | | | | |
| alcohol use disorder – control | 0.9331 | 0.5225 | 1.7856 | 0.087 |
| opioid use disorder – control | 1.1882 | 0.4888 | 2.4307 | 0.023 |
| opioid+alcohol use disorder – control | 0.0146 | 0.5781 | 0.0252 | 0.980 |
| Age Cateogry: | | | | |
| 50-64 – 30-49 | -0.1991 | 0.5208 | -0.3823 | 0.706 |
| 50-65 – 30-49 | -0.3331 | 0.8001 | -0.4163 | 0.681 |
| 65+ – 30-49 | 0.5852 | 0.6034 | 0.9698 | 0.342 |
| <30 – 30-49 | -1.0218 | 0.6309 | -1.6196 | 0.118 |
| Ethnicity: | | | | |
| Asian – White | -1.3553 | 1.0505 | -1.2902 | 0.209 |
| Black – White | 0.9414 | 0.4277 | 2.2014 | 0.038 |
| Hispanic – White | 0.6451 | 0.8179 | 0.7887 | 0.438 |
| Gender: | | | | |
| Male – Female | 1.2112 | 0.5055 | 2.3961 | 0.025 |
| PMI converted to hours | 0.0182 | 0.0210 | 0.8660 | 0.395 |
| pH | 0.8599 | 0.7251 | 1.1860 | 0.247 |

[a] Represents reference level

#### Linear Regression

Model Fit Measures

| Model | R | $R^2$ |
|---|---|---|
| 1 | 0.557 | 0.310 |

Model Coefficients - NEFM

| Predictor | Estimate | SE | t | p |
|---|---|---|---|---|
| Intercept [a] | 30.5626 | 4.2507 | 7.1900 | < .001 |
| Arm: | | | | |
| alcohol use disorder – control | -0.6429 | 0.4473 | -1.4375 | 0.163 |
| opioid use disorder – control | -0.3036 | 0.4184 | -0.7256 | 0.475 |
| opioid+alcohol use disorder – control | -0.0864 | 0.4948 | -0.1745 | 0.863 |
| Age Cateogry: | | | | |
| 50-64 – 30-49 | 0.4974 | 0.4458 | 1.1158 | 0.276 |
| 50-65 – 30-49 | 0.3810 | 0.6849 | 0.5563 | 0.583 |
| 65+ – 30-49 | 0.6821 | 0.5165 | 1.3206 | 0.199 |
| <30 – 30-49 | 0.4136 | 0.5400 | 0.7660 | 0.451 |
| Ethnicity: | | | | |
| Asian – White | 0.7731 | 0.8991 | 0.8598 | 0.398 |
| Black – White | -0.0334 | 0.3661 | -0.0913 | 0.928 |
| Hispanic – White | 0.7311 | 0.7001 | 1.0443 | 0.307 |
| Gender: | | | | |
| Male – Female | 0.1659 | 0.4327 | 0.3835 | 0.705 |
| PMI converted to hours | -0.0228 | 0.0180 | -1.2718 | 0.216 |
| pH | 0.1235 | 0.6207 | 0.1990 | 0.844 |

[a] Represents reference level

#### Linear Regression

Model Fit Measures

| Model | R | $R^2$ |
|---|---|---|
| 1 | 0.791 | 0.625 |

Model Coefficients - EEF1G

| Predictor | Estimate | SE | t | p |
|---|---|---|---|---|
| Intercept [a] | 30.52434 | 0.70373 | 43.375 | < .001 |
| Arm: | | | | |
| alcohol use disorder – control | -0.15922 | 0.07405 | -2.150 | 0.042 |
| opioid use disorder – control | -0.11266 | 0.06927 | -1.626 | 0.117 |
| opioid+alcohol use disorder – control | 0.12890 | 0.08192 | 1.574 | 0.129 |
| Age Cateogry: | | | | |
| 50-64 – 30-49 | 0.05941 | 0.07381 | 0.805 | 0.429 |
| 50-65 – 30-49 | 0.05020 | 0.11339 | 0.443 | 0.662 |
| 65+ – 30-49 | 0.13420 | 0.08551 | 1.570 | 0.130 |
| <30 – 30-49 | 0.10788 | 0.08940 | 1.207 | 0.239 |
| Ethnicity: | | | | |
| Asian – White | 0.20297 | 0.14886 | 1.363 | 0.185 |
| Black – White | 0.05918 | 0.06060 | 0.976 | 0.339 |
| Hispanic – White | -0.06644 | 0.11590 | -0.573 | 0.572 |
| Gender: | | | | |
| Male – Female | -0.06105 | 0.07163 | -0.852 | 0.402 |
| PMI converted to hours | -0.00249 | 0.00297 | -0.837 | 0.411 |
| pH | -0.11039 | 0.10275 | -1.074 | 0.293 |

[a] Represents reference level

#### Linear Regression

Model Fit Measures

| Model | R | $R^2$ |
|---|---|---|
| 1 | 0.677 | 0.459 |

Model Coefficients - RPS27

| Predictor | Estimate | SE | t | p |
|---|---|---|---|---|
| Intercept [a] | 25.56393 | 5.2230 | 4.89450 | < .001 |
| Arm: | | | | |
| alcohol use disorder – control | 1.48406 | 0.5496 | 2.70033 | 0.012 |
| opioid use disorder – control | 0.47848 | 0.5141 | 0.93069 | 0.361 |
| opioid+alcohol use disorder – control | -0.55657 | 0.6080 | -0.91544 | 0.369 |
| Age Cateogry: | | | | |
| 50-64 – 30-49 | 0.00231 | 0.5478 | 0.00422 | 0.997 |
| 50-65 – 30-49 | 0.09920 | 0.8415 | 0.11788 | 0.907 |
| 65+ – 30-49 | -0.63512 | 0.6346 | -1.00079 | 0.327 |
| <30 – 30-49 | -0.19369 | 0.6635 | -0.29193 | 0.773 |
| Ethnicity: | | | | |
| Asian – White | -0.45256 | 1.1048 | -0.40963 | 0.686 |
| Black – White | -0.10395 | 0.4498 | -0.23111 | 0.819 |
| Hispanic – White | 0.52762 | 0.8602 | 0.61335 | 0.545 |
| Gender: | | | | |
| Male – Female | -0.55203 | 0.5317 | -1.03833 | 0.309 |
| PMI converted to hours | -0.01239 | 0.0221 | -0.56158 | 0.580 |
| pH | -0.17534 | 0.7626 | -0.22992 | 0.820 |

[a] Represents reference level

### Linear Regression

Model Fit Measures

| Model | R | $R^2$ |
|---|---|---|
| 1 | 0.638 | 0.407 |

Model Coefficients - GGT5

| Predictor | Estimate | SE | t | p |
|---|---|---|---|---|
| Intercept [a] | 22.0633 | 3.5146 | 6.278 | < .001 |
| Arm: | | | | |
| alcohol use disorder – control | 0.6682 | 0.3698 | 1.807 | 0.083 |
| opioid use disorder – control | -0.0496 | 0.3460 | -0.143 | 0.887 |
| opioid+alcohol use disorder – control | -0.1614 | 0.4091 | -0.395 | 0.697 |
| Age Cateogry: | | | | |
| 50-64 – 30-49 | -0.1862 | 0.3686 | -0.505 | 0.618 |
| 50-65 – 30-49 | -0.2034 | 0.5663 | -0.359 | 0.723 |
| 65+ – 30-49 | 0.2717 | 0.4270 | 0.636 | 0.531 |
| <30 – 30-49 | -0.1280 | 0.4465 | -0.287 | 0.777 |
| Ethnicity: | | | | |
| Asian – White | -0.8447 | 0.7434 | -1.136 | 0.267 |
| Black – White | 0.2431 | 0.3027 | 0.803 | 0.430 |
| Hispanic – White | 0.2221 | 0.5788 | 0.384 | 0.705 |
| Gender: | | | | |
| Male – Female | -0.1834 | 0.3578 | -0.513 | 0.613 |
| PMI converted to hours | -0.0148 | 0.0148 | -0.998 | 0.328 |
| pH | 0.3128 | 0.5132 | 0.610 | 0.548 |

[a] Represents reference level

### Linear Regression

Model Fit Measures

| Model | R | $R^2$ |
|---|---|---|
| 1 | 0.677 | 0.458 |

Model Coefficients - ALDH4A1

| Predictor | Estimate | SE | t | p |
|---|---|---|---|---|
| Intercept [a] | 27.6962 | 2.01736 | 13.7289 | < .001 |
| Arm: | | | | |
| alcohol use disorder – control | 0.4259 | 0.21227 | 2.0061 | 0.056 |
| opioid use disorder – control | 0.1596 | 0.19858 | 0.8035 | 0.430 |
| opioid+alcohol use disorder – control | 0.0627 | 0.23483 | 0.2670 | 0.792 |
| Age Cateogry: | | | | |
| 50-64 – 30-49 | -0.0476 | 0.21158 | -0.2248 | 0.824 |
| 50-65 – 30-49 | -0.0132 | 0.32504 | -0.0405 | 0.968 |
| 65+ – 30-49 | -0.0712 | 0.24512 | -0.2904 | 0.774 |
| <30 – 30-49 | -0.1516 | 0.25627 | -0.5916 | 0.560 |
| Ethnicity: | | | | |
| Asian – White | 0.0827 | 0.42673 | 0.1938 | 0.848 |
| Black – White | -0.4347 | 0.17373 | -2.5020 | 0.020 |
| Hispanic – White | -0.0294 | 0.33226 | -0.0886 | 0.930 |
| Gender: | | | | |
| Male – Female | 0.0101 | 0.20535 | 0.0493 | 0.961 |
| PMI converted to hours | 5.79e-4 | 0.00852 | 0.0680 | 0.946 |
| pH | 0.3295 | 0.29456 | 1.1185 | 0.274 |

[a] Represents reference level

#### Linear Regression

Model Fit Measures

| Model | R | $R^2$ |
|---|---|---|
| 1 | 0.730 | 0.533 |

Model Coefficients - SLC16A1

| Predictor | Estimate | SE | t | p |
|---|---|---|---|---|
| Intercept [a] | 24.40325 | 3.6108 | 6.7584 | < .001 |
| Arm: | | | | |
| alcohol use disorder – control | 1.21643 | 0.3799 | 3.2016 | 0.004 |
| opioid use disorder – control | 0.73092 | 0.3554 | 2.0565 | 0.051 |
| opioid+alcohol use disorder – control | 0.03280 | 0.4203 | 0.0780 | 0.938 |
| Age Cateogry: | | | | |
| 50-64 – 30-49 | 0.29737 | 0.3787 | 0.7853 | 0.440 |
| 50-65 – 30-49 | -0.54182 | 0.5818 | -0.9313 | 0.361 |
| 65+ – 30-49 | -0.04158 | 0.4387 | -0.0948 | 0.925 |
| <30 – 30-49 | 0.11551 | 0.4587 | 0.2518 | 0.803 |
| Ethnicity: | | | | |
| Asian – White | -1.12751 | 0.7638 | -1.4762 | 0.153 |
| Black – White | -0.31980 | 0.3109 | -1.0285 | 0.314 |
| Hispanic – White | -0.70758 | 0.5947 | -1.1898 | 0.246 |
| Gender: | | | | |
| Male – Female | -0.26787 | 0.3675 | -0.7288 | 0.473 |
| PMI converted to hours | -0.00945 | 0.0153 | -0.6198 | 0.541 |
| pH | 0.03583 | 0.5272 | 0.0680 | 0.946 |

[a] Represents reference level

#### Linear Regression

Model Fit Measures

| Model | R | $R^2$ |
|---|---|---|
| 1 | 0.757 | 0.573 |

Model Coefficients - RUFY3

| Predictor | Estimate | SE | t | p |
|---|---|---|---|---|
| Intercept [a] | 27.16509 | 2.14677 | 12.65391 | < .001 |
| Arm: | | | | |
| alcohol use disorder – control | -0.34716 | 0.22589 | -1.53682 | 0.137 |
| opioid use disorder – control | -0.35676 | 0.21131 | -1.68827 | 0.104 |
| opioid+alcohol use disorder – control | 0.25354 | 0.24989 | 1.01459 | 0.320 |
| Age Cateogry: | | | | |
| 50-64 – 30-49 | 0.45506 | 0.22515 | 2.02115 | 0.055 |
| 50-65 – 30-49 | 0.29346 | 0.34589 | 0.84843 | 0.405 |
| 65+ – 30-49 | 0.44104 | 0.26084 | 1.69082 | 0.104 |
| <30 – 30-49 | -0.04089 | 0.27271 | -0.14992 | 0.882 |
| Ethnicity: | | | | |
| Asian – White | 0.35477 | 0.45410 | 0.78124 | 0.442 |
| Black – White | 0.31588 | 0.18487 | 1.70865 | 0.100 |
| Hispanic – White | 0.19270 | 0.35357 | 0.54501 | 0.591 |
| Gender: | | | | |
| Male – Female | 0.00203 | 0.21852 | 0.00927 | 0.993 |
| PMI converted to hours | -0.00201 | 0.00907 | -0.22142 | 0.827 |
| pH | 0.07932 | 0.31345 | 0.25304 | 0.802 |

[a] Represents reference level

### Linear Regression

Model Fit Measures

| Model | R | $R^2$ |
|---|---|---|
| 1 | 0.518 | 0.269 |

Model Coefficients - GUCY1B3

| Predictor | Estimate | SE | t | p |
|---|---|---|---|---|
| Intercept [a] | 32.31875 | 7.8690 | 4.1071 | < .001 |
| Arm: | | | | |
| alcohol use disorder – control | -1.38272 | 0.8280 | -1.6699 | 0.108 |
| opioid use disorder – control | -1.08496 | 0.7746 | -1.4007 | 0.174 |
| opioid+alcohol use disorder – control | -1.20862 | 0.9160 | -1.3195 | 0.199 |
| Age Cateogry: | | | | |
| 50-64 – 30-49 | 0.08352 | 0.8253 | 0.1012 | 0.920 |
| 50-65 – 30-49 | -0.33523 | 1.2679 | -0.2644 | 0.794 |
| 65+ – 30-49 | -0.84984 | 0.9561 | -0.8888 | 0.383 |
| <30 – 30-49 | 0.12595 | 0.9996 | 0.1260 | 0.901 |
| Ethnicity: | | | | |
| Asian – White | -0.13588 | 1.6645 | -0.0816 | 0.936 |
| Black – White | -0.02270 | 0.6776 | -0.0335 | 0.974 |
| Hispanic – White | -0.99545 | 1.2960 | -0.7681 | 0.450 |
| Gender: | | | | |
| Male – Female | -0.97954 | 0.8010 | -1.2229 | 0.233 |
| PMI converted to hours | 0.00225 | 0.0332 | 0.0677 | 0.947 |
| pH | -0.72778 | 1.1490 | -0.6334 | 0.532 |

[a] Represents reference level

### Linear Regression

Model Fit Measures

| Model | R | $R^2$ |
|---|---|---|
| 1 | 0.675 | 0.455 |

Model Coefficients - RAB3GAP1

| Predictor | Estimate | SE | t | p |
|---|---|---|---|---|
| Intercept [a] | 28.1255 | 4.0388 | 6.964 | < .001 |
| Arm: | | | | |
| alcohol use disorder – control | -0.4828 | 0.4250 | -1.136 | 0.267 |
| opioid use disorder – control | 0.1562 | 0.3976 | 0.393 | 0.698 |
| opioid+alcohol use disorder – control | -0.1346 | 0.4701 | -0.286 | 0.777 |
| Age Cateogry: | | | | |
| 50-64 – 30-49 | 1.0174 | 0.4236 | 2.402 | 0.024 |
| 50-65 – 30-49 | 0.3994 | 0.6507 | 0.614 | 0.545 |
| 65+ – 30-49 | 0.8950 | 0.4907 | 1.824 | 0.081 |
| <30 – 30-49 | 0.4117 | 0.5131 | 0.802 | 0.430 |
| Ethnicity: | | | | |
| Asian – White | 0.9512 | 0.8543 | 1.113 | 0.277 |
| Black – White | 0.1535 | 0.3478 | 0.441 | 0.663 |
| Hispanic – White | -0.3856 | 0.6652 | -0.580 | 0.568 |
| Gender: | | | | |
| Male – Female | -0.1759 | 0.4111 | -0.428 | 0.673 |
| PMI converted to hours | 0.0191 | 0.0171 | 1.119 | 0.274 |
| pH | -0.7393 | 0.5897 | -1.254 | 0.222 |

[a] Represents reference level

#### Linear Regression

Model Fit Measures

| Model | R | $R^2$ |
|---|---|---|
| 1 | 0.715 | 0.511 |

Model Coefficients - HNRNPH2

| Predictor | Estimate | SE | t | p |
|---|---|---|---|---|
| Intercept [a] | 27.75093 | 4.5403 | 6.1121 | < .001 |
| Arm: | | | | |
| alcohol use disorder – control | -0.44850 | 0.4778 | -0.9388 | 0.357 |
| opioid use disorder – control | -0.34420 | 0.4469 | -0.7702 | 0.449 |
| opioid+alcohol use disorder – control | -1.00590 | 0.5285 | -1.9033 | 0.069 |
| Age Cateogry: | | | | |
| 50-64 – 30-49 | 1.12602 | 0.4762 | 2.3647 | 0.026 |
| 50-65 – 30-49 | -0.08432 | 0.7315 | -0.1153 | 0.909 |
| 65+ – 30-49 | 0.38679 | 0.5517 | 0.7011 | 0.490 |
| <30 – 30-49 | 0.01406 | 0.5768 | 0.0244 | 0.981 |
| Ethnicity: | | | | |
| Asian – White | 1.77474 | 0.9604 | 1.8479 | 0.077 |
| Black – White | 0.03285 | 0.3910 | 0.0840 | 0.934 |
| Hispanic – White | -0.18504 | 0.7478 | -0.2474 | 0.807 |
| Gender: | | | | |
| Male – Female | -0.85312 | 0.4622 | -1.8459 | 0.077 |
| PMI converted to hours | 0.00906 | 0.0192 | 0.4723 | 0.641 |
| pH | -0.54702 | 0.6629 | -0.8251 | 0.417 |

[a] Represents reference level

#### Linear Regression

Model Fit Measures

| Model | R | $R^2$ |
|---|---|---|
| 1 | 0.776 | 0.602 |

Model Coefficients - RPS11

| Predictor | Estimate | SE | t | p |
|---|---|---|---|---|
| Intercept [a] | 35.80560 | 4.8224 | 7.4249 | < .001 |
| Arm: | | | | |
| alcohol use disorder – control | -1.38388 | 0.5074 | -2.7272 | 0.012 |
| opioid use disorder – control | -0.03950 | 0.4747 | -0.0832 | 0.934 |
| opioid+alcohol use disorder – control | -0.89255 | 0.5613 | -1.5900 | 0.125 |
| Age Cateogry: | | | | |
| 50-64 – 30-49 | 0.98886 | 0.5058 | 1.9552 | 0.062 |
| 50-65 – 30-49 | 0.73679 | 0.7770 | 0.9483 | 0.352 |
| 65+ – 30-49 | 0.50963 | 0.5859 | 0.8698 | 0.393 |
| <30 – 30-49 | 1.17083 | 0.6126 | 1.9112 | 0.068 |
| Ethnicity: | | | | |
| Asian – White | 0.50328 | 1.0201 | 0.4934 | 0.626 |
| Black – White | -0.34968 | 0.4153 | -0.8420 | 0.408 |
| Hispanic – White | 0.47283 | 0.7942 | 0.5953 | 0.557 |
| Gender: | | | | |
| Male – Female | -0.73021 | 0.4909 | -1.4876 | 0.150 |
| PMI converted to hours | -0.00781 | 0.0204 | -0.3833 | 0.705 |
| pH | -1.47573 | 0.7041 | -2.0958 | 0.047 |

[a] Represents reference level

### Linear Regression

Model Fit Measures

| Model | R | $R^2$ |
|---|---|---|
| 1 | 0.657 | 0.432 |

Model Coefficients - AUH

| Predictor | Estimate | SE | t | p |
|---|---|---|---|---|
| Intercept [a] | 30.2022 | 1.01670 | 29.7061 | < .001 |
| Arm: | | | | |
| alcohol use disorder – control | 0.2371 | 0.10698 | 2.2163 | 0.036 |
| opioid use disorder – control | 0.1383 | 0.10008 | 1.3819 | 0.180 |
| opioid+alcohol use disorder – control | -0.1971 | 0.11835 | -1.6658 | 0.109 |
| Age Cateogry: | | | | |
| 50-64 – 30-49 | 0.0364 | 0.10663 | 0.3416 | 0.736 |
| 50-65 – 30-49 | -0.0198 | 0.16381 | -0.1207 | 0.905 |
| 65+ – 30-49 | -0.0151 | 0.12353 | -0.1223 | 0.904 |
| <30 – 30-49 | 0.1506 | 0.12916 | 1.1661 | 0.255 |
| Ethnicity: | | | | |
| Asian – White | -0.0511 | 0.21506 | -0.2377 | 0.814 |
| Black – White | -0.1382 | 0.08755 | -1.5784 | 0.128 |
| Hispanic – White | -0.1426 | 0.16745 | -0.8519 | 0.403 |
| Gender: | | | | |
| Male – Female | -0.1480 | 0.10349 | -1.4305 | 0.165 |
| PMI converted to hours | 1.18e-4 | 0.00429 | 0.0275 | 0.978 |
| pH | -0.2281 | 0.14845 | -1.5367 | 0.137 |

[a] Represents reference level

### Linear Regression

Model Fit Measures

| Model | R | $R^2$ |
|---|---|---|
| 1 | 0.842 | 0.708 |

Model Coefficients - PCBP2

| Predictor | Estimate | SE | t | p |
|---|---|---|---|---|
| Intercept [a] | 28.94712 | 1.12819 | 25.65796 | < .001 |
| Arm: | | | | |
| alcohol use disorder – control | -5.12e−4 | 0.11871 | -0.00431 | 0.997 |
| opioid use disorder – control | -0.26037 | 0.11105 | -2.34456 | 0.028 |
| opioid+alcohol use disorder – control | 0.28924 | 0.13133 | 2.20243 | 0.037 |
| Age Cateogry: | | | | |
| 50-64 – 30-49 | 0.27717 | 0.11832 | 2.34251 | 0.028 |
| 50-65 – 30-49 | -0.15056 | 0.18177 | -0.82826 | 0.416 |
| 65+ – 30-49 | -0.07083 | 0.13708 | -0.51669 | 0.610 |
| <30 – 30-49 | 0.39560 | 0.14332 | 2.76029 | 0.011 |
| Ethnicity: | | | | |
| Asian – White | 0.42269 | 0.23864 | 1.77119 | 0.089 |
| Black – White | 0.02115 | 0.09715 | 0.21774 | 0.829 |
| Hispanic – White | -0.04177 | 0.18581 | -0.22478 | 0.824 |
| Gender: | | | | |
| Male – Female | -0.25055 | 0.11484 | -2.18178 | 0.039 |
| PMI converted to hours | -0.00262 | 0.00477 | -0.54938 | 0.588 |
| pH | -0.12927 | 0.16473 | -0.78471 | 0.440 |

[a] Represents reference level

#### Linear Regression

Model Fit Measures

| Model | R | $R^2$ |
|---|---|---|
| 1 | 0.801 | 0.641 |

Model Coefficients - EPM2AIP1

| Predictor | Estimate | SE | t | p |
|---|---|---|---|---|
| Intercept [a] | 26.5024 | 3.2578 | 8.135 | < .001 |
| Arm: | | | | |
| alcohol use disorder – control | -0.5243 | 0.3428 | -1.529 | 0.139 |
| opioid use disorder – control | -0.2337 | 0.3207 | -0.729 | 0.473 |
| opioid+alcohol use disorder – control | 0.4831 | 0.3792 | 1.274 | 0.215 |
| Age Cateogry: | | | | |
| 50-64 – 30-49 | 0.4880 | 0.3417 | 1.428 | 0.166 |
| 50-65 – 30-49 | -0.4215 | 0.5249 | -0.803 | 0.430 |
| 65+ – 30-49 | 0.6061 | 0.3958 | 1.531 | 0.139 |
| <30 – 30-49 | 0.3548 | 0.4139 | 0.857 | 0.400 |
| Ethnicity: | | | | |
| Asian – White | 1.3750 | 0.6891 | 1.995 | 0.057 |
| Black – White | 0.1249 | 0.2805 | 0.445 | 0.660 |
| Hispanic – White | -0.5514 | 0.5366 | -1.028 | 0.314 |
| Gender: | | | | |
| Male – Female | -0.5555 | 0.3316 | -1.675 | 0.107 |
| PMI converted to hours | 0.0185 | 0.0138 | 1.345 | 0.191 |
| pH | -0.4281 | 0.4757 | -0.900 | 0.377 |

[a] Represents reference level

#### Linear Regression

Model Fit Measures

| Model | R | $R^2$ |
|---|---|---|
| 1 | 0.733 | 0.538 |

Model Coefficients - RDH11

| Predictor | Estimate | SE | t | p |
|---|---|---|---|---|
| Intercept [a] | 24.4989 | 4.2842 | 5.718 | < .001 |
| Arm: | | | | |
| alcohol use disorder – control | -0.8373 | 0.4508 | -1.857 | 0.076 |
| opioid use disorder – control | -1.0561 | 0.4217 | -2.504 | 0.019 |
| opioid+alcohol use disorder – control | -1.1732 | 0.4987 | -2.353 | 0.027 |
| Age Cateogry: | | | | |
| 50-64 – 30-49 | 0.6720 | 0.4493 | 1.496 | 0.148 |
| 50-65 – 30-49 | -0.1700 | 0.6903 | -0.246 | 0.808 |
| 65+ – 30-49 | 0.2782 | 0.5205 | 0.534 | 0.598 |
| <30 – 30-49 | 0.2305 | 0.5442 | 0.424 | 0.676 |
| Ethnicity: | | | | |
| Asian – White | 0.5008 | 0.9062 | 0.553 | 0.586 |
| Black – White | 0.2768 | 0.3689 | 0.750 | 0.460 |
| Hispanic – White | 0.8776 | 0.7056 | 1.244 | 0.226 |
| Gender: | | | | |
| Male – Female | -0.9047 | 0.4361 | -2.075 | 0.049 |
| PMI converted to hours | -0.0124 | 0.0181 | -0.688 | 0.498 |
| pH | 0.1899 | 0.6255 | 0.304 | 0.764 |

[a] Represents reference level

### Linear Regression

Model Fit Measures

| Model | R | $R^2$ |
|---|---|---|
| 1 | 0.674 | 0.455 |

Model Coefficients - CCDC124

| Predictor | Estimate | SE | t | p |
|---|---|---|---|---|
| Intercept [a] | 24.9985 | 3.2333 | 7.7315 | < .001 |
| Arm: | | | | |
| alcohol use disorder – control | -0.7182 | 0.3402 | -2.1109 | 0.045 |
| opioid use disorder – control | -0.0560 | 0.3183 | -0.1760 | 0.862 |
| opioid+alcohol use disorder – control | -1.0592 | 0.3764 | -2.8143 | 0.010 |
| Age Cateogry: | | | | |
| 50-64 – 30-49 | 0.0853 | 0.3391 | 0.2514 | 0.804 |
| 50-65 – 30-49 | -0.1019 | 0.5210 | -0.1956 | 0.847 |
| 65+ – 30-49 | 0.3257 | 0.3929 | 0.8290 | 0.415 |
| <30 – 30-49 | 0.0349 | 0.4107 | 0.0849 | 0.933 |
| Ethnicity: | | | | |
| Asian – White | -0.0297 | 0.6839 | -0.0434 | 0.966 |
| Black – White | 0.0704 | 0.2784 | 0.2530 | 0.802 |
| Hispanic – White | -0.3169 | 0.5325 | -0.5951 | 0.557 |
| Gender: | | | | |
| Male – Female | -0.2205 | 0.3291 | -0.6701 | 0.509 |
| PMI converted to hours | 0.0171 | 0.0137 | 1.2505 | 0.223 |
| pH | -0.2344 | 0.4721 | -0.4966 | 0.624 |

[a] Represents reference level

### Linear Regression

Model Fit Measures

| Model | R | $R^2$ |
|---|---|---|
| 1 | 0.739 | 0.547 |

Model Coefficients - ARHGEF2

| Predictor | Estimate | SE | t | p |
|---|---|---|---|---|
| Intercept [a] | 29.9385 | 5.6774 | 5.273 | < .001 |
| Arm: | | | | |
| alcohol use disorder – control | -0.6572 | 0.5974 | -1.100 | 0.282 |
| opioid use disorder – control | -0.6976 | 0.5588 | -1.248 | 0.224 |
| opioid+alcohol use disorder – control | 0.5808 | 0.6609 | 0.879 | 0.388 |
| Age Cateogry: | | | | |
| 50-64 – 30-49 | 1.1677 | 0.5954 | 1.961 | 0.062 |
| 50-65 – 30-49 | -0.9073 | 0.9147 | -0.992 | 0.331 |
| 65+ – 30-49 | 0.8093 | 0.6898 | 1.173 | 0.252 |
| <30 – 30-49 | 0.2488 | 0.7212 | 0.345 | 0.733 |
| Ethnicity: | | | | |
| Asian – White | 1.5591 | 1.2009 | 1.298 | 0.207 |
| Black – White | 0.7356 | 0.4889 | 1.505 | 0.145 |
| Hispanic – White | 0.8441 | 0.9351 | 0.903 | 0.376 |
| Gender: | | | | |
| Male – Female | -0.4589 | 0.5779 | -0.794 | 0.435 |
| PMI converted to hours | -0.0239 | 0.0240 | -0.996 | 0.329 |
| pH | -0.5749 | 0.8290 | -0.693 | 0.495 |

[a] Represents reference level

### Linear Regression

Model Fit Measures

| Model | R | $R^2$ |
|---|---|---|
| 1 | 0.671 | 0.450 |

Model Coefficients - IGSF8

| Predictor | Estimate | SE | t | p |
|---|---|---|---|---|
| Intercept [a] | 31.91108 | 1.10820 | 28.796 | < .001 |
| Arm: | | | | |
| alcohol use disorder – control | 0.14317 | 0.11661 | 1.228 | 0.231 |
| opioid use disorder – control | 0.16468 | 0.10908 | 1.510 | 0.144 |
| opioid+alcohol use disorder – control | -0.06149 | 0.12900 | -0.477 | 0.638 |
| Age Cateogry: | | | | |
| 50-64 – 30-49 | -0.07366 | 0.11623 | -0.634 | 0.532 |
| 50-65 – 30-49 | -0.02255 | 0.17855 | -0.126 | 0.901 |
| 65+ – 30-49 | -0.14245 | 0.13465 | -1.058 | 0.301 |
| <30 – 30-49 | -0.06228 | 0.14078 | -0.442 | 0.662 |
| Ethnicity: | | | | |
| Asian – White | -0.34840 | 0.23441 | -1.486 | 0.150 |
| Black – White | -0.14080 | 0.09543 | -1.475 | 0.153 |
| Hispanic – White | -0.40009 | 0.18252 | -2.192 | 0.038 |
| Gender: | | | | |
| Male – Female | 0.04646 | 0.11280 | 0.412 | 0.684 |
| PMI converted to hours | 0.00158 | 0.00468 | 0.338 | 0.739 |
| pH | -0.05706 | 0.16181 | -0.353 | 0.727 |

[a] Represents reference level

### Linear Regression

Model Fit Measures

| Model | R | $R^2$ |
|---|---|---|
| 1 | 0.547 | 0.299 |

Model Coefficients - WASF1

| Predictor | Estimate | SE | t | p |
|---|---|---|---|---|
| Intercept [a] | 27.05959 | 2.12109 | 12.7574 | < .001 |
| Arm: | | | | |
| alcohol use disorder – control | -0.42450 | 0.22319 | -1.9020 | 0.069 |
| opioid use disorder – control | 0.03259 | 0.20879 | 0.1561 | 0.877 |
| opioid+alcohol use disorder – control | 0.10415 | 0.24690 | 0.4218 | 0.677 |
| Age Cateogry: | | | | |
| 50-64 – 30-49 | -0.11140 | 0.22245 | -0.5008 | 0.621 |
| 50-65 – 30-49 | -0.02293 | 0.34175 | -0.0671 | 0.947 |
| 65+ – 30-49 | -0.17880 | 0.25772 | -0.6938 | 0.494 |
| <30 – 30-49 | -0.32701 | 0.26945 | -1.2136 | 0.237 |
| Ethnicity: | | | | |
| Asian – White | -0.24437 | 0.44867 | -0.5447 | 0.591 |
| Black – White | 0.03512 | 0.18266 | 0.1923 | 0.849 |
| Hispanic – White | 0.33071 | 0.34934 | 0.9467 | 0.353 |
| Gender: | | | | |
| Male – Female | -0.02317 | 0.21591 | -0.1073 | 0.915 |
| PMI converted to hours | 0.00449 | 0.00896 | 0.5015 | 0.621 |
| pH | 0.13402 | 0.30970 | 0.4327 | 0.669 |

[a] Represents reference level

### Linear Regression

Model Fit Measures

| Model | R | $R^2$ |
|---|---|---|
| 1 | 0.647 | 0.419 |

Model Coefficients - RABGGTA

| Predictor | Estimate | SE | t | p |
|---|---|---|---|---|
| Intercept [a] | 25.9140 | 4.0029 | 6.474 | < .001 |
| Arm: | | | | |
| alcohol use disorder – control | -0.4709 | 0.4212 | -1.118 | 0.275 |
| opioid use disorder – control | -0.1561 | 0.3940 | -0.396 | 0.696 |
| opioid+alcohol use disorder – control | -0.6588 | 0.4660 | -1.414 | 0.170 |
| Age Cateogry: | | | | |
| 50-64 – 30-49 | 0.4591 | 0.4198 | 1.094 | 0.285 |
| 50-65 – 30-49 | 0.8240 | 0.6449 | 1.278 | 0.214 |
| 65+ – 30-49 | -0.2160 | 0.4864 | -0.444 | 0.661 |
| <30 – 30-49 | 1.0001 | 0.5085 | 1.967 | 0.061 |
| Ethnicity: | | | | |
| Asian – White | 0.5557 | 0.8467 | 0.656 | 0.518 |
| Black – White | 0.1140 | 0.3447 | 0.331 | 0.744 |
| Hispanic – White | 0.6796 | 0.6593 | 1.031 | 0.313 |
| Gender: | | | | |
| Male – Female | -0.1685 | 0.4075 | -0.414 | 0.683 |
| PMI converted to hours | -0.0178 | 0.0169 | -1.051 | 0.304 |
| pH | -0.0689 | 0.5845 | -0.118 | 0.907 |

[a] Represents reference level

### Linear Regression

Model Fit Measures

| Model | R | $R^2$ |
|---|---|---|
| 1 | 0.773 | 0.597 |

Model Coefficients - D2HGDH

| Predictor | Estimate | SE | t | p |
|---|---|---|---|---|
| Intercept [a] | 23.6156 | 4.6864 | 5.0392 | < .001 |
| Arm: | | | | |
| alcohol use disorder – control | 0.8931 | 0.4931 | 1.8112 | 0.083 |
| opioid use disorder – control | 0.7569 | 0.4613 | 1.6408 | 0.114 |
| opioid+alcohol use disorder – control | -1.2802 | 0.5455 | -2.3467 | 0.028 |
| Age Cateogry: | | | | |
| 50-64 – 30-49 | -0.4817 | 0.4915 | -0.9801 | 0.337 |
| 50-65 – 30-49 | -0.6320 | 0.7551 | -0.8370 | 0.411 |
| 65+ – 30-49 | 0.0114 | 0.5694 | 0.0201 | 0.984 |
| <30 – 30-49 | -0.1817 | 0.5953 | -0.3053 | 0.763 |
| Ethnicity: | | | | |
| Asian – White | -0.6156 | 0.9913 | -0.6210 | 0.540 |
| Black – White | 0.0674 | 0.4036 | 0.1671 | 0.869 |
| Hispanic – White | -0.6891 | 0.7718 | -0.8929 | 0.381 |
| Gender: | | | | |
| Male – Female | -0.1569 | 0.4770 | -0.3289 | 0.745 |
| PMI converted to hours | 0.0150 | 0.0198 | 0.7565 | 0.457 |
| pH | 0.1024 | 0.6843 | 0.1497 | 0.882 |

[a] Represents reference level

### Linear Regression

Model Fit Measures

| Model | R | $R^2$ |
|---|---|---|
| 1 | 0.726 | 0.527 |

Model Coefficients - UCHL5

| Predictor | Estimate | SE | t | p |
|---|---|---|---|---|
| Intercept [a] | 21.5088 | 2.3737 | 9.0612 | < .001 |
| Arm: | | | | |
| alcohol use disorder – control | -0.6649 | 0.2498 | -2.6622 | 0.014 |
| opioid use disorder – control | -0.2443 | 0.2337 | -1.0456 | 0.306 |
| opioid+alcohol use disorder – control | -0.2498 | 0.2763 | -0.9040 | 0.375 |
| Age Cateogry: | | | | |
| 50-64 – 30-49 | 0.2198 | 0.2490 | 0.8831 | 0.386 |
| 50-65 – 30-49 | -0.0708 | 0.3825 | -0.1852 | 0.855 |
| 65+ – 30-49 | 0.3691 | 0.2884 | 1.2797 | 0.213 |
| <30 – 30-49 | -0.0156 | 0.3015 | -0.0517 | 0.959 |
| Ethnicity: | | | | |
| Asian – White | 0.1619 | 0.5021 | 0.3225 | 0.750 |
| Black – White | -0.2922 | 0.2044 | -1.4297 | 0.166 |
| Hispanic – White | -0.5789 | 0.3910 | -1.4808 | 0.152 |
| Gender: | | | | |
| Male – Female | -0.0219 | 0.2416 | -0.0906 | 0.929 |
| PMI converted to hours | 0.0153 | 0.0100 | 1.5301 | 0.139 |
| pH | 0.3205 | 0.3466 | 0.9248 | 0.364 |

[a] Represents reference level

### Linear Regression

Model Fit Measures

| Model | R | $R^2$ |
|---|---|---|
| 1 | 0.485 | 0.236 |

Model Coefficients - CAMK2A

| Predictor | Estimate | SE | t | p |
|---|---|---|---|---|
| Intercept [a] | 32.7687 | 1.73094 | 18.9311 | < .001 |
| Arm: | | | | |
| alcohol use disorder – control | -0.0987 | 0.18214 | -0.5421 | 0.593 |
| opioid use disorder – control | -0.0374 | 0.17038 | -0.2193 | 0.828 |
| opioid+alcohol use disorder – control | 0.1363 | 0.20149 | 0.6764 | 0.505 |
| Age Cateogry: | | | | |
| 50-64 – 30-49 | 0.2020 | 0.18154 | 1.1130 | 0.277 |
| 50-65 – 30-49 | 0.0510 | 0.27889 | 0.1830 | 0.856 |
| 65+ – 30-49 | 0.2013 | 0.21032 | 0.9570 | 0.348 |
| <30 – 30-49 | 0.0869 | 0.21989 | 0.3953 | 0.696 |
| Ethnicity: | | | | |
| Asian – White | 0.1148 | 0.36614 | 0.3134 | 0.757 |
| Black – White | 0.1061 | 0.14906 | 0.7116 | 0.484 |
| Hispanic – White | -0.0625 | 0.28508 | -0.2194 | 0.828 |
| Gender: | | | | |
| Male – Female | -0.0702 | 0.17619 | -0.3985 | 0.694 |
| PMI converted to hours | -6.04e−4 | 0.00731 | -0.0827 | 0.935 |
| pH | -0.1956 | 0.25274 | -0.7740 | 0.447 |

[a] Represents reference level

#### Linear Regression

Model Fit Measures

| Model | R | $R^2$ |
|---|---|---|
| 1 | 0.668 | 0.447 |

Model Coefficients - RPH3A

| Predictor | Estimate | SE | t | p |
|---|---|---|---|---|
| Intercept [a] | 30.68485 | 1.55208 | 19.7701 | < .001 |
| Arm: | | | | |
| alcohol use disorder – control | -0.09930 | 0.16332 | -0.6080 | 0.549 |
| opioid use disorder – control | -0.17014 | 0.15278 | -1.1137 | 0.276 |
| opioid+alcohol use disorder – control | 0.05151 | 0.18067 | 0.2851 | 0.778 |
| Age Cateogry: | | | | |
| 50-64 – 30-49 | 0.35382 | 0.16278 | 2.1736 | 0.040 |
| 50-65 – 30-49 | -0.01739 | 0.25007 | -0.0695 | 0.945 |
| 65+ – 30-49 | 0.23656 | 0.18858 | 1.2544 | 0.222 |
| <30 – 30-49 | 0.26660 | 0.19717 | 1.3521 | 0.189 |
| Ethnicity: | | | | |
| Asian – White | 0.16110 | 0.32831 | 0.4907 | 0.628 |
| Black – White | 0.07906 | 0.13366 | 0.5915 | 0.560 |
| Hispanic – White | 0.16123 | 0.25563 | 0.6307 | 0.534 |
| Gender: | | | | |
| Male – Female | -0.20997 | 0.15799 | -1.3290 | 0.196 |
| PMI converted to hours | -0.00958 | 0.00656 | -1.4620 | 0.157 |
| pH | -0.09344 | 0.22662 | -0.4123 | 0.684 |

[a] Represents reference level

#### Linear Regression

Model Fit Measures

| Model | R | $R^2$ |
|---|---|---|
| 1 | 0.657 | 0.432 |

Model Coefficients - PPP1R12B

| Predictor | Estimate | SE | t | p |
|---|---|---|---|---|
| Intercept [a] | 24.85061 | 3.2674 | 7.6057 | < .001 |
| Arm: | | | | |
| alcohol use disorder – control | -0.09445 | 0.3438 | -0.2747 | 0.786 |
| opioid use disorder – control | 0.80750 | 0.3216 | 2.5107 | 0.019 |
| opioid+alcohol use disorder – control | -0.41519 | 0.3803 | -1.0916 | 0.286 |
| Age Cateogry: | | | | |
| 50-64 – 30-49 | 0.15111 | 0.3427 | 0.4410 | 0.663 |
| 50-65 – 30-49 | -0.05692 | 0.5264 | -0.1081 | 0.915 |
| 65+ – 30-49 | 0.28555 | 0.3970 | 0.7193 | 0.479 |
| <30 – 30-49 | -0.23174 | 0.4151 | -0.5583 | 0.582 |
| Ethnicity: | | | | |
| Asian – White | 0.13686 | 0.6911 | 0.1980 | 0.845 |
| Black – White | -0.25483 | 0.2814 | -0.9057 | 0.374 |
| Hispanic – White | -0.50719 | 0.5381 | -0.9425 | 0.355 |
| Gender: | | | | |
| Male – Female | 0.00377 | 0.3326 | 0.0113 | 0.991 |
| PMI converted to hours | 0.00690 | 0.0138 | 0.5003 | 0.621 |
| pH | -0.26326 | 0.4771 | -0.5518 | 0.586 |

[a] Represents reference level

#### Linear Regression

Model Fit Measures

| Model | R | $R^2$ |
|---|---|---|
| 1 | 0.810 | 0.655 |

Model Coefficients - ATP6V1G2

| Predictor | Estimate | SE | t | p |
|---|---|---|---|---|
| Intercept [a] | 30.8987 | 1.73939 | 17.76411 | < .001 |
| Arm: | | | | |
| alcohol use disorder – control | 0.0761 | 0.18303 | 0.41553 | 0.681 |
| opioid use disorder – control | 0.4898 | 0.17121 | 2.86075 | 0.009 |
| opioid+alcohol use disorder – control | -0.5163 | 0.20247 | -2.55002 | 0.018 |
| Age Cateogry: | | | | |
| 50-64 – 30-49 | 0.0393 | 0.18242 | 0.21542 | 0.831 |
| 50-65 – 30-49 | 0.0227 | 0.28025 | 0.08086 | 0.936 |
| 65+ – 30-49 | -0.0444 | 0.21134 | -0.21014 | 0.835 |
| <30 – 30-49 | -0.0479 | 0.22096 | -0.21677 | 0.830 |
| Ethnicity: | | | | |
| Asian – White | -0.1624 | 0.36793 | -0.44140 | 0.663 |
| Black – White | -0.2363 | 0.14979 | -1.57770 | 0.128 |
| Hispanic – White | -0.5354 | 0.28648 | -1.86878 | 0.074 |
| Gender: | | | | |
| Male – Female | -0.0931 | 0.17705 | -0.52608 | 0.604 |
| PMI converted to hours | 0.0241 | 0.00735 | 3.27549 | 0.003 |
| pH | -3.43e−4 | 0.25397 | -0.00135 | 0.999 |

[a] Represents reference level

#### Linear Regression

Model Fit Measures

| Model | R | $R^2$ |
|---|---|---|
| 1 | 0.785 | 0.616 |

Model Coefficients - XPO1

| Predictor | Estimate | SE | t | p |
|---|---|---|---|---|
| Intercept [a] | 21.4555 | 4.6525 | 4.6116 | < .001 |
| Arm: | | | | |
| alcohol use disorder – control | 0.1117 | 0.4896 | 0.2281 | 0.821 |
| opioid use disorder – control | -0.9290 | 0.4580 | -2.0286 | 0.054 |
| opioid+alcohol use disorder – control | 0.8668 | 0.5416 | 1.6005 | 0.123 |
| Age Cateogry: | | | | |
| 50-64 – 30-49 | 0.9821 | 0.4879 | 2.0128 | 0.055 |
| 50-65 – 30-49 | -0.0178 | 0.7496 | -0.0238 | 0.981 |
| 65+ – 30-49 | 0.2388 | 0.5653 | 0.4224 | 0.676 |
| <30 – 30-49 | 0.5432 | 0.5910 | 0.9191 | 0.367 |
| Ethnicity: | | | | |
| Asian – White | 1.3973 | 0.9841 | 1.4198 | 0.169 |
| Black – White | 0.7105 | 0.4007 | 1.7732 | 0.089 |
| Hispanic – White | 1.5507 | 0.7663 | 2.0237 | 0.054 |
| Gender: | | | | |
| Male – Female | -0.2635 | 0.4736 | -0.5563 | 0.583 |
| PMI converted to hours | -0.0395 | 0.0197 | -2.0121 | 0.056 |
| pH | 0.7400 | 0.6793 | 1.0894 | 0.287 |

[a] Represents reference level

#### Linear Regression

Model Fit Measures

| Model | R | $R^2$ |
|---|---|---|
| 1 | 0.715 | 0.511 |

Model Coefficients - PRKRA

| Predictor | Estimate | SE | t | p |
|---|---|---|---|---|
| Intercept [a] | 26.80738 | 2.7565 | 9.725 | < .001 |
| Arm: | | | | |
| alcohol use disorder – control | 0.03868 | 0.2900 | 0.133 | 0.895 |
| opioid use disorder – control | -0.40950 | 0.2713 | -1.509 | 0.144 |
| opioid+alcohol use disorder – control | 0.60583 | 0.3209 | 1.888 | 0.071 |
| Age Cateogry: | | | | |
| 50-64 – 30-49 | 0.32822 | 0.2891 | 1.135 | 0.267 |
| 50-65 – 30-49 | -0.21431 | 0.4441 | -0.483 | 0.634 |
| 65+ – 30-49 | 0.05264 | 0.3349 | 0.157 | 0.876 |
| <30 – 30-49 | 0.31461 | 0.3502 | 0.898 | 0.378 |
| Ethnicity: | | | | |
| Asian – White | 0.87058 | 0.5831 | 1.493 | 0.148 |
| Black – White | 0.26131 | 0.2374 | 1.101 | 0.282 |
| Hispanic – White | 0.31253 | 0.4540 | 0.688 | 0.498 |
| Gender: | | | | |
| Male – Female | -0.11499 | 0.2806 | -0.410 | 0.686 |
| PMI converted to hours | 0.00358 | 0.0116 | 0.307 | 0.761 |
| pH | -0.10202 | 0.4025 | -0.253 | 0.802 |

[a] Represents reference level

#### Linear Regression

Model Fit Measures

| Model | R | $R^2$ |
|---|---|---|
| 1 | 0.739 | 0.545 |

Model Coefficients - GLS

| Predictor | Estimate | SE | t | p |
|---|---|---|---|---|
| Intercept [a] | 31.6550 | 3.0605 | 10.343 | < .001 |
| Arm: | | | | |
| alcohol use disorder – control | -0.1062 | 0.3220 | -0.330 | 0.744 |
| opioid use disorder – control | -0.4157 | 0.3013 | -1.380 | 0.180 |
| opioid+alcohol use disorder – control | 0.4429 | 0.3563 | 1.243 | 0.226 |
| Age Cateogry: | | | | |
| 50-64 – 30-49 | 0.6909 | 0.3210 | 2.152 | 0.042 |
| 50-65 – 30-49 | -0.2018 | 0.4931 | -0.409 | 0.686 |
| 65+ – 30-49 | 0.5333 | 0.3719 | 1.434 | 0.164 |
| <30 – 30-49 | 0.3338 | 0.3888 | 0.859 | 0.399 |
| Ethnicity: | | | | |
| Asian – White | 1.0237 | 0.6474 | 1.581 | 0.127 |
| Black – White | 0.2964 | 0.2636 | 1.125 | 0.272 |
| Hispanic – White | 0.7677 | 0.5041 | 1.523 | 0.141 |
| Gender: | | | | |
| Male – Female | -0.0818 | 0.3115 | -0.263 | 0.795 |
| PMI converted to hours | -0.0160 | 0.0129 | -1.241 | 0.227 |
| pH | -0.0565 | 0.4469 | -0.126 | 0.900 |

[a] Represents reference level

#### Linear Regression

Model Fit Measures

| Model | R | $R^2$ |
|---|---|---|
| 1 | 0.647 | 0.418 |

Model Coefficients - GAPDH

| Predictor | Estimate | SE | t | p |
|---|---|---|---|---|
| Intercept [a] | 35.85194 | 1.05823 | 33.8792 | < .001 |
| Arm: | | | | |
| alcohol use disorder – control | -0.01256 | 0.11135 | -0.1128 | 0.911 |
| opioid use disorder – control | 0.19649 | 0.10416 | 1.8863 | 0.071 |
| opioid+alcohol use disorder – control | -0.15419 | 0.12318 | -1.2517 | 0.223 |
| Age Cateogry: | | | | |
| 50-64 – 30-49 | -0.06700 | 0.11098 | -0.6037 | 0.552 |
| 50-65 – 30-49 | 0.02775 | 0.17050 | 0.1628 | 0.872 |
| 65+ – 30-49 | 0.02963 | 0.12858 | 0.2305 | 0.820 |
| <30 – 30-49 | -0.08560 | 0.13443 | -0.6368 | 0.530 |
| Ethnicity: | | | | |
| Asian – White | -0.12882 | 0.22385 | -0.5755 | 0.570 |
| Black – White | 0.06532 | 0.09113 | 0.7168 | 0.480 |
| Hispanic – White | 0.00853 | 0.17429 | 0.0489 | 0.961 |
| Gender: | | | | |
| Male – Female | 0.06857 | 0.10772 | 0.6366 | 0.530 |
| PMI converted to hours | 0.00222 | 0.00447 | 0.4959 | 0.624 |
| pH | 0.01516 | 0.15451 | 0.0981 | 0.923 |

[a] Represents reference level

#### Linear Regression

Model Fit Measures

| Model | R | $R^2$ |
|---|---|---|
| 1 | 0.725 | 0.525 |

Model Coefficients - AIFM1

| Predictor | Estimate | SE | t | p |
|---|---|---|---|---|
| Intercept [a] | 30.17650 | 1.08384 | 27.842 | < .001 |
| Arm: | | | | |
| alcohol use disorder – control | 0.02916 | 0.11405 | 0.256 | 0.800 |
| opioid use disorder – control | 0.11868 | 0.10669 | 1.112 | 0.277 |
| opioid+alcohol use disorder – control | -0.27948 | 0.12616 | -2.215 | 0.036 |
| Age Cateogry: | | | | |
| 50-64 – 30-49 | -0.12621 | 0.11367 | -1.110 | 0.278 |
| 50-65 – 30-49 | -0.28193 | 0.17463 | -1.614 | 0.120 |
| 65+ – 30-49 | 0.07025 | 0.13169 | 0.533 | 0.599 |
| <30 – 30-49 | -0.05390 | 0.13769 | -0.391 | 0.699 |
| Ethnicity: | | | | |
| Asian – White | -0.36153 | 0.22926 | -1.577 | 0.128 |
| Black – White | -0.05007 | 0.09334 | -0.536 | 0.597 |
| Hispanic – White | -0.13034 | 0.17851 | -0.730 | 0.472 |
| Gender: | | | | |
| Male – Female | -0.09549 | 0.11032 | -0.866 | 0.395 |
| PMI converted to hours | 0.00311 | 0.00458 | 0.679 | 0.504 |
| pH | -0.24536 | 0.15825 | -1.550 | 0.134 |

[a] Represents reference level

#### Linear Regression

Model Fit Measures

| Model | R | $R^2$ |
|---|---|---|
| 1 | 0.755 | 0.570 |

Model Coefficients - EML1

| Predictor | Estimate | SE | t | p |
|---|---|---|---|---|
| Intercept [a] | 23.6204 | 2.11188 | 11.185 | < .001 |
| Arm: | | | | |
| alcohol use disorder – control | -0.0963 | 0.22222 | -0.433 | 0.669 |
| opioid use disorder – control | 0.5585 | 0.20788 | 2.687 | 0.013 |
| opioid+alcohol use disorder – control | -0.4118 | 0.24583 | -1.675 | 0.107 |
| Age Cateogry: | | | | |
| 50-64 – 30-49 | 0.2577 | 0.22149 | 1.163 | 0.256 |
| 50-65 – 30-49 | 0.3071 | 0.34027 | 0.903 | 0.376 |
| 65+ – 30-49 | 0.3114 | 0.25660 | 1.213 | 0.237 |
| <30 – 30-49 | -0.1792 | 0.26828 | -0.668 | 0.511 |
| Ethnicity: | | | | |
| Asian – White | -0.3255 | 0.44672 | -0.729 | 0.473 |
| Black – White | -0.2355 | 0.18186 | -1.295 | 0.208 |
| Hispanic – White | -0.5296 | 0.34783 | -1.523 | 0.141 |
| Gender: | | | | |
| Male – Female | -0.1284 | 0.21497 | -0.597 | 0.556 |
| PMI converted to hours | 0.0135 | 0.00892 | 1.510 | 0.144 |
| pH | -0.0915 | 0.30836 | -0.297 | 0.769 |

[a] Represents reference level

#### Linear Regression

Model Fit Measures

| Model | R | $R^2$ |
|---|---|---|
| 1 | 0.676 | 0.456 |

Model Coefficients - DENR

| Predictor | Estimate | SE | t | p |
|---|---|---|---|---|
| Intercept [a] | 24.8713 | 2.27722 | 10.922 | < .001 |
| Arm: | | | | |
| alcohol use disorder – control | 0.0827 | 0.23962 | 0.345 | 0.733 |
| opioid use disorder – control | 0.5337 | 0.22415 | 2.381 | 0.026 |
| opioid+alcohol use disorder – control | -0.2758 | 0.26508 | -1.040 | 0.309 |
| Age Cateogry: | | | | |
| 50-64 – 30-49 | 0.0288 | 0.23883 | 0.121 | 0.905 |
| 50-65 – 30-49 | 0.1573 | 0.36691 | 0.429 | 0.672 |
| 65+ – 30-49 | 0.2460 | 0.27669 | 0.889 | 0.383 |
| <30 – 30-49 | -0.2384 | 0.28929 | -0.824 | 0.418 |
| Ethnicity: | | | | |
| Asian – White | -0.2684 | 0.48170 | -0.557 | 0.583 |
| Black – White | -0.0812 | 0.19610 | -0.414 | 0.683 |
| Hispanic – White | -0.2307 | 0.37506 | -0.615 | 0.544 |
| Gender: | | | | |
| Male – Female | 0.0613 | 0.23180 | 0.264 | 0.794 |
| PMI converted to hours | 0.0206 | 0.00962 | 2.144 | 0.042 |
| pH | 0.1826 | 0.33250 | 0.549 | 0.588 |

[a] Represents reference level

#### Linear Regression

Model Fit Measures

| Model | R | $R^2$ |
|---|---|---|
| 1 | 0.628 | 0.394 |

Model Coefficients - CDS2

| Predictor | Estimate | SE | t | p |
|---|---|---|---|---|
| Intercept [a] | 27.8186 | 2.3810 | 11.6837 | < .001 |
| Arm: | | | | |
| alcohol use disorder – control | -0.2862 | 0.2505 | -1.1423 | 0.265 |
| opioid use disorder – control | -0.4891 | 0.2344 | -2.0867 | 0.048 |
| opioid+alcohol use disorder – control | 0.4132 | 0.2772 | 1.4909 | 0.149 |
| Age Cateogry: | | | | |
| 50-64 – 30-49 | -0.3262 | 0.2497 | -1.3061 | 0.204 |
| 50-65 – 30-49 | -0.0223 | 0.3836 | -0.0582 | 0.954 |
| 65+ – 30-49 | -0.1365 | 0.2893 | -0.4717 | 0.641 |
| <30 – 30-49 | -0.2065 | 0.3025 | -0.6826 | 0.501 |
| Ethnicity: | | | | |
| Asian – White | -0.2466 | 0.5036 | -0.4897 | 0.629 |
| Black – White | 0.1040 | 0.2050 | 0.5074 | 0.617 |
| Hispanic – White | 0.2021 | 0.3921 | 0.5153 | 0.611 |
| Gender: | | | | |
| Male – Female | 0.1213 | 0.2424 | 0.5007 | 0.621 |
| PMI converted to hours | -3.57e−4 | 0.0101 | -0.0355 | 0.972 |
| pH | -0.0168 | 0.3476 | -0.0484 | 0.962 |

[a] Represents reference level

#### Linear Regression

Model Fit Measures

| Model | R | $R^2$ |
|---|---|---|
| 1 | 0.579 | 0.335 |

Model Coefficients - MYL12A

| Predictor | Estimate | SE | t | p |
|---|---|---|---|---|
| Intercept [a] | 24.5281 | 7.2911 | 3.364 | 0.003 |
| Arm: | | | | |
| alcohol use disorder – control | -0.6180 | 0.7672 | -0.806 | 0.428 |
| opioid use disorder – control | -0.4654 | 0.7177 | -0.648 | 0.523 |
| opioid+alcohol use disorder – control | 1.2112 | 0.8487 | 1.427 | 0.166 |
| Age Cateogry: | | | | |
| 50-64 – 30-49 | -0.2824 | 0.7647 | -0.369 | 0.715 |
| 50-65 – 30-49 | 1.0778 | 1.1747 | 0.917 | 0.368 |
| 65+ – 30-49 | 0.9477 | 0.8859 | 1.070 | 0.295 |
| <30 – 30-49 | -0.3268 | 0.9262 | -0.353 | 0.727 |
| Ethnicity: | | | | |
| Asian – White | 0.9540 | 1.5423 | 0.619 | 0.542 |
| Black – White | 1.2352 | 0.6279 | 1.967 | 0.061 |
| Hispanic – White | 2.2517 | 1.2008 | 1.875 | 0.073 |
| Gender: | | | | |
| Male – Female | 0.3573 | 0.7422 | 0.481 | 0.635 |
| PMI converted to hours | -0.0345 | 0.0308 | -1.122 | 0.273 |
| pH | 0.3415 | 1.0646 | 0.321 | 0.751 |

[a] Represents reference level

### Linear Regression

Model Fit Measures

| Model | R | $R^2$ |
|---|---|---|
| 1 | 0.651 | 0.423 |

Model Coefficients - NPTXR

| Predictor | Estimate | SE | t | p |
|---|---|---|---|---|
| Intercept [a] | 28.1059 | 4.1512 | 6.7706 | < .001 |
| Arm: | | | | |
| alcohol use disorder – control | 0.3032 | 0.4368 | 0.6942 | 0.494 |
| opioid use disorder – control | 0.9128 | 0.4086 | 2.2339 | 0.035 |
| opioid+alcohol use disorder – control | 0.3546 | 0.4832 | 0.7338 | 0.470 |
| Age Cateogry: | | | | |
| 50-64 – 30-49 | -0.1845 | 0.4354 | -0.4239 | 0.675 |
| 50-65 – 30-49 | -0.0730 | 0.6688 | -0.1091 | 0.914 |
| 65+ – 30-49 | 0.3444 | 0.5044 | 0.6827 | 0.501 |
| <30 – 30-49 | -1.0645 | 0.5273 | -2.0187 | 0.055 |
| Ethnicity: | | | | |
| Asian – White | 0.1076 | 0.8781 | 0.1226 | 0.903 |
| Black – White | 0.2145 | 0.3575 | 0.6001 | 0.554 |
| Hispanic – White | -1.0032 | 0.6837 | -1.4673 | 0.155 |
| Gender: | | | | |
| Male – Female | 0.0212 | 0.4225 | 0.0501 | 0.960 |
| PMI converted to hours | 0.0214 | 0.0175 | 1.2189 | 0.235 |
| pH | -0.4850 | 0.6061 | -0.8002 | 0.431 |

[a] Represents reference level

### Linear Regression

Model Fit Measures

| Model | R | $R^2$ |
|---|---|---|
| 1 | 0.686 | 0.470 |

Model Coefficients - DDX3X (2)

| Predictor | Estimate | SE | t | p |
|---|---|---|---|---|
| Intercept [a] | 25.8596 | 3.8402 | 6.734 | < .001 |
| Arm: | | | | |
| alcohol use disorder – control | -0.2936 | 0.4041 | -0.727 | 0.474 |
| opioid use disorder – control | -0.6210 | 0.3780 | -1.643 | 0.113 |
| opioid+alcohol use disorder – control | 0.0750 | 0.4470 | 0.168 | 0.868 |
| Age Cateogry: | | | | |
| 50-64 – 30-49 | 0.6367 | 0.4028 | 1.581 | 0.127 |
| 50-65 – 30-49 | -0.2111 | 0.6187 | -0.341 | 0.736 |
| 65+ – 30-49 | 0.7610 | 0.4666 | 1.631 | 0.116 |
| <30 – 30-49 | 0.3272 | 0.4878 | 0.671 | 0.509 |
| Ethnicity: | | | | |
| Asian – White | 1.2202 | 0.8123 | 1.502 | 0.146 |
| Black – White | 0.2651 | 0.3307 | 0.802 | 0.431 |
| Hispanic – White | 0.7548 | 0.6325 | 1.193 | 0.244 |
| Gender: | | | | |
| Male – Female | -0.3433 | 0.3909 | -0.878 | 0.388 |
| PMI converted to hours | -0.0158 | 0.0162 | -0.972 | 0.341 |
| pH | -0.1522 | 0.5607 | -0.271 | 0.788 |

[a] Represents reference level

#### Linear Regression

Model Fit Measures

| Model | R | $R^2$ |
|---|---|---|
| 1 | 0.648 | 0.420 |

Model Coefficients - OAT

| Predictor | Estimate | SE | t | p |
|---|---|---|---|---|
| Intercept [a] | 33.2877 | 3.2079 | 10.377 | < .001 |
| Arm: | | | | |
| alcohol use disorder – control | 0.2981 | 0.3375 | 0.883 | 0.386 |
| opioid use disorder – control | -0.7805 | 0.3158 | -2.472 | 0.021 |
| opioid+alcohol use disorder – control | -0.3508 | 0.3734 | -0.940 | 0.357 |
| Age Cateogry: | | | | |
| 50-64 – 30-49 | 0.4798 | 0.3364 | 1.426 | 0.167 |
| 50-65 – 30-49 | -0.6549 | 0.5169 | -1.267 | 0.217 |
| 65+ – 30-49 | 0.0549 | 0.3898 | 0.141 | 0.889 |
| <30 – 30-49 | 0.5832 | 0.4075 | 1.431 | 0.165 |
| Ethnicity: | | | | |
| Asian – White | 0.3459 | 0.6786 | 0.510 | 0.615 |
| Black – White | 0.2131 | 0.2762 | 0.771 | 0.448 |
| Hispanic – White | -0.0572 | 0.5283 | -0.108 | 0.915 |
| Gender: | | | | |
| Male – Female | -0.6610 | 0.3265 | -2.024 | 0.054 |
| PMI converted to hours | -0.0121 | 0.0135 | -0.896 | 0.379 |
| pH | -0.8459 | 0.4684 | -1.806 | 0.083 |

[a] Represents reference level

#### Linear Regression

Model Fit Measures

| Model | R | $R^2$ |
|---|---|---|
| 1 | 0.635 | 0.403 |

Model Coefficients - SCAMP3

| Predictor | Estimate | SE | t | p |
|---|---|---|---|---|
| Intercept [a] | 21.0201 | 6.4250 | 3.272 | 0.003 |
| Arm: | | | | |
| alcohol use disorder – control | 0.5591 | 0.6761 | 0.827 | 0.416 |
| opioid use disorder – control | 0.3253 | 0.6324 | 0.514 | 0.612 |
| opioid+alcohol use disorder – control | -0.7817 | 0.7479 | -1.045 | 0.306 |
| Age Cateogry: | | | | |
| 50-64 – 30-49 | -0.1940 | 0.6738 | -0.288 | 0.776 |
| 50-65 – 30-49 | -0.1222 | 1.0352 | -0.118 | 0.907 |
| 65+ – 30-49 | -1.6814 | 0.7807 | -2.154 | 0.042 |
| <30 – 30-49 | 0.1077 | 0.8162 | 0.132 | 0.896 |
| Ethnicity: | | | | |
| Asian – White | -2.0988 | 1.3591 | -1.544 | 0.136 |
| Black – White | -0.1720 | 0.5533 | -0.311 | 0.759 |
| Hispanic – White | 0.3128 | 1.0582 | 0.296 | 0.770 |
| Gender: | | | | |
| Male – Female | -0.5478 | 0.6540 | -0.838 | 0.410 |
| PMI converted to hours | 0.0164 | 0.0271 | 0.603 | 0.552 |
| pH | 0.6774 | 0.9381 | 0.722 | 0.477 |

[a] Represents reference level

#### Linear Regression

Model Fit Measures

| Model | R | $R^2$ |
|---|---|---|
| 1 | 0.672 | 0.451 |

Model Coefficients - PRPSAP2

| Predictor | Estimate | SE | t | p |
|---|---|---|---|---|
| Intercept [a] | 27.40457 | 1.19005 | 23.028 | < .001 |
| Arm: | | | | |
| alcohol use disorder – control | 0.13508 | 0.12522 | 1.079 | 0.291 |
| opioid use disorder – control | 0.22190 | 0.11714 | 1.894 | 0.070 |
| opioid+alcohol use disorder – control | -0.11962 | 0.13853 | -0.864 | 0.396 |
| Age Cateogry: | | | | |
| 50-64 – 30-49 | -0.02843 | 0.12481 | -0.228 | 0.822 |
| 50-65 – 30-49 | 0.05699 | 0.19174 | 0.297 | 0.769 |
| 65+ – 30-49 | -0.05945 | 0.14460 | -0.411 | 0.685 |
| <30 – 30-49 | -0.22191 | 0.15118 | -1.468 | 0.155 |
| Ethnicity: | | | | |
| Asian – White | -0.14323 | 0.25173 | -0.569 | 0.575 |
| Black – White | 0.03298 | 0.10248 | 0.322 | 0.750 |
| Hispanic – White | -0.34042 | 0.19600 | -1.737 | 0.095 |
| Gender: | | | | |
| Male – Female | 0.06090 | 0.12114 | 0.503 | 0.620 |
| PMI converted to hours | 0.00865 | 0.00503 | 1.720 | 0.098 |
| pH | -0.14793 | 0.17376 | -0.851 | 0.403 |

[a] Represents reference level

#### Linear Regression

Model Fit Measures

| Model | R | $R^2$ |
|---|---|---|
| 1 | 0.595 | 0.354 |

Model Coefficients - RASAL1

| Predictor | Estimate | SE | t | p |
|---|---|---|---|---|
| Intercept [a] | 24.21838 | 6.4353 | 3.7634 | < .001 |
| Arm: | | | | |
| alcohol use disorder – control | 1.10068 | 0.6771 | 1.6255 | 0.117 |
| opioid use disorder – control | 1.43333 | 0.6334 | 2.2628 | 0.033 |
| opioid+alcohol use disorder – control | 1.10696 | 0.7491 | 1.4777 | 0.152 |
| Age Cateogry: | | | | |
| 50-64 – 30-49 | -0.29253 | 0.6749 | -0.4334 | 0.669 |
| 50-65 – 30-49 | -1.11510 | 1.0369 | -1.0755 | 0.293 |
| 65+ – 30-49 | -0.38209 | 0.7819 | -0.4887 | 0.630 |
| <30 – 30-49 | -0.01283 | 0.8175 | -0.0157 | 0.988 |
| Ethnicity: | | | | |
| Asian – White | 1.31775 | 1.3612 | 0.9680 | 0.343 |
| Black – White | 0.24234 | 0.5542 | 0.4373 | 0.666 |
| Hispanic – White | -0.26525 | 1.0599 | -0.2503 | 0.805 |
| Gender: | | | | |
| Male – Female | 0.48306 | 0.6550 | 0.7374 | 0.468 |
| PMI converted to hours | -0.00334 | 0.0272 | -0.1228 | 0.903 |
| pH | 0.03034 | 0.9396 | 0.0323 | 0.975 |

[a] Represents reference level

### Linear Regression

Model Fit Measures

| Model | R | $R^2$ |
|---|---|---|
| 1 | 0.767 | 0.589 |

Model Coefficients - HSPA12A (2)

| Predictor | Estimate | SE | t | p |
|---|---|---|---|---|
| Intercept [a] | 29.96345 | 1.56526 | 19.1428 | < .001 |
| Arm: | | | | |
| alcohol use disorder – control | -0.20744 | 0.16470 | -1.2594 | 0.220 |
| opioid use disorder – control | -0.40841 | 0.15407 | -2.6508 | 0.014 |
| opioid+alcohol use disorder – control | 0.09654 | 0.18220 | 0.5299 | 0.601 |
| Age Cateogry: | | | | |
| 50-64 – 30-49 | 0.31293 | 0.16416 | 1.9062 | 0.069 |
| 50-65 – 30-49 | 0.04877 | 0.25220 | 0.1934 | 0.848 |
| 65+ – 30-49 | 0.15284 | 0.19019 | 0.8036 | 0.429 |
| <30 – 30-49 | 0.16547 | 0.19884 | 0.8321 | 0.414 |
| Ethnicity: | | | | |
| Asian – White | 0.21610 | 0.33110 | 0.6527 | 0.520 |
| Black – White | -0.00703 | 0.13479 | -0.0522 | 0.959 |
| Hispanic – White | 0.29992 | 0.25780 | 1.1634 | 0.256 |
| Gender: | | | | |
| Male – Female | -0.18154 | 0.15933 | -1.1394 | 0.266 |
| PMI converted to hours | -0.00811 | 0.00661 | -1.2274 | 0.232 |
| pH | 0.04630 | 0.22855 | 0.2026 | 0.841 |

[a] Represents reference level

### Linear Regression

Model Fit Measures

| Model | R | $R^2$ |
|---|---|---|
| 1 | 0.788 | 0.621 |

Model Coefficients - PPM1B

| Predictor | Estimate | SE | t | p |
|---|---|---|---|---|
| Intercept [a] | 20.0282 | 4.0674 | 4.9240 | < .001 |
| Arm: | | | | |
| alcohol use disorder – control | 0.0341 | 0.4280 | 0.0797 | 0.937 |
| opioid use disorder – control | 0.7254 | 0.4004 | 1.8118 | 0.083 |
| opioid+alcohol use disorder – control | -0.9335 | 0.4735 | -1.9717 | 0.060 |
| Age Cateogry: | | | | |
| 50-64 – 30-49 | -0.7917 | 0.4266 | -1.8558 | 0.076 |
| 50-65 – 30-49 | -0.7589 | 0.6553 | -1.1579 | 0.258 |
| 65+ – 30-49 | -0.6098 | 0.4942 | -1.2338 | 0.229 |
| <30 – 30-49 | -0.9096 | 0.5167 | -1.7604 | 0.091 |
| Ethnicity: | | | | |
| Asian – White | -1.5881 | 0.8604 | -1.8459 | 0.077 |
| Black – White | -0.0248 | 0.3503 | -0.0709 | 0.944 |
| Hispanic – White | 0.4561 | 0.6699 | 0.6808 | 0.503 |
| Gender: | | | | |
| Male – Female | 0.1891 | 0.4140 | 0.4567 | 0.652 |
| PMI converted to hours | 0.0186 | 0.0172 | 1.0821 | 0.290 |
| pH | 0.6297 | 0.5939 | 1.0604 | 0.300 |

[a] Represents reference level

#### Linear Regression

Model Fit Measures

| Model | R | $R^2$ |
|---|---|---|
| 1 | 0.768 | 0.590 |

Model Coefficients - PGK1

| Predictor | Estimate | SE | t | p |
|---|---|---|---|---|
| Intercept [a] | 33.49055 | 0.83018 | 40.342 | < .001 |
| Arm: | | | | |
| alcohol use disorder – control | 0.06059 | 0.08735 | 0.694 | 0.495 |
| opioid use disorder – control | 0.27519 | 0.08172 | 3.368 | 0.003 |
| opioid+alcohol use disorder – control | -0.13683 | 0.09664 | -1.416 | 0.170 |
| Age Cateogry: | | | | |
| 50-64 – 30-49 | 0.01488 | 0.08707 | 0.171 | 0.866 |
| 50-65 – 30-49 | 0.10119 | 0.13376 | 0.757 | 0.457 |
| 65+ – 30-49 | 0.09745 | 0.10087 | 0.966 | 0.344 |
| <30 – 30-49 | -0.07157 | 0.10546 | -0.679 | 0.504 |
| Ethnicity: | | | | |
| Asian – White | -0.07695 | 0.17561 | -0.438 | 0.665 |
| Black – White | -0.02717 | 0.07149 | -0.380 | 0.707 |
| Hispanic – White | -0.20440 | 0.13673 | -1.495 | 0.148 |
| Gender: | | | | |
| Male – Female | 0.07322 | 0.08450 | 0.866 | 0.395 |
| PMI converted to hours | 0.00824 | 0.00351 | 2.351 | 0.027 |
| pH | 0.02552 | 0.12122 | 0.210 | 0.835 |

[a] Represents reference level

#### Linear Regression

Model Fit Measures

| Model | R | $R^2$ |
|---|---|---|
| 1 | 0.748 | 0.559 |

Model Coefficients - ATP6V1B2

| Predictor | Estimate | SE | t | p |
|---|---|---|---|---|
| Intercept [a] | 33.04627 | 0.79811 | 41.406 | < .001 |
| Arm: | | | | |
| alcohol use disorder – control | 0.05905 | 0.08398 | 0.703 | 0.489 |
| opioid use disorder – control | 0.11041 | 0.07856 | 1.405 | 0.173 |
| opioid+alcohol use disorder – control | -0.25686 | 0.09290 | -2.765 | 0.011 |
| Age Cateogry: | | | | |
| 50-64 – 30-49 | -0.02104 | 0.08370 | -0.251 | 0.804 |
| 50-65 – 30-49 | -0.04240 | 0.12859 | -0.330 | 0.744 |
| 65+ – 30-49 | -0.09069 | 0.09697 | -0.935 | 0.359 |
| <30 – 30-49 | -0.03593 | 0.10139 | -0.354 | 0.726 |
| Ethnicity: | | | | |
| Asian – White | -0.23785 | 0.16882 | -1.409 | 0.172 |
| Black – White | -0.12785 | 0.06873 | -1.860 | 0.075 |
| Hispanic – White | -0.26474 | 0.13145 | -2.014 | 0.055 |
| Gender: | | | | |
| Male – Female | 0.00988 | 0.08124 | 0.122 | 0.904 |
| PMI converted to hours | 0.00507 | 0.00337 | 1.505 | 0.145 |
| pH | -0.04408 | 0.11653 | -0.378 | 0.709 |

[a] Represents reference level

### Linear Regression

Model Fit Measures

| Model | R | $R^2$ |
|---|---|---|
| 1 | 0.628 | 0.394 |

Model Coefficients - RPS12

| Predictor | Estimate | SE | t | p |
|---|---|---|---|---|
| Intercept [a] | 22.0826 | 4.0088 | 5.5085 | < .001 |
| Arm: | | | | |
| alcohol use disorder – control | 0.3781 | 0.4218 | 0.8964 | 0.379 |
| opioid use disorder – control | 0.8970 | 0.3946 | 2.2731 | 0.032 |
| opioid+alcohol use disorder – control | 0.3847 | 0.4666 | 0.8244 | 0.418 |
| Age Cateogry: | | | | |
| 50-64 – 30-49 | -0.2128 | 0.4204 | -0.5061 | 0.617 |
| 50-65 – 30-49 | -0.1669 | 0.6459 | -0.2584 | 0.798 |
| 65+ – 30-49 | -0.6949 | 0.4871 | -1.4267 | 0.167 |
| <30 – 30-49 | -0.5578 | 0.5093 | -1.0953 | 0.284 |
| Ethnicity: | | | | |
| Asian – White | -0.0913 | 0.8480 | -0.1076 | 0.915 |
| Black – White | -0.0586 | 0.3452 | -0.1697 | 0.867 |
| Hispanic – White | 0.0430 | 0.6602 | 0.0651 | 0.949 |
| Gender: | | | | |
| Male – Female | 0.5900 | 0.4081 | 1.4460 | 0.161 |
| PMI converted to hours | 0.0185 | 0.0169 | 1.0904 | 0.286 |
| pH | 0.6796 | 0.5853 | 1.1610 | 0.257 |

[a] Represents reference level

### Linear Regression

Model Fit Measures

| Model | R | $R^2$ |
|---|---|---|
| 1 | 0.676 | 0.457 |

Model Coefficients - MT3

| Predictor | Estimate | SE | t | p |
|---|---|---|---|---|
| Intercept [a] | 18.3611 | 8.1600 | 2.2501 | 0.034 |
| Arm: | | | | |
| alcohol use disorder – control | 0.6081 | 0.8586 | 0.7082 | 0.486 |
| opioid use disorder – control | 1.9393 | 0.8032 | 2.4145 | 0.024 |
| opioid+alcohol use disorder – control | -1.0237 | 0.9499 | -1.0777 | 0.292 |
| Age Cateogry: | | | | |
| 50-64 – 30-49 | 0.3054 | 0.8558 | 0.3569 | 0.724 |
| 50-65 – 30-49 | 1.2072 | 1.3147 | 0.9182 | 0.368 |
| 65+ – 30-49 | -0.0819 | 0.9915 | -0.0826 | 0.935 |
| <30 – 30-49 | -0.6012 | 1.0366 | -0.5800 | 0.567 |
| Ethnicity: | | | | |
| Asian – White | 0.0660 | 1.7261 | 0.0382 | 0.970 |
| Black – White | -0.3240 | 0.7027 | -0.4611 | 0.649 |
| Hispanic – White | -0.4240 | 1.3439 | -0.3155 | 0.755 |
| Gender: | | | | |
| Male – Female | 0.6589 | 0.8306 | 0.7933 | 0.435 |
| PMI converted to hours | 0.0482 | 0.0345 | 1.3977 | 0.175 |
| pH | 1.1664 | 1.1915 | 0.9790 | 0.337 |

[a] Represents reference level

### Linear Regression

Model Fit Measures

| Model | R | $R^2$ |
|---|---|---|
| 1 | 0.728 | 0.530 |

Model Coefficients - UQCRH

| Predictor | Estimate | SE | t | p |
|---|---|---|---|---|
| Intercept [a] | 23.8550 | 2.8738 | 8.301 | < .001 |
| Arm: | | | | |
| alcohol use disorder – control | 0.3678 | 0.3024 | 1.216 | 0.236 |
| opioid use disorder – control | 0.5313 | 0.2829 | 1.878 | 0.073 |
| opioid+alcohol use disorder – control | -0.4343 | 0.3345 | -1.298 | 0.207 |
| Age Cateogry: | | | | |
| 50-64 – 30-49 | -0.1099 | 0.3014 | -0.365 | 0.719 |
| 50-65 – 30-49 | 0.2040 | 0.4630 | 0.441 | 0.663 |
| 65+ – 30-49 | -0.4246 | 0.3492 | -1.216 | 0.236 |
| <30 – 30-49 | -0.0424 | 0.3651 | -0.116 | 0.909 |
| Ethnicity: | | | | |
| Asian – White | -0.5823 | 0.6079 | -0.958 | 0.348 |
| Black – White | -0.3226 | 0.2475 | -1.303 | 0.205 |
| Hispanic – White | -0.4038 | 0.4733 | -0.853 | 0.402 |
| Gender: | | | | |
| Male – Female | 0.1037 | 0.2925 | 0.355 | 0.726 |
| PMI converted to hours | 0.0133 | 0.0121 | 1.092 | 0.286 |
| pH | 0.6063 | 0.4196 | 1.445 | 0.161 |

[a] Represents reference level

### Linear Regression

Model Fit Measures

| Model | R | $R^2$ |
|---|---|---|
| 1 | 0.709 | 0.502 |

Model Coefficients - GNAI3

| Predictor | Estimate | SE | t | p |
|---|---|---|---|---|
| Intercept [a] | 24.61056 | 1.35406 | 18.175 | < .001 |
| Arm: | | | | |
| alcohol use disorder – control | -0.25843 | 0.14248 | -1.814 | 0.082 |
| opioid use disorder – control | -0.43290 | 0.13328 | -3.248 | 0.003 |
| opioid+alcohol use disorder – control | -0.03673 | 0.15762 | -0.233 | 0.818 |
| Age Cateogry: | | | | |
| 50-64 – 30-49 | 0.03674 | 0.14201 | 0.259 | 0.798 |
| 50-65 – 30-49 | 0.05528 | 0.21817 | 0.253 | 0.802 |
| 65+ – 30-49 | -0.06410 | 0.16452 | -0.390 | 0.700 |
| <30 – 30-49 | 0.05389 | 0.17201 | 0.313 | 0.757 |
| Ethnicity: | | | | |
| Asian – White | -0.23050 | 0.28642 | -0.805 | 0.429 |
| Black – White | -0.04744 | 0.11661 | -0.407 | 0.688 |
| Hispanic – White | 0.07340 | 0.22301 | 0.329 | 0.745 |
| Gender: | | | | |
| Male – Female | 0.12738 | 0.13783 | 0.924 | 0.365 |
| PMI converted to hours | -0.00758 | 0.00572 | -1.325 | 0.197 |
| pH | 0.22275 | 0.19771 | 1.127 | 0.271 |

[a] Represents reference level

#### Linear Regression

Model Fit Measures

| Model | R | $R^2$ |
|---|---|---|
| 1 | 0.552 | 0.305 |

Model Coefficients - DARS

| Predictor | Estimate | SE | t | p |
|---|---|---|---|---|
| Intercept [a] | 28.51060 | 3.9121 | 7.28780 | < .001 |
| Arm: | | | | |
| alcohol use disorder – control | -0.05489 | 0.4116 | -0.13335 | 0.895 |
| opioid use disorder – control | -0.46506 | 0.3851 | -1.20769 | 0.239 |
| opioid+alcohol use disorder – control | 0.30021 | 0.4554 | 0.65924 | 0.516 |
| Age Cateogry: | | | | |
| 50-64 – 30-49 | 0.20789 | 0.4103 | 0.50670 | 0.617 |
| 50-65 – 30-49 | -0.45306 | 0.6303 | -0.71879 | 0.479 |
| 65+ – 30-49 | 0.53259 | 0.4753 | 1.12045 | 0.274 |
| <30 – 30-49 | -0.06732 | 0.4970 | -0.13545 | 0.893 |
| Ethnicity: | | | | |
| Asian – White | 0.36210 | 0.8275 | 0.43757 | 0.666 |
| Black – White | 0.17872 | 0.3369 | 0.53051 | 0.601 |
| Hispanic – White | -0.00174 | 0.6443 | -0.00270 | 0.998 |
| Gender: | | | | |
| Male – Female | 0.25197 | 0.3982 | 0.63276 | 0.533 |
| PMI converted to hours | -0.00329 | 0.0165 | -0.19887 | 0.844 |
| pH | -0.30633 | 0.5712 | -0.53628 | 0.597 |

[a] Represents reference level

#### Linear Regression

Model Fit Measures

| Model | R | $R^2$ |
|---|---|---|
| 1 | 0.657 | 0.432 |

Model Coefficients - HSPD1

| Predictor | Estimate | SE | t | p |
|---|---|---|---|---|
| Intercept [a] | 33.97424 | 0.82685 | 41.08877 | < .001 |
| Arm: | | | | |
| alcohol use disorder – control | 0.10950 | 0.08700 | 1.25860 | 0.220 |
| opioid use disorder – control | 0.15245 | 0.08139 | 1.87312 | 0.073 |
| opioid+alcohol use disorder – control | -0.09350 | 0.09625 | -0.97144 | 0.341 |
| Age Cateogry: | | | | |
| 50-64 – 30-49 | -0.01502 | 0.08672 | -0.17317 | 0.864 |
| 50-65 – 30-49 | -0.03549 | 0.13322 | -0.26641 | 0.792 |
| 65+ – 30-49 | -0.10347 | 0.10047 | -1.02986 | 0.313 |
| <30 – 30-49 | -0.03515 | 0.10504 | -0.33461 | 0.741 |
| Ethnicity: | | | | |
| Asian – White | -0.13987 | 0.17490 | -0.79973 | 0.432 |
| Black – White | -0.04618 | 0.07120 | -0.64851 | 0.523 |
| Hispanic – White | -0.26228 | 0.13618 | -1.92596 | 0.066 |
| Gender: | | | | |
| Male – Female | -2.02e−4 | 0.08417 | -0.00240 | 0.998 |
| PMI converted to hours | 0.00209 | 0.00349 | 0.59956 | 0.554 |
| pH | -0.11555 | 0.12073 | -0.95710 | 0.348 |

[a] Represents reference level

#### Linear Regression

Model Fit Measures

| Model | R | $R^2$ |
|---|---|---|
| 1 | 0.681 | 0.463 |

Model Coefficients - HSPA8

| Predictor | Estimate | SE | t | p |
|---|---|---|---|---|
| Intercept [a] | 34.05617 | 0.73923 | 46.0701 | < .001 |
| Arm: | | | | |
| alcohol use disorder – control | -0.07269 | 0.07778 | -0.9345 | 0.359 |
| opioid use disorder – control | 0.09522 | 0.07276 | 1.3086 | 0.203 |
| opioid+alcohol use disorder – control | -0.15019 | 0.08605 | -1.7454 | 0.094 |
| Age Cateogry: | | | | |
| 50-64 – 30-49 | -0.06662 | 0.07753 | -0.8593 | 0.399 |
| 50-65 – 30-49 | 0.08323 | 0.11910 | 0.6988 | 0.491 |
| 65+ – 30-49 | -0.05335 | 0.08982 | -0.5940 | 0.558 |
| <30 – 30-49 | -0.01277 | 0.09391 | -0.1360 | 0.893 |
| Ethnicity: | | | | |
| Asian – White | 0.01525 | 0.15637 | 0.0975 | 0.923 |
| Black – White | -0.06360 | 0.06366 | -0.9990 | 0.328 |
| Hispanic – White | -0.16797 | 0.12175 | -1.3796 | 0.180 |
| Gender: | | | | |
| Male – Female | 0.02369 | 0.07525 | 0.3148 | 0.756 |
| PMI converted to hours | 0.00264 | 0.00312 | 0.8448 | 0.407 |
| pH | 0.03910 | 0.10794 | 0.3622 | 0.720 |

[a] Represents reference level

#### Linear Regression

Model Fit Measures

| Model | R | $R^2$ |
|---|---|---|
| 1 | 0.656 | 0.430 |

Model Coefficients - NME1

| Predictor | Estimate | SE | t | p |
|---|---|---|---|---|
| Intercept [a] | 29.0767 | 11.2807 | 2.5776 | 0.017 |
| Arm: | | | | |
| alcohol use disorder – control | -0.2228 | 1.1870 | -0.1877 | 0.853 |
| opioid use disorder – control | -1.7008 | 1.1104 | -1.5317 | 0.139 |
| opioid+alcohol use disorder – control | 2.1239 | 1.3131 | 1.6174 | 0.119 |
| Age Cateogry: | | | | |
| 50-64 – 30-49 | -0.0451 | 1.1831 | -0.0382 | 0.970 |
| 50-65 – 30-49 | 1.6940 | 1.8175 | 0.9320 | 0.361 |
| 65+ – 30-49 | 0.3269 | 1.3707 | 0.2385 | 0.813 |
| <30 – 30-49 | 1.6995 | 1.4330 | 1.1859 | 0.247 |
| Ethnicity: | | | | |
| Asian – White | 1.7100 | 2.3862 | 0.7166 | 0.481 |
| Black – White | -0.2459 | 0.9714 | -0.2531 | 0.802 |
| Hispanic – White | 0.8521 | 1.8579 | 0.4586 | 0.651 |
| Gender: | | | | |
| Male – Female | 0.1893 | 1.1483 | 0.1648 | 0.870 |
| PMI converted to hours | -0.0449 | 0.0476 | -0.9417 | 0.356 |
| pH | -0.2770 | 1.6471 | -0.1682 | 0.868 |

[a] Represents reference level

#### Linear Regression

Model Fit Measures

| Model | R | $R^2$ |
|---|---|---|
| 1 | 0.790 | 0.625 |

Model Coefficients - HMBS

| Predictor | Estimate | SE | t | p |
|---|---|---|---|---|
| Intercept [a] | 28.71419 | 2.6623 | 10.7855 | < .001 |
| Arm: | | | | |
| alcohol use disorder – control | -0.16985 | 0.2801 | -0.6063 | 0.550 |
| opioid use disorder – control | 0.35276 | 0.2621 | 1.3461 | 0.191 |
| opioid+alcohol use disorder – control | -0.87592 | 0.3099 | -2.8265 | 0.009 |
| Age Cateogry: | | | | |
| 50-64 – 30-49 | -0.31257 | 0.2792 | -1.1194 | 0.274 |
| 50-65 – 30-49 | -0.65118 | 0.4289 | -1.5181 | 0.142 |
| 65+ – 30-49 | -0.01615 | 0.3235 | -0.0499 | 0.961 |
| <30 – 30-49 | -0.06573 | 0.3382 | -0.1943 | 0.848 |
| Ethnicity: | | | | |
| Asian – White | -1.59205 | 0.5631 | -2.8270 | 0.009 |
| Black – White | -0.22717 | 0.2293 | -0.9909 | 0.332 |
| Hispanic – White | 0.12088 | 0.4385 | 0.2757 | 0.785 |
| Gender: | | | | |
| Male – Female | -0.15997 | 0.2710 | -0.5903 | 0.561 |
| PMI converted to hours | -0.00360 | 0.0112 | -0.3199 | 0.752 |
| pH | -0.70552 | 0.3887 | -1.8150 | 0.082 |

[a] Represents reference level

#### Linear Regression

Model Fit Measures

| Model | R | $R^2$ |
|---|---|---|
| 1 | 0.648 | 0.420 |

Model Coefficients - RPL7

| Predictor | Estimate | SE | t | p |
|---|---|---|---|---|
| Intercept [a] | 29.23825 | 1.30151 | 22.4649 | < .001 |
| Arm: | | | | |
| alcohol use disorder – control | 0.06753 | 0.13695 | 0.4931 | 0.626 |
| opioid use disorder – control | 0.28844 | 0.12811 | 2.2515 | 0.034 |
| opioid+alcohol use disorder – control | -0.21745 | 0.15150 | -1.4353 | 0.164 |
| Age Cateogry: | | | | |
| 50-64 – 30-49 | 0.10632 | 0.13650 | 0.7789 | 0.444 |
| 50-65 – 30-49 | 0.08179 | 0.20970 | 0.3900 | 0.700 |
| 65+ – 30-49 | 0.08159 | 0.15814 | 0.5159 | 0.611 |
| <30 – 30-49 | -0.07947 | 0.16534 | -0.4807 | 0.635 |
| Ethnicity: | | | | |
| Asian – White | -0.00398 | 0.27531 | -0.0145 | 0.989 |
| Black – White | -0.05691 | 0.11208 | -0.5077 | 0.616 |
| Hispanic – White | -0.27462 | 0.21436 | -1.2812 | 0.212 |
| Gender: | | | | |
| Male – Female | -0.01787 | 0.13248 | -0.1349 | 0.894 |
| PMI converted to hours | 0.00734 | 0.00550 | 1.3360 | 0.194 |
| pH | -0.07507 | 0.19004 | -0.3950 | 0.696 |

[a] Represents reference level

#### Linear Regression

Model Fit Measures

| Model | R | $R^2$ |
|---|---|---|
| 1 | 0.763 | 0.581 |

Model Coefficients - JMJD7

| Predictor | Estimate | SE | t | p |
|---|---|---|---|---|
| Intercept [a] | 21.84823 | 2.27757 | 9.593 | < .001 |
| Arm: | | | | |
| alcohol use disorder – control | -0.04922 | 0.23966 | -0.205 | 0.839 |
| opioid use disorder – control | 0.38189 | 0.22419 | 1.703 | 0.101 |
| opioid+alcohol use disorder – control | -0.52915 | 0.26512 | -1.996 | 0.057 |
| Age Cateogry: | | | | |
| 50-64 – 30-49 | -0.17286 | 0.23887 | -0.724 | 0.476 |
| 50-65 – 30-49 | 0.06158 | 0.36696 | 0.168 | 0.868 |
| 65+ – 30-49 | 0.38730 | 0.27673 | 1.400 | 0.174 |
| <30 – 30-49 | -0.23575 | 0.28933 | -0.815 | 0.423 |
| Ethnicity: | | | | |
| Asian – White | -0.13395 | 0.48177 | -0.278 | 0.783 |
| Black – White | -0.23295 | 0.19613 | -1.188 | 0.247 |
| Hispanic – White | -0.82072 | 0.37511 | -2.188 | 0.039 |
| Gender: | | | | |
| Male – Female | -0.05982 | 0.23183 | -0.258 | 0.799 |
| PMI converted to hours | 0.00373 | 0.00962 | 0.388 | 0.701 |
| pH | 0.25714 | 0.33255 | 0.773 | 0.447 |

[a] Represents reference level

#### Linear Regression

Model Fit Measures

| Model | R | $R^2$ |
|---|---|---|
| 1 | 0.634 | 0.402 |

Model Coefficients - PCMT1

| Predictor | Estimate | SE | t | p |
|---|---|---|---|---|
| Intercept [a] | 32.07292 | 1.09934 | 29.1748 | < .001 |
| Arm: | | | | |
| alcohol use disorder – control | -0.04552 | 0.11568 | -0.3935 | 0.697 |
| opioid use disorder – control | 0.17908 | 0.10821 | 1.6550 | 0.111 |
| opioid+alcohol use disorder – control | -0.21213 | 0.12797 | -1.6576 | 0.110 |
| Age Cateogry: | | | | |
| 50-64 – 30-49 | -0.00562 | 0.11530 | -0.0487 | 0.962 |
| 50-65 – 30-49 | -0.09692 | 0.17713 | -0.5472 | 0.589 |
| 65+ – 30-49 | 0.06492 | 0.13357 | 0.4861 | 0.631 |
| <30 – 30-49 | -0.08024 | 0.13965 | -0.5746 | 0.571 |
| Ethnicity: | | | | |
| Asian – White | -0.13504 | 0.23254 | -0.5807 | 0.567 |
| Black – White | -0.08116 | 0.09467 | -0.8573 | 0.400 |
| Hispanic – White | -0.24614 | 0.18106 | -1.3594 | 0.187 |
| Gender: | | | | |
| Male – Female | 0.03212 | 0.11190 | 0.2870 | 0.777 |
| PMI converted to hours | 0.00656 | 0.00464 | 1.4118 | 0.171 |
| pH | -0.03674 | 0.16052 | -0.2289 | 0.821 |

[a] Represents reference level

### Linear Regression

Model Fit Measures

| Model | R | $R^2$ |
|---|---|---|
| 1 | 0.772 | 0.596 |

Model Coefficients - HIST1H2BO

| Predictor | Estimate | SE | t | p |
|---|---|---|---|---|
| Intercept [a] | 35.7376 | 5.8640 | 6.094 | < .001 |
| Arm: | | | | |
| alcohol use disorder – control | 0.4475 | 0.6170 | 0.725 | 0.475 |
| opioid use disorder – control | -1.9506 | 0.5772 | -3.379 | 0.002 |
| opioid+alcohol use disorder – control | -0.1055 | 0.6826 | -0.155 | 0.878 |
| Age Cateogry: | | | | |
| 50-64 – 30-49 | 0.4868 | 0.6150 | 0.792 | 0.436 |
| 50-65 – 30-49 | 0.4330 | 0.9448 | 0.458 | 0.651 |
| 65+ – 30-49 | 0.5169 | 0.7125 | 0.725 | 0.475 |
| <30 – 30-49 | 1.5652 | 0.7449 | 2.101 | 0.046 |
| Ethnicity: | | | | |
| Asian – White | 1.6302 | 1.2404 | 1.314 | 0.201 |
| Black – White | 0.5276 | 0.5050 | 1.045 | 0.307 |
| Hispanic – White | -0.6841 | 0.9658 | -0.708 | 0.486 |
| Gender: | | | | |
| Male – Female | -0.9006 | 0.5969 | -1.509 | 0.144 |
| PMI converted to hours | -0.0239 | 0.0248 | -0.964 | 0.345 |
| pH | -1.4624 | 0.8562 | -1.708 | 0.101 |

[a] Represents reference level

### Linear Regression

Model Fit Measures

| Model | R | $R^2$ |
|---|---|---|
| 1 | 0.695 | 0.484 |

Model Coefficients - PDGFRB

| Predictor | Estimate | SE | t | p |
|---|---|---|---|---|
| Intercept [a] | 27.58340 | 2.7391 | 10.0703 | < .001 |
| Arm: | | | | |
| alcohol use disorder – control | 0.23254 | 0.2882 | 0.8068 | 0.428 |
| opioid use disorder – control | 0.41677 | 0.2696 | 1.5458 | 0.135 |
| opioid+alcohol use disorder – control | -0.58876 | 0.3188 | -1.8466 | 0.077 |
| Age Cateogry: | | | | |
| 50-64 – 30-49 | 0.14109 | 0.2873 | 0.4912 | 0.628 |
| 50-65 – 30-49 | -0.66565 | 0.4413 | -1.5083 | 0.145 |
| 65+ – 30-49 | -0.17023 | 0.3328 | -0.5115 | 0.614 |
| <30 – 30-49 | 0.01015 | 0.3480 | 0.0292 | 0.977 |
| Ethnicity: | | | | |
| Asian – White | -0.13039 | 0.5794 | -0.2251 | 0.824 |
| Black – White | -0.31708 | 0.2359 | -1.3443 | 0.191 |
| Hispanic – White | 0.50221 | 0.4511 | 1.1132 | 0.277 |
| Gender: | | | | |
| Male – Female | 0.05149 | 0.2788 | 0.1847 | 0.855 |
| PMI converted to hours | -0.00403 | 0.0116 | -0.3483 | 0.731 |
| pH | -0.61916 | 0.3999 | -1.5481 | 0.135 |

[a] Represents reference level

### Linear Regression

Model Fit Measures

| Model | R | $R^2$ |
|---|---|---|
| 1 | 0.765 | 0.585 |

Model Coefficients - ATP2B1

| Predictor | Estimate | SE | t | p |
|---|---|---|---|---|
| Intercept [a] | 30.84853 | 0.54680 | 56.4169 | < .001 |
| Arm: | | | | |
| alcohol use disorder – control | 0.02811 | 0.05754 | 0.4885 | 0.630 |
| opioid use disorder – control | 0.12302 | 0.05382 | 2.2856 | 0.031 |
| opioid+alcohol use disorder – control | 0.14442 | 0.06365 | 2.2690 | 0.033 |
| Age Cateogry: | | | | |
| 50-64 – 30-49 | -0.12997 | 0.05735 | -2.2664 | 0.033 |
| 50-65 – 30-49 | 0.10516 | 0.08810 | 1.1937 | 0.244 |
| 65+ – 30-49 | -0.12437 | 0.06644 | -1.8719 | 0.073 |
| <30 – 30-49 | 0.04337 | 0.06946 | 0.6243 | 0.538 |
| Ethnicity: | | | | |
| Asian – White | -0.00993 | 0.11566 | -0.0859 | 0.932 |
| Black – White | 0.01000 | 0.04709 | 0.2123 | 0.834 |
| Hispanic – White | 0.01487 | 0.09006 | 0.1651 | 0.870 |
| Gender: | | | | |
| Male – Female | 0.00114 | 0.05566 | 0.0204 | 0.984 |
| PMI converted to hours | 0.00324 | 0.00231 | 1.4044 | 0.173 |
| pH | 0.02964 | 0.07984 | 0.3713 | 0.714 |

[a] Represents reference level

### Linear Regression

Model Fit Measures

| Model | R | $R^2$ |
|---|---|---|
| 1 | 0.750 | 0.562 |

Model Coefficients - ATP2B4

| Predictor | Estimate | SE | t | p |
|---|---|---|---|---|
| Intercept [a] | 32.21589 | 0.57949 | 55.59349 | < .001 |
| Arm: | | | | |
| alcohol use disorder – control | 0.04589 | 0.06098 | 0.75253 | 0.459 |
| opioid use disorder – control | 0.15645 | 0.05704 | 2.74270 | 0.011 |
| opioid+alcohol use disorder – control | 0.04919 | 0.06746 | 0.72918 | 0.473 |
| Age Cateogry: | | | | |
| 50-64 – 30-49 | -0.09347 | 0.06078 | -1.53801 | 0.137 |
| 50-65 – 30-49 | -0.05705 | 0.09337 | -0.61104 | 0.547 |
| 65+ – 30-49 | -1.28e−4 | 0.07041 | -0.00182 | 0.999 |
| <30 – 30-49 | 0.04465 | 0.07362 | 0.60658 | 0.550 |
| Ethnicity: | | | | |
| Asian – White | 0.10468 | 0.12258 | 0.85399 | 0.402 |
| Black – White | -0.07232 | 0.04990 | -1.44920 | 0.160 |
| Hispanic – White | 5.26e-4 | 0.09544 | 0.00551 | 0.996 |
| Gender: | | | | |
| Male – Female | -0.05807 | 0.05899 | -0.98443 | 0.335 |
| PMI converted to hours | 0.00279 | 0.00245 | 1.14020 | 0.265 |
| pH | -0.08471 | 0.08461 | -1.00110 | 0.327 |

[a] Represents reference level

#### Linear Regression

Model Fit Measures

| Model | R | $R^2$ |
|---|---|---|
| 1 | 0.618 | 0.382 |

Model Coefficients - PDHA1

| Predictor | Estimate | SE | t | p |
|---|---|---|---|---|
| Intercept [a] | 31.71167 | 1.34007 | 23.6642 | < .001 |
| Arm: | | | | |
| alcohol use disorder – control | -0.12165 | 0.14101 | -0.8627 | 0.397 |
| opioid use disorder – control | -0.32753 | 0.13191 | -2.4831 | 0.020 |
| opioid+alcohol use disorder – control | 0.11470 | 0.15599 | 0.7353 | 0.469 |
| Age Cateogry: | | | | |
| 50-64 – 30-49 | -0.09119 | 0.14054 | -0.6488 | 0.523 |
| 50-65 – 30-49 | 0.09390 | 0.21591 | 0.4349 | 0.668 |
| 65+ – 30-49 | -0.14991 | 0.16282 | -0.9207 | 0.366 |
| <30 – 30-49 | 0.07562 | 0.17024 | 0.4442 | 0.661 |
| Ethnicity: | | | | |
| Asian – White | -0.08560 | 0.28346 | -0.3020 | 0.765 |
| Black – White | 0.00204 | 0.11540 | 0.0177 | 0.986 |
| Hispanic – White | 0.02692 | 0.22071 | 0.1220 | 0.904 |
| Gender: | | | | |
| Male – Female | -0.03920 | 0.13641 | -0.2874 | 0.776 |
| PMI converted to hours | -0.00442 | 0.00566 | -0.7808 | 0.443 |
| pH | -0.04553 | 0.19567 | -0.2327 | 0.818 |

[a] Represents reference level

#### Linear Regression

Model Fit Measures

| Model | R | $R^2$ |
|---|---|---|
| 1 | 0.725 | 0.525 |

Model Coefficients - GNAO1

| Predictor | Estimate | SE | t | p |
|---|---|---|---|---|
| Intercept [a] | 34.49183 | 0.60353 | 57.1503 | < .001 |
| Arm: | | | | |
| alcohol use disorder – control | 0.13535 | 0.06351 | 2.1312 | 0.044 |
| opioid use disorder – control | 0.15170 | 0.05941 | 2.5535 | 0.017 |
| opioid+alcohol use disorder – control | 0.10845 | 0.07025 | 1.5437 | 0.136 |
| Age Cateogry: | | | | |
| 50-64 – 30-49 | 0.00659 | 0.06330 | 0.1041 | 0.918 |
| 50-65 – 30-49 | -0.10156 | 0.09724 | -1.0444 | 0.307 |
| 65+ – 30-49 | -0.10697 | 0.07333 | -1.4587 | 0.158 |
| <30 – 30-49 | 0.14881 | 0.07667 | 1.9409 | 0.064 |
| Ethnicity: | | | | |
| Asian – White | 0.14099 | 0.12766 | 1.1044 | 0.280 |
| Black – White | -0.04967 | 0.05197 | -0.9557 | 0.349 |
| Hispanic – White | -0.06798 | 0.09940 | -0.6839 | 0.501 |
| Gender: | | | | |
| Male – Female | -0.00783 | 0.06143 | -0.1274 | 0.900 |
| PMI converted to hours | -3.57e−4 | 0.00255 | -0.1402 | 0.890 |
| pH | -0.00247 | 0.08812 | -0.0280 | 0.978 |

[a] Represents reference level

### Linear Regression

Model Fit Measures

| Model | R | $R^2$ |
|---|---|---|
| 1 | 0.518 | 0.268 |

Model Coefficients - NEFH

| Predictor | Estimate | SE | t | p |
|---|---|---|---|---|
| Intercept [a] | 26.5780 | 7.4234 | 3.580 | 0.002 |
| Arm: | | | | |
| alcohol use disorder – control | -0.7328 | 0.7811 | -0.938 | 0.358 |
| opioid use disorder – control | -0.5221 | 0.7307 | -0.714 | 0.482 |
| opioid+alcohol use disorder – control | -1.3730 | 0.8641 | -1.589 | 0.125 |
| Age Cateogry: | | | | |
| 50-64 – 30-49 | 0.9641 | 0.7786 | 1.238 | 0.228 |
| 50-65 – 30-49 | 0.9139 | 1.1961 | 0.764 | 0.452 |
| 65+ – 30-49 | 1.0341 | 0.9020 | 1.146 | 0.263 |
| <30 – 30-49 | 0.7163 | 0.9430 | 0.760 | 0.455 |
| Ethnicity: | | | | |
| Asian – White | 0.5526 | 1.5703 | 0.352 | 0.728 |
| Black – White | -0.2339 | 0.6393 | -0.366 | 0.718 |
| Hispanic – White | 0.8121 | 1.2226 | 0.664 | 0.513 |
| Gender: | | | | |
| Male – Female | 0.1206 | 0.7556 | 0.160 | 0.875 |
| PMI converted to hours | -0.0194 | 0.0314 | -0.618 | 0.542 |
| pH | 0.1092 | 1.0839 | 0.101 | 0.921 |

[a] Represents reference level

### Linear Regression

Model Fit Measures

| Model | R | $R^2$ |
|---|---|---|
| 1 | 0.681 | 0.463 |

Model Coefficients - PC

| Predictor | Estimate | SE | t | p |
|---|---|---|---|---|
| Intercept [a] | 25.4573 | 7.7909 | 3.2676 | 0.003 |
| Arm: | | | | |
| alcohol use disorder – control | 0.0458 | 0.8198 | 0.0559 | 0.956 |
| opioid use disorder – control | -1.1332 | 0.7669 | -1.4777 | 0.152 |
| opioid+alcohol use disorder – control | -0.5215 | 0.9069 | -0.5751 | 0.571 |
| Age Cateogry: | | | | |
| 50-64 – 30-49 | 1.5393 | 0.8171 | 1.8838 | 0.072 |
| 50-65 – 30-49 | -1.5825 | 1.2553 | -1.2607 | 0.220 |
| 65+ – 30-49 | 1.6238 | 0.9466 | 1.7154 | 0.099 |
| <30 – 30-49 | 0.4452 | 0.9897 | 0.4498 | 0.657 |
| Ethnicity: | | | | |
| Asian – White | 1.3841 | 1.6480 | 0.8399 | 0.409 |
| Black – White | 0.0220 | 0.6709 | 0.0328 | 0.974 |
| Hispanic – White | 0.3212 | 1.2831 | 0.2504 | 0.804 |
| Gender: | | | | |
| Male – Female | 0.0742 | 0.7930 | 0.0936 | 0.926 |
| PMI converted to hours | -0.0157 | 0.0329 | -0.4775 | 0.637 |
| pH | 0.3263 | 1.1376 | 0.2868 | 0.777 |

[a] Represents reference level

#### Linear Regression

Model Fit Measures

| Model | R | $R^2$ |
|---|---|---|
| 1 | 0.625 | 0.391 |

Model Coefficients - PYGB

| Predictor | Estimate | SE | t | p |
|---|---|---|---|---|
| Intercept [a] | 32.40810 | 2.30532 | 14.0579 | < .001 |
| Arm: | | | | |
| alcohol use disorder – control | -0.01301 | 0.24258 | -0.0536 | 0.958 |
| opioid use disorder – control | -0.38128 | 0.22692 | -1.6802 | 0.106 |
| opioid+alcohol use disorder – control | 0.33693 | 0.26835 | 1.2555 | 0.221 |
| Age Cateogry: | | | | |
| 50-64 – 30-49 | 0.22397 | 0.24178 | 0.9263 | 0.363 |
| 50-65 – 30-49 | -0.02682 | 0.37143 | -0.0722 | 0.943 |
| 65+ – 30-49 | -0.15371 | 0.28011 | -0.5487 | 0.588 |
| <30 – 30-49 | 0.13551 | 0.29286 | 0.4627 | 0.648 |
| Ethnicity: | | | | |
| Asian – White | 0.03830 | 0.48764 | 0.0785 | 0.938 |
| Black – White | 0.11717 | 0.19852 | 0.5902 | 0.561 |
| Hispanic – White | 0.11815 | 0.37968 | 0.3112 | 0.758 |
| Gender: | | | | |
| Male – Female | 0.03521 | 0.23466 | 0.1501 | 0.882 |
| PMI converted to hours | -0.00828 | 0.00974 | -0.8505 | 0.403 |
| pH | -0.15092 | 0.33660 | -0.4484 | 0.658 |

[a] Represents reference level

#### Linear Regression

Model Fit Measures

| Model | R | $R^2$ |
|---|---|---|
| 1 | 0.741 | 0.550 |

Model Coefficients - CKB

| Predictor | Estimate | SE | t | p |
|---|---|---|---|---|
| Intercept [a] | 34.3925 | 2.8775 | 11.952 | < .001 |
| Arm: | | | | |
| alcohol use disorder – control | -0.1088 | 0.3028 | -0.359 | 0.723 |
| opioid use disorder – control | -0.4296 | 0.2832 | -1.517 | 0.142 |
| opioid+alcohol use disorder – control | 0.4583 | 0.3350 | 1.368 | 0.184 |
| Age Cateogry: | | | | |
| 50-64 – 30-49 | 0.5411 | 0.3018 | 1.793 | 0.086 |
| 50-65 – 30-49 | -0.2360 | 0.4636 | -0.509 | 0.615 |
| 65+ – 30-49 | 0.4535 | 0.3496 | 1.297 | 0.207 |
| <30 – 30-49 | 0.5024 | 0.3655 | 1.374 | 0.182 |
| Ethnicity: | | | | |
| Asian – White | 1.0670 | 0.6087 | 1.753 | 0.092 |
| Black – White | 0.1843 | 0.2478 | 0.744 | 0.464 |
| Hispanic – White | 0.7097 | 0.4739 | 1.497 | 0.147 |
| Gender: | | | | |
| Male – Female | -0.1408 | 0.2929 | -0.481 | 0.635 |
| PMI converted to hours | -0.0136 | 0.0122 | -1.123 | 0.273 |
| pH | 0.0908 | 0.4202 | 0.216 | 0.831 |

[a] Represents reference level

### Linear Regression

Model Fit Measures

| Model | R | $R^2$ |
|---|---|---|
| 1 | 0.695 | 0.483 |

Model Coefficients - UQCRB

| Predictor | Estimate | SE | t | p |
|---|---|---|---|---|
| Intercept [a] | 30.93873 | 1.27223 | 24.3185 | < .001 |
| Arm: | | | | |
| alcohol use disorder – control | 0.05525 | 0.13387 | 0.4127 | 0.684 |
| opioid use disorder – control | 0.19738 | 0.12523 | 1.5762 | 0.128 |
| opioid+alcohol use disorder – control | -0.34357 | 0.14809 | -2.3199 | 0.029 |
| Age Cateogry: | | | | |
| 50-64 – 30-49 | 0.05203 | 0.13343 | 0.3900 | 0.700 |
| 50-65 – 30-49 | 0.16856 | 0.20498 | 0.8223 | 0.419 |
| 65+ – 30-49 | 0.00412 | 0.15458 | 0.0267 | 0.979 |
| <30 – 30-49 | 0.03187 | 0.16162 | 0.1972 | 0.845 |
| Ethnicity: | | | | |
| Asian – White | -0.14057 | 0.26911 | -0.5223 | 0.606 |
| Black – White | -0.03577 | 0.10956 | -0.3265 | 0.747 |
| Hispanic – White | -0.29533 | 0.20954 | -1.4095 | 0.172 |
| Gender: | | | | |
| Male – Female | -0.12760 | 0.12950 | -0.9853 | 0.334 |
| PMI converted to hours | 0.00129 | 0.00537 | 0.2394 | 0.813 |
| pH | -0.07427 | 0.18576 | -0.3998 | 0.693 |

[a] Represents reference level

### Linear Regression

Model Fit Measures

| Model | R | $R^2$ |
|---|---|---|
| 1 | 0.697 | 0.485 |

Model Coefficients - ATP2A2

| Predictor | Estimate | SE | t | p |
|---|---|---|---|---|
| Intercept [a] | 31.85936 | 0.70768 | 45.0198 | < .001 |
| Arm: | | | | |
| alcohol use disorder – control | 0.04633 | 0.07446 | 0.6222 | 0.540 |
| opioid use disorder – control | 0.14989 | 0.06966 | 2.1518 | 0.042 |
| opioid+alcohol use disorder – control | 0.09374 | 0.08238 | 1.1379 | 0.266 |
| Age Cateogry: | | | | |
| 50-64 – 30-49 | -0.03336 | 0.07422 | -0.4495 | 0.657 |
| 50-65 – 30-49 | -0.04374 | 0.11402 | -0.3836 | 0.705 |
| 65+ – 30-49 | -0.14747 | 0.08599 | -1.7150 | 0.099 |
| <30 – 30-49 | 0.08479 | 0.08990 | 0.9431 | 0.355 |
| Ethnicity: | | | | |
| Asian – White | 0.04601 | 0.14969 | 0.3074 | 0.761 |
| Black – White | 0.00446 | 0.06094 | 0.0732 | 0.942 |
| Hispanic – White | -0.06561 | 0.11655 | -0.5629 | 0.579 |
| Gender: | | | | |
| Male – Female | -0.06931 | 0.07203 | -0.9621 | 0.346 |
| PMI converted to hours | 0.00194 | 0.00299 | 0.6499 | 0.522 |
| pH | -0.06237 | 0.10333 | -0.6036 | 0.552 |

[a] Represents reference level

#### Linear Regression

Model Fit Measures

| Model | R | $R^2$ |
|---|---|---|
| 1 | 0.792 | 0.627 |

Model Coefficients - WARS

| Predictor | Estimate | SE | t | p |
|---|---|---|---|---|
| Intercept [a] | 26.53286 | 1.68185 | 15.776 | < .001 |
| Arm: | | | | |
| alcohol use disorder – control | 0.21711 | 0.17697 | 1.227 | 0.232 |
| opioid use disorder – control | 0.65942 | 0.16555 | 3.983 | < .001 |
| opioid+alcohol use disorder – control | -0.15839 | 0.19577 | -0.809 | 0.426 |
| Age Cateogry: | | | | |
| 50-64 – 30-49 | -0.07834 | 0.17639 | -0.444 | 0.661 |
| 50-65 – 30-49 | 0.08781 | 0.27098 | 0.324 | 0.749 |
| 65+ – 30-49 | -0.04382 | 0.20435 | -0.214 | 0.832 |
| <30 – 30-49 | -0.35051 | 0.21365 | -1.641 | 0.114 |
| Ethnicity: | | | | |
| Asian – White | 0.06730 | 0.35576 | 0.189 | 0.852 |
| Black – White | -0.02635 | 0.14483 | -0.182 | 0.857 |
| Hispanic – White | -0.10482 | 0.27700 | -0.378 | 0.708 |
| Gender: | | | | |
| Male – Female | 0.15028 | 0.17120 | 0.878 | 0.389 |
| PMI converted to hours | 0.00887 | 0.00710 | 1.249 | 0.224 |
| pH | 0.24077 | 0.24557 | 0.980 | 0.337 |

[a] Represents reference level

#### Linear Regression

Model Fit Measures

| Model | R | $R^2$ |
|---|---|---|
| 1 | 0.564 | 0.318 |

Model Coefficients - DDX6

| Predictor | Estimate | SE | t | p |
|---|---|---|---|---|
| Intercept [a] | 29.4118 | 3.2257 | 9.118 | < .001 |
| Arm: | | | | |
| alcohol use disorder – control | 0.4262 | 0.3394 | 1.256 | 0.221 |
| opioid use disorder – control | 0.4906 | 0.3175 | 1.545 | 0.135 |
| opioid+alcohol use disorder – control | 0.2020 | 0.3755 | 0.538 | 0.596 |
| Age Cateogry: | | | | |
| 50-64 – 30-49 | -0.0765 | 0.3383 | -0.226 | 0.823 |
| 50-65 – 30-49 | 0.1529 | 0.5197 | 0.294 | 0.771 |
| 65+ – 30-49 | 0.1614 | 0.3919 | 0.412 | 0.684 |
| <30 – 30-49 | 0.4001 | 0.4098 | 0.976 | 0.339 |
| Ethnicity: | | | | |
| Asian – White | 0.2503 | 0.6823 | 0.367 | 0.717 |
| Black – White | 0.1595 | 0.2778 | 0.574 | 0.571 |
| Hispanic – White | 0.4884 | 0.5313 | 0.919 | 0.367 |
| Gender: | | | | |
| Male – Female | -0.2024 | 0.3283 | -0.616 | 0.543 |
| PMI converted to hours | -0.0198 | 0.0136 | -1.456 | 0.158 |
| pH | -0.3040 | 0.4710 | -0.646 | 0.525 |

[a] Represents reference level

### Linear Regression

Model Fit Measures

| Model | R | $R^2$ |
|---|---|---|
| 1 | 0.763 | 0.583 |

Model Coefficients - GPD2

| Predictor | Estimate | SE | t | p |
|---|---|---|---|---|
| Intercept [a] | 30.28073 | 0.94384 | 32.0825 | < .001 |
| Arm: | | | | |
| alcohol use disorder – control | -0.05844 | 0.09931 | -0.5885 | 0.562 |
| opioid use disorder – control | 0.15078 | 0.09291 | 1.6229 | 0.118 |
| opioid+alcohol use disorder – control | -0.23250 | 0.10987 | -2.1162 | 0.045 |
| Age Cateogry: | | | | |
| 50-64 – 30-49 | -0.04670 | 0.09899 | -0.4718 | 0.641 |
| 50-65 – 30-49 | -0.36733 | 0.15207 | -2.4155 | 0.024 |
| 65+ – 30-49 | -0.05732 | 0.11468 | -0.4998 | 0.622 |
| <30 – 30-49 | -0.15310 | 0.11990 | -1.2769 | 0.214 |
| Ethnicity: | | | | |
| Asian – White | -0.01623 | 0.19965 | -0.0813 | 0.936 |
| Black – White | 0.00657 | 0.08128 | 0.0809 | 0.936 |
| Hispanic – White | -0.36310 | 0.15545 | -2.3358 | 0.028 |
| Gender: | | | | |
| Male – Female | -0.01156 | 0.09607 | -0.1203 | 0.905 |
| PMI converted to hours | 0.00601 | 0.00399 | 1.5074 | 0.145 |
| pH | -0.07006 | 0.13781 | -0.5084 | 0.616 |

[a] Represents reference level

### Linear Regression

Model Fit Measures

| Model | R | $R^2$ |
|---|---|---|
| 1 | 0.787 | 0.619 |

Model Coefficients - GSS

| Predictor | Estimate | SE | t | p |
|---|---|---|---|---|
| Intercept [a] | 27.06002 | 1.03313 | 26.19219 | < .001 |
| Arm: | | | | |
| alcohol use disorder – control | -0.04800 | 0.10871 | -0.44153 | 0.663 |
| opioid use disorder – control | 0.17878 | 0.10169 | 1.75805 | 0.091 |
| opioid+alcohol use disorder – control | -0.25144 | 0.12026 | -2.09078 | 0.047 |
| Age Cateogry: | | | | |
| 50-64 – 30-49 | -0.18573 | 0.10835 | -1.71409 | 0.099 |
| 50-65 – 30-49 | -0.00152 | 0.16646 | -0.00911 | 0.993 |
| 65+ – 30-49 | -0.12253 | 0.12553 | -0.97611 | 0.339 |
| <30 – 30-49 | -0.00540 | 0.13124 | -0.04111 | 0.968 |
| Ethnicity: | | | | |
| Asian – White | -0.25471 | 0.21854 | -1.16553 | 0.255 |
| Black – White | -0.08355 | 0.08897 | -0.93905 | 0.357 |
| Hispanic – White | -0.17119 | 0.17016 | -1.00610 | 0.324 |
| Gender: | | | | |
| Male – Female | -0.10929 | 0.10516 | -1.03926 | 0.309 |
| PMI converted to hours | 0.00724 | 0.00436 | 1.65978 | 0.110 |
| pH | 0.15628 | 0.15085 | 1.03599 | 0.311 |

[a] Represents reference level

#### Linear Regression

Model Fit Measures

| Model | R | $R^2$ |
|---|---|---|
| 1 | 0.789 | 0.623 |

Model Coefficients - IDH3A

| Predictor | Estimate | SE | t | p |
|---|---|---|---|---|
| Intercept [a] | 32.88684 | 1.04275 | 31.5387 | < .001 |
| Arm: | | | | |
| alcohol use disorder – control | -0.27169 | 0.10972 | -2.4761 | 0.021 |
| opioid use disorder – control | -0.25195 | 0.10264 | -2.4546 | 0.022 |
| opioid+alcohol use disorder – control | 0.18237 | 0.12138 | 1.5025 | 0.146 |
| Age Cateogry: | | | | |
| 50-64 – 30-49 | -0.01042 | 0.10936 | -0.0953 | 0.925 |
| 50-65 – 30-49 | 0.07506 | 0.16801 | 0.4467 | 0.659 |
| 65+ – 30-49 | 0.02006 | 0.12670 | 0.1583 | 0.876 |
| <30 – 30-49 | 0.06532 | 0.13246 | 0.4931 | 0.626 |
| Ethnicity: | | | | |
| Asian – White | -0.15173 | 0.22057 | -0.6879 | 0.498 |
| Black – White | 0.02826 | 0.08980 | 0.3147 | 0.756 |
| Hispanic – White | -0.07026 | 0.17174 | -0.4091 | 0.686 |
| Gender: | | | | |
| Male – Female | 0.10886 | 0.10614 | 1.0256 | 0.315 |
| PMI converted to hours | -0.00612 | 0.00440 | -1.3899 | 0.177 |
| pH | -0.24699 | 0.15225 | -1.6222 | 0.118 |

[a] Represents reference level

#### Linear Regression

Model Fit Measures

| Model | R | $R^2$ |
|---|---|---|
| 1 | 0.785 | 0.617 |

Model Coefficients - DNM2

| Predictor | Estimate | SE | t | p |
|---|---|---|---|---|
| Intercept [a] | 32.7867 | 9.8132 | 3.341 | 0.003 |
| Arm: | | | | |
| alcohol use disorder – control | 0.3633 | 1.0326 | 0.352 | 0.728 |
| opioid use disorder – control | -2.1317 | 0.9659 | -2.207 | 0.037 |
| opioid+alcohol use disorder – control | 2.1742 | 1.1423 | 1.903 | 0.069 |
| Age Cateogry: | | | | |
| 50-64 – 30-49 | 1.7180 | 1.0292 | 1.669 | 0.108 |
| 50-65 – 30-49 | -1.1855 | 1.5811 | -0.750 | 0.461 |
| 65+ – 30-49 | 0.4802 | 1.1923 | 0.403 | 0.691 |
| <30 – 30-49 | 2.7632 | 1.2466 | 2.217 | 0.036 |
| Ethnicity: | | | | |
| Asian – White | 3.7266 | 2.0758 | 1.795 | 0.085 |
| Black – White | 0.6493 | 0.8451 | 0.768 | 0.450 |
| Hispanic – White | 1.3136 | 1.6162 | 0.813 | 0.424 |
| Gender: | | | | |
| Male – Female | -1.1001 | 0.9989 | -1.101 | 0.282 |
| PMI converted to hours | -0.0897 | 0.0414 | -2.165 | 0.041 |
| pH | -0.7090 | 1.4328 | -0.495 | 0.625 |

[a] Represents reference level

### Linear Regression

Model Fit Measures

| Model | R | $R^2$ |
|---|---|---|
| 1 | 0.698 | 0.487 |

Model Coefficients - PRDX6

| Predictor | Estimate | SE | t | p |
|---|---|---|---|---|
| Intercept [a] | 33.2492 | 2.13769 | 15.554 | < .001 |
| Arm: | | | | |
| alcohol use disorder – control | -0.1882 | 0.22494 | -0.837 | 0.411 |
| opioid use disorder – control | -0.4697 | 0.21042 | -2.232 | 0.035 |
| opioid+alcohol use disorder – control | 0.3350 | 0.24884 | 1.346 | 0.191 |
| Age Cateogry: | | | | |
| 50-64 – 30-49 | -0.1169 | 0.22420 | -0.521 | 0.607 |
| 50-65 – 30-49 | -0.2014 | 0.34443 | -0.585 | 0.564 |
| 65+ – 30-49 | 0.0636 | 0.25974 | 0.245 | 0.809 |
| <30 – 30-49 | 0.0489 | 0.27156 | 0.180 | 0.859 |
| Ethnicity: | | | | |
| Asian – White | 0.1059 | 0.45218 | 0.234 | 0.817 |
| Black – White | 0.0419 | 0.18409 | 0.228 | 0.822 |
| Hispanic – White | 0.1539 | 0.35208 | 0.437 | 0.666 |
| Gender: | | | | |
| Male – Female | 0.1952 | 0.21760 | 0.897 | 0.379 |
| PMI converted to hours | -0.0112 | 0.00903 | -1.241 | 0.227 |
| pH | -0.2518 | 0.31213 | -0.807 | 0.428 |

[a] Represents reference level

### Linear Regression

Model Fit Measures

| Model | R | $R^2$ |
|---|---|---|
| 1 | 0.723 | 0.522 |

Model Coefficients - PPP2R1A

| Predictor | Estimate | SE | t | p |
|---|---|---|---|---|
| Intercept [a] | 32.26490 | 1.42515 | 22.6396 | < .001 |
| Arm: | | | | |
| alcohol use disorder – control | -0.11624 | 0.14996 | -0.7752 | 0.446 |
| opioid use disorder – control | -0.30840 | 0.14028 | -2.1984 | 0.038 |
| opioid+alcohol use disorder – control | 0.30676 | 0.16589 | 1.8491 | 0.077 |
| Age Cateogry: | | | | |
| 50-64 – 30-49 | 0.01064 | 0.14947 | 0.0712 | 0.944 |
| 50-65 – 30-49 | 0.00651 | 0.22962 | 0.0283 | 0.978 |
| 65+ – 30-49 | 0.04366 | 0.17316 | 0.2521 | 0.803 |
| <30 – 30-49 | 0.19357 | 0.18104 | 1.0692 | 0.296 |
| Ethnicity: | | | | |
| Asian – White | 0.28111 | 0.30146 | 0.9325 | 0.360 |
| Black – White | 0.02589 | 0.12273 | 0.2110 | 0.835 |
| Hispanic – White | 0.15942 | 0.23472 | 0.6792 | 0.504 |
| Gender: | | | | |
| Male – Female | -0.07064 | 0.14507 | -0.4869 | 0.631 |
| PMI converted to hours | -0.00400 | 0.00602 | -0.6650 | 0.512 |
| pH | -0.08974 | 0.20809 | -0.4313 | 0.670 |

[a] Represents reference level

#### Linear Regression

Model Fit Measures

| Model | R | $R^2$ |
|---|---|---|
| 1 | 0.855 | 0.731 |

Model Coefficients - RPL3

| Predictor | Estimate | SE | t | p |
|---|---|---|---|---|
| Intercept [a] | 28.66087 | 0.88750 | 32.2938 | < .001 |
| Arm: | | | | |
| alcohol use disorder – control | 0.10251 | 0.09339 | 1.0976 | 0.283 |
| opioid use disorder – control | 0.07871 | 0.08736 | 0.9010 | 0.377 |
| opioid+alcohol use disorder – control | -0.46002 | 0.10331 | -4.4528 | < .001 |
| Age Cateogry: | | | | |
| 50-64 – 30-49 | -0.11793 | 0.09308 | -1.2670 | 0.217 |
| 50-65 – 30-49 | -0.18514 | 0.14299 | -1.2947 | 0.208 |
| 65+ – 30-49 | -0.00552 | 0.10784 | -0.0512 | 0.960 |
| <30 – 30-49 | 0.07549 | 0.11274 | 0.6696 | 0.510 |
| Ethnicity: | | | | |
| Asian – White | -0.15188 | 0.18773 | -0.8090 | 0.426 |
| Black – White | -0.04286 | 0.07643 | -0.5608 | 0.580 |
| Hispanic – White | -0.41842 | 0.14617 | -2.8625 | 0.009 |
| Gender: | | | | |
| Male – Female | -0.26125 | 0.09034 | -2.8919 | 0.008 |
| PMI converted to hours | 0.00613 | 0.00375 | 1.6361 | 0.115 |
| pH | -0.17734 | 0.12959 | -1.3685 | 0.184 |

[a] Represents reference level

#### Linear Regression

Model Fit Measures

| Model | R | $R^2$ |
|---|---|---|
| 1 | 0.597 | 0.357 |

Model Coefficients - RANBP1

| Predictor | Estimate | SE | t | p |
|---|---|---|---|---|
| Intercept [a] | 23.18715 | 5.2300 | 4.433 | < .001 |
| Arm: | | | | |
| alcohol use disorder – control | -0.44703 | 0.5503 | -0.812 | 0.425 |
| opioid use disorder – control | -0.32965 | 0.5148 | -0.640 | 0.528 |
| opioid+alcohol use disorder – control | 0.57345 | 0.6088 | 0.942 | 0.356 |
| Age Cateogry: | | | | |
| 50-64 – 30-49 | 0.54115 | 0.5485 | 0.987 | 0.334 |
| 50-65 – 30-49 | 0.64985 | 0.8427 | 0.771 | 0.448 |
| 65+ – 30-49 | 0.95515 | 0.6355 | 1.503 | 0.146 |
| <30 – 30-49 | 0.09344 | 0.6644 | 0.141 | 0.889 |
| Ethnicity: | | | | |
| Asian – White | 1.12275 | 1.1063 | 1.015 | 0.320 |
| Black – White | 0.50212 | 0.4504 | 1.115 | 0.276 |
| Hispanic – White | 0.75086 | 0.8614 | 0.872 | 0.392 |
| Gender: | | | | |
| Male – Female | -0.08220 | 0.5324 | -0.154 | 0.879 |
| PMI converted to hours | 0.00973 | 0.0221 | 0.441 | 0.664 |
| pH | 0.51347 | 0.7636 | 0.672 | 0.508 |

[a] Represents reference level

### Linear Regression

Model Fit Measures

| Model | R | $R^2$ |
|---|---|---|
| 1 | 0.641 | 0.411 |

Model Coefficients - RAP1GAP

| Predictor | Estimate | SE | t | p |
|---|---|---|---|---|
| Intercept [a] | 23.5710 | 5.6369 | 4.182 | < .001 |
| Arm: | | | | |
| alcohol use disorder – control | -0.7956 | 0.5931 | -1.341 | 0.192 |
| opioid use disorder – control | -0.7464 | 0.5549 | -1.345 | 0.191 |
| opioid+alcohol use disorder – control | 0.8396 | 0.6562 | 1.280 | 0.213 |
| Age Cateogry: | | | | |
| 50-64 – 30-49 | 0.3517 | 0.5912 | 0.595 | 0.557 |
| 50-65 – 30-49 | 0.6924 | 0.9082 | 0.762 | 0.453 |
| 65+ – 30-49 | 0.5061 | 0.6849 | 0.739 | 0.467 |
| <30 – 30-49 | 0.1639 | 0.7161 | 0.229 | 0.821 |
| Ethnicity: | | | | |
| Asian – White | 1.1987 | 1.1924 | 1.005 | 0.325 |
| Black – White | 0.7559 | 0.4854 | 1.557 | 0.133 |
| Hispanic – White | 1.1244 | 0.9284 | 1.211 | 0.238 |
| Gender: | | | | |
| Male – Female | 0.3436 | 0.5738 | 0.599 | 0.555 |
| PMI converted to hours | -0.0203 | 0.0238 | -0.851 | 0.403 |
| pH | 0.4204 | 0.8231 | 0.511 | 0.614 |

[a] Represents reference level

### Linear Regression

Model Fit Measures

| Model | R | $R^2$ |
|---|---|---|
| 1 | 0.726 | 0.527 |

Model Coefficients - NSF

| Predictor | Estimate | SE | t | p |
|---|---|---|---|---|
| Intercept [a] | 32.73778 | 1.23375 | 26.5353 | < .001 |
| Arm: | | | | |
| alcohol use disorder – control | -0.03872 | 0.12982 | -0.2983 | 0.768 |
| opioid use disorder – control | -0.28132 | 0.12144 | -2.3165 | 0.029 |
| opioid+alcohol use disorder – control | -0.01211 | 0.14361 | -0.0843 | 0.934 |
| Age Cateogry: | | | | |
| 50-64 – 30-49 | 0.26273 | 0.12939 | 2.0305 | 0.054 |
| 50-65 – 30-49 | 0.08159 | 0.19878 | 0.4104 | 0.685 |
| 65+ – 30-49 | 0.22332 | 0.14991 | 1.4898 | 0.149 |
| <30 – 30-49 | 0.31280 | 0.15673 | 1.9958 | 0.057 |
| Ethnicity: | | | | |
| Asian – White | 0.11034 | 0.26097 | 0.4228 | 0.676 |
| Black – White | 0.00576 | 0.10624 | 0.0542 | 0.957 |
| Hispanic – White | 0.24562 | 0.20320 | 1.2088 | 0.239 |
| Gender: | | | | |
| Male – Female | -0.17530 | 0.12558 | -1.3958 | 0.176 |
| PMI converted to hours | -0.00297 | 0.00521 | -0.5692 | 0.575 |
| pH | -0.10382 | 0.18014 | -0.5763 | 0.570 |

[a] Represents reference level

### Linear Regression

Model Fit Measures

| Model | R | $R^2$ |
|---|---|---|
| 1 | 0.779 | 0.607 |

Model Coefficients - AARS

| Predictor | Estimate | SE | t | p |
|---|---|---|---|---|
| Intercept [a] | 28.95878 | 1.46071 | 19.8251 | < .001 |
| Arm: | | | | |
| alcohol use disorder – control | -0.16690 | 0.15370 | -1.0858 | 0.288 |
| opioid use disorder – control | -0.31594 | 0.14378 | -2.1974 | 0.038 |
| opioid+alcohol use disorder – control | 0.04691 | 0.17003 | 0.2759 | 0.785 |
| Age Cateogry: | | | | |
| 50-64 – 30-49 | 0.21735 | 0.15320 | 1.4188 | 0.169 |
| 50-65 – 30-49 | -0.31498 | 0.23535 | -1.3383 | 0.193 |
| 65+ – 30-49 | 0.47133 | 0.17748 | 2.6557 | 0.014 |
| <30 – 30-49 | 0.04387 | 0.18556 | 0.2364 | 0.815 |
| Ethnicity: | | | | |
| Asian – White | 0.31433 | 0.30898 | 1.0173 | 0.319 |
| Black – White | 0.13474 | 0.12579 | 1.0711 | 0.295 |
| Hispanic – White | 0.43341 | 0.24058 | 1.8015 | 0.084 |
| Gender: | | | | |
| Male – Female | 0.00803 | 0.14869 | 0.0540 | 0.957 |
| PMI converted to hours | -0.00820 | 0.00617 | -1.3293 | 0.196 |
| pH | 0.11742 | 0.21328 | 0.5505 | 0.587 |

[a] Represents reference level

### Linear Regression

Model Fit Measures

| Model | R | $R^2$ |
|---|---|---|
| 1 | 0.731 | 0.534 |

Model Coefficients - PAFAH1B1

| Predictor | Estimate | SE | t | p |
|---|---|---|---|---|
| Intercept [a] | 29.73610 | 1.56237 | 19.0326 | < .001 |
| Arm: | | | | |
| alcohol use disorder – control | -0.05672 | 0.16440 | -0.3450 | 0.733 |
| opioid use disorder – control | -0.25684 | 0.15379 | -1.6701 | 0.108 |
| opioid+alcohol use disorder – control | 0.36325 | 0.18187 | 1.9973 | 0.057 |
| Age Cateogry: | | | | |
| 50-64 – 30-49 | 0.13042 | 0.16386 | 0.7960 | 0.434 |
| 50-65 – 30-49 | 0.16258 | 0.25173 | 0.6458 | 0.525 |
| 65+ – 30-49 | 0.14974 | 0.18983 | 0.7888 | 0.438 |
| <30 – 30-49 | 0.22198 | 0.19848 | 1.1184 | 0.274 |
| Ethnicity: | | | | |
| Asian – White | 0.54495 | 0.33049 | 1.6489 | 0.112 |
| Black – White | 0.07499 | 0.13454 | 0.5574 | 0.582 |
| Hispanic – White | 0.29542 | 0.25732 | 1.1481 | 0.262 |
| Gender: | | | | |
| Male – Female | 0.00545 | 0.15903 | 0.0342 | 0.973 |
| PMI converted to hours | -0.01259 | 0.00660 | -1.9083 | 0.068 |
| pH | -0.06092 | 0.22813 | -0.2670 | 0.792 |

[a] Represents reference level

#### Linear Regression

Model Fit Measures

| Model | R | $R^2$ |
|---|---|---|
| 1 | 0.833 | 0.694 |

Model Coefficients - ACSL1

| Predictor | Estimate | SE | t | p |
|---|---|---|---|---|
| Intercept [a] | 20.0907 | 2.4739 | 8.121 | < .001 |
| Arm: | | | | |
| alcohol use disorder – control | -0.1373 | 0.2603 | -0.527 | 0.603 |
| opioid use disorder – control | 0.4721 | 0.2435 | 1.939 | 0.064 |
| opioid+alcohol use disorder – control | -0.3837 | 0.2880 | -1.332 | 0.195 |
| Age Cateogry: | | | | |
| 50-64 – 30-49 | -0.3766 | 0.2595 | -1.451 | 0.160 |
| 50-65 – 30-49 | -0.2395 | 0.3986 | -0.601 | 0.554 |
| 65+ – 30-49 | 0.3498 | 0.3006 | 1.164 | 0.256 |
| <30 – 30-49 | -0.5163 | 0.3143 | -1.643 | 0.113 |
| Ethnicity: | | | | |
| Asian – White | -1.3062 | 0.5233 | -2.496 | 0.020 |
| Black – White | 0.2353 | 0.2130 | 1.104 | 0.280 |
| Hispanic – White | -0.7456 | 0.4075 | -1.830 | 0.080 |
| Gender: | | | | |
| Male – Female | 0.5556 | 0.2518 | 2.206 | 0.037 |
| PMI converted to hours | -0.0124 | 0.0104 | -1.185 | 0.248 |
| pH | 0.5899 | 0.3612 | 1.633 | 0.115 |

[a] Represents reference level

#### Linear Regression

Model Fit Measures

| Model | R | $R^2$ |
|---|---|---|
| 1 | 0.734 | 0.539 |

Model Coefficients - SHMT2

| Predictor | Estimate | SE | t | p |
|---|---|---|---|---|
| Intercept [a] | 24.7655 | 4.6613 | 5.313 | < .001 |
| Arm: | | | | |
| alcohol use disorder – control | -0.4564 | 0.4905 | -0.931 | 0.361 |
| opioid use disorder – control | -1.3707 | 0.4588 | -2.987 | 0.006 |
| opioid+alcohol use disorder – control | -1.0567 | 0.5426 | -1.948 | 0.063 |
| Age Categry: | | | | |
| 50-64 – 30-49 | 0.2770 | 0.4889 | 0.567 | 0.576 |
| 50-65 – 30-49 | -1.0108 | 0.7510 | -1.346 | 0.191 |
| 65+ – 30-49 | 0.6002 | 0.5664 | 1.060 | 0.300 |
| <30 – 30-49 | 0.1934 | 0.5921 | 0.327 | 0.747 |
| Ethnicity: | | | | |
| Asian – White | 0.9478 | 0.9860 | 0.961 | 0.346 |
| Black – White | 0.3825 | 0.4014 | 0.953 | 0.350 |
| Hispanic – White | 1.2322 | 0.7677 | 1.605 | 0.122 |
| Gender: | | | | |
| Male – Female | 0.0555 | 0.4745 | 0.117 | 0.908 |
| PMI converted to hours | -0.0229 | 0.0197 | -1.164 | 0.256 |
| pH | 0.1018 | 0.6806 | 0.150 | 0.882 |

[a] Represents reference level

#### Linear Regression

Model Fit Measures

| Model | R | $R^2$ |
|---|---|---|
| 1 | 0.782 | 0.612 |

Model Coefficients - USP5

| Predictor | Estimate | SE | t | p |
|---|---|---|---|---|
| Intercept [a] | 29.00233 | 1.80473 | 16.070 | < .001 |
| Arm: | | | | |
| alcohol use disorder – control | -0.19332 | 0.18990 | -1.018 | 0.319 |
| opioid use disorder – control | -0.41981 | 0.17765 | -2.363 | 0.027 |
| opioid+alcohol use disorder – control | -0.10507 | 0.21008 | -0.500 | 0.622 |
| Age Categry: | | | | |
| 50-64 – 30-49 | 0.44471 | 0.18928 | 2.350 | 0.027 |
| 50-65 – 30-49 | -0.35663 | 0.29078 | -1.226 | 0.232 |
| 65+ – 30-49 | 0.43585 | 0.21928 | 1.988 | 0.058 |
| <30 – 30-49 | -0.02396 | 0.22926 | -0.104 | 0.918 |
| Ethnicity: | | | | |
| Asian – White | 0.24454 | 0.38175 | 0.641 | 0.528 |
| Black – White | 0.10372 | 0.15541 | 0.667 | 0.511 |
| Hispanic – White | -0.06772 | 0.29724 | -0.228 | 0.822 |
| Gender: | | | | |
| Male – Female | -0.19283 | 0.18370 | -1.050 | 0.304 |
| PMI converted to hours | -0.00104 | 0.00762 | -0.136 | 0.893 |
| pH | -0.04923 | 0.26351 | -0.187 | 0.853 |

[a] Represents reference level

#### Linear Regression

Model Fit Measures

| Model | R | $R^2$ |
|---|---|---|
| 1 | 0.651 | 0.424 |

Model Coefficients - EIF2S3

| Predictor | Estimate | SE | t | p |
| --- | --- | --- | --- | --- |
| Intercept [a] | 26.24796 | 0.82388 | 31.8589 | < .001 |
| Arm: | | | | |
| alcohol use disorder – control | -0.01811 | 0.08669 | -0.2089 | 0.836 |
| opioid use disorder – control | 0.19950 | 0.08110 | 2.4601 | 0.021 |
| opioid+alcohol use disorder – control | -0.02602 | 0.09590 | -0.2714 | 0.788 |
| Age Cateogry: | | | | |
| 50-64 – 30-49 | -0.04738 | 0.08641 | -0.5483 | 0.589 |
| 50-65 – 30-49 | 0.17485 | 0.13274 | 1.3172 | 0.200 |
| 65+ – 30-49 | 0.03480 | 0.10011 | 0.3476 | 0.731 |
| <30 – 30-49 | 0.00387 | 0.10466 | 0.0370 | 0.971 |
| Ethnicity: | | | | |
| Asian – White | 0.14726 | 0.17427 | 0.8450 | 0.406 |
| Black – White | -0.11764 | 0.07095 | -1.6582 | 0.110 |
| Hispanic – White | 0.08714 | 0.13569 | 0.6422 | 0.527 |
| Gender: | | | | |
| Male – Female | 0.01332 | 0.08386 | 0.1588 | 0.875 |
| PMI converted to hours | 0.00365 | 0.00348 | 1.0477 | 0.305 |
| pH | 0.06927 | 0.12030 | 0.5758 | 0.570 |

[a] Represents reference level

### Linear Regression

Model Fit Measures

| Model | R | $R^2$ |
| --- | --- | --- |
| 1 | 0.607 | 0.369 |

Model Coefficients - GNA11

| Predictor | Estimate | SE | t | p |
| --- | --- | --- | --- | --- |
| Intercept [a] | 26.35725 | 3.4419 | 7.658 | < .001 |
| Arm: | | | | |
| alcohol use disorder – control | 0.13966 | 0.3622 | 0.386 | 0.703 |
| opioid use disorder – control | -0.30961 | 0.3388 | -0.914 | 0.370 |
| opioid+alcohol use disorder – control | 0.52166 | 0.4007 | 1.302 | 0.205 |
| Age Cateogry: | | | | |
| 50-64 – 30-49 | 0.23841 | 0.3610 | 0.660 | 0.515 |
| 50-65 – 30-49 | -0.37777 | 0.5546 | -0.681 | 0.502 |
| 65+ – 30-49 | 0.30912 | 0.4182 | 0.739 | 0.467 |
| <30 – 30-49 | 0.10743 | 0.4372 | 0.246 | 0.808 |
| Ethnicity: | | | | |
| Asian – White | 0.47765 | 0.7281 | 0.656 | 0.518 |
| Black – White | 0.42415 | 0.2964 | 1.431 | 0.165 |
| Hispanic – White | 0.40511 | 0.5669 | 0.715 | 0.482 |
| Gender: | | | | |
| Male – Female | 0.43775 | 0.3504 | 1.249 | 0.224 |
| PMI converted to hours | -0.00993 | 0.0145 | -0.683 | 0.501 |
| pH | -0.07167 | 0.5026 | -0.143 | 0.888 |

[a] Represents reference level

### Linear Regression

Model Fit Measures

| Model | R | $R^2$ |
| --- | --- | --- |
| 1 | 0.628 | 0.394 |

Model Coefficients - DLST

| Predictor | Estimate | SE | t | p |
|---|---|---|---|---|
| Intercept [a] | 30.21706 | 0.99274 | 30.438 | < .001 |
| Arm: | | | | |
| alcohol use disorder – control | -0.03561 | 0.10446 | -0.341 | 0.736 |
| opioid use disorder – control | -0.19659 | 0.09772 | -2.012 | 0.056 |
| opioid+alcohol use disorder – control | 0.01861 | 0.11556 | 0.161 | 0.873 |
| Age Cateogry: | | | | |
| 50-64 – 30-49 | 0.02400 | 0.10412 | 0.231 | 0.820 |
| 50-65 – 30-49 | 0.03883 | 0.15995 | 0.243 | 0.810 |
| 65+ – 30-49 | -0.02350 | 0.12062 | -0.195 | 0.847 |
| <30 – 30-49 | 0.17139 | 0.12611 | 1.359 | 0.187 |
| Ethnicity: | | | | |
| Asian – White | 0.07435 | 0.20999 | 0.354 | 0.726 |
| Black – White | -0.07039 | 0.08549 | -0.823 | 0.418 |
| Hispanic – White | 0.13143 | 0.16350 | 0.804 | 0.429 |
| Gender: | | | | |
| Male – Female | 0.04230 | 0.10105 | 0.419 | 0.679 |
| PMI converted to hours | -0.00484 | 0.00419 | -1.154 | 0.260 |
| pH | -0.04563 | 0.14495 | -0.315 | 0.756 |

[a] Represents reference level

### Linear Regression

Model Fit Measures

| Model | R | $R^2$ |
|---|---|---|
| 1 | 0.736 | 0.542 |

Model Coefficients - BSG

| Predictor | Estimate | SE | t | p |
|---|---|---|---|---|
| Intercept [a] | 29.7935 | 1.06436 | 27.99184 | < .001 |
| Arm: | | | | |
| alcohol use disorder – control | 0.0655 | 0.11200 | 0.58459 | 0.564 |
| opioid use disorder – control | 0.3206 | 0.10477 | 3.05982 | 0.005 |
| opioid+alcohol use disorder – control | 0.1727 | 0.12390 | 1.39397 | 0.176 |
| Age Cateogry: | | | | |
| 50-64 – 30-49 | -0.1217 | 0.11163 | -1.09008 | 0.287 |
| 50-65 – 30-49 | 8.14e-4 | 0.17149 | 0.00475 | 0.996 |
| 65+ – 30-49 | -0.1980 | 0.12932 | -1.53091 | 0.139 |
| <30 – 30-49 | -7.16e−4 | 0.13521 | -0.00530 | 0.996 |
| Ethnicity: | | | | |
| Asian – White | 0.0499 | 0.22514 | 0.22146 | 0.827 |
| Black – White | -0.0343 | 0.09166 | -0.37476 | 0.711 |
| Hispanic – White | -0.0649 | 0.17530 | -0.37001 | 0.715 |
| Gender: | | | | |
| Male – Female | -0.0547 | 0.10834 | -0.50491 | 0.618 |
| PMI converted to hours | 4.55e-4 | 0.00450 | 0.10127 | 0.920 |
| pH | -0.0174 | 0.15541 | -0.11199 | 0.912 |

[a] Represents reference level

### Linear Regression

Model Fit Measures

| Model | R | $R^2$ |
|---|---|---|
| 1 | 0.829 | 0.687 |

Model Coefficients - GNL1

| Predictor | Estimate | SE | t | p |
|---|---|---|---|---|
| Intercept [a] | 29.4040 | 3.5166 | 8.361 | < .001 |
| Arm: | | | | |
| alcohol use disorder – control | -0.1069 | 0.3700 | -0.289 | 0.775 |
| opioid use disorder – control | -0.5178 | 0.3462 | -1.496 | 0.148 |
| opioid+alcohol use disorder – control | 0.7902 | 0.4093 | 1.930 | 0.065 |
| Age Cateogry: | | | | |
| 50-64 – 30-49 | 1.1729 | 0.3688 | 3.180 | 0.004 |
| 50-65 – 30-49 | 0.4943 | 0.5666 | 0.872 | 0.392 |
| 65+ – 30-49 | 1.2817 | 0.4273 | 3.000 | 0.006 |
| <30 – 30-49 | 0.3121 | 0.4467 | 0.699 | 0.491 |
| Ethnicity: | | | | |
| Asian – White | 1.3943 | 0.7439 | 1.874 | 0.073 |
| Black – White | 0.5619 | 0.3028 | 1.856 | 0.076 |
| Hispanic – White | 0.6421 | 0.5792 | 1.109 | 0.279 |
| Gender: | | | | |
| Male – Female | 0.2989 | 0.3580 | 0.835 | 0.412 |
| PMI converted to hours | -0.0267 | 0.0149 | -1.797 | 0.085 |
| pH | -0.4461 | 0.5135 | -0.869 | 0.394 |

[a] Represents reference level

### Linear Regression

Model Fit Measures

| Model | R | $R^2$ |
|---|---|---|
| 1 | 0.753 | 0.568 |

Model Coefficients - TUFM

| Predictor | Estimate | SE | t | p |
|---|---|---|---|---|
| Intercept [a] | 30.7357 | 2.5844 | 11.893 | < .001 |
| Arm: | | | | |
| alcohol use disorder – control | -0.1456 | 0.2719 | -0.535 | 0.597 |
| opioid use disorder – control | -0.6099 | 0.2544 | -2.398 | 0.025 |
| opioid+alcohol use disorder – control | 0.0415 | 0.3008 | 0.138 | 0.891 |
| Age Cateogry: | | | | |
| 50-64 – 30-49 | 0.6220 | 0.2710 | 2.295 | 0.031 |
| 50-65 – 30-49 | -0.0588 | 0.4164 | -0.141 | 0.889 |
| 65+ – 30-49 | 0.5227 | 0.3140 | 1.665 | 0.109 |
| <30 – 30-49 | 0.4411 | 0.3283 | 1.344 | 0.192 |
| Ethnicity: | | | | |
| Asian – White | 0.8444 | 0.5467 | 1.545 | 0.136 |
| Black – White | 0.0936 | 0.2226 | 0.420 | 0.678 |
| Hispanic – White | 0.1980 | 0.4256 | 0.465 | 0.646 |
| Gender: | | | | |
| Male – Female | -0.2403 | 0.2631 | -0.914 | 0.370 |
| PMI converted to hours | -0.0106 | 0.0109 | -0.970 | 0.342 |
| pH | -0.0719 | 0.3773 | -0.191 | 0.850 |

[a] Represents reference level

### Linear Regression

Model Fit Measures

| Model | R | $R^2$ |
|---|---|---|
| 1 | 0.744 | 0.553 |

Model Coefficients - VCP

| Predictor | Estimate | SE | t | p |
|---|---|---|---|---|
| Intercept [a] | 31.77884 | 0.90893 | 34.9627 | < .001 |
| Arm: | | | | |
| alcohol use disorder – control | 0.11486 | 0.09564 | 1.2009 | 0.242 |
| opioid use disorder – control | 0.20468 | 0.08947 | 2.2877 | 0.031 |
| opioid+alcohol use disorder – control | -0.22183 | 0.10580 | -2.0966 | 0.047 |
| Age Cateogry: | | | | |
| 50-64 – 30-49 | -0.02206 | 0.09533 | -0.2314 | 0.819 |
| 50-65 – 30-49 | -0.04771 | 0.14645 | -0.3258 | 0.747 |
| 65+ – 30-49 | -0.03133 | 0.11044 | -0.2837 | 0.779 |
| <30 – 30-49 | -0.06818 | 0.11547 | -0.5904 | 0.560 |
| Ethnicity: | | | | |
| Asian – White | -0.07162 | 0.19227 | -0.3725 | 0.713 |
| Black – White | -0.08599 | 0.07827 | -1.0986 | 0.283 |
| Hispanic – White | -0.26009 | 0.14970 | -1.7374 | 0.095 |
| Gender: | | | | |
| Male – Female | -0.00562 | 0.09252 | -0.0608 | 0.952 |
| PMI converted to hours | 0.00666 | 0.00384 | 1.7355 | 0.095 |
| pH | -0.02105 | 0.13272 | -0.1586 | 0.875 |

[a] Represents reference level

### Linear Regression

Model Fit Measures

| Model | R | $R^2$ |
|---|---|---|
| 1 | 0.683 | 0.466 |

Model Coefficients - ARHGDIA

| Predictor | Estimate | SE | t | p |
|---|---|---|---|---|
| Intercept [a] | 31.21203 | 0.88058 | 35.445 | < .001 |
| Arm: | | | | |
| alcohol use disorder – control | -0.03979 | 0.09266 | -0.429 | 0.671 |
| opioid use disorder – control | 0.11224 | 0.08668 | 1.295 | 0.208 |
| opioid+alcohol use disorder – control | -0.19409 | 0.10250 | -1.893 | 0.070 |
| Age Cateogry: | | | | |
| 50-64 – 30-49 | -0.11283 | 0.09235 | -1.222 | 0.234 |
| 50-65 – 30-49 | -0.05134 | 0.14188 | -0.362 | 0.721 |
| 65+ – 30-49 | -0.02863 | 0.10699 | -0.268 | 0.791 |
| <30 – 30-49 | -0.04870 | 0.11186 | -0.435 | 0.667 |
| Ethnicity: | | | | |
| Asian – White | -0.15477 | 0.18627 | -0.831 | 0.414 |
| Black – White | -0.07676 | 0.07583 | -1.012 | 0.322 |
| Hispanic – White | -0.19898 | 0.14503 | -1.372 | 0.183 |
| Gender: | | | | |
| Male – Female | 0.03430 | 0.08963 | 0.383 | 0.705 |
| PMI converted to hours | 0.00330 | 0.00372 | 0.889 | 0.383 |
| pH | 0.05485 | 0.12858 | 0.427 | 0.673 |

[a] Represents reference level

### Linear Regression

Model Fit Measures

| Model | R | $R^2$ |
|---|---|---|
| 1 | 0.692 | 0.479 |

Model Coefficients - PPP5C

| Predictor | Estimate | SE | t | p |
|---|---|---|---|---|
| Intercept [a] | 28.14062 | 0.72964 | 38.5680 | < .001 |
| Arm: | | | | |
| alcohol use disorder – control | 0.06782 | 0.07678 | 0.8834 | 0.386 |
| opioid use disorder – control | 0.15132 | 0.07182 | 2.1070 | 0.046 |
| opioid+alcohol use disorder – control | -0.06836 | 0.08493 | -0.8049 | 0.429 |
| Age Cateogry: | | | | |
| 50-64 – 30-49 | -0.03071 | 0.07652 | -0.4013 | 0.692 |
| 50-65 – 30-49 | 0.12199 | 0.11756 | 1.0377 | 0.310 |
| 65+ – 30-49 | -0.12738 | 0.08865 | -1.4368 | 0.164 |
| <30 – 30-49 | 0.02496 | 0.09269 | 0.2693 | 0.790 |
| Ethnicity: | | | | |
| Asian – White | -0.04833 | 0.15434 | -0.3131 | 0.757 |
| Black – White | 0.01698 | 0.06283 | 0.2703 | 0.789 |
| Hispanic – White | -0.02493 | 0.12017 | -0.2075 | 0.837 |
| Gender: | | | | |
| Male – Female | -0.02478 | 0.07427 | -0.3337 | 0.741 |
| PMI converted to hours | 2.39e-4 | 0.00308 | 0.0774 | 0.939 |
| pH | 0.00193 | 0.10654 | 0.0181 | 0.986 |

[a] Represents reference level

#### Linear Regression

Model Fit Measures

| Model | R | $R^2$ |
|---|---|---|
| 1 | 0.730 | 0.533 |

Model Coefficients - SLC16A1 (2)

| Predictor | Estimate | SE | t | p |
|---|---|---|---|---|
| Intercept [a] | 24.40325 | 3.6108 | 6.7584 | < .001 |
| Arm: | | | | |
| alcohol use disorder – control | 1.21643 | 0.3799 | 3.2016 | 0.004 |
| opioid use disorder – control | 0.73092 | 0.3554 | 2.0565 | 0.051 |
| opioid+alcohol use disorder – control | 0.03280 | 0.4203 | 0.0780 | 0.938 |
| Age Cateogry: | | | | |
| 50-64 – 30-49 | 0.29737 | 0.3787 | 0.7853 | 0.440 |
| 50-65 – 30-49 | -0.54182 | 0.5818 | -0.9313 | 0.361 |
| 65+ – 30-49 | -0.04158 | 0.4387 | -0.0948 | 0.925 |
| <30 – 30-49 | 0.11551 | 0.4587 | 0.2518 | 0.803 |
| Ethnicity: | | | | |
| Asian – White | -1.12751 | 0.7638 | -1.4762 | 0.153 |
| Black – White | -0.31980 | 0.3109 | -1.0285 | 0.314 |
| Hispanic – White | -0.70758 | 0.5947 | -1.1898 | 0.246 |
| Gender: | | | | |
| Male – Female | -0.26787 | 0.3675 | -0.7288 | 0.473 |
| PMI converted to hours | -0.00945 | 0.0153 | -0.6198 | 0.541 |
| pH | 0.03583 | 0.5272 | 0.0680 | 0.946 |

[a] Represents reference level

#### Linear Regression

Model Fit Measures

| Model | R | $R^2$ |
|---|---|---|
| 1 | 0.619 | 0.383 |

Model Coefficients - EPHA4

| Predictor | Estimate | SE | t | p |
|---|---|---|---|---|
| Intercept [a] | 28.1850 | 1.07187 | 26.29523 | < .001 |
| Arm: | | | | |
| alcohol use disorder – control | 0.1123 | 0.11279 | 0.99612 | 0.329 |
| opioid use disorder – control | 0.1979 | 0.10551 | 1.87593 | 0.073 |
| opioid+alcohol use disorder – control | -0.0796 | 0.12477 | -0.63806 | 0.529 |
| Age Cateogry: | | | | |
| 50-64 – 30-49 | -8.97e−4 | 0.11242 | -0.00798 | 0.994 |
| 50-65 – 30-49 | 0.0474 | 0.17270 | 0.27436 | 0.786 |
| 65+ – 30-49 | 0.0681 | 0.13024 | 0.52261 | 0.606 |
| <30 – 30-49 | 0.1252 | 0.13616 | 0.91912 | 0.367 |
| Ethnicity: | | | | |
| Asian – White | -0.0149 | 0.22673 | -0.06563 | 0.948 |
| Black – White | 0.0192 | 0.09230 | 0.20782 | 0.837 |
| Hispanic – White | 0.1026 | 0.17654 | 0.58096 | 0.567 |
| Gender: | | | | |
| Male – Female | -0.1545 | 0.10911 | -1.41601 | 0.170 |
| PMI converted to hours | 4.90e-4 | 0.00453 | 0.10830 | 0.915 |
| pH | -0.0298 | 0.15651 | -0.19063 | 0.850 |

[a] Represents reference level

### Linear Regression

Model Fit Measures

| Model | R | $R^2$ |
|---|---|---|
| 1 | 0.691 | 0.477 |

Model Coefficients - CACNA2D1

| Predictor | Estimate | SE | t | p |
|---|---|---|---|---|
| Intercept [a] | 28.8606 | 1.46625 | 19.6834 | < .001 |
| Arm: | | | | |
| alcohol use disorder – control | -0.0101 | 0.15428 | -0.0655 | 0.948 |
| opioid use disorder – control | 0.2042 | 0.14433 | 1.4150 | 0.170 |
| opioid+alcohol use disorder – control | -0.2968 | 0.17068 | -1.7388 | 0.095 |
| Age Cateogry: | | | | |
| 50-64 – 30-49 | -0.1494 | 0.15378 | -0.9718 | 0.341 |
| 50-65 – 30-49 | -0.0902 | 0.23624 | -0.3818 | 0.706 |
| 65+ – 30-49 | -0.0490 | 0.17815 | -0.2752 | 0.786 |
| <30 – 30-49 | -0.0539 | 0.18626 | -0.2894 | 0.775 |
| Ethnicity: | | | | |
| Asian – White | -0.1631 | 0.31015 | -0.5259 | 0.604 |
| Black – White | 0.0230 | 0.12627 | 0.1820 | 0.857 |
| Hispanic – White | -0.3413 | 0.24149 | -1.4133 | 0.170 |
| Gender: | | | | |
| Male – Female | -0.0277 | 0.14925 | -0.1854 | 0.855 |
| PMI converted to hours | 0.0105 | 0.00619 | 1.6999 | 0.102 |
| pH | 0.0420 | 0.21409 | 0.1964 | 0.846 |

[a] Represents reference level

### Linear Regression

Model Fit Measures

| Model | R | $R^2$ |
|---|---|---|
| 1 | 0.665 | 0.442 |

Model Coefficients - CSE1L

| Predictor | Estimate | SE | t | p |
|---|---|---|---|---|
| Intercept [a] | 23.95805 | 6.1117 | 3.920 | < .001 |
| Arm: | | | | |
| alcohol use disorder – control | -0.39180 | 0.6431 | -0.609 | 0.548 |
| opioid use disorder – control | -0.57672 | 0.6016 | -0.959 | 0.347 |
| opioid+alcohol use disorder – control | 0.21359 | 0.7114 | 0.300 | 0.767 |
| Age Cateogry: | | | | |
| 50-64 – 30-49 | 1.23242 | 0.6410 | 1.923 | 0.066 |
| 50-65 – 30-49 | 1.59208 | 0.9847 | 1.617 | 0.119 |
| 65+ – 30-49 | 1.25418 | 0.7426 | 1.689 | 0.104 |
| <30 – 30-49 | 1.15972 | 0.7764 | 1.494 | 0.148 |
| Ethnicity: | | | | |
| Asian – White | 1.60771 | 1.2928 | 1.244 | 0.226 |
| Black – White | 0.18767 | 0.5263 | 0.357 | 0.725 |
| Hispanic – White | 1.33236 | 1.0066 | 1.324 | 0.198 |
| Gender: | | | | |
| Male – Female | -0.25310 | 0.6221 | -0.407 | 0.688 |
| PMI converted to hours | -0.00686 | 0.0258 | -0.266 | 0.793 |
| pH | 0.33353 | 0.8924 | 0.374 | 0.712 |

[a] Represents reference level

### Linear Regression

Model Fit Measures

| Model | R | $R^2$ |
|---|---|---|
| 1 | 0.601 | 0.361 |

Model Coefficients - RABGGTB

| Predictor | Estimate | SE | t | p |
|---|---|---|---|---|
| Intercept [a] | 21.9085 | 8.1102 | 2.7014 | 0.012 |
| Arm: | | | | |
| alcohol use disorder – control | -0.5925 | 0.8534 | -0.6943 | 0.494 |
| opioid use disorder – control | 0.9974 | 0.7983 | 1.2494 | 0.224 |
| opioid+alcohol use disorder – control | -0.5124 | 0.9441 | -0.5427 | 0.592 |
| Age Cateogry: | | | | |
| 50-64 – 30-49 | -0.7214 | 0.8506 | -0.8481 | 0.405 |
| 50-65 – 30-49 | -0.7075 | 1.3067 | -0.5414 | 0.593 |
| 65+ – 30-49 | -0.4449 | 0.9854 | -0.4515 | 0.656 |
| <30 – 30-49 | -0.9252 | 1.0303 | -0.8980 | 0.378 |
| Ethnicity: | | | | |
| Asian – White | -1.2401 | 1.7155 | -0.7229 | 0.477 |
| Black – White | -0.9084 | 0.6984 | -1.3006 | 0.206 |
| Hispanic – White | -0.1904 | 1.3357 | -0.1425 | 0.888 |
| Gender: | | | | |
| Male – Female | -0.0447 | 0.8255 | -0.0541 | 0.957 |
| PMI converted to hours | -0.0267 | 0.0343 | -0.7797 | 0.443 |
| pH | 0.5157 | 1.1842 | 0.4355 | 0.667 |

[a] Represents reference level

### Linear Regression

Model Fit Measures

| Model | R | $R^2$ |
|---|---|---|
| 1 | 0.770 | 0.594 |

Model Coefficients - SUCLG1

| Predictor | Estimate | SE | t | p |
|---|---|---|---|---|
| Intercept [a] | 29.51194 | 1.97292 | 14.95850 | < .001 |
| Arm: | | | | |
| alcohol use disorder – control | 0.11540 | 0.20760 | 0.55587 | 0.583 |
| opioid use disorder – control | -0.24551 | 0.19420 | -1.26421 | 0.218 |
| opioid+alcohol use disorder – control | -0.11355 | 0.22966 | -0.49446 | 0.625 |
| Age Cateogry: | | | | |
| 50-64 – 30-49 | 0.60654 | 0.20692 | 2.93133 | 0.007 |
| 50-65 – 30-49 | 0.15894 | 0.31788 | 0.49999 | 0.622 |
| 65+ – 30-49 | 0.69042 | 0.23972 | 2.88015 | 0.008 |
| <30 – 30-49 | 0.57863 | 0.25063 | 2.30869 | 0.030 |
| Ethnicity: | | | | |
| Asian – White | 0.99231 | 0.41733 | 2.37777 | 0.026 |
| Black – White | 0.07920 | 0.16990 | 0.46618 | 0.645 |
| Hispanic – White | 0.35852 | 0.32494 | 1.10335 | 0.281 |
| Gender: | | | | |
| Male – Female | -0.29271 | 0.20082 | -1.45756 | 0.158 |
| PMI converted to hours | 2.63e-4 | 0.00833 | 0.03156 | 0.975 |
| pH | 0.00120 | 0.28807 | 0.00417 | 0.997 |

[a] Represents reference level

#### Linear Regression

Model Fit Measures

| Model | R | $R^2$ |
|---|---|---|
| 1 | 0.588 | 0.346 |

Model Coefficients - PDAP1

| Predictor | Estimate | SE | t | p |
|---|---|---|---|---|
| Intercept [a] | 23.35502 | 4.1818 | 5.5849 | < .001 |
| Arm: | | | | |
| alcohol use disorder – control | 0.26922 | 0.4400 | 0.6118 | 0.546 |
| opioid use disorder – control | 0.57413 | 0.4116 | 1.3948 | 0.176 |
| opioid+alcohol use disorder – control | -0.57507 | 0.4868 | -1.1814 | 0.249 |
| Age Cateogry: | | | | |
| 50-64 – 30-49 | -0.16518 | 0.4386 | -0.3766 | 0.710 |
| 50-65 – 30-49 | 0.59405 | 0.6738 | 0.8817 | 0.387 |
| 65+ – 30-49 | 0.43585 | 0.5081 | 0.8578 | 0.399 |
| <30 – 30-49 | 0.11802 | 0.5312 | 0.2222 | 0.826 |
| Ethnicity: | | | | |
| Asian – White | 0.33595 | 0.8846 | 0.3798 | 0.707 |
| Black – White | 0.07007 | 0.3601 | 0.1946 | 0.847 |
| Hispanic – White | 0.40697 | 0.6887 | 0.5909 | 0.560 |
| Gender: | | | | |
| Male – Female | -0.01848 | 0.4257 | -0.0434 | 0.966 |
| PMI converted to hours | 0.00103 | 0.0177 | 0.0584 | 0.954 |
| pH | 0.33994 | 0.6106 | 0.5567 | 0.583 |

[a] Represents reference level

#### Linear Regression

Model Fit Measures

| Model | R | $R^2$ |
|---|---|---|
| 1 | 0.757 | 0.573 |

Model Coefficients - RUFY3 (2)

| Predictor | Estimate | SE | t | p |
|---|---|---|---|---|
| Intercept [a] | 27.16509 | 2.14677 | 12.65391 | < .001 |
| Arm: | | | | |
| alcohol use disorder – control | -0.34716 | 0.22589 | -1.53682 | 0.137 |
| opioid use disorder – control | -0.35676 | 0.21131 | -1.68827 | 0.104 |
| opioid+alcohol use disorder – control | 0.25354 | 0.24989 | 1.01459 | 0.320 |
| Age Cateogry: | | | | |
| 50-64 – 30-49 | 0.45506 | 0.22515 | 2.02115 | 0.055 |
| 50-65 – 30-49 | 0.29346 | 0.34589 | 0.84843 | 0.405 |
| 65+ – 30-49 | 0.44104 | 0.26084 | 1.69082 | 0.104 |
| <30 – 30-49 | -0.04089 | 0.27271 | -0.14992 | 0.882 |
| Ethnicity: | | | | |
| Asian – White | 0.35477 | 0.45410 | 0.78124 | 0.442 |
| Black – White | 0.31588 | 0.18487 | 1.70865 | 0.100 |
| Hispanic – White | 0.19270 | 0.35357 | 0.54501 | 0.591 |
| Gender: | | | | |
| Male – Female | 0.00203 | 0.21852 | 0.00927 | 0.993 |
| PMI converted to hours | -0.00201 | 0.00907 | -0.22142 | 0.827 |
| pH | 0.07932 | 0.31345 | 0.25304 | 0.802 |

[a] Represents reference level

#### Linear Regression

Model Fit Measures

| Model | R | $R^2$ |
|---|---|---|
| 1 | 0.765 | 0.585 |

Model Coefficients - PPP2R5D

| Predictor | Estimate | SE | t | p |
|---|---|---|---|---|
| Intercept [a] | 29.2990 | 4.6198 | 6.342 | < .001 |
| Arm: | | | | |
| alcohol use disorder – control | -0.3085 | 0.4861 | -0.635 | 0.532 |
| opioid use disorder – control | -0.9760 | 0.4547 | -2.146 | 0.042 |
| opioid+alcohol use disorder – control | 0.8043 | 0.5378 | 1.496 | 0.148 |
| Age Cateogry: | | | | |
| 50-64 – 30-49 | 0.7006 | 0.4845 | 1.446 | 0.161 |
| 50-65 – 30-49 | -0.1945 | 0.7443 | -0.261 | 0.796 |
| 65+ – 30-49 | 0.6984 | 0.5613 | 1.244 | 0.225 |
| <30 – 30-49 | 0.3168 | 0.5869 | 0.540 | 0.594 |
| Ethnicity: | | | | |
| Asian – White | 0.9321 | 0.9772 | 0.954 | 0.350 |
| Black – White | 1.0204 | 0.3978 | 2.565 | 0.017 |
| Hispanic – White | 0.8960 | 0.7609 | 1.178 | 0.250 |
| Gender: | | | | |
| Male – Female | 0.2465 | 0.4703 | 0.524 | 0.605 |
| PMI converted to hours | -0.0330 | 0.0195 | -1.689 | 0.104 |
| pH | -0.4498 | 0.6746 | -0.667 | 0.511 |

[a] Represents reference level

#### Linear Regression

Model Fit Measures

| Model | R | $R^2$ |
|---|---|---|
| 1 | 0.748 | 0.559 |

Model Coefficients - RAB5B

| Predictor | Estimate | SE | t | p |
|---|---|---|---|---|
| Intercept [a] | 28.96049 | 0.95355 | 30.3713 | < .001 |
| Arm: | | | | |
| alcohol use disorder – control | 0.33101 | 0.10034 | 3.2990 | 0.003 |
| opioid use disorder – control | 0.31364 | 0.09386 | 3.3415 | 0.003 |
| opioid+alcohol use disorder – control | 0.00161 | 0.11100 | 0.0145 | 0.989 |
| Age Cateogry: | | | | |
| 50-64 – 30-49 | 0.10116 | 0.10001 | 1.0115 | 0.322 |
| 50-65 – 30-49 | 0.15573 | 0.15364 | 1.0136 | 0.321 |
| 65+ – 30-49 | -0.00743 | 0.11586 | -0.0641 | 0.949 |
| <30 – 30-49 | -0.05186 | 0.12113 | -0.4281 | 0.672 |
| Ethnicity: | | | | |
| Asian – White | 0.08847 | 0.20170 | 0.4386 | 0.665 |
| Black – White | 0.04559 | 0.08211 | 0.5552 | 0.584 |
| Hispanic – White | -0.19913 | 0.15705 | -1.2679 | 0.217 |
| Gender: | | | | |
| Male – Female | -0.01451 | 0.09706 | -0.1495 | 0.882 |
| PMI converted to hours | 0.00588 | 0.00403 | 1.4600 | 0.157 |
| pH | -0.24791 | 0.13923 | -1.7806 | 0.088 |

[a] Represents reference level

### Linear Regression

Model Fit Measures

| Model | R | $R^2$ |
|---|---|---|
| 1 | 0.739 | 0.546 |

Model Coefficients - HNRNPK

| Predictor | Estimate | SE | t | p |
|---|---|---|---|---|
| Intercept [a] | 30.56168 | 0.76605 | 39.895 | < .001 |
| Arm: | | | | |
| alcohol use disorder – control | -0.07634 | 0.08061 | -0.947 | 0.353 |
| opioid use disorder – control | -0.24368 | 0.07541 | -3.232 | 0.004 |
| opioid+alcohol use disorder – control | 0.03154 | 0.08917 | 0.354 | 0.727 |
| Age Cateogry: | | | | |
| 50-64 – 30-49 | -0.01713 | 0.08034 | -0.213 | 0.833 |
| 50-65 – 30-49 | -0.24086 | 0.12343 | -1.951 | 0.063 |
| 65+ – 30-49 | -0.03529 | 0.09308 | -0.379 | 0.708 |
| <30 – 30-49 | 0.11178 | 0.09732 | 1.149 | 0.262 |
| Ethnicity: | | | | |
| Asian – White | 0.16230 | 0.16204 | 1.002 | 0.327 |
| Black – White | -0.03827 | 0.06597 | -0.580 | 0.567 |
| Hispanic – White | 0.09624 | 0.12617 | 0.763 | 0.453 |
| Gender: | | | | |
| Male – Female | -0.09014 | 0.07798 | -1.156 | 0.259 |
| PMI converted to hours | -0.00105 | 0.00324 | -0.325 | 0.748 |
| pH | -0.03945 | 0.11185 | -0.353 | 0.727 |

[a] Represents reference level

### Linear Regression

Model Fit Measures

| Model | R | $R^2$ |
|---|---|---|
| 1 | 0.677 | 0.458 |

Model Coefficients - RPS29

| Predictor | Estimate | SE | t | p |
|---|---|---|---|---|
| Intercept [a] | 20.6099 | 3.9693 | 5.192 | < .001 |
| Arm: | | | | |
| alcohol use disorder – control | -0.1706 | 0.4177 | -0.409 | 0.686 |
| opioid use disorder – control | 1.0089 | 0.3907 | 2.582 | 0.016 |
| opioid+alcohol use disorder – control | -0.2854 | 0.4620 | -0.618 | 0.543 |
| Age Cateogry: | | | | |
| 50-64 – 30-49 | 0.1773 | 0.4163 | 0.426 | 0.674 |
| 50-65 – 30-49 | 0.2575 | 0.6395 | 0.403 | 0.691 |
| 65+ – 30-49 | 0.3743 | 0.4823 | 0.776 | 0.445 |
| <30 – 30-49 | -0.1162 | 0.5042 | -0.230 | 0.820 |
| Ethnicity: | | | | |
| Asian – White | -0.7466 | 0.8396 | -0.889 | 0.383 |
| Black – White | 0.0815 | 0.3418 | 0.238 | 0.814 |
| Hispanic – White | 0.2939 | 0.6537 | 0.450 | 0.657 |
| Gender: | | | | |
| Male – Female | 0.2813 | 0.4040 | 0.696 | 0.493 |
| PMI converted to hours | -0.0106 | 0.0168 | -0.635 | 0.531 |
| pH | 0.4336 | 0.5796 | 0.748 | 0.462 |

[a] Represents reference level

### Linear Regression

Model Fit Measures

| Model | R | $R^2$ |
|---|---|---|
| 1 | 0.750 | 0.562 |

Model Coefficients - SLC7A5

| Predictor | Estimate | SE | t | p |
|---|---|---|---|---|
| Intercept [a] | 26.3631 | 3.8588 | 6.8320 | < .001 |
| Arm: | | | | |
| alcohol use disorder – control | 0.6561 | 0.4060 | 1.6160 | 0.119 |
| opioid use disorder – control | 0.4351 | 0.3798 | 1.1454 | 0.263 |
| opioid+alcohol use disorder – control | -1.0807 | 0.4492 | -2.4059 | 0.024 |
| Age Cateogry: | | | | |
| 50-64 – 30-49 | 0.5436 | 0.4047 | 1.3432 | 0.192 |
| 50-65 – 30-49 | 0.1328 | 0.6217 | 0.2135 | 0.833 |
| 65+ – 30-49 | -0.1267 | 0.4689 | -0.2702 | 0.789 |
| <30 – 30-49 | 0.9462 | 0.4902 | 1.9303 | 0.065 |
| Ethnicity: | | | | |
| Asian – White | -2.1143 | 0.8162 | -2.5903 | 0.016 |
| Black – White | -0.5970 | 0.3323 | -1.7967 | 0.085 |
| Hispanic – White | -0.0228 | 0.6355 | -0.0359 | 0.972 |
| Gender: | | | | |
| Male – Female | -0.4503 | 0.3928 | -1.1464 | 0.263 |
| PMI converted to hours | -0.0127 | 0.0163 | -0.7782 | 0.444 |
| pH | -0.0283 | 0.5634 | -0.0503 | 0.960 |

[a] Represents reference level

### Linear Regression

Model Fit Measures

| Model | R | $R^2$ |
|---|---|---|
| 1 | 0.732 | 0.535 |

Model Coefficients - DPYSL3

| Predictor | Estimate | SE | t | p |
|---|---|---|---|---|
| Intercept [a] | 31.37013 | 0.75377 | 41.6176 | < .001 |
| Arm: | | | | |
| alcohol use disorder – control | 0.05053 | 0.07931 | 0.6370 | 0.530 |
| opioid use disorder – control | 0.20252 | 0.07420 | 2.7295 | 0.012 |
| opioid+alcohol use disorder – control | 0.06957 | 0.08774 | 0.7929 | 0.436 |
| Age Cateogry: | | | | |
| 50-64 – 30-49 | -0.05717 | 0.07905 | -0.7231 | 0.477 |
| 50-65 – 30-49 | -0.00756 | 0.12145 | -0.0622 | 0.951 |
| 65+ – 30-49 | -0.05818 | 0.09159 | -0.6352 | 0.531 |
| <30 – 30-49 | 0.05879 | 0.09575 | 0.6139 | 0.545 |
| Ethnicity: | | | | |
| Asian – White | -0.09953 | 0.15944 | -0.6242 | 0.538 |
| Black – White | -0.03051 | 0.06491 | -0.4700 | 0.643 |
| Hispanic – White | -0.15577 | 0.12415 | -1.2547 | 0.222 |
| Gender: | | | | |
| Male – Female | -0.04936 | 0.07673 | -0.6433 | 0.526 |
| PMI converted to hours | 0.00724 | 0.00318 | 2.2733 | 0.032 |
| pH | -0.00992 | 0.11006 | -0.0901 | 0.929 |

[a] Represents reference level

#### Linear Regression

Model Fit Measures

| Model | R | $R^2$ |
|---|---|---|
| 1 | 0.768 | 0.590 |

Model Coefficients - DYNC1H1

| Predictor | Estimate | SE | t | p |
|---|---|---|---|---|
| Intercept [a] | 31.43985 | 1.78040 | 17.659 | < .001 |
| Arm: | | | | |
| alcohol use disorder – control | -0.15051 | 0.18734 | -0.803 | 0.430 |
| opioid use disorder – control | -0.32863 | 0.17525 | -1.875 | 0.073 |
| opioid+alcohol use disorder – control | 0.09685 | 0.20725 | 0.467 | 0.644 |
| Age Cateogry: | | | | |
| 50-64 – 30-49 | 0.43332 | 0.18672 | 2.321 | 0.029 |
| 50-65 – 30-49 | -0.09567 | 0.28686 | -0.334 | 0.742 |
| 65+ – 30-49 | 0.49973 | 0.21633 | 2.310 | 0.030 |
| <30 – 30-49 | 0.09078 | 0.22617 | 0.401 | 0.692 |
| Ethnicity: | | | | |
| Asian – White | 0.51712 | 0.37660 | 1.373 | 0.182 |
| Black – White | 0.15645 | 0.15332 | 1.020 | 0.318 |
| Hispanic – White | 0.21710 | 0.29323 | 0.740 | 0.466 |
| Gender: | | | | |
| Male – Female | -0.02548 | 0.18123 | -0.141 | 0.889 |
| PMI converted to hours | -0.00446 | 0.00752 | -0.594 | 0.558 |
| pH | 0.09655 | 0.25996 | 0.371 | 0.714 |

[a] Represents reference level

#### Linear Regression

Model Fit Measures

| Model | R | $R^2$ |
|---|---|---|
| 1 | 0.745 | 0.555 |

Model Coefficients - PRPSAP1

| Predictor | Estimate | SE | t | p |
|---|---|---|---|---|
| Intercept [a] | 25.7942 | 5.3807 | 4.794 | < .001 |
| Arm: | | | | |
| alcohol use disorder – control | -0.7412 | 0.5662 | -1.309 | 0.203 |
| opioid use disorder – control | 1.4177 | 0.5296 | 2.677 | 0.013 |
| opioid+alcohol use disorder – control | 0.3818 | 0.6263 | 0.610 | 0.548 |
| Age Cateogry: | | | | |
| 50-64 – 30-49 | -0.1856 | 0.5643 | -0.329 | 0.745 |
| 50-65 – 30-49 | 0.8872 | 0.8669 | 1.023 | 0.316 |
| 65+ – 30-49 | 1.2970 | 0.6538 | 1.984 | 0.059 |
| <30 – 30-49 | -1.5923 | 0.6835 | -2.329 | 0.029 |
| Ethnicity: | | | | |
| Asian – White | 0.4320 | 1.1382 | 0.380 | 0.708 |
| Black – White | 0.4144 | 0.4634 | 0.894 | 0.380 |
| Hispanic – White | -1.9667 | 0.8862 | -2.219 | 0.036 |
| Gender: | | | | |
| Male – Female | 0.8795 | 0.5477 | 1.606 | 0.121 |
| PMI converted to hours | 0.0672 | 0.0227 | 2.957 | 0.007 |
| pH | -0.4737 | 0.7856 | -0.603 | 0.552 |

[a] Represents reference level

### Linear Regression

Model Fit Measures

| Model | R | $R^2$ |
|---|---|---|
| 1 | 0.726 | 0.526 |

Model Coefficients - WDR44

| Predictor | Estimate | SE | t | p |
|---|---|---|---|---|
| Intercept [a] | 25.1927 | 2.9666 | 8.4921 | < .001 |
| Arm: | | | | |
| alcohol use disorder – control | 0.4332 | 0.3122 | 1.3878 | 0.178 |
| opioid use disorder – control | 0.6444 | 0.2920 | 2.2068 | 0.037 |
| opioid+alcohol use disorder – control | -0.0219 | 0.3453 | -0.0634 | 0.950 |
| Age Cateogry: | | | | |
| 50-64 – 30-49 | -0.1018 | 0.3111 | -0.3273 | 0.746 |
| 50-65 – 30-49 | 0.1000 | 0.4780 | 0.2092 | 0.836 |
| 65+ – 30-49 | 0.0598 | 0.3605 | 0.1659 | 0.870 |
| <30 – 30-49 | -0.5307 | 0.3769 | -1.4081 | 0.172 |
| Ethnicity: | | | | |
| Asian – White | 0.4366 | 0.6275 | 0.6957 | 0.493 |
| Black – White | 0.2090 | 0.2555 | 0.8182 | 0.421 |
| Hispanic – White | -1.3988 | 0.4886 | -2.8629 | 0.009 |
| Gender: | | | | |
| Male – Female | -0.0901 | 0.3020 | -0.2985 | 0.768 |
| PMI converted to hours | 0.0144 | 0.0125 | 1.1459 | 0.263 |
| pH | 0.0924 | 0.4332 | 0.2132 | 0.833 |

[a] Represents reference level

### Linear Regression

Model Fit Measures

| Model | R | $R^2$ |
|---|---|---|
| 1 | 0.740 | 0.547 |

Model Coefficients - ETF1

| Predictor | Estimate | SE | t | p |
|---|---|---|---|---|
| Intercept [a] | 23.80589 | 3.6335 | 6.552 | < .001 |
| Arm: | | | | |
| alcohol use disorder – control | 0.24092 | 0.3823 | 0.630 | 0.535 |
| opioid use disorder – control | 0.60191 | 0.3577 | 1.683 | 0.105 |
| opioid+alcohol use disorder – control | -0.72144 | 0.4230 | -1.706 | 0.101 |
| Age Cateogry: | | | | |
| 50-64 – 30-49 | -0.04153 | 0.3811 | -0.109 | 0.914 |
| 50-65 – 30-49 | -0.83889 | 0.5854 | -1.433 | 0.165 |
| 65+ – 30-49 | -0.18299 | 0.4415 | -0.414 | 0.682 |
| <30 – 30-49 | -0.45137 | 0.4616 | -0.978 | 0.338 |
| Ethnicity: | | | | |
| Asian – White | -1.50700 | 0.7686 | -1.961 | 0.062 |
| Black – White | 0.09927 | 0.3129 | 0.317 | 0.754 |
| Hispanic – White | -1.07943 | 0.5984 | -1.804 | 0.084 |
| Gender: | | | | |
| Male – Female | 0.12785 | 0.3699 | 0.346 | 0.733 |
| PMI converted to hours | 0.00378 | 0.0153 | 0.246 | 0.808 |
| pH | 0.07177 | 0.5305 | 0.135 | 0.894 |

[a] Represents reference level

### Linear Regression

Model Fit Measures

| Model | R | $R^2$ |
|---|---|---|
| 1 | 0.750 | 0.562 |

Model Coefficients - PFKP

| Predictor | Estimate | SE | t | p |
|---|---|---|---|---|
| Intercept [a] | 31.49643 | 1.70186 | 18.507 | < .001 |
| Arm: | | | | |
| alcohol use disorder – control | -0.07041 | 0.17908 | -0.393 | 0.698 |
| opioid use disorder – control | -0.36344 | 0.16752 | -2.170 | 0.040 |
| opioid+alcohol use disorder – control | 0.03814 | 0.19810 | 0.193 | 0.849 |
| Age Cateogry: | | | | |
| 50-64 – 30-49 | 0.44197 | 0.17849 | 2.476 | 0.021 |
| 50-65 – 30-49 | -0.30717 | 0.27420 | -1.120 | 0.274 |
| 65+ – 30-49 | 0.27132 | 0.20678 | 1.312 | 0.202 |
| <30 – 30-49 | 0.07822 | 0.21619 | 0.362 | 0.721 |
| Ethnicity: | | | | |
| Asian – White | 0.08854 | 0.35999 | 0.246 | 0.808 |
| Black – White | 0.13005 | 0.14656 | 0.887 | 0.384 |
| Hispanic – White | 0.08862 | 0.28029 | 0.316 | 0.755 |
| Gender: | | | | |
| Male – Female | -0.04944 | 0.17323 | -0.285 | 0.778 |
| PMI converted to hours | -0.00481 | 0.00719 | -0.669 | 0.510 |
| pH | -0.13349 | 0.24849 | -0.537 | 0.596 |

[a] Represents reference level

### Linear Regression

Model Fit Measures

| Model | R | $R^2$ |
|---|---|---|
| 1 | 0.712 | 0.507 |

Model Coefficients - HNRNPD

| Predictor | Estimate | SE | t | p |
|---|---|---|---|---|
| Intercept [a] | 27.96025 | 1.22647 | 22.797 | < .001 |
| Arm: | | | | |
| alcohol use disorder – control | -0.09613 | 0.12905 | -0.745 | 0.464 |
| opioid use disorder – control | -0.19725 | 0.12073 | -1.634 | 0.115 |
| opioid+alcohol use disorder – control | 0.14954 | 0.14277 | 1.047 | 0.305 |
| Age Cateogry: | | | | |
| 50-64 – 30-49 | 0.18964 | 0.12863 | 1.474 | 0.153 |
| 50-65 – 30-49 | -0.02323 | 0.19761 | -0.118 | 0.907 |
| 65+ – 30-49 | 0.25201 | 0.14902 | 1.691 | 0.104 |
| <30 – 30-49 | 0.18401 | 0.15580 | 1.181 | 0.249 |
| Ethnicity: | | | | |
| Asian – White | 0.39438 | 0.25943 | 1.520 | 0.142 |
| Black – White | 0.10528 | 0.10562 | 0.997 | 0.329 |
| Hispanic – White | 0.11271 | 0.20200 | 0.558 | 0.582 |
| Gender: | | | | |
| Male – Female | 0.04678 | 0.12484 | 0.375 | 0.711 |
| PMI converted to hours | -0.00522 | 0.00518 | -1.007 | 0.324 |
| pH | 0.06936 | 0.17908 | 0.387 | 0.702 |

[a] Represents reference level

#### Linear Regression

Model Fit Measures

| Model | R | $R^2$ |
|---|---|---|
| 1 | 0.673 | 0.453 |

Model Coefficients - EIF6

| Predictor | Estimate | SE | t | p |
|---|---|---|---|---|
| Intercept [a] | 23.6499 | 2.4832 | 9.524 | < .001 |
| Arm: | | | | |
| alcohol use disorder – control | 0.4372 | 0.2613 | 1.673 | 0.107 |
| opioid use disorder – control | 0.5947 | 0.2444 | 2.433 | 0.023 |
| opioid+alcohol use disorder – control | 0.0665 | 0.2891 | 0.230 | 0.820 |
| Age Cateogry: | | | | |
| 50-64 – 30-49 | -0.0575 | 0.2604 | -0.221 | 0.827 |
| 50-65 – 30-49 | 0.2717 | 0.4001 | 0.679 | 0.504 |
| 65+ – 30-49 | -0.3652 | 0.3017 | -1.210 | 0.238 |
| <30 – 30-49 | -0.3497 | 0.3155 | -1.108 | 0.279 |
| Ethnicity: | | | | |
| Asian – White | -0.3802 | 0.5253 | -0.724 | 0.476 |
| Black – White | -0.0654 | 0.2138 | -0.306 | 0.762 |
| Hispanic – White | -0.3324 | 0.4090 | -0.813 | 0.424 |
| Gender: | | | | |
| Male – Female | 0.1724 | 0.2528 | 0.682 | 0.502 |
| PMI converted to hours | 0.0179 | 0.0105 | 1.703 | 0.101 |
| pH | 0.1710 | 0.3626 | 0.472 | 0.642 |

[a] Represents reference level

#### Linear Regression

Model Fit Measures

| Model | R | $R^2$ |
|---|---|---|
| 1 | 0.732 | 0.536 |

Model Coefficients - SET

| Predictor | Estimate | SE | t | p |
|---|---|---|---|---|
| Intercept [a] | 27.9511 | 2.8125 | 9.938 | < .001 |
| Arm: | | | | |
| alcohol use disorder – control | 0.0356 | 0.2959 | 0.120 | 0.905 |
| opioid use disorder – control | 0.5913 | 0.2768 | 2.136 | 0.043 |
| opioid+alcohol use disorder – control | -0.5221 | 0.3274 | -1.595 | 0.124 |
| Age Cateogry: | | | | |
| 50-64 – 30-49 | 0.0923 | 0.2950 | 0.313 | 0.757 |
| 50-65 – 30-49 | -0.7881 | 0.4531 | -1.739 | 0.095 |
| 65+ – 30-49 | -0.1428 | 0.3417 | -0.418 | 0.680 |
| <30 – 30-49 | -0.1680 | 0.3573 | -0.470 | 0.643 |
| Ethnicity: | | | | |
| Asian – White | -0.3569 | 0.5949 | -0.600 | 0.554 |
| Black – White | -0.2540 | 0.2422 | -1.049 | 0.305 |
| Hispanic – White | -1.1402 | 0.4632 | -2.462 | 0.021 |
| Gender: | | | | |
| Male – Female | 0.3012 | 0.2863 | 1.052 | 0.303 |
| PMI converted to hours | 0.0227 | 0.0119 | 1.908 | 0.068 |
| pH | -0.8087 | 0.4107 | -1.969 | 0.061 |

[a] Represents reference level

### Linear Regression

Model Fit Measures

| Model | R | $R^2$ |
|---|---|---|
| 1 | 0.636 | 0.405 |

Model Coefficients - SSBP1

| Predictor | Estimate | SE | t | p |
|---|---|---|---|---|
| Intercept [a] | 28.7554 | 0.88629 | 32.4445 | < .001 |
| Arm: | | | | |
| alcohol use disorder – control | -0.2108 | 0.09326 | -2.2607 | 0.033 |
| opioid use disorder – control | -0.2111 | 0.08724 | -2.4201 | 0.023 |
| opioid+alcohol use disorder – control | -0.0530 | 0.10317 | -0.5134 | 0.612 |
| Age Cateogry: | | | | |
| 50-64 – 30-49 | -0.0484 | 0.09295 | -0.5212 | 0.607 |
| 50-65 – 30-49 | -0.0234 | 0.14280 | -0.1638 | 0.871 |
| 65+ – 30-49 | 0.0256 | 0.10769 | 0.2376 | 0.814 |
| <30 – 30-49 | -0.0583 | 0.11259 | -0.5179 | 0.609 |
| Ethnicity: | | | | |
| Asian – White | 0.1010 | 0.18748 | 0.5388 | 0.595 |
| Black – White | -0.0248 | 0.07632 | -0.3247 | 0.748 |
| Hispanic – White | -0.1800 | 0.14597 | -1.2331 | 0.229 |
| Gender: | | | | |
| Male – Female | -0.0298 | 0.09022 | -0.3301 | 0.744 |
| PMI converted to hours | 2.10e-4 | 0.00374 | 0.0560 | 0.956 |
| pH | -0.0550 | 0.12941 | -0.4252 | 0.675 |

[a] Represents reference level

### Linear Regression

Model Fit Measures

| Model | R | $R^2$ |
|---|---|---|
| 1 | 0.799 | 0.638 |

Model Coefficients - CAMK2G

| Predictor | Estimate | SE | t | p |
|---|---|---|---|---|
| Intercept [a] | 25.16124 | 5.1583 | 4.8778 | < .001 |
| Arm: | | | | |
| alcohol use disorder – control | -0.83487 | 0.5428 | -1.5382 | 0.137 |
| opioid use disorder – control | -0.79632 | 0.5077 | -1.5683 | 0.130 |
| opioid+alcohol use disorder – control | 0.34431 | 0.6004 | 0.5734 | 0.572 |
| Age Cateogry: | | | | |
| 50-64 – 30-49 | 1.13714 | 0.5410 | 2.1020 | 0.046 |
| 50-65 – 30-49 | 0.09502 | 0.8311 | 0.1143 | 0.910 |
| 65+ – 30-49 | 1.85832 | 0.6268 | 2.9650 | 0.007 |
| <30 – 30-49 | 1.33277 | 0.6553 | 2.0339 | 0.053 |
| Ethnicity: | | | | |
| Asian – White | 0.88918 | 1.0911 | 0.8149 | 0.423 |
| Black – White | -0.08315 | 0.4442 | -0.1872 | 0.853 |
| Hispanic – White | 1.03992 | 0.8496 | 1.2241 | 0.233 |
| Gender: | | | | |
| Male – Female | 0.85025 | 0.5251 | 1.6193 | 0.118 |
| PMI converted to hours | -0.00467 | 0.0218 | -0.2142 | 0.832 |
| pH | -0.04524 | 0.7532 | -0.0601 | 0.953 |

[a] Represents reference level

### Linear Regression

Model Fit Measures

| Model | R | $R^2$ |
|---|---|---|
| 1 | 0.802 | 0.643 |

Model Coefficients - NCS1

| Predictor | Estimate | SE | t | p |
|---|---|---|---|---|
| Intercept [a] | 23.2380 | 3.8898 | 5.974 | < .001 |
| Arm: | | | | |
| alcohol use disorder – control | -0.6642 | 0.4093 | -1.623 | 0.118 |
| opioid use disorder – control | -1.4902 | 0.3829 | -3.892 | < .001 |
| opioid+alcohol use disorder – control | 0.5386 | 0.4528 | 1.189 | 0.246 |
| Age Cateogry: | | | | |
| 50-64 – 30-49 | -0.6824 | 0.4080 | -1.673 | 0.107 |
| 50-65 – 30-49 | -0.3668 | 0.6267 | -0.585 | 0.564 |
| 65+ – 30-49 | -0.9235 | 0.4726 | -1.954 | 0.062 |
| <30 – 30-49 | 0.5692 | 0.4941 | 1.152 | 0.261 |
| Ethnicity: | | | | |
| Asian – White | 0.5554 | 0.8228 | 0.675 | 0.506 |
| Black – White | -0.2690 | 0.3350 | -0.803 | 0.430 |
| Hispanic – White | 1.4972 | 0.6406 | 2.337 | 0.028 |
| Gender: | | | | |
| Male – Female | -0.3012 | 0.3959 | -0.761 | 0.454 |
| PMI converted to hours | -0.0278 | 0.0164 | -1.694 | 0.103 |
| pH | 0.4282 | 0.5680 | 0.754 | 0.458 |

[a] Represents reference level

### Linear Regression

Model Fit Measures

| Model | R | $R^2$ |
|---|---|---|
| 1 | 0.738 | 0.545 |

Model Coefficients - CCBL2

| Predictor | Estimate | SE | t | p |
|---|---|---|---|---|
| Intercept [a] | 20.7383 | 4.7822 | 4.3366 | < .001 |
| Arm: | | | | |
| alcohol use disorder – control | -0.2624 | 0.5032 | -0.5214 | 0.607 |
| opioid use disorder – control | -1.2648 | 0.4707 | -2.6869 | 0.013 |
| opioid+alcohol use disorder – control | -1.4566 | 0.5567 | -2.6167 | 0.015 |
| Age Cateogry: | | | | |
| 50-64 – 30-49 | -0.1991 | 0.5015 | -0.3969 | 0.695 |
| 50-65 – 30-49 | 0.1550 | 0.7705 | 0.2011 | 0.842 |
| 65+ – 30-49 | -0.1481 | 0.5811 | -0.2548 | 0.801 |
| <30 – 30-49 | 0.2484 | 0.6075 | 0.4089 | 0.686 |
| Ethnicity: | | | | |
| Asian – White | -0.0203 | 1.0116 | -0.0201 | 0.984 |
| Black – White | -0.5336 | 0.4118 | -1.2956 | 0.207 |
| Hispanic – White | -0.8877 | 0.7876 | -1.1270 | 0.271 |
| Gender: | | | | |
| Male – Female | -0.6024 | 0.4868 | -1.2375 | 0.228 |
| PMI converted to hours | 0.0602 | 0.0202 | 2.9806 | 0.006 |
| pH | 0.8574 | 0.6983 | 1.2279 | 0.231 |

[a] Represents reference level

#### Linear Regression

Model Fit Measures

| Model | R | $R^2$ |
|---|---|---|
| 1 | 0.773 | 0.598 |

Model Coefficients - CDH13

| Predictor | Estimate | SE | t | p |
|---|---|---|---|---|
| Intercept [a] | 24.3337 | 1.57175 | 15.482 | < .001 |
| Arm: | | | | |
| alcohol use disorder – control | -0.0896 | 0.16539 | -0.542 | 0.593 |
| opioid use disorder – control | 0.3291 | 0.15471 | 2.127 | 0.044 |
| opioid+alcohol use disorder – control | -0.1253 | 0.18296 | -0.685 | 0.500 |
| Age Cateogry: | | | | |
| 50-64 – 30-49 | -0.3083 | 0.16484 | -1.870 | 0.074 |
| 50-65 – 30-49 | 0.1828 | 0.25324 | 0.722 | 0.477 |
| 65+ – 30-49 | 0.0694 | 0.19097 | 0.364 | 0.719 |
| <30 – 30-49 | -0.3040 | 0.19967 | -1.523 | 0.141 |
| Ethnicity: | | | | |
| Asian – White | -0.3620 | 0.33247 | -1.089 | 0.287 |
| Black – White | 0.0240 | 0.13535 | 0.177 | 0.861 |
| Hispanic – White | -0.4256 | 0.25887 | -1.644 | 0.113 |
| Gender: | | | | |
| Male – Female | 0.1485 | 0.15999 | 0.928 | 0.363 |
| PMI converted to hours | 0.0218 | 0.00664 | 3.278 | 0.003 |
| pH | 0.4938 | 0.22949 | 2.152 | 0.042 |

[a] Represents reference level

#### Linear Regression

Model Fit Measures

| Model | R | $R^2$ |
|---|---|---|
| 1 | 0.666 | 0.443 |

Model Coefficients - PITRM1

| Predictor | Estimate | SE | t | p |
|---|---|---|---|---|
| Intercept [a] | 28.7977 | 5.4106 | 5.322 | < .001 |
| Arm: | | | | |
| alcohol use disorder – control | 1.0372 | 0.5693 | 1.822 | 0.081 |
| opioid use disorder – control | 1.8149 | 0.5326 | 3.408 | 0.002 |
| opioid+alcohol use disorder – control | 0.9408 | 0.6298 | 1.494 | 0.148 |
| Age Cateogry: | | | | |
| 50-64 – 30-49 | 0.2209 | 0.5675 | 0.389 | 0.700 |
| 50-65 – 30-49 | -0.2786 | 0.8718 | -0.320 | 0.752 |
| 65+ – 30-49 | -0.2611 | 0.6574 | -0.397 | 0.695 |
| <30 – 30-49 | -0.6173 | 0.6873 | -0.898 | 0.378 |
| Ethnicity: | | | | |
| Asian – White | 1.5894 | 1.1445 | 1.389 | 0.178 |
| Black – White | 0.5786 | 0.4659 | 1.242 | 0.226 |
| Hispanic – White | 0.3790 | 0.8911 | 0.425 | 0.674 |
| Gender: | | | | |
| Male – Female | 0.5513 | 0.5507 | 1.001 | 0.327 |
| PMI converted to hours | 0.0134 | 0.0229 | 0.588 | 0.562 |
| pH | -0.5216 | 0.7900 | -0.660 | 0.515 |

[a] Represents reference level

#### Linear Regression

Model Fit Measures

| Model | R | $R^2$ |
|---|---|---|
| 1 | 0.624 | 0.389 |

Model Coefficients - OGDH

| Predictor | Estimate | SE | t | p |
|---|---|---|---|---|
| Intercept [a] | 29.40336 | 1.25694 | 23.3928 | < .001 |
| Arm: | | | | |
| alcohol use disorder – control | -0.16769 | 0.13226 | -1.2679 | 0.217 |
| opioid use disorder – control | -0.17195 | 0.12372 | -1.3898 | 0.177 |
| opioid+alcohol use disorder – control | 0.08716 | 0.14631 | 0.5957 | 0.557 |
| Age Cateogry: | | | | |
| 50-64 – 30-49 | 0.05086 | 0.13183 | 0.3858 | 0.703 |
| 50-65 – 30-49 | 0.10762 | 0.20252 | 0.5314 | 0.600 |
| 65+ – 30-49 | 0.21124 | 0.15272 | 1.3832 | 0.179 |
| <30 – 30-49 | 0.13206 | 0.15967 | 0.8271 | 0.416 |
| Ethnicity: | | | | |
| Asian – White | 0.15420 | 0.26588 | 0.5800 | 0.567 |
| Black – White | 0.00219 | 0.10824 | 0.0203 | 0.984 |
| Hispanic – White | 0.33681 | 0.20702 | 1.6270 | 0.117 |
| Gender: | | | | |
| Male – Female | 0.12448 | 0.12794 | 0.9729 | 0.340 |
| PMI converted to hours | -0.00489 | 0.00531 | -0.9204 | 0.367 |
| pH | 0.16304 | 0.18353 | 0.8883 | 0.383 |

[a] Represents reference level

#### Linear Regression

Model Fit Measures

| Model | R | $R^2$ |
|---|---|---|
| 1 | 0.610 | 0.372 |

Model Coefficients - ASPH

| Predictor | Estimate | SE | t | p |
|---|---|---|---|---|
| Intercept [a] | 24.39083 | 3.3148 | 7.35821 | < .001 |
| Arm: | | | | |
| alcohol use disorder – control | -0.20627 | 0.3488 | -0.59138 | 0.560 |
| opioid use disorder – control | 0.50106 | 0.3263 | 1.53567 | 0.138 |
| opioid+alcohol use disorder – control | -0.29866 | 0.3859 | -0.77403 | 0.446 |
| Age Cateogry: | | | | |
| 50-64 – 30-49 | -0.36747 | 0.3476 | -1.05702 | 0.301 |
| 50-65 – 30-49 | -0.50891 | 0.5341 | -0.95287 | 0.350 |
| 65+ – 30-49 | 0.25920 | 0.4028 | 0.64356 | 0.526 |
| <30 – 30-49 | -0.48060 | 0.4211 | -1.14133 | 0.265 |
| Ethnicity: | | | | |
| Asian – White | -0.46727 | 0.7012 | -0.66641 | 0.512 |
| Black – White | 0.00141 | 0.2855 | 0.00494 | 0.996 |
| Hispanic – White | 0.08055 | 0.5459 | 0.14754 | 0.884 |
| Gender: | | | | |
| Male – Female | 0.06227 | 0.3374 | 0.18456 | 0.855 |
| PMI converted to hours | 0.00192 | 0.0140 | 0.13725 | 0.892 |
| pH | -0.14200 | 0.4840 | -0.29339 | 0.772 |

[a] Represents reference level

#### Linear Regression

Model Fit Measures

| Model | R | $R^2$ |
|---|---|---|
| 1 | 0.595 | 0.354 |

Model Coefficients - ALCAM

| Predictor | Estimate | SE | t | p |
|---|---|---|---|---|
| Intercept [a] | 26.44951 | 1.39889 | 18.9075 | < .001 |
| Arm: | | | | |
| alcohol use disorder – control | 0.02390 | 0.14720 | 0.1624 | 0.872 |
| opioid use disorder – control | 0.23518 | 0.13770 | 1.7079 | 0.101 |
| opioid+alcohol use disorder – control | -0.01141 | 0.16284 | -0.0701 | 0.945 |
| Age Cateogry: | | | | |
| 50-64 – 30-49 | -0.19115 | 0.14671 | -1.3029 | 0.205 |
| 50-65 – 30-49 | 0.05365 | 0.22539 | 0.2381 | 0.814 |
| 65+ – 30-49 | 0.09874 | 0.16997 | 0.5809 | 0.567 |
| <30 – 30-49 | -0.16413 | 0.17771 | -0.9236 | 0.365 |
| Ethnicity: | | | | |
| Asian – White | -0.09540 | 0.29591 | -0.3224 | 0.750 |
| Black – White | 0.05907 | 0.12047 | 0.4904 | 0.628 |
| Hispanic – White | -0.11110 | 0.23040 | -0.4822 | 0.634 |
| Gender: | | | | |
| Male – Female | 0.09480 | 0.14239 | 0.6658 | 0.512 |
| PMI converted to hours | 0.00636 | 0.00591 | 1.0757 | 0.293 |
| pH | 0.25041 | 0.20425 | 1.2260 | 0.232 |

[a] Represents reference level

#### Linear Regression

Model Fit Measures

| Model | R | $R^2$ |
|---|---|---|
| 1 | 0.743 | 0.553 |

Model Coefficients - VPS13C

| Predictor | Estimate | SE | t | p |
|---|---|---|---|---|
| Intercept [a] | 19.5235 | 7.5683 | 2.580 | 0.016 |
| Arm: | | | | |
| alcohol use disorder – control | -0.7896 | 0.7964 | -0.992 | 0.331 |
| opioid use disorder – control | -1.9902 | 0.7450 | -2.672 | 0.013 |
| opioid+alcohol use disorder – control | 0.6962 | 0.8810 | 0.790 | 0.437 |
| Age Cateogry: | | | | |
| 50-64 – 30-49 | 0.2158 | 0.7937 | 0.272 | 0.788 |
| 50-65 – 30-49 | 1.4869 | 1.2194 | 1.219 | 0.235 |
| 65+ – 30-49 | 0.2558 | 0.9196 | 0.278 | 0.783 |
| <30 – 30-49 | 1.5170 | 0.9614 | 1.578 | 0.128 |
| Ethnicity: | | | | |
| Asian – White | 2.1086 | 1.6009 | 1.317 | 0.200 |
| Black – White | -0.4635 | 0.6517 | -0.711 | 0.484 |
| Hispanic – White | 1.4480 | 1.2465 | 1.162 | 0.257 |
| Gender: | | | | |
| Male – Female | -1.0285 | 0.7704 | -1.335 | 0.194 |
| PMI converted to hours | -0.0619 | 0.0320 | -1.936 | 0.065 |
| pH | 1.2457 | 1.1051 | 1.127 | 0.271 |

[a] Represents reference level

### Linear Regression

Model Fit Measures

| Model | R | $R^2$ |
|---|---|---|
| 1 | 0.717 | 0.514 |

Model Coefficients - SORD

| Predictor | Estimate | SE | t | p |
|---|---|---|---|---|
| Intercept [a] | 28.32557 | 2.7891 | 10.156 | < .001 |
| Arm: | | | | |
| alcohol use disorder – control | -0.72778 | 0.2935 | -2.480 | 0.021 |
| opioid use disorder – control | -0.57480 | 0.2745 | -2.094 | 0.047 |
| opioid+alcohol use disorder – control | -0.23625 | 0.3247 | -0.728 | 0.474 |
| Age Cateogry: | | | | |
| 50-64 – 30-49 | -0.16501 | 0.2925 | -0.564 | 0.578 |
| 50-65 – 30-49 | -1.54143 | 0.4494 | -3.430 | 0.002 |
| 65+ – 30-49 | 0.33468 | 0.3389 | 0.988 | 0.333 |
| <30 – 30-49 | -0.12331 | 0.3543 | -0.348 | 0.731 |
| Ethnicity: | | | | |
| Asian – White | -0.10838 | 0.5900 | -0.184 | 0.856 |
| Black – White | -0.05687 | 0.2402 | -0.237 | 0.815 |
| Hispanic – White | -0.07743 | 0.4594 | -0.169 | 0.868 |
| Gender: | | | | |
| Male – Female | 0.13070 | 0.2839 | 0.460 | 0.649 |
| PMI converted to hours | -0.00788 | 0.0118 | -0.669 | 0.510 |
| pH | -0.09822 | 0.4072 | -0.241 | 0.811 |

[a] Represents reference level

### Linear Regression

Model Fit Measures

| Model | R | $R^2$ |
|---|---|---|
| 1 | 0.735 | 0.541 |

Model Coefficients - GNB1

| Predictor | Estimate | SE | t | p |
|---|---|---|---|---|
| Intercept [a] | 33.16207 | 0.98890 | 33.5341 | < .001 |
| Arm: | | | | |
| alcohol use disorder – control | 0.10709 | 0.10406 | 1.0291 | 0.314 |
| opioid use disorder – control | 0.29901 | 0.09734 | 3.0718 | 0.005 |
| opioid+alcohol use disorder – control | 0.12032 | 0.11511 | 1.0452 | 0.306 |
| Age Cateogry: | | | | |
| 50-64 – 30-49 | -0.07501 | 0.10371 | -0.7232 | 0.477 |
| 50-65 – 30-49 | -0.04859 | 0.15933 | -0.3049 | 0.763 |
| 65+ – 30-49 | -0.27767 | 0.12016 | -2.3109 | 0.030 |
| <30 – 30-49 | -0.16651 | 0.12563 | -1.3254 | 0.198 |
| Ethnicity: | | | | |
| Asian – White | 0.04369 | 0.20918 | 0.2088 | 0.836 |
| Black – White | 0.00459 | 0.08516 | 0.0539 | 0.957 |
| Hispanic – White | 0.00528 | 0.16287 | 0.0324 | 0.974 |
| Gender: | | | | |
| Male – Female | 0.17818 | 0.10066 | 1.7701 | 0.089 |
| PMI converted to hours | 0.00487 | 0.00418 | 1.1669 | 0.255 |
| pH | 0.13628 | 0.14439 | 0.9439 | 0.355 |

[a] Represents reference level

#### Linear Regression

Model Fit Measures

| Model | R | $R^2$ |
|---|---|---|
| 1 | 0.764 | 0.583 |

Model Coefficients - SIRPA

| Predictor | Estimate | SE | t | p |
|---|---|---|---|---|
| Intercept [a] | 31.73640 | 0.97993 | 32.38638 | < .001 |
| Arm: | | | | |
| alcohol use disorder – control | 0.13725 | 0.10311 | 1.33103 | 0.196 |
| opioid use disorder – control | 0.33751 | 0.09646 | 3.49900 | 0.002 |
| opioid+alcohol use disorder – control | -0.00661 | 0.11407 | -0.05799 | 0.954 |
| Age Cateogry: | | | | |
| 50-64 – 30-49 | 0.02450 | 0.10277 | 0.23844 | 0.814 |
| 50-65 – 30-49 | 0.33275 | 0.15789 | 2.10750 | 0.046 |
| 65+ – 30-49 | -0.06761 | 0.11907 | -0.56783 | 0.575 |
| <30 – 30-49 | 0.00109 | 0.12449 | 0.00872 | 0.993 |
| Ethnicity: | | | | |
| Asian – White | 0.14084 | 0.20728 | 0.67944 | 0.503 |
| Black – White | -0.08558 | 0.08439 | -1.01410 | 0.321 |
| Hispanic – White | -0.22814 | 0.16139 | -1.41357 | 0.170 |
| Gender: | | | | |
| Male – Female | -0.01935 | 0.09975 | -0.19398 | 0.848 |
| PMI converted to hours | 0.00108 | 0.00414 | 0.26021 | 0.797 |
| pH | 0.12057 | 0.14308 | 0.84266 | 0.408 |

[a] Represents reference level

#### Linear Regression

Model Fit Measures

| Model | R | $R^2$ |
|---|---|---|
| 1 | 0.690 | 0.476 |

Model Coefficients - EEF1A2

| Predictor | Estimate | SE | t | p |
|---|---|---|---|---|
| Intercept [a] | 32.49167 | 1.64709 | 19.7267 | < .001 |
| Arm: | | | | |
| alcohol use disorder – control | -0.11543 | 0.17331 | -0.6660 | 0.512 |
| opioid use disorder – control | -0.30337 | 0.16213 | -1.8712 | 0.074 |
| opioid+alcohol use disorder – control | 0.34735 | 0.19173 | 1.8117 | 0.083 |
| Age Cateogry: | | | | |
| 50-64 – 30-49 | 0.01239 | 0.17274 | 0.0717 | 0.943 |
| 50-65 – 30-49 | -0.11488 | 0.26538 | -0.4329 | 0.669 |
| 65+ – 30-49 | 0.12841 | 0.20013 | 0.6417 | 0.527 |
| <30 – 30-49 | -0.04576 | 0.20924 | -0.2187 | 0.829 |
| Ethnicity: | | | | |
| Asian – White | 0.16587 | 0.34841 | 0.4761 | 0.638 |
| Black – White | 0.14298 | 0.14184 | 1.0081 | 0.323 |
| Hispanic – White | 0.21344 | 0.27127 | 0.7868 | 0.439 |
| Gender: | | | | |
| Male – Female | -0.00398 | 0.16766 | -0.0238 | 0.981 |
| PMI converted to hours | -0.00882 | 0.00696 | -1.2679 | 0.217 |
| pH | -0.10977 | 0.24049 | -0.4565 | 0.652 |

[a] Represents reference level

### Linear Regression

Model Fit Measures

| Model | R | $R^2$ |
|---|---|---|
| 1 | 0.657 | 0.432 |

Model Coefficients - AUH (2)

| Predictor | Estimate | SE | t | p |
|---|---|---|---|---|
| Intercept [a] | 30.2022 | 1.01670 | 29.7061 | < .001 |
| Arm: | | | | |
| alcohol use disorder – control | 0.2371 | 0.10698 | 2.2163 | 0.036 |
| opioid use disorder – control | 0.1383 | 0.10008 | 1.3819 | 0.180 |
| opioid+alcohol use disorder – control | -0.1971 | 0.11835 | -1.6658 | 0.109 |
| Age Cateogry: | | | | |
| 50-64 – 30-49 | 0.0364 | 0.10663 | 0.3416 | 0.736 |
| 50-65 – 30-49 | -0.0198 | 0.16381 | -0.1207 | 0.905 |
| 65+ – 30-49 | -0.0151 | 0.12353 | -0.1223 | 0.904 |
| <30 – 30-49 | 0.1506 | 0.12916 | 1.1661 | 0.255 |
| Ethnicity: | | | | |
| Asian – White | -0.0511 | 0.21506 | -0.2377 | 0.814 |
| Black – White | -0.1382 | 0.08755 | -1.5784 | 0.128 |
| Hispanic – White | -0.1426 | 0.16745 | -0.8519 | 0.403 |
| Gender: | | | | |
| Male – Female | -0.1480 | 0.10349 | -1.4305 | 0.165 |
| PMI converted to hours | 1.18e-4 | 0.00429 | 0.0275 | 0.978 |
| pH | -0.2281 | 0.14845 | -1.5367 | 0.137 |

[a] Represents reference level

### Linear Regression

Model Fit Measures

| Model | R | $R^2$ |
|---|---|---|
| 1 | 0.721 | 0.520 |

Model Coefficients - CRMP1

| Predictor | Estimate | SE | t | p |
|---|---|---|---|---|
| Intercept [a] | 32.64969 | 0.71857 | 45.4372 | < .001 |
| Arm: | | | | |
| alcohol use disorder – control | -0.05058 | 0.07561 | -0.6689 | 0.510 |
| opioid use disorder – control | 0.16399 | 0.07073 | 2.3185 | 0.029 |
| opioid+alcohol use disorder – control | 0.00169 | 0.08364 | 0.0201 | 0.984 |
| Age Cateogry: | | | | |
| 50-64 – 30-49 | -0.08713 | 0.07536 | -1.1561 | 0.259 |
| 50-65 – 30-49 | -0.01650 | 0.11578 | -0.1425 | 0.888 |
| 65+ – 30-49 | -0.01105 | 0.08731 | -0.1266 | 0.900 |
| <30 – 30-49 | -0.13080 | 0.09128 | -1.4330 | 0.165 |
| Ethnicity: | | | | |
| Asian – White | 0.04583 | 0.15200 | 0.3015 | 0.766 |
| Black – White | 0.04716 | 0.06188 | 0.7621 | 0.453 |
| Hispanic – White | -0.08741 | 0.11835 | -0.7386 | 0.467 |
| Gender: | | | | |
| Male – Female | -0.06111 | 0.07314 | -0.8354 | 0.412 |
| PMI converted to hours | 0.00572 | 0.00303 | 1.8859 | 0.071 |
| pH | 0.03383 | 0.10492 | 0.3225 | 0.750 |

[a] Represents reference level

### Linear Regression

Model Fit Measures

| Model | R | $R^2$ |
|---|---|---|
| 1 | 0.847 | 0.717 |

Model Coefficients - EIF4A2

| Predictor | Estimate | SE | t | p |
|---|---|---|---|---|
| Intercept [a] | 30.26094 | 0.72271 | 41.871 | < .001 |
| Arm: | | | | |
| alcohol use disorder – control | -0.10130 | 0.07605 | -1.332 | 0.195 |
| opioid use disorder – control | -0.16913 | 0.07114 | -2.377 | 0.026 |
| opioid+alcohol use disorder – control | 0.11856 | 0.08413 | 1.409 | 0.172 |
| Age Cateogry: | | | | |
| 50-64 – 30-49 | 0.14381 | 0.07580 | 1.897 | 0.070 |
| 50-65 – 30-49 | -0.20269 | 0.11644 | -1.741 | 0.095 |
| 65+ – 30-49 | 0.23996 | 0.08781 | 2.733 | 0.012 |
| <30 – 30-49 | 0.08076 | 0.09181 | 0.880 | 0.388 |
| Ethnicity: | | | | |
| Asian – White | 0.31723 | 0.15287 | 2.075 | 0.049 |
| Black – White | 0.04107 | 0.06224 | 0.660 | 0.516 |
| Hispanic – White | -0.11892 | 0.11903 | -0.999 | 0.328 |
| Gender: | | | | |
| Male – Female | -0.03703 | 0.07357 | -0.503 | 0.619 |
| PMI converted to hours | 0.00151 | 0.00305 | 0.493 | 0.626 |
| pH | -0.09606 | 0.10552 | -0.910 | 0.372 |

[a] Represents reference level

### Linear Regression

Model Fit Measures

| Model | R | $R^2$ |
|---|---|---|
| 1 | 0.832 | 0.692 |

Model Coefficients - TCEB1

| Predictor | Estimate | SE | t | p |
|---|---|---|---|---|
| Intercept [a] | 27.46039 | 2.5008 | 10.9805 | < .001 |
| Arm: | | | | |
| alcohol use disorder – control | -0.29783 | 0.2631 | -1.1318 | 0.269 |
| opioid use disorder – control | -0.73950 | 0.2462 | -3.0041 | 0.006 |
| opioid+alcohol use disorder – control | 0.44994 | 0.2911 | 1.5456 | 0.135 |
| Age Cateogry: | | | | |
| 50-64 – 30-49 | 0.36254 | 0.2623 | 1.3823 | 0.180 |
| 50-65 – 30-49 | -0.36266 | 0.4029 | -0.9001 | 0.377 |
| 65+ – 30-49 | 0.22730 | 0.3039 | 0.7480 | 0.462 |
| <30 – 30-49 | 0.00587 | 0.3177 | 0.0185 | 0.985 |
| Ethnicity: | | | | |
| Asian – White | 0.24855 | 0.5290 | 0.4698 | 0.643 |
| Black – White | 0.32293 | 0.2154 | 1.4995 | 0.147 |
| Hispanic – White | 0.87357 | 0.4119 | 2.1209 | 0.044 |
| Gender: | | | | |
| Male – Female | -0.06368 | 0.2546 | -0.2502 | 0.805 |
| PMI converted to hours | -0.04226 | 0.0106 | -4.0011 | < .001 |
| pH | 0.05876 | 0.3652 | 0.1609 | 0.873 |

[a] Represents reference level

### Linear Regression

Model Fit Measures

| Model | R | $R^2$ |
|---|---|---|
| 1 | 0.845 | 0.714 |

Model Coefficients - UBE2V2

| Predictor | Estimate | SE | t | p |
|---|---|---|---|---|
| Intercept [a] | 28.06422 | 1.75306 | 16.0087 | < .001 |
| Arm: | | | | |
| alcohol use disorder – control | -0.00541 | 0.18446 | -0.0293 | 0.977 |
| opioid use disorder – control | 0.45231 | 0.17256 | 2.6212 | 0.015 |
| opioid+alcohol use disorder – control | -0.49518 | 0.20406 | -2.4266 | 0.023 |
| Age Cateogry: | | | | |
| 50-64 – 30-49 | -0.40030 | 0.18386 | -2.1772 | 0.040 |
| 50-65 – 30-49 | -0.43615 | 0.28245 | -1.5441 | 0.136 |
| 65+ – 30-49 | -0.00995 | 0.21300 | -0.0467 | 0.963 |
| <30 – 30-49 | -0.23304 | 0.22270 | -1.0464 | 0.306 |
| Ethnicity: | | | | |
| Asian – White | -1.00582 | 0.37082 | -2.7124 | 0.012 |
| Black – White | -0.12033 | 0.15097 | -0.7970 | 0.433 |
| Hispanic – White | 0.02016 | 0.28873 | 0.0698 | 0.945 |
| Gender: | | | | |
| Male – Female | 0.04877 | 0.17844 | 0.2733 | 0.787 |
| PMI converted to hours | 0.01606 | 0.00740 | 2.1692 | 0.040 |
| pH | 0.08162 | 0.25597 | 0.3188 | 0.753 |

[a] Represents reference level

### Linear Regression

Model Fit Measures

| Model | R | $R^2$ |
|---|---|---|
| 1 | 0.671 | 0.450 |

Model Coefficients - NEDD8

| Predictor | Estimate | SE | t | p |
|---|---|---|---|---|
| Intercept [a] | 25.5548 | 5.0190 | 5.092 | < .001 |
| Arm: | | | | |
| alcohol use disorder – control | 0.6587 | 0.5281 | 1.247 | 0.224 |
| opioid use disorder – control | 0.6561 | 0.4940 | 1.328 | 0.197 |
| opioid+alcohol use disorder – control | -1.3055 | 0.5842 | -2.235 | 0.035 |
| Age Cateogry: | | | | |
| 50-64 – 30-49 | 0.5785 | 0.5264 | 1.099 | 0.283 |
| 50-65 – 30-49 | 0.5718 | 0.8087 | 0.707 | 0.486 |
| 65+ – 30-49 | 0.1989 | 0.6098 | 0.326 | 0.747 |
| <30 – 30-49 | 0.4818 | 0.6376 | 0.756 | 0.457 |
| Ethnicity: | | | | |
| Asian – White | 0.7264 | 1.0617 | 0.684 | 0.500 |
| Black – White | -0.2495 | 0.4322 | -0.577 | 0.569 |
| Hispanic – White | 0.1797 | 0.8266 | 0.217 | 0.830 |
| Gender: | | | | |
| Male – Female | -0.5835 | 0.5109 | -1.142 | 0.265 |
| PMI converted to hours | 0.0181 | 0.0212 | 0.855 | 0.401 |
| pH | 0.3551 | 0.7328 | 0.485 | 0.632 |

[a] Represents reference level

#### Linear Regression

Model Fit Measures

| Model | R | $R^2$ |
|---|---|---|
| 1 | 0.498 | 0.248 |

Model Coefficients - COPS6

| Predictor | Estimate | SE | t | p |
|---|---|---|---|---|
| Intercept [a] | 28.0818 | 3.5374 | 7.9385 | < .001 |
| Arm: | | | | |
| alcohol use disorder – control | 0.0754 | 0.3722 | 0.2025 | 0.841 |
| opioid use disorder – control | -0.3532 | 0.3482 | -1.0143 | 0.321 |
| opioid+alcohol use disorder – control | 0.4695 | 0.4118 | 1.1402 | 0.265 |
| Age Cateogry: | | | | |
| 50-64 – 30-49 | -0.2160 | 0.3710 | -0.5823 | 0.566 |
| 50-65 – 30-49 | 0.0155 | 0.5700 | 0.0272 | 0.978 |
| 65+ – 30-49 | -0.2338 | 0.4298 | -0.5439 | 0.592 |
| <30 – 30-49 | 0.0791 | 0.4494 | 0.1761 | 0.862 |
| Ethnicity: | | | | |
| Asian – White | 0.4837 | 0.7483 | 0.6464 | 0.524 |
| Black – White | 0.3110 | 0.3046 | 1.0210 | 0.317 |
| Hispanic – White | 0.3039 | 0.5826 | 0.5216 | 0.607 |
| Gender: | | | | |
| Male – Female | -0.2635 | 0.3601 | -0.7317 | 0.471 |
| PMI converted to hours | -0.0104 | 0.0149 | -0.6982 | 0.492 |
| pH | -0.1542 | 0.5165 | -0.2985 | 0.768 |

[a] Represents reference level

#### Linear Regression

Model Fit Measures

| Model | R | $R^2$ |
|---|---|---|
| 1 | 0.733 | 0.538 |

Model Coefficients - RDH11 (2)

| Predictor | Estimate | SE | t | p |
|---|---|---|---|---|
| Intercept [a] | 24.4989 | 4.2842 | 5.718 | < .001 |
| Arm: | | | | |
| alcohol use disorder – control | -0.8373 | 0.4508 | -1.857 | 0.076 |
| opioid use disorder – control | -1.0561 | 0.4217 | -2.504 | 0.019 |
| opioid+alcohol use disorder – control | -1.1732 | 0.4987 | -2.353 | 0.027 |
| Age Cateogry: | | | | |
| 50-64 – 30-49 | 0.6720 | 0.4493 | 1.496 | 0.148 |
| 50-65 – 30-49 | -0.1700 | 0.6903 | -0.246 | 0.808 |
| 65+ – 30-49 | 0.2782 | 0.5205 | 0.534 | 0.598 |
| <30 – 30-49 | 0.2305 | 0.5442 | 0.424 | 0.676 |
| Ethnicity: | | | | |
| Asian – White | 0.5008 | 0.9062 | 0.553 | 0.586 |
| Black – White | 0.2768 | 0.3689 | 0.750 | 0.460 |
| Hispanic – White | 0.8776 | 0.7056 | 1.244 | 0.226 |
| Gender: | | | | |
| Male – Female | -0.9047 | 0.4361 | -2.075 | 0.049 |
| PMI converted to hours | -0.0124 | 0.0181 | -0.688 | 0.498 |
| pH | 0.1899 | 0.6255 | 0.304 | 0.764 |

[a] Represents reference level

### Linear Regression

Model Fit Measures

| Model | R | $R^2$ |
|---|---|---|
| 1 | 0.695 | 0.482 |

Model Coefficients - MAL2

| Predictor | Estimate | SE | t | p |
|---|---|---|---|---|
| Intercept [a] | 19.96038 | 7.4295 | 2.687 | 0.013 |
| Arm: | | | | |
| alcohol use disorder – control | 1.54932 | 0.7818 | 1.982 | 0.059 |
| opioid use disorder – control | 1.61711 | 0.7313 | 2.211 | 0.037 |
| opioid+alcohol use disorder – control | 0.61553 | 0.8648 | 0.712 | 0.483 |
| Age Cateogry: | | | | |
| 50-64 – 30-49 | -0.13380 | 0.7792 | -0.172 | 0.865 |
| 50-65 – 30-49 | -0.55502 | 1.1970 | -0.464 | 0.647 |
| 65+ – 30-49 | 0.54833 | 0.9027 | 0.607 | 0.549 |
| <30 – 30-49 | 0.25025 | 0.9438 | 0.265 | 0.793 |
| Ethnicity: | | | | |
| Asian – White | -2.13822 | 1.5715 | -1.361 | 0.186 |
| Black – White | 0.08792 | 0.6398 | 0.137 | 0.892 |
| Hispanic – White | 1.36873 | 1.2236 | 1.119 | 0.274 |
| Gender: | | | | |
| Male – Female | -0.57034 | 0.7562 | -0.754 | 0.458 |
| PMI converted to hours | -0.00319 | 0.0314 | -0.102 | 0.920 |
| pH | 0.97404 | 1.0848 | 0.898 | 0.378 |

[a] Represents reference level

### Linear Regression

Model Fit Measures

| Model | R | $R^2$ |
|---|---|---|
| 1 | 0.718 | 0.515 |

Model Coefficients - NRCAM

| Predictor | Estimate | SE | t | p |
|---|---|---|---|---|
| Intercept [a] | 32.14171 | 0.76798 | 41.852 | < .001 |
| Arm: | | | | |
| alcohol use disorder – control | 0.05604 | 0.08081 | 0.694 | 0.495 |
| opioid use disorder – control | 0.19568 | 0.07559 | 2.589 | 0.016 |
| opioid+alcohol use disorder – control | -0.04446 | 0.08940 | -0.497 | 0.623 |
| Age Categry: | | | | |
| 50-64 – 30-49 | -0.02575 | 0.08054 | -0.320 | 0.752 |
| 50-65 – 30-49 | 0.01333 | 0.12374 | 0.108 | 0.915 |
| 65+ – 30-49 | -0.01156 | 0.09331 | -0.124 | 0.902 |
| <30 – 30-49 | 0.07639 | 0.09756 | 0.783 | 0.441 |
| Ethnicity: | | | | |
| Asian – White | 0.02072 | 0.16245 | 0.128 | 0.900 |
| Black – White | -0.07165 | 0.06613 | -1.083 | 0.289 |
| Hispanic – White | -0.25714 | 0.12649 | -2.033 | 0.053 |
| Gender: | | | | |
| Male – Female | -0.06176 | 0.07817 | -0.790 | 0.437 |
| PMI converted to hours | 0.00582 | 0.00324 | 1.794 | 0.085 |
| pH | -0.08486 | 0.11213 | -0.757 | 0.457 |

[a] Represents reference level

### Linear Regression

Model Fit Measures

| Model | R | $R^2$ |
|---|---|---|
| 1 | 0.705 | 0.497 |

Model Coefficients - SCCPDH

| Predictor | Estimate | SE | t | p |
|---|---|---|---|---|
| Intercept [a] | 29.97220 | 1.19372 | 25.1083 | < .001 |
| Arm: | | | | |
| alcohol use disorder – control | 0.06225 | 0.12561 | 0.4956 | 0.625 |
| opioid use disorder – control | 0.13828 | 0.11750 | 1.1769 | 0.251 |
| opioid+alcohol use disorder – control | -0.29484 | 0.13895 | -2.1219 | 0.044 |
| Age Categry: | | | | |
| 50-64 – 30-49 | 0.05650 | 0.12519 | 0.4513 | 0.656 |
| 50-65 – 30-49 | -0.20404 | 0.19233 | -1.0609 | 0.299 |
| 65+ – 30-49 | -0.11684 | 0.14504 | -0.8056 | 0.428 |
| <30 – 30-49 | 0.01905 | 0.15164 | 0.1256 | 0.901 |
| Ethnicity: | | | | |
| Asian – White | -0.37621 | 0.25250 | -1.4899 | 0.149 |
| Black – White | 0.00180 | 0.10280 | 0.0175 | 0.986 |
| Hispanic – White | -0.47340 | 0.19660 | -2.4079 | 0.024 |
| Gender: | | | | |
| Male – Female | -0.07060 | 0.12151 | -0.5810 | 0.567 |
| PMI converted to hours | 0.00141 | 0.00504 | 0.2793 | 0.782 |
| pH | -0.11737 | 0.17430 | -0.6734 | 0.507 |

[a] Represents reference level

### Linear Regression

Model Fit Measures

| Model | R | $R^2$ |
|---|---|---|
| 1 | 0.783 | 0.613 |

Model Coefficients - CPNE4

| Predictor | Estimate | SE | t | p |
|---|---|---|---|---|
| Intercept [a] | 21.0534 | 3.4433 | 6.114 | < .001 |
| Arm: | | | | |
| alcohol use disorder – control | -0.3564 | 0.3623 | -0.984 | 0.335 |
| opioid use disorder – control | 0.5084 | 0.3389 | 1.500 | 0.147 |
| opioid+alcohol use disorder – control | -0.4556 | 0.4008 | -1.137 | 0.267 |
| Age Cateogry: | | | | |
| 50-64 – 30-49 | -0.7951 | 0.3611 | -2.202 | 0.038 |
| 50-65 – 30-49 | -1.7913 | 0.5548 | -3.229 | 0.004 |
| 65+ – 30-49 | -0.5427 | 0.4184 | -1.297 | 0.207 |
| <30 – 30-49 | -0.8560 | 0.4374 | -1.957 | 0.062 |
| Ethnicity: | | | | |
| Asian – White | -0.1753 | 0.7284 | -0.241 | 0.812 |
| Black – White | -0.2100 | 0.2965 | -0.708 | 0.486 |
| Hispanic – White | -0.3422 | 0.5671 | -0.603 | 0.552 |
| Gender: | | | | |
| Male – Female | 0.6546 | 0.3505 | 1.867 | 0.074 |
| PMI converted to hours | 0.0321 | 0.0145 | 2.208 | 0.037 |
| pH | 0.4173 | 0.5028 | 0.830 | 0.415 |

[a] Represents reference level

### Linear Regression

Model Fit Measures

| Model | R | $R^2$ |
|---|---|---|
| 1 | 0.689 | 0.474 |

Model Coefficients - ISOC2

| Predictor | Estimate | SE | t | p |
|---|---|---|---|---|
| Intercept [a] | 27.40451 | 2.27354 | 12.054 | < .001 |
| Arm: | | | | |
| alcohol use disorder – control | 0.25871 | 0.23923 | 1.081 | 0.290 |
| opioid use disorder – control | 0.45261 | 0.22379 | 2.022 | 0.054 |
| opioid+alcohol use disorder – control | -0.02656 | 0.26465 | -0.100 | 0.921 |
| Age Cateogry: | | | | |
| 50-64 – 30-49 | -0.26418 | 0.23844 | -1.108 | 0.279 |
| 50-65 – 30-49 | -0.53305 | 0.36631 | -1.455 | 0.159 |
| 65+ – 30-49 | -0.27002 | 0.27624 | -0.977 | 0.338 |
| <30 – 30-49 | -0.41080 | 0.28882 | -1.422 | 0.168 |
| Ethnicity: | | | | |
| Asian – White | -0.52320 | 0.48092 | -1.088 | 0.287 |
| Black – White | 0.09472 | 0.19579 | 0.484 | 0.633 |
| Hispanic – White | -0.11889 | 0.37445 | -0.318 | 0.754 |
| Gender: | | | | |
| Male – Female | 0.35124 | 0.23142 | 1.518 | 0.142 |
| PMI converted to hours | 0.00841 | 0.00960 | 0.875 | 0.390 |
| pH | 0.08134 | 0.33196 | 0.245 | 0.809 |

[a] Represents reference level

### Linear Regression

Model Fit Measures

| Model | R | $R^2$ |
|---|---|---|
| 1 | 0.678 | 0.460 |

Model Coefficients - MRPL41

| Predictor | Estimate | SE | t | p |
|---|---|---|---|---|
| Intercept [a] | 18.93622 | 2.22104 | 8.5258 | < .001 |
| Arm: | | | | |
| alcohol use disorder – control | 0.01896 | 0.23371 | 0.0811 | 0.936 |
| opioid use disorder – control | 0.27843 | 0.21862 | 1.2736 | 0.215 |
| opioid+alcohol use disorder – control | -0.11544 | 0.25854 | -0.4465 | 0.659 |
| Age Cateogry: | | | | |
| 50-64 – 30-49 | -0.46487 | 0.23294 | -1.9957 | 0.057 |
| 50-65 – 30-49 | -0.07395 | 0.35785 | -0.2067 | 0.838 |
| 65+ – 30-49 | -0.20192 | 0.26987 | -0.7482 | 0.462 |
| <30 – 30-49 | -0.45944 | 0.28215 | -1.6284 | 0.117 |
| Ethnicity: | | | | |
| Asian – White | -0.27312 | 0.46981 | -0.5813 | 0.566 |
| Black – White | 0.13299 | 0.19127 | 0.6953 | 0.494 |
| Hispanic – White | -0.19955 | 0.36580 | -0.5455 | 0.590 |
| Gender: | | | | |
| Male – Female | 0.23652 | 0.22608 | 1.0462 | 0.306 |
| PMI converted to hours | 0.00987 | 0.00938 | 1.0525 | 0.303 |
| pH | 0.65646 | 0.32430 | 2.0242 | 0.054 |

[a] Represents reference level

### Linear Regression

Model Fit Measures

| Model | R | $R^2$ |
|---|---|---|
| 1 | 0.760 | 0.577 |

Model Coefficients - EXOC8

| Predictor | Estimate | SE | t | p |
|---|---|---|---|---|
| Intercept [a] | 24.14399 | 3.1020 | 7.783 | < .001 |
| Arm: | | | | |
| alcohol use disorder – control | -0.08591 | 0.3264 | -0.263 | 0.795 |
| opioid use disorder – control | -1.01789 | 0.3053 | -3.334 | 0.003 |
| opioid+alcohol use disorder – control | -0.58734 | 0.3611 | -1.627 | 0.117 |
| Age Cateogry: | | | | |
| 50-64 – 30-49 | 0.15165 | 0.3253 | 0.466 | 0.645 |
| 50-65 – 30-49 | -0.62776 | 0.4998 | -1.256 | 0.221 |
| 65+ – 30-49 | 0.43913 | 0.3769 | 1.165 | 0.255 |
| <30 – 30-49 | 0.07993 | 0.3941 | 0.203 | 0.841 |
| Ethnicity: | | | | |
| Asian – White | 0.73576 | 0.6562 | 1.121 | 0.273 |
| Black – White | 0.45730 | 0.2671 | 1.712 | 0.100 |
| Hispanic – White | 0.45161 | 0.5109 | 0.884 | 0.385 |
| Gender: | | | | |
| Male – Female | -0.65483 | 0.3158 | -2.074 | 0.049 |
| PMI converted to hours | -0.00521 | 0.0131 | -0.397 | 0.695 |
| pH | 0.16505 | 0.4529 | 0.364 | 0.719 |

[a] Represents reference level

### Linear Regression

Model Fit Measures

| Model | R | $R^2$ |
|---|---|---|
| 1 | 0.617 | 0.380 |

Model Coefficients - NLGN2

| Predictor | Estimate | SE | t | p |
|---|---|---|---|---|
| Intercept [a] | 24.0158 | 2.4898 | 9.6458 | < .001 |
| Arm: | | | | |
| alcohol use disorder – control | 0.4585 | 0.2620 | 1.7500 | 0.093 |
| opioid use disorder – control | 0.6885 | 0.2451 | 2.8094 | 0.010 |
| opioid+alcohol use disorder – control | 0.1978 | 0.2898 | 0.6824 | 0.502 |
| Age Cateogry: | | | | |
| 50-64 – 30-49 | 0.1789 | 0.2611 | 0.6853 | 0.500 |
| 50-65 – 30-49 | 0.1709 | 0.4012 | 0.4259 | 0.674 |
| 65+ – 30-49 | -0.0942 | 0.3025 | -0.3113 | 0.758 |
| <30 – 30-49 | -0.0956 | 0.3163 | -0.3023 | 0.765 |
| Ethnicity: | | | | |
| Asian – White | 0.0192 | 0.5267 | 0.0365 | 0.971 |
| Black – White | 0.1857 | 0.2144 | 0.8660 | 0.395 |
| Hispanic – White | -0.1870 | 0.4101 | -0.4560 | 0.652 |
| Gender: | | | | |
| Male – Female | 0.1811 | 0.2534 | 0.7145 | 0.482 |
| PMI converted to hours | 0.0126 | 0.0105 | 1.1984 | 0.242 |
| pH | 0.0298 | 0.3635 | 0.0820 | 0.935 |

[a] Represents reference level

### Linear Regression

Model Fit Measures

| Model | R | $R^2$ |
|---|---|---|
| 1 | 0.756 | 0.571 |

Model Coefficients - CCDC132

| Predictor | Estimate | SE | t | p |
|---|---|---|---|---|
| Intercept [a] | 19.79332 | 3.4545 | 5.7298 | < .001 |
| Arm: | | | | |
| alcohol use disorder – control | 0.19084 | 0.3635 | 0.5250 | 0.604 |
| opioid use disorder – control | -0.76311 | 0.3400 | -2.2442 | 0.034 |
| opioid+alcohol use disorder – control | -0.01202 | 0.4021 | -0.0299 | 0.976 |
| Age Cateogry: | | | | |
| 50-64 – 30-49 | 0.50465 | 0.3623 | 1.3929 | 0.176 |
| 50-65 – 30-49 | 0.62989 | 0.5566 | 1.1317 | 0.269 |
| 65+ – 30-49 | 0.22784 | 0.4197 | 0.5428 | 0.592 |
| <30 – 30-49 | 1.21638 | 0.4388 | 2.7718 | 0.011 |
| Ethnicity: | | | | |
| Asian – White | 0.83336 | 0.7307 | 1.1405 | 0.265 |
| Black – White | -0.27309 | 0.2975 | -0.9180 | 0.368 |
| Hispanic – White | 0.57581 | 0.5689 | 1.0121 | 0.322 |
| Gender: | | | | |
| Male – Female | -0.37693 | 0.3516 | -1.0720 | 0.294 |
| PMI converted to hours | 0.00327 | 0.0146 | 0.2244 | 0.824 |
| pH | 0.69340 | 0.5044 | 1.3747 | 0.182 |

[a] Represents reference level

### Linear Regression

Model Fit Measures

| Model | R | $R^2$ |
|---|---|---|
| 1 | 0.779 | 0.607 |

Model Coefficients - NEGR1

| Predictor | Estimate | SE | t | p |
|---|---|---|---|---|
| Intercept [a] | 28.13633 | 1.20318 | 23.3850 | < .001 |
| Arm: | | | | |
| alcohol use disorder – control | 0.16307 | 0.12660 | 1.2881 | 0.210 |
| opioid use disorder – control | 0.31651 | 0.11843 | 2.6725 | 0.013 |
| opioid+alcohol use disorder – control | -0.10293 | 0.14006 | -0.7349 | 0.470 |
| Age Cateogry: | | | | |
| 50-64 – 30-49 | -0.18461 | 0.12619 | -1.4630 | 0.156 |
| 50-65 – 30-49 | 0.00758 | 0.19386 | 0.0391 | 0.969 |
| 65+ – 30-49 | -0.17656 | 0.14619 | -1.2077 | 0.239 |
| <30 – 30-49 | -0.14075 | 0.15285 | -0.9208 | 0.366 |
| Ethnicity: | | | | |
| Asian – White | 0.15340 | 0.25451 | 0.6027 | 0.552 |
| Black – White | 0.03068 | 0.10361 | 0.2961 | 0.770 |
| Hispanic – White | -0.27238 | 0.19816 | -1.3745 | 0.182 |
| Gender: | | | | |
| Male – Female | 0.17076 | 0.12247 | 1.3943 | 0.176 |
| PMI converted to hours | 0.01128 | 0.00508 | 2.2201 | 0.036 |
| pH | 0.16910 | 0.17568 | 0.9626 | 0.345 |

[a] Represents reference level

#### Linear Regression

Model Fit Measures

| Model | R | $R^2$ |
|---|---|---|
| 1 | 0.744 | 0.553 |

Model Coefficients - STX12

| Predictor | Estimate | SE | t | p |
|---|---|---|---|---|
| Intercept [a] | 25.64409 | 1.81337 | 14.1417 | < .001 |
| Arm: | | | | |
| alcohol use disorder – control | 0.00654 | 0.19081 | 0.0343 | 0.973 |
| opioid use disorder – control | 0.43210 | 0.17850 | 2.4208 | 0.023 |
| opioid+alcohol use disorder – control | -0.11821 | 0.21108 | -0.5600 | 0.581 |
| Age Cateogry: | | | | |
| 50-64 – 30-49 | -0.22037 | 0.19018 | -1.1587 | 0.258 |
| 50-65 – 30-49 | -0.32716 | 0.29217 | -1.1197 | 0.274 |
| 65+ – 30-49 | 0.08985 | 0.22033 | 0.4078 | 0.687 |
| <30 – 30-49 | -0.50747 | 0.23036 | -2.2029 | 0.037 |
| Ethnicity: | | | | |
| Asian – White | -0.09509 | 0.38358 | -0.2479 | 0.806 |
| Black – White | 0.16085 | 0.15616 | 1.0300 | 0.313 |
| Hispanic – White | 0.07358 | 0.29866 | 0.2464 | 0.808 |
| Gender: | | | | |
| Male – Female | 0.02063 | 0.18458 | 0.1118 | 0.912 |
| PMI converted to hours | 0.01669 | 0.00766 | 2.1788 | 0.039 |
| pH | 0.26613 | 0.26477 | 1.0051 | 0.325 |

[a] Represents reference level

#### Linear Regression

Model Fit Measures

| Model | R | $R^2$ |
|---|---|---|
| 1 | 0.773 | 0.597 |

Model Coefficients - D2HGDH (2)

| Predictor | Estimate | SE | t | p |
|---|---|---|---|---|
| Intercept [a] | 23.6156 | 4.6864 | 5.0392 | < .001 |
| Arm: | | | | |
| alcohol use disorder – control | 0.8931 | 0.4931 | 1.8112 | 0.083 |
| opioid use disorder – control | 0.7569 | 0.4613 | 1.6408 | 0.114 |
| opioid+alcohol use disorder – control | -1.2802 | 0.5455 | -2.3467 | 0.028 |
| Age Cateogry: | | | | |
| 50-64 – 30-49 | -0.4817 | 0.4915 | -0.9801 | 0.337 |
| 50-65 – 30-49 | -0.6320 | 0.7551 | -0.8370 | 0.411 |
| 65+ – 30-49 | 0.0114 | 0.5694 | 0.0201 | 0.984 |
| <30 – 30-49 | -0.1817 | 0.5953 | -0.3053 | 0.763 |
| Ethnicity: | | | | |
| Asian – White | -0.6156 | 0.9913 | -0.6210 | 0.540 |
| Black – White | 0.0674 | 0.4036 | 0.1671 | 0.869 |
| Hispanic – White | -0.6891 | 0.7718 | -0.8929 | 0.381 |
| Gender: | | | | |
| Male – Female | -0.1569 | 0.4770 | -0.3289 | 0.745 |
| PMI converted to hours | 0.0150 | 0.0198 | 0.7565 | 0.457 |
| pH | 0.1024 | 0.6843 | 0.1497 | 0.882 |

[a] Represents reference level

#### Linear Regression

Model Fit Measures

| Model | R | $R^2$ |
|---|---|---|
| 1 | 0.810 | 0.656 |

Model Coefficients - LRRC57

| Predictor | Estimate | SE | t | p |
|---|---|---|---|---|
| Intercept [a] | 27.63666 | 1.47663 | 18.716 | < .001 |
| Arm: | | | | |
| alcohol use disorder – control | 0.17627 | 0.15538 | 1.134 | 0.268 |
| opioid use disorder – control | 0.42110 | 0.14535 | 2.897 | 0.008 |
| opioid+alcohol use disorder – control | -0.29420 | 0.17189 | -1.712 | 0.100 |
| Age Cateogry: | | | | |
| 50-64 – 30-49 | 0.10428 | 0.15487 | 0.673 | 0.507 |
| 50-65 – 30-49 | -0.07195 | 0.23792 | -0.302 | 0.765 |
| 65+ – 30-49 | 0.14056 | 0.17942 | 0.783 | 0.441 |
| <30 – 30-49 | 0.07274 | 0.18758 | 0.388 | 0.702 |
| Ethnicity: | | | | |
| Asian – White | -0.79414 | 0.31235 | -2.542 | 0.018 |
| Black – White | 0.08672 | 0.12716 | 0.682 | 0.502 |
| Hispanic – White | -0.24174 | 0.24320 | -0.994 | 0.330 |
| Gender: | | | | |
| Male – Female | -0.17925 | 0.15031 | -1.193 | 0.245 |
| PMI converted to hours | 0.00486 | 0.00624 | 0.778 | 0.444 |
| pH | -0.08387 | 0.21561 | -0.389 | 0.701 |

[a] Represents reference level

#### Linear Regression

Model Fit Measures

| Model | R | $R^2$ |
|---|---|---|
| 1 | 0.657 | 0.432 |

Model Coefficients - SYNPR

| Predictor | Estimate | SE | t | p |
|---|---|---|---|---|
| Intercept [a] | 24.3831 | 4.4106 | 5.528 | < .001 |
| Arm: | | | | |
| alcohol use disorder – control | 0.7745 | 0.4641 | 1.669 | 0.108 |
| opioid use disorder – control | 0.9432 | 0.4342 | 2.172 | 0.040 |
| opioid+alcohol use disorder – control | -0.5189 | 0.5134 | -1.011 | 0.322 |
| Age Cateogry: | | | | |
| 50-64 – 30-49 | 0.3616 | 0.4626 | 0.782 | 0.442 |
| 50-65 – 30-49 | 0.7357 | 0.7106 | 1.035 | 0.311 |
| 65+ – 30-49 | 0.1666 | 0.5359 | 0.311 | 0.759 |
| <30 – 30-49 | 0.4047 | 0.5603 | 0.722 | 0.477 |
| Ethnicity: | | | | |
| Asian – White | 0.1443 | 0.9330 | 0.155 | 0.878 |
| Black – White | 0.0850 | 0.3798 | 0.224 | 0.825 |
| Hispanic – White | 0.5060 | 0.7264 | 0.697 | 0.493 |
| Gender: | | | | |
| Male – Female | -0.2162 | 0.4490 | -0.481 | 0.635 |
| PMI converted to hours | 0.0108 | 0.0186 | 0.579 | 0.568 |
| pH | 0.4176 | 0.6440 | 0.649 | 0.523 |

[a] Represents reference level

### Linear Regression

Model Fit Measures

| Model | R | $R^2$ |
|---|---|---|
| 1 | 0.758 | 0.575 |

Model Coefficients - TNR

| Predictor | Estimate | SE | t | p |
|---|---|---|---|---|
| Intercept [a] | 33.50008 | 0.63997 | 52.3466 | < .001 |
| Arm: | | | | |
| alcohol use disorder – control | -0.00237 | 0.06734 | -0.0352 | 0.972 |
| opioid use disorder – control | 0.13610 | 0.06299 | 2.1605 | 0.041 |
| opioid+alcohol use disorder – control | -0.17479 | 0.07449 | -2.3463 | 0.028 |
| Age Cateogry: | | | | |
| 50-64 – 30-49 | -0.05075 | 0.06712 | -0.7561 | 0.457 |
| 50-65 – 30-49 | -0.05370 | 0.10311 | -0.5207 | 0.607 |
| 65+ – 30-49 | 0.04972 | 0.07776 | 0.6395 | 0.529 |
| <30 – 30-49 | -0.00367 | 0.08130 | -0.0452 | 0.964 |
| Ethnicity: | | | | |
| Asian – White | -0.13212 | 0.13537 | -0.9760 | 0.339 |
| Black – White | -0.06612 | 0.05511 | -1.1997 | 0.242 |
| Hispanic – White | -0.15956 | 0.10540 | -1.5139 | 0.143 |
| Gender: | | | | |
| Male – Female | -0.01877 | 0.06514 | -0.2882 | 0.776 |
| PMI converted to hours | 0.00228 | 0.00270 | 0.8453 | 0.406 |
| pH | -0.12873 | 0.09344 | -1.3777 | 0.181 |

[a] Represents reference level

### Linear Regression

Model Fit Measures

| Model | R | $R^2$ |
|---|---|---|
| 1 | 0.723 | 0.522 |

Model Coefficients - PDXP

| Predictor | Estimate | SE | t | p |
|---|---|---|---|---|
| Intercept [a] | 29.3654 | 0.90197 | 32.557 | < .001 |
| Arm: | | | | |
| alcohol use disorder – control | 0.1323 | 0.09491 | 1.394 | 0.176 |
| opioid use disorder – control | 0.1901 | 0.08878 | 2.141 | 0.043 |
| opioid+alcohol use disorder – control | -0.1269 | 0.10499 | -1.208 | 0.239 |
| Age Categry: | | | | |
| 50-64 – 30-49 | -0.0755 | 0.09460 | -0.799 | 0.432 |
| 50-65 – 30-49 | -0.0536 | 0.14533 | -0.369 | 0.715 |
| 65+ – 30-49 | -0.0629 | 0.10959 | -0.574 | 0.571 |
| <30 – 30-49 | -0.0329 | 0.11458 | -0.287 | 0.777 |
| Ethnicity: | | | | |
| Asian – White | -0.1651 | 0.19079 | -0.865 | 0.395 |
| Black – White | 0.0332 | 0.07767 | 0.427 | 0.673 |
| Hispanic – White | -0.1024 | 0.14855 | -0.689 | 0.497 |
| Gender: | | | | |
| Male – Female | -0.0412 | 0.09181 | -0.449 | 0.657 |
| PMI converted to hours | 8.62e-4 | 0.00381 | 0.226 | 0.823 |
| pH | 0.0205 | 0.13170 | 0.156 | 0.878 |

[a] Represents reference level

### Linear Regression

Model Fit Measures

| Model | R | $R^2$ |
|---|---|---|
| 1 | 0.820 | 0.672 |

Model Coefficients - CPPED1

| Predictor | Estimate | SE | t | p |
|---|---|---|---|---|
| Intercept [a] | 31.4350 | 4.0746 | 7.715 | < .001 |
| Arm: | | | | |
| alcohol use disorder – control | 0.4316 | 0.4287 | 1.007 | 0.324 |
| opioid use disorder – control | -1.3675 | 0.4011 | -3.409 | 0.002 |
| opioid+alcohol use disorder – control | 0.4725 | 0.4743 | 0.996 | 0.329 |
| Age Categry: | | | | |
| 50-64 – 30-49 | 0.3494 | 0.4273 | 0.818 | 0.422 |
| 50-65 – 30-49 | 0.2081 | 0.6565 | 0.317 | 0.754 |
| 65+ – 30-49 | 0.3730 | 0.4951 | 0.753 | 0.459 |
| <30 – 30-49 | 0.2973 | 0.5176 | 0.574 | 0.571 |
| Ethnicity: | | | | |
| Asian – White | 0.1135 | 0.8619 | 0.132 | 0.896 |
| Black – White | 0.5207 | 0.3509 | 1.484 | 0.151 |
| Hispanic – White | -0.8160 | 0.6711 | -1.216 | 0.236 |
| Gender: | | | | |
| Male – Female | 0.2198 | 0.4148 | 0.530 | 0.601 |
| PMI converted to hours | -0.0112 | 0.0172 | -0.650 | 0.522 |
| pH | -0.9224 | 0.5949 | -1.550 | 0.134 |

[a] Represents reference level

### Linear Regression

Model Fit Measures

| Model | R | $R^2$ |
|---|---|---|
| 1 | 0.697 | 0.486 |

Model Coefficients - SH3GL3

| Predictor | Estimate | SE | t | p |
|---|---|---|---|---|
| Intercept [a] | 29.41897 | 1.26139 | 23.3227 | < .001 |
| Arm: | | | | |
| alcohol use disorder – control | 0.02936 | 0.13273 | 0.2212 | 0.827 |
| opioid use disorder – control | 0.11894 | 0.12416 | 0.9580 | 0.348 |
| opioid+alcohol use disorder – control | -0.28980 | 0.14683 | -1.9737 | 0.060 |
| Age Cateogry: | | | | |
| 50-64 – 30-49 | -0.07867 | 0.13229 | -0.5947 | 0.558 |
| 50-65 – 30-49 | 0.02863 | 0.20324 | 0.1409 | 0.889 |
| 65+ – 30-49 | -0.17591 | 0.15326 | -1.1477 | 0.262 |
| <30 – 30-49 | 0.17392 | 0.16024 | 1.0854 | 0.289 |
| Ethnicity: | | | | |
| Asian – White | 0.01841 | 0.26682 | 0.0690 | 0.946 |
| Black – White | -0.08529 | 0.10862 | -0.7851 | 0.440 |
| Hispanic – White | -0.28149 | 0.20775 | -1.3549 | 0.188 |
| Gender: | | | | |
| Male – Female | -0.20697 | 0.12840 | -1.6120 | 0.120 |
| PMI converted to hours | 0.00891 | 0.00533 | 1.6731 | 0.107 |
| pH | -0.08456 | 0.18418 | -0.4591 | 0.650 |

[a] Represents reference level

### Linear Regression

Model Fit Measures

| Model | R | $R^2$ |
|---|---|---|
| 1 | 0.636 | 0.405 |

Model Coefficients - CNNM2

| Predictor | Estimate | SE | t | p |
|---|---|---|---|---|
| Intercept [a] | 22.0537 | 2.4758 | 8.9076 | < .001 |
| Arm: | | | | |
| alcohol use disorder – control | -0.1203 | 0.2605 | -0.4618 | 0.648 |
| opioid use disorder – control | 0.3381 | 0.2437 | 1.3872 | 0.178 |
| opioid+alcohol use disorder – control | -0.4842 | 0.2882 | -1.6802 | 0.106 |
| Age Cateogry: | | | | |
| 50-64 – 30-49 | -0.0585 | 0.2597 | -0.2251 | 0.824 |
| 50-65 – 30-49 | 0.2392 | 0.3989 | 0.5996 | 0.554 |
| 65+ – 30-49 | 0.0248 | 0.3008 | 0.0824 | 0.935 |
| <30 – 30-49 | -0.1034 | 0.3145 | -0.3288 | 0.745 |
| Ethnicity: | | | | |
| Asian – White | -0.5815 | 0.5237 | -1.1104 | 0.278 |
| Black – White | 0.0211 | 0.2132 | 0.0988 | 0.922 |
| Hispanic – White | 0.0421 | 0.4078 | 0.1033 | 0.919 |
| Gender: | | | | |
| Male – Female | 0.0608 | 0.2520 | 0.2411 | 0.812 |
| PMI converted to hours | 8.62e-4 | 0.0105 | 0.0825 | 0.935 |
| pH | 0.2102 | 0.3615 | 0.5814 | 0.566 |

[a] Represents reference level

### Linear Regression

Model Fit Measures

| Model | R | $R^2$ |
|---|---|---|
| 1 | 0.726 | 0.527 |

Model Coefficients - ACTR3B

| Predictor | Estimate | SE | t | p |
|---|---|---|---|---|
| Intercept [a] | 28.78831 | 1.42848 | 20.153 | < .001 |
| Arm: | | | | |
| alcohol use disorder – control | 0.01763 | 0.15031 | 0.117 | 0.908 |
| opioid use disorder – control | 0.37262 | 0.14061 | 2.650 | 0.014 |
| opioid+alcohol use disorder – control | 0.19382 | 0.16628 | 1.166 | 0.255 |
| Age Cateogry: | | | | |
| 50-64 – 30-49 | 0.06861 | 0.14982 | 0.458 | 0.651 |
| 50-65 – 30-49 | 0.03000 | 0.23016 | 0.130 | 0.897 |
| 65+ – 30-49 | -0.15365 | 0.17357 | -0.885 | 0.385 |
| <30 – 30-49 | 0.12267 | 0.18147 | 0.676 | 0.506 |
| Ethnicity: | | | | |
| Asian – White | 0.32984 | 0.30216 | 1.092 | 0.286 |
| Black – White | 0.20936 | 0.12301 | 1.702 | 0.102 |
| Hispanic – White | -0.08880 | 0.23527 | -0.377 | 0.709 |
| Gender: | | | | |
| Male – Female | -0.02577 | 0.14541 | -0.177 | 0.861 |
| PMI converted to hours | 0.00557 | 0.00603 | 0.923 | 0.365 |
| pH | -0.08893 | 0.20857 | -0.426 | 0.674 |

[a] Represents reference level

### Linear Regression

Model Fit Measures

| Model | R | $R^2$ |
|---|---|---|
| 1 | 0.682 | 0.465 |

Model Coefficients - ACAT2

| Predictor | Estimate | SE | t | p |
|---|---|---|---|---|
| Intercept [a] | 27.9487 | 1.41643 | 19.732 | < .001 |
| Arm: | | | | |
| alcohol use disorder – control | 0.0224 | 0.14904 | 0.150 | 0.882 |
| opioid use disorder – control | 0.3292 | 0.13942 | 2.361 | 0.027 |
| opioid+alcohol use disorder – control | -0.1625 | 0.16488 | -0.986 | 0.334 |
| Age Cateogry: | | | | |
| 50-64 – 30-49 | -0.0676 | 0.14855 | -0.455 | 0.653 |
| 50-65 – 30-49 | 0.1877 | 0.22822 | 0.822 | 0.419 |
| 65+ – 30-49 | 0.0345 | 0.17210 | 0.201 | 0.843 |
| <30 – 30-49 | -0.1383 | 0.17994 | -0.769 | 0.450 |
| Ethnicity: | | | | |
| Asian – White | -0.1956 | 0.29961 | -0.653 | 0.520 |
| Black – White | -0.0740 | 0.12198 | -0.606 | 0.550 |
| Hispanic – White | -0.0426 | 0.23328 | -0.183 | 0.857 |
| Gender: | | | | |
| Male – Female | 0.0272 | 0.14418 | 0.188 | 0.852 |
| PMI converted to hours | 0.0125 | 0.00598 | 2.096 | 0.047 |
| pH | 0.1629 | 0.20682 | 0.788 | 0.439 |

[a] Represents reference level

### Linear Regression

Model Fit Measures

| Model | R | $R^2$ |
|---|---|---|
| 1 | 0.778 | 0.605 |

Model Coefficients - SLC25A22

| Predictor | Estimate | SE | t | p |
|---|---|---|---|---|
| Intercept [a] | 30.78671 | 0.95950 | 32.0861 | < .001 |
| Arm: | | | | |
| alcohol use disorder – control | 0.02140 | 0.10096 | 0.2120 | 0.834 |
| opioid use disorder – control | 0.23790 | 0.09445 | 2.5189 | 0.019 |
| opioid+alcohol use disorder – control | 0.06503 | 0.11169 | 0.5823 | 0.566 |
| Age Cateogry: | | | | |
| 50-64 – 30-49 | -0.20919 | 0.10063 | -2.0788 | 0.048 |
| 50-65 – 30-49 | -0.15416 | 0.15460 | -0.9972 | 0.329 |
| 65+ – 30-49 | -0.36860 | 0.11658 | -3.1617 | 0.004 |
| <30 – 30-49 | -0.27520 | 0.12189 | -2.2577 | 0.033 |
| Ethnicity: | | | | |
| Asian – White | 0.00644 | 0.20296 | 0.0317 | 0.975 |
| Black – White | -0.02416 | 0.08263 | -0.2924 | 0.772 |
| Hispanic – White | -0.08784 | 0.15803 | -0.5558 | 0.583 |
| Gender: | | | | |
| Male – Female | 0.14555 | 0.09767 | 1.4903 | 0.149 |
| PMI converted to hours | 0.00752 | 0.00405 | 1.8557 | 0.076 |
| pH | -0.01715 | 0.14010 | -0.1224 | 0.904 |

[a] Represents reference level

#### Linear Regression

Model Fit Measures

| Model | R | $R^2$ |
|---|---|---|
| 1 | 0.615 | 0.378 |

Model Coefficients - GNB4

| Predictor | Estimate | SE | t | p |
|---|---|---|---|---|
| Intercept [a] | 28.29590 | 1.90811 | 14.8293 | < .001 |
| Arm: | | | | |
| alcohol use disorder – control | -0.01692 | 0.20078 | -0.0843 | 0.934 |
| opioid use disorder – control | 0.34038 | 0.18782 | 1.8122 | 0.082 |
| opioid+alcohol use disorder – control | -0.20754 | 0.22211 | -0.9344 | 0.359 |
| Age Cateogry: | | | | |
| 50-64 – 30-49 | -0.09235 | 0.20012 | -0.4615 | 0.649 |
| 50-65 – 30-49 | -0.18167 | 0.30744 | -0.5909 | 0.560 |
| 65+ – 30-49 | 0.05565 | 0.23184 | 0.2400 | 0.812 |
| <30 – 30-49 | -0.06808 | 0.24240 | -0.2808 | 0.781 |
| Ethnicity: | | | | |
| Asian – White | 0.09100 | 0.40362 | 0.2255 | 0.824 |
| Black – White | 0.02493 | 0.16432 | 0.1517 | 0.881 |
| Hispanic – White | -0.28732 | 0.31426 | -0.9143 | 0.370 |
| Gender: | | | | |
| Male – Female | 0.00648 | 0.19423 | 0.0334 | 0.974 |
| PMI converted to hours | 0.01286 | 0.00806 | 1.5960 | 0.124 |
| pH | -0.06922 | 0.27861 | -0.2485 | 0.806 |

[a] Represents reference level

#### Linear Regression

Model Fit Measures

| Model | R | $R^2$ |
|---|---|---|
| 1 | 0.597 | 0.356 |

Model Coefficients - GDAP1L1

| Predictor | Estimate | SE | t | p |
|---|---|---|---|---|
| Intercept [a] | 22.3135 | 4.1746 | 5.345 | < .001 |
| Arm: | | | | |
| alcohol use disorder – control | -0.6842 | 0.4393 | -1.558 | 0.132 |
| opioid use disorder – control | -0.3914 | 0.4109 | -0.953 | 0.350 |
| opioid+alcohol use disorder – control | 0.3816 | 0.4859 | 0.785 | 0.440 |
| Age Cateogry: | | | | |
| 50-64 – 30-49 | 0.1791 | 0.4378 | 0.409 | 0.686 |
| 50-65 – 30-49 | 0.5010 | 0.6726 | 0.745 | 0.464 |
| 65+ – 30-49 | 0.2104 | 0.5072 | 0.415 | 0.682 |
| <30 – 30-49 | 0.3811 | 0.5303 | 0.719 | 0.479 |
| Ethnicity: | | | | |
| Asian – White | 0.3552 | 0.8830 | 0.402 | 0.691 |
| Black – White | -0.3004 | 0.3595 | -0.836 | 0.412 |
| Hispanic – White | 0.6848 | 0.6875 | 0.996 | 0.329 |
| Gender: | | | | |
| Male – Female | 0.1089 | 0.4249 | 0.256 | 0.800 |
| PMI converted to hours | -0.0165 | 0.0176 | -0.938 | 0.358 |
| pH | 0.7516 | 0.6095 | 1.233 | 0.230 |

[a] Represents reference level

#### Linear Regression

Model Fit Measures

| Model | R | $R^2$ |
|---|---|---|
| 1 | 0.572 | 0.327 |

Model Coefficients - C6orf203

| Predictor | Estimate | SE | t | p |
|---|---|---|---|---|
| Intercept [a] | 23.2293 | 3.0873 | 7.5241 | < .001 |
| Arm: | | | | |
| alcohol use disorder – control | -0.1287 | 0.3249 | -0.3962 | 0.695 |
| opioid use disorder – control | 0.4517 | 0.3039 | 1.4863 | 0.150 |
| opioid+alcohol use disorder – control | -0.3990 | 0.3594 | -1.1103 | 0.278 |
| Age Cateogry: | | | | |
| 50-64 – 30-49 | -0.0194 | 0.3238 | -0.0600 | 0.953 |
| 50-65 – 30-49 | 0.2927 | 0.4974 | 0.5885 | 0.562 |
| 65+ – 30-49 | 0.3406 | 0.3751 | 0.9080 | 0.373 |
| <30 – 30-49 | 0.0466 | 0.3922 | 0.1188 | 0.906 |
| Ethnicity: | | | | |
| Asian – White | -0.0817 | 0.6531 | -0.1250 | 0.902 |
| Black – White | -0.0711 | 0.2659 | -0.2673 | 0.792 |
| Hispanic – White | -0.1996 | 0.5085 | -0.3925 | 0.698 |
| Gender: | | | | |
| Male – Female | -0.1542 | 0.3143 | -0.4907 | 0.628 |
| PMI converted to hours | 8.46e-4 | 0.0130 | 0.0649 | 0.949 |
| pH | 0.0335 | 0.4508 | 0.0743 | 0.941 |

[a] Represents reference level

#### Linear Regression

Model Fit Measures

| Model | R | $R^2$ |
|---|---|---|
| 1 | 0.727 | 0.529 |

Model Coefficients - UQCC2

| Predictor | Estimate | SE | t | p |
|---|---|---|---|---|
| Intercept [a] | 24.9161 | 2.7977 | 8.906 | < .001 |
| Arm: | | | | |
| alcohol use disorder – control | -0.2398 | 0.2944 | -0.815 | 0.423 |
| opioid use disorder – control | 0.5882 | 0.2754 | 2.136 | 0.043 |
| opioid+alcohol use disorder – control | -0.4671 | 0.3257 | -1.434 | 0.164 |
| Age Cateogry: | | | | |
| 50-64 – 30-49 | -0.2197 | 0.2934 | -0.749 | 0.461 |
| 50-65 – 30-49 | -0.1194 | 0.4508 | -0.265 | 0.793 |
| 65+ – 30-49 | -0.0771 | 0.3399 | -0.227 | 0.822 |
| <30 – 30-49 | -0.6628 | 0.3554 | -1.865 | 0.074 |
| Ethnicity: | | | | |
| Asian – White | -0.5795 | 0.5918 | -0.979 | 0.337 |
| Black – White | -0.2778 | 0.2409 | -1.153 | 0.260 |
| Hispanic – White | -0.8784 | 0.4608 | -1.906 | 0.069 |
| Gender: | | | | |
| Male – Female | 0.3998 | 0.2848 | 1.404 | 0.173 |
| PMI converted to hours | 0.0191 | 0.0118 | 1.617 | 0.119 |
| pH | -0.3057 | 0.4085 | -0.748 | 0.462 |

[a] Represents reference level

### Linear Regression

Model Fit Measures

| Model | R | $R^2$ |
|---|---|---|
| 1 | 0.557 | 0.310 |

Model Coefficients - UGGT1

| Predictor | Estimate | SE | t | p |
|---|---|---|---|---|
| Intercept [a] | 28.6754 | 3.5995 | 7.9665 | < .001 |
| Arm: | | | | |
| alcohol use disorder – control | 0.0343 | 0.3788 | 0.0905 | 0.929 |
| opioid use disorder – control | 0.6070 | 0.3543 | 1.7132 | 0.100 |
| opioid+alcohol use disorder – control | 0.0927 | 0.4190 | 0.2212 | 0.827 |
| Age Cateogry: | | | | |
| 50-64 – 30-49 | -0.3191 | 0.3775 | -0.8452 | 0.406 |
| 50-65 – 30-49 | -0.1704 | 0.5800 | -0.2937 | 0.771 |
| 65+ – 30-49 | -0.0418 | 0.4374 | -0.0956 | 0.925 |
| <30 – 30-49 | -0.9221 | 0.4573 | -2.0166 | 0.055 |
| Ethnicity: | | | | |
| Asian – White | -0.6195 | 0.7614 | -0.8137 | 0.424 |
| Black – White | -0.0538 | 0.3100 | -0.1736 | 0.864 |
| Hispanic – White | -0.3077 | 0.5928 | -0.5190 | 0.609 |
| Gender: | | | | |
| Male – Female | 0.3942 | 0.3664 | 1.0758 | 0.293 |
| PMI converted to hours | 0.0142 | 0.0152 | 0.9319 | 0.361 |
| pH | 0.2913 | 0.5256 | 0.5542 | 0.585 |

[a] Represents reference level

### Linear Regression

Model Fit Measures

| Model | R | $R^2$ |
|---|---|---|
| 1 | 0.586 | 0.343 |

Model Coefficients - FAM49B

| Predictor | Estimate | SE | t | p |
|---|---|---|---|---|
| Intercept [a] | 28.93269 | 1.23292 | 23.46686 | < .001 |
| Arm: | | | | |
| alcohol use disorder – control | 0.02034 | 0.12973 | 0.15681 | 0.877 |
| opioid use disorder – control | 0.29159 | 0.12136 | 2.40266 | 0.024 |
| opioid+alcohol use disorder – control | 0.12347 | 0.14352 | 0.86033 | 0.398 |
| Age Cateogry: | | | | |
| 50-64 – 30-49 | -7.40e−4 | 0.12931 | -0.00572 | 0.995 |
| 50-65 – 30-49 | 0.13598 | 0.19865 | 0.68455 | 0.500 |
| 65+ – 30-49 | -0.10600 | 0.14980 | -0.70761 | 0.486 |
| <30 – 30-49 | -0.10754 | 0.15662 | -0.68663 | 0.499 |
| Ethnicity: | | | | |
| Asian – White | 0.12282 | 0.26080 | 0.47096 | 0.642 |
| Black – White | 0.05153 | 0.10617 | 0.48529 | 0.632 |
| Hispanic – White | -0.18111 | 0.20306 | -0.89189 | 0.381 |
| Gender: | | | | |
| Male – Female | 0.14781 | 0.12550 | 1.17775 | 0.250 |
| PMI converted to hours | 0.00412 | 0.00521 | 0.79078 | 0.437 |
| pH | 0.00328 | 0.18002 | 0.01820 | 0.986 |

[a] Represents reference level

### Linear Regression

Model Fit Measures

| Model | R | $R^2$ |
|---|---|---|
| 1 | 0.749 | 0.561 |

Model Coefficients - NLGN3

| Predictor | Estimate | SE | t | p |
|---|---|---|---|---|
| Intercept [a] | 19.3753 | 2.8291 | 6.848 | < .001 |
| Arm: | | | | |
| alcohol use disorder – control | -0.2748 | 0.2977 | -0.923 | 0.365 |
| opioid use disorder – control | 0.6295 | 0.2785 | 2.260 | 0.033 |
| opioid+alcohol use disorder – control | -0.1670 | 0.3293 | -0.507 | 0.617 |
| Age Cateogry: | | | | |
| 50-64 – 30-49 | -0.6766 | 0.2967 | -2.280 | 0.032 |
| 50-65 – 30-49 | -0.3640 | 0.4558 | -0.799 | 0.432 |
| 65+ – 30-49 | 0.2127 | 0.3438 | 0.619 | 0.542 |
| <30 – 30-49 | -0.6379 | 0.3594 | -1.775 | 0.089 |
| Ethnicity: | | | | |
| Asian – White | -0.5272 | 0.5984 | -0.881 | 0.387 |
| Black – White | -0.1918 | 0.2436 | -0.787 | 0.439 |
| Hispanic – White | -0.0803 | 0.4660 | -0.172 | 0.865 |
| Gender: | | | | |
| Male – Female | 0.0980 | 0.2880 | 0.340 | 0.736 |
| PMI converted to hours | 0.0268 | 0.0119 | 2.241 | 0.035 |
| pH | 0.5910 | 0.4131 | 1.431 | 0.165 |

[a] Represents reference level

### Linear Regression

Model Fit Measures

| Model | R | $R^2$ |
|---|---|---|
| 1 | 0.576 | 0.332 |

Model Coefficients - MYADM

| Predictor | Estimate | SE | t | p |
|---|---|---|---|---|
| Intercept [a] | 29.4662 | 3.7719 | 7.8120 | < .001 |
| Arm: | | | | |
| alcohol use disorder – control | 0.1850 | 0.3969 | 0.4660 | 0.645 |
| opioid use disorder – control | -0.6063 | 0.3713 | -1.6330 | 0.116 |
| opioid+alcohol use disorder – control | -0.8208 | 0.4391 | -1.8694 | 0.074 |
| Age Cateogry: | | | | |
| 50-64 – 30-49 | 0.6342 | 0.3956 | 1.6032 | 0.122 |
| 50-65 – 30-49 | 0.0111 | 0.6077 | 0.0182 | 0.986 |
| 65+ – 30-49 | 0.2034 | 0.4583 | 0.4438 | 0.661 |
| <30 – 30-49 | 0.2759 | 0.4792 | 0.5758 | 0.570 |
| Ethnicity: | | | | |
| Asian – White | 0.2663 | 0.7979 | 0.3338 | 0.741 |
| Black – White | 0.1938 | 0.3248 | 0.5966 | 0.556 |
| Hispanic – White | 0.1412 | 0.6212 | 0.2273 | 0.822 |
| Gender: | | | | |
| Male – Female | -0.5291 | 0.3839 | -1.3781 | 0.181 |
| PMI converted to hours | -0.0177 | 0.0159 | -1.1113 | 0.277 |
| pH | -0.3474 | 0.5507 | -0.6308 | 0.534 |

[a] Represents reference level

#### Linear Regression

Model Fit Measures

| Model | R | $R^2$ |
|---|---|---|
| 1 | 0.620 | 0.384 |

Model Coefficients - SH3GLB2

| Predictor | Estimate | SE | t | p |
|---|---|---|---|---|
| Intercept [a] | 30.57311 | 1.29185 | 23.6661 | < .001 |
| Arm: | | | | |
| alcohol use disorder – control | 0.04878 | 0.13593 | 0.3588 | 0.723 |
| opioid use disorder – control | 0.28322 | 0.12716 | 2.2273 | 0.036 |
| opioid+alcohol use disorder – control | -0.11377 | 0.15038 | -0.7566 | 0.457 |
| Age Cateogry: | | | | |
| 50-64 – 30-49 | -0.01998 | 0.13549 | -0.1474 | 0.884 |
| 50-65 – 30-49 | 0.00254 | 0.20814 | 0.0122 | 0.990 |
| 65+ – 30-49 | 0.04416 | 0.15697 | 0.2813 | 0.781 |
| <30 – 30-49 | -0.11520 | 0.16411 | -0.7020 | 0.489 |
| Ethnicity: | | | | |
| Asian – White | -0.28080 | 0.27326 | -1.0276 | 0.314 |
| Black – White | -0.02801 | 0.11125 | -0.2518 | 0.803 |
| Hispanic – White | -0.16403 | 0.21277 | -0.7709 | 0.448 |
| Gender: | | | | |
| Male – Female | 0.02586 | 0.13150 | 0.1966 | 0.846 |
| PMI converted to hours | 0.00614 | 0.00546 | 1.1256 | 0.271 |
| pH | -0.10054 | 0.18863 | -0.5330 | 0.599 |

[a] Represents reference level

#### Linear Regression

Model Fit Measures

| Model | R | $R^2$ |
|---|---|---|
| 1 | 0.691 | 0.478 |

Model Coefficients - NUCKS1

| Predictor | Estimate | SE | t | p |
|---|---|---|---|---|
| Intercept [a] | 27.1819 | 3.6034 | 7.543 | < .001 |
| Arm: | | | | |
| alcohol use disorder – control | 0.6421 | 0.3792 | 1.693 | 0.103 |
| opioid use disorder – control | 0.6134 | 0.3547 | 1.729 | 0.097 |
| opioid+alcohol use disorder – control | -0.9222 | 0.4195 | -2.198 | 0.038 |
| Age Cateogry: | | | | |
| 50-64 – 30-49 | 0.5464 | 0.3779 | 1.446 | 0.161 |
| 50-65 – 30-49 | 0.5715 | 0.5806 | 0.984 | 0.335 |
| 65+ – 30-49 | 0.4358 | 0.4378 | 0.995 | 0.330 |
| <30 – 30-49 | 0.7772 | 0.4578 | 1.698 | 0.102 |
| Ethnicity: | | | | |
| Asian – White | 0.4479 | 0.7622 | 0.588 | 0.562 |
| Black – White | -0.3709 | 0.3103 | -1.195 | 0.244 |
| Hispanic – White | -0.4536 | 0.5935 | -0.764 | 0.452 |
| Gender: | | | | |
| Male – Female | -0.4110 | 0.3668 | -1.121 | 0.274 |
| PMI converted to hours | 0.0147 | 0.0152 | 0.967 | 0.343 |
| pH | -0.1185 | 0.5261 | -0.225 | 0.824 |

[a] Represents reference level

#### Linear Regression

Model Fit Measures

| Model | R | $R^2$ |
|---|---|---|
| 1 | 0.601 | 0.361 |

Model Coefficients - ABRACL

| Predictor | Estimate | SE | t | p |
|---|---|---|---|---|
| Intercept [a] | 20.58847 | 4.0593 | 5.0719 | < .001 |
| Arm: | | | | |
| alcohol use disorder – control | 0.59573 | 0.4271 | 1.3947 | 0.176 |
| opioid use disorder – control | 0.78635 | 0.3996 | 1.9680 | 0.061 |
| opioid+alcohol use disorder – control | -0.00699 | 0.4725 | -0.0148 | 0.988 |
| Age Cateogry: | | | | |
| 50-64 – 30-49 | 0.09316 | 0.4257 | 0.2188 | 0.829 |
| 50-65 – 30-49 | 0.44348 | 0.6540 | 0.6781 | 0.504 |
| 65+ – 30-49 | -0.43706 | 0.4932 | -0.8861 | 0.384 |
| <30 – 30-49 | -0.10091 | 0.5157 | -0.1957 | 0.847 |
| Ethnicity: | | | | |
| Asian – White | 0.35035 | 0.8587 | 0.4080 | 0.687 |
| Black – White | -0.07376 | 0.3496 | -0.2110 | 0.835 |
| Hispanic – White | 0.09201 | 0.6686 | 0.1376 | 0.892 |
| Gender: | | | | |
| Male – Female | 0.18284 | 0.4132 | 0.4425 | 0.662 |
| PMI converted to hours | 0.01962 | 0.0171 | 1.1442 | 0.264 |
| pH | 0.70876 | 0.5927 | 1.1958 | 0.243 |

[a] Represents reference level

#### Linear Regression

Model Fit Measures

| Model | R | $R^2$ |
|---|---|---|
| 1 | 0.698 | 0.487 |

Model Coefficients - TMX4

| Predictor | Estimate | SE | t | p |
|---|---|---|---|---|
| Intercept [a] | 29.2784 | 6.3966 | 4.577 | < .001 |
| Arm: | | | | |
| alcohol use disorder – control | -0.2744 | 0.6731 | -0.408 | 0.687 |
| opioid use disorder – control | -1.4595 | 0.6296 | -2.318 | 0.029 |
| opioid+alcohol use disorder – control | 0.9232 | 0.7446 | 1.240 | 0.227 |
| Age Cateogry: | | | | |
| 50-64 – 30-49 | 0.2339 | 0.6709 | 0.349 | 0.730 |
| 50-65 – 30-49 | -0.2694 | 1.0306 | -0.261 | 0.796 |
| 65+ – 30-49 | -0.2232 | 0.7772 | -0.287 | 0.776 |
| <30 – 30-49 | 0.8669 | 0.8126 | 1.067 | 0.297 |
| Ethnicity: | | | | |
| Asian – White | 1.0861 | 1.3531 | 0.803 | 0.430 |
| Black – White | -0.1471 | 0.5508 | -0.267 | 0.792 |
| Hispanic – White | 0.8101 | 1.0535 | 0.769 | 0.449 |
| Gender: | | | | |
| Male – Female | -0.3556 | 0.6511 | -0.546 | 0.590 |
| PMI converted to hours | -0.0562 | 0.0270 | -2.081 | 0.048 |
| pH | -0.3734 | 0.9340 | -0.400 | 0.693 |

[a] Represents reference level

#### Linear Regression

Model Fit Measures

| Model | R | $R^2$ |
|---|---|---|
| 1 | 0.583 | 0.340 |

Model Coefficients - RAB6B

| Predictor | Estimate | SE | t | p |
|---|---|---|---|---|
| Intercept [a] | 28.0011 | 1.35637 | 20.6441 | < .001 |
| Arm: | | | | |
| alcohol use disorder – control | 0.1383 | 0.14272 | 0.9688 | 0.342 |
| opioid use disorder – control | 0.2251 | 0.13351 | 1.6858 | 0.105 |
| opioid+alcohol use disorder – control | 0.0435 | 0.15789 | 0.2752 | 0.786 |
| Age Cateogry: | | | | |
| 50-64 – 30-49 | -0.0798 | 0.14225 | -0.5611 | 0.580 |
| 50-65 – 30-49 | 0.0556 | 0.21854 | 0.2542 | 0.801 |
| 65+ – 30-49 | -0.2450 | 0.16480 | -1.4869 | 0.150 |
| <30 – 30-49 | -0.1356 | 0.17231 | -0.7871 | 0.439 |
| Ethnicity: | | | | |
| Asian – White | -0.1849 | 0.28691 | -0.6445 | 0.525 |
| Black – White | 0.0484 | 0.11680 | 0.4140 | 0.683 |
| Hispanic – White | -0.0478 | 0.22339 | -0.2141 | 0.832 |
| Gender: | | | | |
| Male – Female | 0.0659 | 0.13807 | 0.4771 | 0.638 |
| PMI converted to hours | 2.26e-4 | 0.00573 | 0.0394 | 0.969 |
| pH | -0.0691 | 0.19805 | -0.3492 | 0.730 |

[a] Represents reference level

#### Linear Regression

Model Fit Measures

| Model | R | $R^2$ |
|---|---|---|
| 1 | 0.639 | 0.408 |

Model Coefficients - VAT1

| Predictor | Estimate | SE | t | p |
|---|---|---|---|---|
| Intercept [a] | 27.4813 | 1.84629 | 14.8846 | < .001 |
| Arm: | | | | |
| alcohol use disorder – control | -0.3917 | 0.19427 | -2.0165 | 0.055 |
| opioid use disorder – control | -0.3976 | 0.18174 | -2.1876 | 0.039 |
| opioid+alcohol use disorder – control | 0.0490 | 0.21492 | 0.2282 | 0.821 |
| Age Cateogry: | | | | |
| 50-64 – 30-49 | -0.1394 | 0.19364 | -0.7197 | 0.479 |
| 50-65 – 30-49 | -0.0109 | 0.29747 | -0.0367 | 0.971 |
| 65+ – 30-49 | -0.0323 | 0.22433 | -0.1441 | 0.887 |
| <30 – 30-49 | -0.2131 | 0.23454 | -0.9087 | 0.373 |
| Ethnicity: | | | | |
| Asian – White | -0.0840 | 0.39054 | -0.2150 | 0.832 |
| Black – White | 0.0539 | 0.15899 | 0.3390 | 0.738 |
| Hispanic – White | 0.6931 | 0.30408 | 2.2794 | 0.032 |
| Gender: | | | | |
| Male – Female | 0.1056 | 0.18793 | 0.5620 | 0.579 |
| PMI converted to hours | -0.0109 | 0.00780 | -1.3948 | 0.176 |
| pH | 0.2268 | 0.26958 | 0.8414 | 0.408 |

[a] Represents reference level

### Linear Regression

Model Fit Measures

| Model | R | $R^2$ |
|---|---|---|
| 1 | 0.583 | 0.340 |

Model Coefficients - PACSIN1

| Predictor | Estimate | SE | t | p |
|---|---|---|---|---|
| Intercept [a] | 33.16748 | 1.05192 | 31.53030 | < .001 |
| Arm: | | | | |
| alcohol use disorder – control | 0.12701 | 0.11069 | 1.14750 | 0.262 |
| opioid use disorder – control | 0.22807 | 0.10354 | 2.20261 | 0.037 |
| opioid+alcohol use disorder – control | 0.11596 | 0.12245 | 0.94698 | 0.353 |
| Age Cateogry: | | | | |
| 50-64 – 30-49 | 0.07081 | 0.11032 | 0.64181 | 0.527 |
| 50-65 – 30-49 | 0.03547 | 0.16949 | 0.20925 | 0.836 |
| 65+ – 30-49 | -0.16187 | 0.12781 | -1.26646 | 0.218 |
| <30 – 30-49 | 0.09548 | 0.13363 | 0.71450 | 0.482 |
| Ethnicity: | | | | |
| Asian – White | 0.04825 | 0.22251 | 0.21683 | 0.830 |
| Black – White | -0.05973 | 0.09059 | -0.65937 | 0.516 |
| Hispanic – White | -0.13131 | 0.17325 | -0.75792 | 0.456 |
| Gender: | | | | |
| Male – Female | -1.96e−4 | 0.10708 | -0.00183 | 0.999 |
| PMI converted to hours | 0.00283 | 0.00444 | 0.63689 | 0.530 |
| pH | -0.07600 | 0.15359 | -0.49484 | 0.625 |

[a] Represents reference level

### Linear Regression

Model Fit Measures

| Model | R | $R^2$ |
|---|---|---|
| 1 | 0.592 | 0.351 |

Model Coefficients - GLTP

| Predictor | Estimate | SE | t | p |
|---|---|---|---|---|
| Intercept [a] | 24.74271 | 6.2469 | 3.9608 | < .001 |
| Arm: | | | | |
| alcohol use disorder – control | -0.91511 | 0.6573 | -1.3922 | 0.177 |
| opioid use disorder – control | -1.32583 | 0.6149 | -2.1561 | 0.041 |
| opioid+alcohol use disorder – control | 0.66219 | 0.7272 | 0.9106 | 0.372 |
| Age Cateogry: | | | | |
| 50-64 – 30-49 | -0.76065 | 0.6552 | -1.1610 | 0.257 |
| 50-65 – 30-49 | 0.31656 | 1.0065 | 0.3145 | 0.756 |
| 65+ – 30-49 | -0.62286 | 0.7590 | -0.8206 | 0.420 |
| <30 – 30-49 | -0.53348 | 0.7936 | -0.6722 | 0.508 |
| Ethnicity: | | | | |
| Asian – White | 0.08550 | 1.3214 | 0.0647 | 0.949 |
| Black – White | -4.04e−4 | 0.5380 | -7.51e−4 | 0.999 |
| Hispanic – White | 0.59675 | 1.0289 | 0.5800 | 0.567 |
| Gender: | | | | |
| Male – Female | 0.09851 | 0.6359 | 0.1549 | 0.878 |
| PMI converted to hours | -0.00791 | 0.0264 | -0.2998 | 0.767 |
| pH | 0.20752 | 0.9121 | 0.2275 | 0.822 |

[a] Represents reference level

#### Linear Regression

Model Fit Measures

| Model | R | $R^2$ |
|---|---|---|
| 1 | 0.683 | 0.467 |

Model Coefficients - TBCD

| Predictor | Estimate | SE | t | p |
|---|---|---|---|---|
| Intercept [a] | 22.5187 | 4.8481 | 4.6448 | < .001 |
| Arm: | | | | |
| alcohol use disorder – control | -0.1970 | 0.5101 | -0.3862 | 0.703 |
| opioid use disorder – control | -0.7451 | 0.4772 | -1.5613 | 0.132 |
| opioid+alcohol use disorder – control | 0.2523 | 0.5643 | 0.4470 | 0.659 |
| Age Cateogry: | | | | |
| 50-64 – 30-49 | 0.4363 | 0.5085 | 0.8581 | 0.399 |
| 50-65 – 30-49 | -0.1783 | 0.7811 | -0.2283 | 0.821 |
| 65+ – 30-49 | 1.1915 | 0.5891 | 2.0226 | 0.054 |
| <30 – 30-49 | -0.0251 | 0.6159 | -0.0408 | 0.968 |
| Ethnicity: | | | | |
| Asian – White | 1.1758 | 1.0255 | 1.1465 | 0.263 |
| Black – White | 0.4290 | 0.4175 | 1.0276 | 0.314 |
| Hispanic – White | 1.4748 | 0.7985 | 1.8470 | 0.077 |
| Gender: | | | | |
| Male – Female | 0.0405 | 0.4935 | 0.0821 | 0.935 |
| PMI converted to hours | -0.0119 | 0.0205 | -0.5818 | 0.566 |
| pH | 0.3448 | 0.7079 | 0.4870 | 0.631 |

[a] Represents reference level

#### Linear Regression

Model Fit Measures

| Model | R | $R^2$ |
|---|---|---|
| 1 | 0.704 | 0.495 |

Model Coefficients - CADM1

| Predictor | Estimate | SE | t | p |
|---|---|---|---|---|
| Intercept [a] | 28.86446 | 1.09084 | 26.4607 | < .001 |
| Arm: | | | | |
| alcohol use disorder – control | 0.03839 | 0.11478 | 0.3344 | 0.741 |
| opioid use disorder – control | 0.25866 | 0.10738 | 2.4089 | 0.024 |
| opioid+alcohol use disorder – control | -0.08855 | 0.12698 | -0.6974 | 0.492 |
| Age Cateogry: | | | | |
| 50-64 – 30-49 | -0.10328 | 0.11441 | -0.9028 | 0.376 |
| 50-65 – 30-49 | 0.25503 | 0.17576 | 1.4510 | 0.160 |
| 65+ – 30-49 | -0.00812 | 0.13254 | -0.0613 | 0.952 |
| <30 – 30-49 | -0.09881 | 0.13857 | -0.7131 | 0.483 |
| Ethnicity: | | | | |
| Asian – White | -0.03732 | 0.23074 | -0.1617 | 0.873 |
| Black – White | -0.14118 | 0.09394 | -1.5029 | 0.146 |
| Hispanic – White | -0.19665 | 0.17966 | -1.0946 | 0.285 |
| Gender: | | | | |
| Male – Female | 0.02087 | 0.11104 | 0.1880 | 0.852 |
| PMI converted to hours | 0.01036 | 0.00461 | 2.2495 | 0.034 |
| pH | 0.05996 | 0.15928 | 0.3765 | 0.710 |

[a] Represents reference level

#### Linear Regression

Model Fit Measures

| Model | R | $R^2$ |
|---|---|---|
| 1 | 0.826 | 0.683 |

Model Coefficients - TTYH3

| Predictor | Estimate | SE | t | p |
|---|---|---|---|---|
| Intercept [a] | 22.82033 | 2.3917 | 9.5414 | < .001 |
| Arm: | | | | |
| alcohol use disorder – control | -0.33618 | 0.2517 | -1.3358 | 0.194 |
| opioid use disorder – control | 0.35137 | 0.2354 | 1.4925 | 0.149 |
| opioid+alcohol use disorder – control | -0.85774 | 0.2784 | -3.0809 | 0.005 |
| Age Cateogry: | | | | |
| 50-64 – 30-49 | -0.65713 | 0.2508 | -2.6197 | 0.015 |
| 50-65 – 30-49 | -0.40043 | 0.3854 | -1.0391 | 0.309 |
| 65+ – 30-49 | -0.00742 | 0.2906 | -0.0255 | 0.980 |
| <30 – 30-49 | -0.21099 | 0.3038 | -0.6944 | 0.494 |
| Ethnicity: | | | | |
| Asian – White | -0.59229 | 0.5059 | -1.1707 | 0.253 |
| Black – White | -0.08334 | 0.2060 | -0.4047 | 0.689 |
| Hispanic – White | -0.37321 | 0.3939 | -0.9475 | 0.353 |
| Gender: | | | | |
| Male – Female | 0.14690 | 0.2435 | 0.6034 | 0.552 |
| PMI converted to hours | 0.02115 | 0.0101 | 2.0942 | 0.047 |
| pH | 0.18913 | 0.3492 | 0.5416 | 0.593 |

[a] Represents reference level

#### Linear Regression

Model Fit Measures

| Model | R | $R^2$ |
|---|---|---|
| 1 | 0.814 | 0.662 |

Model Coefficients - RAB1B

| Predictor | Estimate | SE | t | p |
|---|---|---|---|---|
| Intercept [a] | 27.3352 | 1.98121 | 13.7972 | < .001 |
| Arm: | | | | |
| alcohol use disorder – control | 0.1471 | 0.20847 | 0.7055 | 0.487 |
| opioid use disorder – control | 0.4573 | 0.19502 | 2.3447 | 0.028 |
| opioid+alcohol use disorder – control | -0.5581 | 0.23062 | -2.4199 | 0.023 |
| Age Cateogry: | | | | |
| 50-64 – 30-49 | -0.1862 | 0.20779 | -0.8962 | 0.379 |
| 50-65 – 30-49 | -0.2278 | 0.31921 | -0.7137 | 0.482 |
| 65+ – 30-49 | 0.0530 | 0.24073 | 0.2201 | 0.828 |
| <30 – 30-49 | -0.3743 | 0.25168 | -1.4870 | 0.150 |
| Ethnicity: | | | | |
| Asian – White | -0.7372 | 0.41908 | -1.7591 | 0.091 |
| Black – White | 0.1120 | 0.17061 | 0.6566 | 0.518 |
| Hispanic – White | -0.2387 | 0.32630 | -0.7316 | 0.471 |
| Gender: | | | | |
| Male – Female | 0.0114 | 0.20167 | 0.0567 | 0.955 |
| PMI converted to hours | 0.0164 | 0.00837 | 1.9655 | 0.061 |
| pH | 0.2168 | 0.28928 | 0.7495 | 0.461 |

[a] Represents reference level

#### Linear Regression

Model Fit Measures

| Model | R | $R^2$ |
|---|---|---|
| 1 | 0.662 | 0.438 |

Model Coefficients - SH3BGRL3

| Predictor | Estimate | SE | t | p |
|---|---|---|---|---|
| Intercept [a] | 21.90855 | 4.5026 | 4.8657 | < .001 |
| Arm: | | | | |
| alcohol use disorder – control | -0.47336 | 0.4738 | -0.9991 | 0.328 |
| opioid use disorder – control | 0.55132 | 0.4432 | 1.2439 | 0.226 |
| opioid+alcohol use disorder – control | -0.61136 | 0.5241 | -1.1664 | 0.255 |
| Age Cateogry: | | | | |
| 50-64 – 30-49 | -0.75302 | 0.4722 | -1.5946 | 0.124 |
| 50-65 – 30-49 | -0.12093 | 0.7255 | -0.1667 | 0.869 |
| 65+ – 30-49 | 0.35551 | 0.5471 | 0.6498 | 0.522 |
| <30 – 30-49 | -0.76067 | 0.5720 | -1.3299 | 0.196 |
| Ethnicity: | | | | |
| Asian – White | -0.56781 | 0.9524 | -0.5962 | 0.557 |
| Black – White | 0.00968 | 0.3877 | 0.0250 | 0.980 |
| Hispanic – White | 0.20305 | 0.7416 | 0.2738 | 0.787 |
| Gender: | | | | |
| Male – Female | 0.97139 | 0.4583 | 2.1194 | 0.045 |
| PMI converted to hours | 0.01627 | 0.0190 | 0.8553 | 0.401 |
| pH | 0.80836 | 0.6574 | 1.2296 | 0.231 |

[a] Represents reference level

#### Linear Regression

Model Fit Measures

| Model | R | $R^2$ |
|---|---|---|
| 1 | 0.743 | 0.551 |

Model Coefficients - SLC12A5

| Predictor | Estimate | SE | t | p |
|---|---|---|---|---|
| Intercept [a] | 31.8997 | 3.9929 | 7.989 | < .001 |
| Arm: | | | | |
| alcohol use disorder – control | -0.0536 | 0.4201 | -0.128 | 0.899 |
| opioid use disorder – control | -0.5679 | 0.3930 | -1.445 | 0.161 |
| opioid+alcohol use disorder – control | 0.7525 | 0.4648 | 1.619 | 0.119 |
| Age Cateogry: | | | | |
| 50-64 – 30-49 | 0.8690 | 0.4188 | 2.075 | 0.049 |
| 50-65 – 30-49 | 0.2229 | 0.6433 | 0.346 | 0.732 |
| 65+ – 30-49 | 0.5772 | 0.4852 | 1.190 | 0.246 |
| <30 – 30-49 | 0.7818 | 0.5072 | 1.541 | 0.136 |
| Ethnicity: | | | | |
| Asian – White | 1.0487 | 0.8446 | 1.242 | 0.226 |
| Black – White | 0.5292 | 0.3438 | 1.539 | 0.137 |
| Hispanic – White | 1.0238 | 0.6576 | 1.557 | 0.133 |
| Gender: | | | | |
| Male – Female | -0.1778 | 0.4064 | -0.438 | 0.666 |
| PMI converted to hours | -0.0301 | 0.0169 | -1.782 | 0.087 |
| pH | -0.4874 | 0.5830 | -0.836 | 0.411 |

[a] Represents reference level

#### Linear Regression

Model Fit Measures

| Model | R | $R^2$ |
|---|---|---|
| 1 | 0.774 | 0.599 |

Model Coefficients - SFXN1

| Predictor | Estimate | SE | t | p |
|---|---|---|---|---|
| Intercept [a] | 30.76121 | 1.04812 | 29.34882 | < .001 |
| Arm: | | | | |
| alcohol use disorder – control | -0.04303 | 0.11029 | -0.39014 | 0.700 |
| opioid use disorder – control | 0.15999 | 0.10317 | 1.55071 | 0.134 |
| opioid+alcohol use disorder – control | -0.27728 | 0.12201 | -2.27269 | 0.032 |
| Age Cateogry: | | | | |
| 50-64 – 30-49 | -0.16687 | 0.10993 | -1.51804 | 0.142 |
| 50-65 – 30-49 | -0.05968 | 0.16887 | -0.35342 | 0.727 |
| 65+ – 30-49 | -0.05637 | 0.12735 | -0.44264 | 0.662 |
| <30 – 30-49 | -0.15964 | 0.13315 | -1.19896 | 0.242 |
| Ethnicity: | | | | |
| Asian – White | -0.60707 | 0.22171 | -2.73817 | 0.011 |
| Black – White | 4.29e-4 | 0.09026 | 0.00476 | 0.996 |
| Hispanic – White | -0.19150 | 0.17263 | -1.10933 | 0.278 |
| Gender: | | | | |
| Male – Female | 0.03857 | 0.10669 | 0.36153 | 0.721 |
| PMI converted to hours | 0.00449 | 0.00443 | 1.01354 | 0.321 |
| pH | 0.01357 | 0.15304 | 0.08865 | 0.930 |

[a] Represents reference level

#### Linear Regression

Model Fit Measures

| Model | R | $R^2$ |
|---|---|---|
| 1 | 0.779 | 0.607 |

Model Coefficients - CPNE5

| Predictor | Estimate | SE | t | p |
|---|---|---|---|---|
| Intercept [a] | 29.32716 | 1.14764 | 25.554 | < .001 |
| Arm: | | | | |
| alcohol use disorder – control | 0.12530 | 0.12076 | 1.038 | 0.310 |
| opioid use disorder – control | 0.24194 | 0.11297 | 2.142 | 0.043 |
| opioid+alcohol use disorder – control | -0.39108 | 0.13359 | -2.927 | 0.007 |
| Age Cateogry: | | | | |
| 50-64 – 30-49 | -0.02503 | 0.12036 | -0.208 | 0.837 |
| 50-65 – 30-49 | -0.20424 | 0.18491 | -1.105 | 0.280 |
| 65+ – 30-49 | 0.01798 | 0.13944 | 0.129 | 0.898 |
| <30 – 30-49 | 0.08735 | 0.14579 | 0.599 | 0.555 |
| Ethnicity: | | | | |
| Asian – White | 0.03316 | 0.24276 | 0.137 | 0.892 |
| Black – White | -0.06664 | 0.09883 | -0.674 | 0.507 |
| Hispanic – White | -0.29187 | 0.18902 | -1.544 | 0.136 |
| Gender: | | | | |
| Male – Female | -0.12134 | 0.11682 | -1.039 | 0.309 |
| PMI converted to hours | 0.00215 | 0.00485 | 0.444 | 0.661 |
| pH | -0.14905 | 0.16757 | -0.889 | 0.383 |

[a] Represents reference level

#### Linear Regression

Model Fit Measures

| Model | R | $R^2$ |
|---|---|---|
| 1 | 0.686 | 0.471 |

Model Coefficients - ICAM5

| Predictor | Estimate | SE | t | p |
|---|---|---|---|---|
| Intercept [a] | 30.03131 | 2.01206 | 14.9257 | < .001 |
| Arm: | | | | |
| alcohol use disorder – control | 0.09190 | 0.21172 | 0.4341 | 0.668 |
| opioid use disorder – control | 0.52010 | 0.19805 | 2.6261 | 0.015 |
| opioid+alcohol use disorder – control | -0.06125 | 0.23421 | -0.2615 | 0.796 |
| Age Cateogry: | | | | |
| 50-64 – 30-49 | -0.10304 | 0.21102 | -0.4883 | 0.630 |
| 50-65 – 30-49 | -0.10573 | 0.32418 | -0.3261 | 0.747 |
| 65+ – 30-49 | -0.16748 | 0.24447 | -0.6851 | 0.500 |
| <30 – 30-49 | -0.03376 | 0.25560 | -0.1321 | 0.896 |
| Ethnicity: | | | | |
| Asian – White | -0.07814 | 0.42561 | -0.1836 | 0.856 |
| Black – White | 0.01254 | 0.17327 | 0.0724 | 0.943 |
| Hispanic – White | -0.41326 | 0.33138 | -1.2471 | 0.224 |
| Gender: | | | | |
| Male – Female | 0.07330 | 0.20481 | 0.3579 | 0.724 |
| PMI converted to hours | 0.00649 | 0.00850 | 0.7641 | 0.452 |
| pH | 0.05180 | 0.29378 | 0.1763 | 0.862 |

[a] Represents reference level

#### Linear Regression

Model Fit Measures

| Model | R | $R^2$ |
|---|---|---|
| 1 | 0.696 | 0.485 |

Model Coefficients - CDV3

| Predictor | Estimate | SE | t | p |
|---|---|---|---|---|
| Intercept [a] | 25.43050 | 2.05524 | 12.3735 | < .001 |
| Arm: | | | | |
| alcohol use disorder – control | 0.19285 | 0.21626 | 0.8917 | 0.381 |
| opioid use disorder – control | 0.47814 | 0.20230 | 2.3635 | 0.027 |
| opioid+alcohol use disorder – control | -0.21269 | 0.23924 | -0.8890 | 0.383 |
| Age Cateogry: | | | | |
| 50-64 – 30-49 | -0.10692 | 0.21555 | -0.4960 | 0.624 |
| 50-65 – 30-49 | -0.09088 | 0.33114 | -0.2744 | 0.786 |
| 65+ – 30-49 | 0.00976 | 0.24972 | 0.0391 | 0.969 |
| <30 – 30-49 | -0.11150 | 0.26109 | -0.4271 | 0.673 |
| Ethnicity: | | | | |
| Asian – White | -0.51003 | 0.43474 | -1.1732 | 0.252 |
| Black – White | -0.07523 | 0.17699 | -0.4251 | 0.675 |
| Hispanic – White | -0.21932 | 0.33850 | -0.6479 | 0.523 |
| Gender: | | | | |
| Male – Female | 0.09674 | 0.20920 | 0.4624 | 0.648 |
| PMI converted to hours | 0.01360 | 0.00868 | 1.5664 | 0.130 |
| pH | 0.22563 | 0.30009 | 0.7519 | 0.459 |

[a] Represents reference level

### Linear Regression

Model Fit Measures

| Model | R | $R^2$ |
|---|---|---|
| 1 | 0.742 | 0.550 |

Model Coefficients - NRXN1

| Predictor | Estimate | SE | t | p |
|---|---|---|---|---|
| Intercept [a] | 26.8813 | 2.12077 | 12.6752 | < .001 |
| Arm: | | | | |
| alcohol use disorder – control | 0.0647 | 0.22316 | 0.2900 | 0.774 |
| opioid use disorder – control | 0.3916 | 0.20875 | 1.8760 | 0.073 |
| opioid+alcohol use disorder – control | -0.3999 | 0.24687 | -1.6198 | 0.118 |
| Age Cateogry: | | | | |
| 50-64 – 30-49 | -0.2482 | 0.22242 | -1.1160 | 0.275 |
| 50-65 – 30-49 | 0.3020 | 0.34170 | 0.8839 | 0.386 |
| 65+ – 30-49 | 0.0723 | 0.25768 | 0.2805 | 0.781 |
| <30 – 30-49 | -0.0422 | 0.26941 | -0.1565 | 0.877 |
| Ethnicity: | | | | |
| Asian – White | -0.4391 | 0.44860 | -0.9788 | 0.337 |
| Black – White | 0.1854 | 0.18263 | 1.0152 | 0.320 |
| Hispanic – White | -0.0130 | 0.34929 | -0.0372 | 0.971 |
| Gender: | | | | |
| Male – Female | 0.0865 | 0.21587 | 0.4005 | 0.692 |
| PMI converted to hours | 0.0128 | 0.00896 | 1.4243 | 0.167 |
| pH | -0.0648 | 0.30966 | -0.2094 | 0.836 |

[a] Represents reference level

### Linear Regression

Model Fit Measures

| Model | R | $R^2$ |
|---|---|---|
| 1 | 0.724 | 0.524 |

Model Coefficients - G3BP2

| Predictor | Estimate | SE | t | p |
|---|---|---|---|---|
| Intercept [a] | 30.57645 | 3.3975 | 9.000 | < .001 |
| Arm: | | | | |
| alcohol use disorder – control | -0.14228 | 0.3575 | -0.398 | 0.694 |
| opioid use disorder – control | -0.75290 | 0.3344 | -2.251 | 0.034 |
| opioid+alcohol use disorder – control | 0.04737 | 0.3955 | 0.120 | 0.906 |
| Age Cateogry: | | | | |
| 50-64 – 30-49 | 0.65297 | 0.3563 | 1.833 | 0.079 |
| 50-65 – 30-49 | -0.15373 | 0.5474 | -0.281 | 0.781 |
| 65+ – 30-49 | 0.74773 | 0.4128 | 1.811 | 0.083 |
| <30 – 30-49 | 0.58876 | 0.4316 | 1.364 | 0.185 |
| Ethnicity: | | | | |
| Asian – White | 0.94341 | 0.7187 | 1.313 | 0.202 |
| Black – White | 0.32120 | 0.2926 | 1.098 | 0.283 |
| Hispanic – White | 0.16503 | 0.5596 | 0.295 | 0.771 |
| Gender: | | | | |
| Male – Female | -0.31910 | 0.3458 | -0.923 | 0.365 |
| PMI converted to hours | -0.00870 | 0.0143 | -0.607 | 0.550 |
| pH | -0.57226 | 0.4961 | -1.154 | 0.260 |

[a] Represents reference level

### Linear Regression

Model Fit Measures

| Model | R | $R^2$ |
|---|---|---|
| 1 | 0.766 | 0.587 |

Model Coefficients - TMED7

| Predictor | Estimate | SE | t | p |
|---|---|---|---|---|
| Intercept [a] | 16.56781 | 4.1770 | 3.9665 | < .001 |
| Arm: | | | | |
| alcohol use disorder – control | 0.00946 | 0.4395 | 0.0215 | 0.983 |
| opioid use disorder – control | 0.70824 | 0.4112 | 1.7226 | 0.098 |
| opioid+alcohol use disorder – control | -0.76641 | 0.4862 | -1.5763 | 0.128 |
| Age Cateogry: | | | | |
| 50-64 – 30-49 | -0.41645 | 0.4381 | -0.9506 | 0.351 |
| 50-65 – 30-49 | -0.65060 | 0.6730 | -0.9667 | 0.343 |
| 65+ – 30-49 | -0.06701 | 0.5075 | -0.1320 | 0.896 |
| <30 – 30-49 | -0.95104 | 0.5306 | -1.7923 | 0.086 |
| Ethnicity: | | | | |
| Asian – White | -0.52847 | 0.8835 | -0.5981 | 0.555 |
| Black – White | -0.05863 | 0.3597 | -0.1630 | 0.872 |
| Hispanic – White | -0.27893 | 0.6879 | -0.4055 | 0.689 |
| Gender: | | | | |
| Male – Female | 0.15856 | 0.4252 | 0.3729 | 0.712 |
| PMI converted to hours | -0.01065 | 0.0176 | -0.6038 | 0.552 |
| pH | 1.17530 | 0.6099 | 1.9271 | 0.066 |

[a] Represents reference level

### Linear Regression

Model Fit Measures

| Model | R | $R^2$ |
|---|---|---|
| 1 | 0.631 | 0.398 |

Model Coefficients - CSNK1G3

| Predictor | Estimate | SE | t | p |
|---|---|---|---|---|
| Intercept [a] | 21.74315 | 3.0342 | 7.166 | < .001 |
| Arm: | | | | |
| alcohol use disorder – control | -0.04744 | 0.3193 | -0.149 | 0.883 |
| opioid use disorder – control | 0.55589 | 0.2987 | 1.861 | 0.075 |
| opioid+alcohol use disorder – control | -0.36623 | 0.3532 | -1.037 | 0.310 |
| Age Cateogry: | | | | |
| 50-64 – 30-49 | 0.28212 | 0.3182 | 0.887 | 0.384 |
| 50-65 – 30-49 | 0.52196 | 0.4889 | 1.068 | 0.296 |
| 65+ – 30-49 | 0.25229 | 0.3687 | 0.684 | 0.500 |
| <30 – 30-49 | 0.26774 | 0.3855 | 0.695 | 0.494 |
| Ethnicity: | | | | |
| Asian – White | 0.41495 | 0.6418 | 0.647 | 0.524 |
| Black – White | -0.12125 | 0.2613 | -0.464 | 0.647 |
| Hispanic – White | -0.05525 | 0.4997 | -0.111 | 0.913 |
| Gender: | | | | |
| Male – Female | -0.31243 | 0.3089 | -1.012 | 0.322 |
| PMI converted to hours | 0.00400 | 0.0128 | 0.312 | 0.758 |
| pH | 0.22900 | 0.4430 | 0.517 | 0.610 |

[a] Represents reference level

### Linear Regression

Model Fit Measures

| Model | R | $R^2$ |
|---|---|---|
| 1 | 0.609 | 0.371 |

Model Coefficients - ZNF207

| Predictor | Estimate | SE | t | p |
|---|---|---|---|---|
| Intercept [a] | 21.93789 | 3.3482 | 6.55205 | < .001 |
| Arm: | | | | |
| alcohol use disorder – control | -0.17293 | 0.3523 | -0.49083 | 0.628 |
| opioid use disorder – control | 0.34488 | 0.3296 | 1.04642 | 0.306 |
| opioid+alcohol use disorder – control | -0.79385 | 0.3898 | -2.03682 | 0.053 |
| Age Cateogry: | | | | |
| 50-64 – 30-49 | 0.24928 | 0.3512 | 0.70989 | 0.485 |
| 50-65 – 30-49 | 0.28493 | 0.5395 | 0.52817 | 0.602 |
| 65+ – 30-49 | 0.12473 | 0.4068 | 0.30660 | 0.762 |
| <30 – 30-49 | -0.19667 | 0.4253 | -0.46237 | 0.648 |
| Ethnicity: | | | | |
| Asian – White | -0.09821 | 0.7082 | -0.13866 | 0.891 |
| Black – White | -0.13604 | 0.2883 | -0.47181 | 0.641 |
| Hispanic – White | -0.33433 | 0.5515 | -0.60627 | 0.550 |
| Gender: | | | | |
| Male – Female | -6.73e−4 | 0.3408 | -0.00197 | 0.998 |
| PMI converted to hours | 0.00200 | 0.0141 | 0.14168 | 0.889 |
| pH | 0.22082 | 0.4889 | 0.45168 | 0.656 |

[a] Represents reference level

### Linear Regression

Model Fit Measures

| Model | R | $R^2$ |
|---|---|---|
| 1 | 0.651 | 0.424 |

Model Coefficients - F2

| Predictor | Estimate | SE | t | p |
|---|---|---|---|---|
| Intercept [a] | 23.2969 | 5.0090 | 4.6510 | < .001 |
| Arm: | | | | |
| alcohol use disorder – control | -0.3947 | 0.5271 | -0.7489 | 0.461 |
| opioid use disorder – control | 0.2104 | 0.4930 | 0.4266 | 0.673 |
| opioid+alcohol use disorder – control | -1.5510 | 0.5831 | -2.6600 | 0.014 |
| Age Cateogry: | | | | |
| 50-64 – 30-49 | 0.1898 | 0.5253 | 0.3613 | 0.721 |
| 50-65 – 30-49 | 0.1110 | 0.8070 | 0.1376 | 0.892 |
| 65+ – 30-49 | 0.4576 | 0.6086 | 0.7519 | 0.459 |
| <30 – 30-49 | 0.0385 | 0.6363 | 0.0606 | 0.952 |
| Ethnicity: | | | | |
| Asian – White | -0.2048 | 1.0595 | -0.1933 | 0.848 |
| Black – White | -0.6015 | 0.4313 | -1.3944 | 0.176 |
| Hispanic – White | -1.0556 | 0.8250 | -1.2795 | 0.213 |
| Gender: | | | | |
| Male – Female | -0.6208 | 0.5099 | -1.2175 | 0.235 |
| PMI converted to hours | 0.0221 | 0.0212 | 1.0466 | 0.306 |
| pH | 0.0680 | 0.7314 | 0.0929 | 0.927 |

[a] Represents reference level

#### Linear Regression

Model Fit Measures

| Model | R | $R^2$ |
|---|---|---|
| 1 | 0.623 | 0.388 |

Model Coefficients - FAM171A2

| Predictor | Estimate | SE | t | p |
|---|---|---|---|---|
| Intercept [a] | 29.23980 | 4.5516 | 6.4241 | < .001 |
| Arm: | | | | |
| alcohol use disorder – control | 0.59883 | 0.4789 | 1.2503 | 0.223 |
| opioid use disorder – control | 0.15669 | 0.4480 | 0.3497 | 0.730 |
| opioid+alcohol use disorder – control | -1.34087 | 0.5298 | -2.5308 | 0.018 |
| Age Cateogry: | | | | |
| 50-64 – 30-49 | 0.29432 | 0.4774 | 0.6166 | 0.543 |
| 50-65 – 30-49 | -0.22082 | 0.7334 | -0.3011 | 0.766 |
| 65+ – 30-49 | 0.03661 | 0.5530 | 0.0662 | 0.948 |
| <30 – 30-49 | 0.87245 | 0.5782 | 1.5089 | 0.144 |
| Ethnicity: | | | | |
| Asian – White | -0.41722 | 0.9628 | -0.4333 | 0.669 |
| Black – White | -0.03951 | 0.3920 | -0.1008 | 0.921 |
| Hispanic – White | -0.23263 | 0.7496 | -0.3103 | 0.759 |
| Gender: | | | | |
| Male – Female | -0.88210 | 0.4633 | -1.9039 | 0.069 |
| PMI converted to hours | 0.00186 | 0.0192 | 0.0966 | 0.924 |
| pH | -0.73534 | 0.6646 | -1.1065 | 0.279 |

[a] Represents reference level

#### Linear Regression

Model Fit Measures

| Model | R | $R^2$ |
|---|---|---|
| 1 | 0.613 | 0.375 |

Model Coefficients - COCH

| Predictor | Estimate | SE | t | p |
|---|---|---|---|---|
| Intercept [a] | 22.3753 | 5.2870 | 4.232 | < .001 |
| Arm: | | | | |
| alcohol use disorder – control | 0.3114 | 0.5563 | 0.560 | 0.581 |
| opioid use disorder – control | 0.5406 | 0.5204 | 1.039 | 0.309 |
| opioid+alcohol use disorder – control | -0.8719 | 0.6154 | -1.417 | 0.169 |
| Age Cateogry: | | | | |
| 50-64 – 30-49 | 0.4802 | 0.5545 | 0.866 | 0.395 |
| 50-65 – 30-49 | 0.4078 | 0.8519 | 0.479 | 0.636 |
| 65+ – 30-49 | 1.6944 | 0.6424 | 2.638 | 0.014 |
| <30 – 30-49 | 0.2782 | 0.6716 | 0.414 | 0.682 |
| Ethnicity: | | | | |
| Asian – White | 0.5607 | 1.1184 | 0.501 | 0.621 |
| Black – White | -0.1716 | 0.4553 | -0.377 | 0.710 |
| Hispanic – White | 0.5508 | 0.8708 | 0.633 | 0.533 |
| Gender: | | | | |
| Male – Female | 0.2565 | 0.5382 | 0.477 | 0.638 |
| PMI converted to hours | -0.0129 | 0.0223 | -0.580 | 0.568 |
| pH | 0.1282 | 0.7720 | 0.166 | 0.870 |

[a] Represents reference level

### Linear Regression

Model Fit Measures

| Model | R | $R^2$ |
|---|---|---|
| 1 | 0.650 | 0.423 |

Model Coefficients - COX7A2L

| Predictor | Estimate | SE | t | p |
|---|---|---|---|---|
| Intercept [a] | 30.67335 | 0.99279 | 30.896 | < .001 |
| Arm: | | | | |
| alcohol use disorder – control | -0.07544 | 0.10447 | -0.722 | 0.477 |
| opioid use disorder – control | -0.01294 | 0.09772 | -0.132 | 0.896 |
| opioid+alcohol use disorder – control | -0.28581 | 0.11557 | -2.473 | 0.021 |
| Age Cateogry: | | | | |
| 50-64 – 30-49 | -0.08662 | 0.10412 | -0.832 | 0.414 |
| 50-65 – 30-49 | -0.08855 | 0.15996 | -0.554 | 0.585 |
| 65+ – 30-49 | -0.05283 | 0.12063 | -0.438 | 0.665 |
| <30 – 30-49 | 0.09310 | 0.12612 | 0.738 | 0.468 |
| Ethnicity: | | | | |
| Asian – White | 0.06058 | 0.21000 | 0.288 | 0.775 |
| Black – White | -0.06494 | 0.08549 | -0.760 | 0.455 |
| Hispanic – White | -0.14598 | 0.16351 | -0.893 | 0.381 |
| Gender: | | | | |
| Male – Female | -0.08724 | 0.10106 | -0.863 | 0.397 |
| PMI converted to hours | 0.00281 | 0.00419 | 0.671 | 0.509 |
| pH | -0.23754 | 0.14496 | -1.639 | 0.114 |

[a] Represents reference level

### Linear Regression

Model Fit Measures

| Model | R | $R^2$ |
|---|---|---|
| 1 | 0.574 | 0.329 |

Model Coefficients - OPHN1

| Predictor | Estimate | SE | t | p |
|---|---|---|---|---|
| Intercept [a] | 24.61025 | 3.3385 | 7.3717 | < .001 |
| Arm: | | | | |
| alcohol use disorder – control | -0.13269 | 0.3513 | -0.3777 | 0.709 |
| opioid use disorder – control | 0.09674 | 0.3286 | 0.2944 | 0.771 |
| opioid+alcohol use disorder – control | -0.91779 | 0.3886 | -2.3617 | 0.027 |
| Age Cateogry: | | | | |
| 50-64 – 30-49 | 0.12757 | 0.3501 | 0.3644 | 0.719 |
| 50-65 – 30-49 | 0.28284 | 0.5379 | 0.5258 | 0.604 |
| 65+ – 30-49 | 0.17065 | 0.4056 | 0.4207 | 0.678 |
| <30 – 30-49 | 0.02442 | 0.4241 | 0.0576 | 0.955 |
| Ethnicity: | | | | |
| Asian – White | -0.34411 | 0.7062 | -0.4873 | 0.630 |
| Black – White | -0.18013 | 0.2875 | -0.6265 | 0.537 |
| Hispanic – White | -0.44432 | 0.5498 | -0.8081 | 0.427 |
| Gender: | | | | |
| Male – Female | -0.35117 | 0.3398 | -1.0334 | 0.312 |
| PMI converted to hours | -0.00157 | 0.0141 | -0.1114 | 0.912 |
| pH | -0.11471 | 0.4875 | -0.2353 | 0.816 |

[a] Represents reference level

### Linear Regression

Model Fit Measures

| Model | R | $R^2$ |
|---|---|---|
| 1 | 0.646 | 0.417 |

Model Coefficients - LUC7L3

| Predictor | Estimate | SE | t | p |
|---|---|---|---|---|
| Intercept [a] | 24.10210 | 2.8084 | 8.5823 | < .001 |
| Arm: | | | | |
| alcohol use disorder – control | -0.35039 | 0.2955 | -1.1857 | 0.247 |
| opioid use disorder – control | -0.13895 | 0.2764 | -0.5027 | 0.620 |
| opioid+alcohol use disorder – control | -1.08394 | 0.3269 | -3.3158 | 0.003 |
| Age Cateogry: | | | | |
| 50-64 – 30-49 | -0.08084 | 0.2945 | -0.2745 | 0.786 |
| 50-65 – 30-49 | -0.28662 | 0.4525 | -0.6334 | 0.532 |
| 65+ – 30-49 | 0.12666 | 0.3412 | 0.3712 | 0.714 |
| <30 – 30-49 | -0.12578 | 0.3568 | -0.3526 | 0.727 |
| Ethnicity: | | | | |
| Asian – White | -0.75574 | 0.5940 | -1.2722 | 0.215 |
| Black – White | -0.30291 | 0.2418 | -1.2525 | 0.222 |
| Hispanic – White | -0.45103 | 0.4625 | -0.9751 | 0.339 |
| Gender: | | | | |
| Male – Female | -0.22648 | 0.2859 | -0.7923 | 0.436 |
| PMI converted to hours | 0.00702 | 0.0119 | 0.5917 | 0.560 |
| pH | -0.02407 | 0.4101 | -0.0587 | 0.954 |

[a] Represents reference level

### Linear Regression

Model Fit Measures

| Model | R | $R^2$ |
|---|---|---|
| 1 | 0.717 | 0.514 |

Model Coefficients - KPNA4

| Predictor | Estimate | SE | t | p |
|---|---|---|---|---|
| Intercept [a] | 34.82005 | 4.6403 | 7.5039 | < .001 |
| Arm: | | | | |
| alcohol use disorder – control | 0.23090 | 0.4883 | 0.4729 | 0.641 |
| opioid use disorder – control | 0.87896 | 0.4568 | 1.9244 | 0.066 |
| opioid+alcohol use disorder – control | 1.47094 | 0.5401 | 2.7232 | 0.012 |
| Age Cateogry: | | | | |
| 50-64 – 30-49 | 0.57641 | 0.4867 | 1.1844 | 0.248 |
| 50-65 – 30-49 | -0.78057 | 0.7476 | -1.0441 | 0.307 |
| 65+ – 30-49 | 0.47510 | 0.5638 | 0.8427 | 0.408 |
| <30 – 30-49 | 0.35008 | 0.5895 | 0.5939 | 0.558 |
| Ethnicity: | | | | |
| Asian – White | 1.06653 | 0.9815 | 1.0866 | 0.288 |
| Black – White | -0.01822 | 0.3996 | -0.0456 | 0.964 |
| Hispanic – White | 0.27052 | 0.7642 | 0.3540 | 0.726 |
| Gender: | | | | |
| Male – Female | 0.41304 | 0.4723 | 0.8745 | 0.391 |
| PMI converted to hours | -0.00163 | 0.0196 | -0.0831 | 0.934 |
| pH | -1.63734 | 0.6775 | -2.4166 | 0.024 |

[a] Represents reference level

### Linear Regression

Model Fit Measures

| Model | R | $R^2$ |
|---|---|---|
| 1 | 0.709 | 0.503 |

Model Coefficients - ATG13

| Predictor | Estimate | SE | t | p |
|---|---|---|---|---|
| Intercept [a] | 25.80242 | 2.7695 | 9.317 | < .001 |
| Arm: | | | | |
| alcohol use disorder – control | -0.45336 | 0.2914 | -1.556 | 0.133 |
| opioid use disorder – control | 0.32865 | 0.2726 | 1.206 | 0.240 |
| opioid+alcohol use disorder – control | -0.92769 | 0.3224 | -2.878 | 0.008 |
| Age Cateogry: | | | | |
| 50-64 – 30-49 | 0.18095 | 0.2905 | 0.623 | 0.539 |
| 50-65 – 30-49 | 0.22378 | 0.4462 | 0.502 | 0.621 |
| 65+ – 30-49 | 0.32727 | 0.3365 | 0.973 | 0.340 |
| <30 – 30-49 | -0.07345 | 0.3518 | -0.209 | 0.836 |
| Ethnicity: | | | | |
| Asian – White | -0.45503 | 0.5858 | -0.777 | 0.445 |
| Black – White | -0.10930 | 0.2385 | -0.458 | 0.651 |
| Hispanic – White | -0.06812 | 0.4561 | -0.149 | 0.883 |
| Gender: | | | | |
| Male – Female | -0.04197 | 0.2819 | -0.149 | 0.883 |
| PMI converted to hours | 0.00888 | 0.0117 | 0.759 | 0.455 |
| pH | -0.38478 | 0.4044 | -0.952 | 0.351 |

[a] Represents reference level

### Linear Regression

Model Fit Measures

| Model | R | $R^2$ |
|---|---|---|
| 1 | 0.769 | 0.591 |

Model Coefficients - PRKCG (2)

| Predictor | Estimate | SE | t | p |
|---|---|---|---|---|
| Intercept [a] | 30.72391 | 1.34268 | 22.882 | < .001 |
| Arm: | | | | |
| alcohol use disorder – control | -0.16468 | 0.14128 | -1.166 | 0.255 |
| opioid use disorder – control | -0.08404 | 0.13216 | -0.636 | 0.531 |
| opioid+alcohol use disorder – control | 0.24645 | 0.15629 | 1.577 | 0.128 |
| Age Cateogry: | | | | |
| 50-64 – 30-49 | 0.20642 | 0.14082 | 1.466 | 0.156 |
| 50-65 – 30-49 | -0.17942 | 0.21633 | -0.829 | 0.415 |
| 65+ – 30-49 | 0.10709 | 0.16314 | 0.656 | 0.518 |
| <30 – 30-49 | 0.27206 | 0.17057 | 1.595 | 0.124 |
| Ethnicity: | | | | |
| Asian – White | 0.19233 | 0.28402 | 0.677 | 0.505 |
| Black – White | 0.06061 | 0.11563 | 0.524 | 0.605 |
| Hispanic – White | -0.09067 | 0.22114 | -0.410 | 0.685 |
| Gender: | | | | |
| Male – Female | -0.23532 | 0.13667 | -1.722 | 0.098 |
| PMI converted to hours | -0.00654 | 0.00567 | -1.154 | 0.260 |
| pH | -0.31928 | 0.19605 | -1.629 | 0.116 |

[a] Represents reference level

#### Linear Regression

Model Fit Measures

| Model | R | $R^2$ |
|---|---|---|
| 1 | 0.705 | 0.497 |

Model Coefficients - CTSB

| Predictor | Estimate | SE | t | p |
|---|---|---|---|---|
| Intercept [a] | 28.66816 | 1.34146 | 21.371 | < .001 |
| Arm: | | | | |
| alcohol use disorder – control | 0.17681 | 0.14115 | 1.253 | 0.222 |
| opioid use disorder – control | -0.10964 | 0.13204 | -0.830 | 0.415 |
| opioid+alcohol use disorder – control | 0.31994 | 0.15615 | 2.049 | 0.052 |
| Age Cateogry: | | | | |
| 50-64 – 30-49 | 0.03081 | 0.14069 | 0.219 | 0.829 |
| 50-65 – 30-49 | 0.05417 | 0.21614 | 0.251 | 0.804 |
| 65+ – 30-49 | -0.18540 | 0.16299 | -1.137 | 0.267 |
| <30 – 30-49 | 0.07686 | 0.17041 | 0.451 | 0.656 |
| Ethnicity: | | | | |
| Asian – White | 0.17071 | 0.28376 | 0.602 | 0.553 |
| Black – White | -0.18311 | 0.11552 | -1.585 | 0.126 |
| Hispanic – White | -0.07519 | 0.22094 | -0.340 | 0.737 |
| Gender: | | | | |
| Male – Female | -0.23101 | 0.13655 | -1.692 | 0.104 |
| PMI converted to hours | -0.00405 | 0.00567 | -0.715 | 0.481 |
| pH | -0.04938 | 0.19587 | -0.252 | 0.803 |

[a] Represents reference level

#### Linear Regression

Model Fit Measures

| Model | R | $R^2$ |
|---|---|---|
| 1 | 0.626 | 0.392 |

Model Coefficients - PTPRA

| Predictor | Estimate | SE | t | p |
|---|---|---|---|---|
| Intercept [a] | 26.47853 | 4.5590 | 5.8079 | < .001 |
| Arm: | | | | |
| alcohol use disorder – control | -0.21769 | 0.4797 | -0.4538 | 0.654 |
| opioid use disorder – control | 0.41009 | 0.4488 | 0.9138 | 0.370 |
| opioid+alcohol use disorder – control | 1.17421 | 0.5307 | 2.2126 | 0.037 |
| Age Cateogry: | | | | |
| 50-64 – 30-49 | -0.25747 | 0.4781 | -0.5385 | 0.595 |
| 50-65 – 30-49 | -0.57513 | 0.7346 | -0.7830 | 0.441 |
| 65+ – 30-49 | -0.83622 | 0.5539 | -1.5096 | 0.144 |
| <30 – 30-49 | 0.02783 | 0.5792 | 0.0481 | 0.962 |
| Ethnicity: | | | | |
| Asian – White | -0.13162 | 0.9644 | -0.1365 | 0.893 |
| Black – White | 0.02926 | 0.3926 | 0.0745 | 0.941 |
| Hispanic – White | 0.11959 | 0.7509 | 0.1593 | 0.875 |
| Gender: | | | | |
| Male – Female | 0.24739 | 0.4641 | 0.5331 | 0.599 |
| PMI converted to hours | 0.00317 | 0.0193 | 0.1644 | 0.871 |
| pH | -0.28801 | 0.6657 | -0.4327 | 0.669 |

[a] Represents reference level

#### Linear Regression

Model Fit Measures

| Model | R | $R^2$ |
|---|---|---|
| 1 | 0.642 | 0.412 |

Model Coefficients - LAMC1

| Predictor | Estimate | SE | t | p |
|---|---|---|---|---|
| Intercept [a] | 24.59494 | 3.1265 | 7.867 | < .001 |
| Arm: | | | | |
| alcohol use disorder – control | -0.14321 | 0.3290 | -0.435 | 0.667 |
| opioid use disorder – control | 0.15460 | 0.3078 | 0.502 | 0.620 |
| opioid+alcohol use disorder – control | -0.94008 | 0.3639 | -2.583 | 0.016 |
| Age Cateogry: | | | | |
| 50-64 – 30-49 | -0.06061 | 0.3279 | -0.185 | 0.855 |
| 50-65 – 30-49 | -0.34753 | 0.5037 | -0.690 | 0.497 |
| 65+ – 30-49 | 0.36306 | 0.3799 | 0.956 | 0.349 |
| <30 – 30-49 | -0.17291 | 0.3972 | -0.435 | 0.667 |
| Ethnicity: | | | | |
| Asian – White | 0.26691 | 0.6613 | 0.404 | 0.690 |
| Black – White | 0.08458 | 0.2692 | 0.314 | 0.756 |
| Hispanic – White | 0.07085 | 0.5149 | 0.138 | 0.892 |
| Gender: | | | | |
| Male – Female | -0.16250 | 0.3182 | -0.511 | 0.614 |
| PMI converted to hours | 0.00638 | 0.0132 | 0.484 | 0.633 |
| pH | -0.17192 | 0.4565 | -0.377 | 0.710 |

[a] Represents reference level

#### Linear Regression

Model Fit Measures

| Model | R | $R^2$ |
|---|---|---|
| 1 | 0.467 | 0.218 |

Model Coefficients - DSP

| Predictor | Estimate | SE | t | p |
|---|---|---|---|---|
| Intercept [a] | 27.01641 | 6.2653 | 4.3121 | < .001 |
| Arm: | | | | |
| alcohol use disorder – control | 0.35762 | 0.6593 | 0.5425 | 0.593 |
| opioid use disorder – control | 0.03768 | 0.6167 | 0.0611 | 0.952 |
| opioid+alcohol use disorder – control | -1.05765 | 0.7293 | -1.4502 | 0.160 |
| Age Cateogry: | | | | |
| 50-64 – 30-49 | 0.03886 | 0.6571 | 0.0591 | 0.953 |
| 50-65 – 30-49 | -0.53713 | 1.0095 | -0.5321 | 0.600 |
| 65+ – 30-49 | 0.49361 | 0.7613 | 0.6484 | 0.523 |
| <30 – 30-49 | -0.03719 | 0.7959 | -0.0467 | 0.963 |
| Ethnicity: | | | | |
| Asian – White | -0.56040 | 1.3253 | -0.4229 | 0.676 |
| Black – White | 0.24680 | 0.5395 | 0.4574 | 0.651 |
| Hispanic – White | -0.93081 | 1.0319 | -0.9020 | 0.376 |
| Gender: | | | | |
| Male – Female | -0.38226 | 0.6377 | -0.5994 | 0.555 |
| PMI converted to hours | 0.00285 | 0.0265 | 0.1075 | 0.915 |
| pH | -0.47097 | 0.9148 | -0.5148 | 0.611 |

[a] Represents reference level

### Results

#### Linear Regression

Model Fit Measures

| Model | R | $R^2$ |
|---|---|---|
| 1 | 0.755 | 0.570 |

Model Coefficients - EPB41L3

| Predictor | Estimate | SE | t | p |
|---|---|---|---|---|
| Intercept [a] | 32.14901 | 1.54095 | 20.8631 | < .001 |
| Arm: | | | | |
| alcohol use disorder – control | -0.10725 | 0.16215 | -0.6614 | 0.515 |
| opioid use disorder – control | -0.31379 | 0.15168 | -2.0687 | 0.050 |
| opioid+alcohol use disorder – control | -0.08446 | 0.17937 | -0.4709 | 0.642 |
| Age Cateogry: | | | | |
| 50-64 – 30-49 | 0.43523 | 0.16161 | 2.6931 | 0.013 |
| 50-65 – 30-49 | 0.05145 | 0.24828 | 0.2072 | 0.838 |
| 65+ – 30-49 | 0.40684 | 0.18723 | 2.1729 | 0.040 |
| <30 – 30-49 | 0.25170 | 0.19575 | 1.2858 | 0.211 |
| Ethnicity: | | | | |
| Asian – White | 0.28505 | 0.32595 | 0.8745 | 0.391 |
| Black – White | -0.09799 | 0.13270 | -0.7385 | 0.467 |
| Hispanic – White | 0.05964 | 0.25379 | 0.2350 | 0.816 |
| Gender: | | | | |
| Male – Female | -0.14803 | 0.15685 | -0.9438 | 0.355 |
| PMI converted to hours | -0.00546 | 0.00651 | -0.8392 | 0.410 |
| pH | 0.00429 | 0.22500 | 0.0191 | 0.985 |

[a] Represents reference level

#### Linear Regression

Model Fit Measures

| Model | R | $R^2$ |
|---|---|---|
| 1 | 0.734 | 0.539 |

Model Coefficients - SH3BGRL2

| Predictor | Estimate | SE | t | p |
|---|---|---|---|---|
| Intercept [a] | 38.8002 | 8.6779 | 4.471 | < .001 |
| Arm: | | | | |
| alcohol use disorder – control | -0.7637 | 0.9131 | -0.836 | 0.411 |
| opioid use disorder – control | -2.4920 | 0.8542 | -2.917 | 0.008 |
| opioid+alcohol use disorder – control | 0.1111 | 1.0101 | 0.110 | 0.913 |
| Age Cateogry: | | | | |
| 50-64 – 30-49 | 1.4226 | 0.9101 | 1.563 | 0.131 |
| 50-65 – 30-49 | -1.3336 | 1.3982 | -0.954 | 0.350 |
| 65+ – 30-49 | -0.6594 | 1.0544 | -0.625 | 0.538 |
| <30 – 30-49 | 1.2076 | 1.1024 | 1.095 | 0.284 |
| Ethnicity: | | | | |
| Asian – White | 0.8803 | 1.8356 | 0.480 | 0.636 |
| Black – White | -0.4564 | 0.7473 | -0.611 | 0.547 |
| Hispanic – White | -0.7364 | 1.4292 | -0.515 | 0.611 |
| Gender: | | | | |
| Male – Female | -0.3285 | 0.8833 | -0.372 | 0.713 |
| PMI converted to hours | -0.0656 | 0.0367 | -1.791 | 0.086 |
| pH | -1.5709 | 1.2671 | -1.240 | 0.227 |

[a] Represents reference level

### Linear Regression

Model Fit Measures

| Model | R | $R^2$ |
|---|---|---|
| 1 | 0.572 | 0.327 |

Model Coefficients - ACSL6

| Predictor | Estimate | SE | t | p |
|---|---|---|---|---|
| Intercept [a] | 27.8660 | 3.6064 | 7.7269 | < .001 |
| Arm: | | | | |
| alcohol use disorder – control | -0.3936 | 0.3795 | -1.0373 | 0.310 |
| opioid use disorder – control | -0.5676 | 0.3550 | -1.5990 | 0.123 |
| opioid+alcohol use disorder – control | 0.2696 | 0.4198 | 0.6422 | 0.527 |
| Age Cateogry: | | | | |
| 50-64 – 30-49 | 0.1595 | 0.3782 | 0.4218 | 0.677 |
| 50-65 – 30-49 | 0.2443 | 0.5811 | 0.4204 | 0.678 |
| 65+ – 30-49 | -0.2817 | 0.4382 | -0.6429 | 0.526 |
| <30 – 30-49 | 0.1267 | 0.4581 | 0.2766 | 0.784 |
| Ethnicity: | | | | |
| Asian – White | 0.1745 | 0.7628 | 0.2288 | 0.821 |
| Black – White | 0.1659 | 0.3106 | 0.5340 | 0.598 |
| Hispanic – White | 0.5387 | 0.5940 | 0.9069 | 0.373 |
| Gender: | | | | |
| Male – Female | -0.1368 | 0.3671 | -0.3725 | 0.713 |
| PMI converted to hours | 5.49e-4 | 0.0152 | 0.0361 | 0.972 |
| pH | 0.0211 | 0.5266 | 0.0401 | 0.968 |

[a] Represents reference level

### Linear Regression

Model Fit Measures

| Model | R | $R^2$ |
|---|---|---|
| 1 | 0.700 | 0.491 |

Model Coefficients - TBC1D24

| Predictor | Estimate | SE | t | p |
|---|---|---|---|---|
| Intercept [a] | 28.42581 | 1.14640 | 24.7957 | < .001 |
| Arm: | | | | |
| alcohol use disorder – control | 0.06029 | 0.12063 | 0.4998 | 0.622 |
| opioid use disorder – control | 0.24527 | 0.11284 | 2.1736 | 0.040 |
| opioid+alcohol use disorder – control | 0.10338 | 0.13345 | 0.7747 | 0.446 |
| Age Cateogry: | | | | |
| 50-64 – 30-49 | -0.18665 | 0.12023 | -1.5524 | 0.134 |
| 50-65 – 30-49 | -0.02089 | 0.18471 | -0.1131 | 0.911 |
| 65+ – 30-49 | -0.26423 | 0.13929 | -1.8969 | 0.070 |
| <30 – 30-49 | -0.22412 | 0.14563 | -1.5390 | 0.137 |
| Ethnicity: | | | | |
| Asian – White | -0.26744 | 0.24250 | -1.1028 | 0.281 |
| Black – White | 0.11102 | 0.09872 | 1.1246 | 0.272 |
| Hispanic – White | 0.01530 | 0.18881 | 0.0810 | 0.936 |
| Gender: | | | | |
| Male – Female | 0.08399 | 0.11669 | 0.7198 | 0.479 |
| PMI converted to hours | 0.00403 | 0.00484 | 0.8314 | 0.414 |
| pH | -0.07363 | 0.16739 | -0.4399 | 0.664 |

[a] Represents reference level

### Linear Regression

Model Fit Measures

| Model | R | $R^2$ |
|---|---|---|
| 1 | 0.585 | 0.342 |

Model Coefficients - OGDHL

| Predictor | Estimate | SE | t | p |
|---|---|---|---|---|
| Intercept [a] | 28.27090 | 3.6298 | 7.7885 | < .001 |
| Arm: | | | | |
| alcohol use disorder – control | -0.33574 | 0.3819 | -0.8790 | 0.388 |
| opioid use disorder – control | -0.46015 | 0.3573 | -1.2879 | 0.210 |
| opioid+alcohol use disorder – control | 0.09944 | 0.4225 | 0.2353 | 0.816 |
| Age Cateogry: | | | | |
| 50-64 – 30-49 | 0.40539 | 0.3807 | 1.0649 | 0.298 |
| 50-65 – 30-49 | -0.04210 | 0.5848 | -0.0720 | 0.943 |
| 65+ – 30-49 | 0.28206 | 0.4410 | 0.6395 | 0.529 |
| <30 – 30-49 | 0.50515 | 0.4611 | 1.0955 | 0.284 |
| Ethnicity: | | | | |
| Asian – White | 0.62610 | 0.7678 | 0.8154 | 0.423 |
| Black – White | 0.20352 | 0.3126 | 0.6511 | 0.521 |
| Hispanic – White | 0.68876 | 0.5978 | 1.1521 | 0.261 |
| Gender: | | | | |
| Male – Female | -0.05602 | 0.3695 | -0.1516 | 0.881 |
| PMI converted to hours | -0.00480 | 0.0153 | -0.3134 | 0.757 |
| pH | 0.04000 | 0.5300 | 0.0755 | 0.940 |

[a] Represents reference level

### Linear Regression

Model Fit Measures

| Model | R | $R^2$ |
|---|---|---|
| 1 | 0.550 | 0.302 |

Model Coefficients - CADPS

| Predictor | Estimate | SE | t | p |
|---|---|---|---|---|
| Intercept [a] | 29.1511 | 1.53520 | 18.9884 | < .001 |
| Arm: | | | | |
| alcohol use disorder – control | -0.0277 | 0.16154 | -0.1715 | 0.865 |
| opioid use disorder – control | -0.2006 | 0.15112 | -1.3273 | 0.197 |
| opioid+alcohol use disorder – control | 0.1130 | 0.17870 | 0.6326 | 0.533 |
| Age Cateogry: | | | | |
| 50-64 – 30-49 | 0.0596 | 0.16101 | 0.3703 | 0.714 |
| 50-65 – 30-49 | 0.0163 | 0.24735 | 0.0658 | 0.948 |
| 65+ – 30-49 | 0.1214 | 0.18653 | 0.6509 | 0.521 |
| <30 – 30-49 | 0.1151 | 0.19502 | 0.5903 | 0.561 |
| Ethnicity: | | | | |
| Asian – White | 0.3116 | 0.32474 | 0.9594 | 0.347 |
| Black – White | 0.0850 | 0.13220 | 0.6431 | 0.526 |
| Hispanic – White | 0.3629 | 0.25285 | 1.4354 | 0.164 |
| Gender: | | | | |
| Male – Female | -0.0411 | 0.15627 | -0.2633 | 0.795 |
| PMI converted to hours | -9.43e−4 | 0.00648 | -0.1454 | 0.886 |
| pH | 0.1023 | 0.22416 | 0.4563 | 0.652 |

[a] Represents reference level

### Linear Regression

Model Fit Measures

| Model | R | $R^2$ |
|---|---|---|
| 1 | 0.709 | 0.503 |

Model Coefficients - PLXNA1

| Predictor | Estimate | SE | t | p |
|---|---|---|---|---|
| Intercept [a] | 28.82020 | 1.42988 | 20.156 | < .001 |
| Arm: | | | | |
| alcohol use disorder – control | 0.23889 | 0.15046 | 1.588 | 0.125 |
| opioid use disorder – control | 0.12536 | 0.14075 | 0.891 | 0.382 |
| opioid+alcohol use disorder – control | -0.26111 | 0.16644 | -1.569 | 0.130 |
| Age Cateogry: | | | | |
| 50-64 – 30-49 | 0.02905 | 0.14996 | 0.194 | 0.848 |
| 50-65 – 30-49 | -0.11523 | 0.23038 | -0.500 | 0.622 |
| 65+ – 30-49 | -0.06914 | 0.17374 | -0.398 | 0.694 |
| <30 – 30-49 | 0.27637 | 0.18164 | 1.522 | 0.141 |
| Ethnicity: | | | | |
| Asian – White | 0.11535 | 0.30246 | 0.381 | 0.706 |
| Black – White | -0.08287 | 0.12313 | -0.673 | 0.507 |
| Hispanic – White | -0.36504 | 0.23550 | -1.550 | 0.134 |
| Gender: | | | | |
| Male – Female | -0.40977 | 0.14555 | -2.815 | 0.010 |
| PMI converted to hours | 0.00577 | 0.00604 | 0.955 | 0.349 |
| pH | 0.02674 | 0.20878 | 0.128 | 0.899 |

[a] Represents reference level

### Linear Regression

Model Fit Measures

| Model | R | $R^2$ |
|---|---|---|
| 1 | 0.655 | 0.429 |

Model Coefficients - LAMTOR3

| Predictor | Estimate | SE | t | p |
|---|---|---|---|---|
| Intercept [a] | 20.0117 | 7.8041 | 2.5643 | 0.017 |
| Arm: | | | | |
| alcohol use disorder – control | 0.1717 | 0.8212 | 0.2090 | 0.836 |
| opioid use disorder – control | 1.8896 | 0.7682 | 2.4598 | 0.021 |
| opioid+alcohol use disorder – control | 0.1137 | 0.9084 | 0.1252 | 0.901 |
| Age Cateogry: | | | | |
| 50-64 – 30-49 | 0.8261 | 0.8185 | 1.0094 | 0.323 |
| 50-65 – 30-49 | 0.0135 | 1.2574 | 0.0108 | 0.991 |
| 65+ – 30-49 | 0.9393 | 0.9482 | 0.9906 | 0.332 |
| <30 – 30-49 | 0.4994 | 0.9914 | 0.5037 | 0.619 |
| Ethnicity: | | | | |
| Asian – White | 0.4836 | 1.6508 | 0.2929 | 0.772 |
| Black – White | -0.6301 | 0.6721 | -0.9376 | 0.358 |
| Hispanic – White | 0.1725 | 1.2853 | 0.1342 | 0.894 |
| Gender: | | | | |
| Male – Female | -0.5556 | 0.7944 | -0.6994 | 0.491 |
| PMI converted to hours | -0.0222 | 0.0330 | -0.6724 | 0.508 |
| pH | 0.6503 | 1.1395 | 0.5707 | 0.574 |

[a] Represents reference level

### Linear Regression

Model Fit Measures

| Model | R | $R^2$ |
|---|---|---|
| 1 | 0.598 | 0.358 |

Model Coefficients - ATP8A1

| Predictor | Estimate | SE | t | p |
|---|---|---|---|---|
| Intercept [a] | 27.32856 | 1.55571 | 17.567 | < .001 |
| Arm: | | | | |
| alcohol use disorder – control | 0.08359 | 0.16370 | 0.511 | 0.614 |
| opioid use disorder – control | 0.19888 | 0.15313 | 1.299 | 0.206 |
| opioid+alcohol use disorder – control | -0.02427 | 0.18109 | -0.134 | 0.895 |
| Age Cateogry: | | | | |
| 50-64 – 30-49 | -0.01915 | 0.16316 | -0.117 | 0.908 |
| 50-65 – 30-49 | -0.09424 | 0.25066 | -0.376 | 0.710 |
| 65+ – 30-49 | -0.25432 | 0.18903 | -1.345 | 0.191 |
| <30 – 30-49 | 0.09599 | 0.19763 | 0.486 | 0.632 |
| Ethnicity: | | | | |
| Asian – White | 0.40288 | 0.32908 | 1.224 | 0.233 |
| Black – White | -0.06534 | 0.13397 | -0.488 | 0.630 |
| Hispanic – White | -0.14599 | 0.25622 | -0.570 | 0.574 |
| Gender: | | | | |
| Male – Female | -0.10885 | 0.15836 | -0.687 | 0.498 |
| PMI converted to hours | 0.00930 | 0.00657 | 1.415 | 0.170 |
| pH | 0.15602 | 0.22715 | 0.687 | 0.499 |

[a] Represents reference level

### Linear Regression

Model Fit Measures

| Model | R | $R^2$ |
|---|---|---|
| 1 | 0.732 | 0.536 |

Model Coefficients - HECTD4

| Predictor | Estimate | SE | t | p |
|---|---|---|---|---|
| Intercept [a] | 18.7944 | 3.9106 | 4.8060 | < .001 |
| Arm: | | | | |
| alcohol use disorder – control | -0.5033 | 0.4115 | -1.2230 | 0.233 |
| opioid use disorder – control | -1.1575 | 0.3849 | -3.0070 | 0.006 |
| opioid+alcohol use disorder – control | 0.0200 | 0.4552 | 0.0439 | 0.965 |
| Age Cateogry: | | | | |
| 50-64 – 30-49 | -0.2341 | 0.4101 | -0.5708 | 0.573 |
| 50-65 – 30-49 | 0.8430 | 0.6301 | 1.3379 | 0.193 |
| 65+ – 30-49 | -0.4451 | 0.4752 | -0.9367 | 0.358 |
| <30 – 30-49 | 0.6958 | 0.4968 | 1.4007 | 0.174 |
| Ethnicity: | | | | |
| Asian – White | 0.2856 | 0.8272 | 0.3453 | 0.733 |
| Black – White | 0.0578 | 0.3368 | 0.1717 | 0.865 |
| Hispanic – White | 1.3548 | 0.6441 | 2.1035 | 0.046 |
| Gender: | | | | |
| Male – Female | -0.9393 | 0.3981 | -2.3597 | 0.027 |
| PMI converted to hours | -0.0180 | 0.0165 | -1.0891 | 0.287 |
| pH | 1.0589 | 0.5710 | 1.8545 | 0.076 |

[a] Represents reference level

### Linear Regression

Model Fit Measures

| Model | R | $R^2$ |
|---|---|---|
| 1 | 0.727 | 0.528 |

Model Coefficients - NDUFA2

| Predictor | Estimate | SE | t | p |
|---|---|---|---|---|
| Intercept [a] | 28.53488 | 1.32804 | 21.4865 | < .001 |
| Arm: | | | | |
| alcohol use disorder – control | 0.11320 | 0.13974 | 0.8101 | 0.426 |
| opioid use disorder – control | 0.07849 | 0.13072 | 0.6004 | 0.554 |
| opioid+alcohol use disorder – control | -0.49746 | 0.15459 | -3.2179 | 0.004 |
| Age Cateogry: | | | | |
| 50-64 – 30-49 | 0.01631 | 0.13928 | 0.1171 | 0.908 |
| 50-65 – 30-49 | 0.11659 | 0.21397 | 0.5449 | 0.591 |
| 65+ – 30-49 | -0.01492 | 0.16136 | -0.0925 | 0.927 |
| <30 – 30-49 | -0.01134 | 0.16871 | -0.0672 | 0.947 |
| Ethnicity: | | | | |
| Asian – White | -0.20005 | 0.28092 | -0.7121 | 0.483 |
| Black – White | -0.17627 | 0.11436 | -1.5413 | 0.136 |
| Hispanic – White | -0.22740 | 0.21873 | -1.0397 | 0.309 |
| Gender: | | | | |
| Male – Female | -0.10814 | 0.13518 | -0.8000 | 0.432 |
| PMI converted to hours | 0.00863 | 0.00561 | 1.5381 | 0.137 |
| pH | 0.04638 | 0.19391 | 0.2392 | 0.813 |

[a] Represents reference level

### Linear Regression

Model Fit Measures

| Model | R | $R^2$ |
|---|---|---|
| 1 | 0.684 | 0.467 |

Model Coefficients - TPPP

| Predictor | Estimate | SE | t | p |
|---|---|---|---|---|
| Intercept [a] | 31.2020 | 1.73894 | 17.9431 | < .001 |
| Arm: | | | | |
| alcohol use disorder – control | 0.1352 | 0.18298 | 0.7389 | 0.467 |
| opioid use disorder – control | 0.2954 | 0.17117 | 1.7258 | 0.097 |
| opioid+alcohol use disorder – control | -0.3743 | 0.20242 | -1.8493 | 0.077 |
| Age Cateogry: | | | | |
| 50-64 – 30-49 | -0.0200 | 0.18238 | -0.1098 | 0.913 |
| 50-65 – 30-49 | 0.0307 | 0.28018 | 0.1095 | 0.914 |
| 65+ – 30-49 | 0.0288 | 0.21129 | 0.1362 | 0.893 |
| <30 – 30-49 | -0.1592 | 0.22091 | -0.7208 | 0.478 |
| Ethnicity: | | | | |
| Asian – White | -0.1402 | 0.36784 | -0.3812 | 0.706 |
| Black – White | -0.0148 | 0.14975 | -0.0988 | 0.922 |
| Hispanic – White | -0.1145 | 0.28640 | -0.3997 | 0.693 |
| Gender: | | | | |
| Male – Female | 0.0926 | 0.17701 | 0.5233 | 0.606 |
| PMI converted to hours | 0.0121 | 0.00734 | 1.6410 | 0.114 |
| pH | 0.1069 | 0.25391 | 0.4211 | 0.677 |

[a] Represents reference level

### Linear Regression

Model Fit Measures

| Model | R | $R^2$ |
|---|---|---|
| 1 | 0.740 | 0.547 |

Model Coefficients - NDUFB6

| Predictor | Estimate | SE | t | p |
|---|---|---|---|---|
| Intercept [a] | 28.74127 | 1.60683 | 17.8870 | < .001 |
| Arm: | | | | |
| alcohol use disorder – control | -0.03286 | 0.16908 | -0.1944 | 0.848 |
| opioid use disorder – control | 0.01406 | 0.15817 | 0.0889 | 0.930 |
| opioid+alcohol use disorder – control | -0.65415 | 0.18704 | -3.4973 | 0.002 |
| Age Cateogry: | | | | |
| 50-64 – 30-49 | -0.14705 | 0.16852 | -0.8726 | 0.392 |
| 50-65 – 30-49 | -0.02196 | 0.25889 | -0.0848 | 0.933 |
| 65+ – 30-49 | -0.03944 | 0.19524 | -0.2020 | 0.842 |
| <30 – 30-49 | -0.03551 | 0.20412 | -0.1740 | 0.863 |
| Ethnicity: | | | | |
| Asian – White | -0.80741 | 0.33989 | -2.3755 | 0.026 |
| Black – White | -0.12642 | 0.13837 | -0.9136 | 0.370 |
| Hispanic – White | -0.27371 | 0.26464 | -1.0342 | 0.311 |
| Gender: | | | | |
| Male – Female | -0.14886 | 0.16356 | -0.9101 | 0.372 |
| PMI converted to hours | 0.00517 | 0.00679 | 0.7619 | 0.454 |
| pH | -0.01176 | 0.23462 | -0.0501 | 0.960 |

[a] Represents reference level

### Linear Regression

Model Fit Measures

| Model | R | $R^2$ |
|---|---|---|
| 1 | 0.694 | 0.482 |

Model Coefficients - KCNQ2

| Predictor | Estimate | SE | t | p |
|---|---|---|---|---|
| Intercept [a] | 21.0771 | 2.5635 | 8.2222 | < .001 |
| Arm: | | | | |
| alcohol use disorder – control | -0.2562 | 0.2697 | -0.9496 | 0.352 |
| opioid use disorder – control | 0.4209 | 0.2523 | 1.6680 | 0.108 |
| opioid+alcohol use disorder – control | -0.6152 | 0.2984 | -2.0615 | 0.050 |
| Age Cateogry: | | | | |
| 50-64 – 30-49 | -0.0199 | 0.2688 | -0.0739 | 0.942 |
| 50-65 – 30-49 | 0.2287 | 0.4130 | 0.5536 | 0.585 |
| 65+ – 30-49 | 0.1124 | 0.3115 | 0.3610 | 0.721 |
| <30 – 30-49 | -0.4122 | 0.3256 | -1.2659 | 0.218 |
| Ethnicity: | | | | |
| Asian – White | 0.1982 | 0.5422 | 0.3654 | 0.718 |
| Black – White | 0.0288 | 0.2208 | 0.1304 | 0.897 |
| Hispanic – White | -0.1353 | 0.4222 | -0.3206 | 0.751 |
| Gender: | | | | |
| Male – Female | 0.1896 | 0.2609 | 0.7265 | 0.475 |
| PMI converted to hours | 0.0123 | 0.0108 | 1.1402 | 0.265 |
| pH | 0.3026 | 0.3743 | 0.8084 | 0.427 |

[a] Represents reference level

### Linear Regression

Model Fit Measures

| Model | R | $R^2$ |
|---|---|---|
| 1 | 0.655 | 0.429 |

Model Coefficients - SPAG9

| Predictor | Estimate | SE | t | p |
|---|---|---|---|---|
| Intercept [a] | 27.51010 | 3.0828 | 8.9238 | < .001 |
| Arm: | | | | |
| alcohol use disorder – control | -0.16696 | 0.3244 | -0.5147 | 0.611 |
| opioid use disorder – control | -0.01919 | 0.3034 | -0.0633 | 0.950 |
| opioid+alcohol use disorder – control | -1.05523 | 0.3588 | -2.9406 | 0.007 |
| Age Cateogry: | | | | |
| 50-64 – 30-49 | 0.18381 | 0.3233 | 0.5685 | 0.575 |
| 50-65 – 30-49 | 0.30340 | 0.4967 | 0.6108 | 0.547 |
| 65+ – 30-49 | 0.15847 | 0.3746 | 0.4231 | 0.676 |
| <30 – 30-49 | 0.30341 | 0.3916 | 0.7748 | 0.446 |
| Ethnicity: | | | | |
| Asian – White | -0.35232 | 0.6521 | -0.5403 | 0.594 |
| Black – White | 0.09907 | 0.2655 | 0.3732 | 0.712 |
| Hispanic – White | 0.49217 | 0.5077 | 0.9694 | 0.342 |
| Gender: | | | | |
| Male – Female | -0.35159 | 0.3138 | -1.1204 | 0.274 |
| PMI converted to hours | -0.00901 | 0.0130 | -0.6920 | 0.496 |
| pH | -0.50712 | 0.4501 | -1.1266 | 0.271 |

[a] Represents reference level

### Linear Regression

Model Fit Measures

| Model | R | $R^2$ |
|---|---|---|
| 1 | 0.794 | 0.630 |

Model Coefficients - HCN1

| Predictor | Estimate | SE | t | p |
|---|---|---|---|---|
| Intercept [a] | 26.9944 | 3.4816 | 7.7535 | < .001 |
| Arm: | | | | |
| alcohol use disorder – control | 0.1160 | 0.3663 | 0.3165 | 0.754 |
| opioid use disorder – control | -0.5038 | 0.3427 | -1.4699 | 0.155 |
| opioid+alcohol use disorder – control | -1.5159 | 0.4053 | -3.7404 | 0.001 |
| Age Cateogry: | | | | |
| 50-64 – 30-49 | -0.5796 | 0.3651 | -1.5873 | 0.126 |
| 50-65 – 30-49 | -0.5869 | 0.5609 | -1.0463 | 0.306 |
| 65+ – 30-49 | -0.7145 | 0.4230 | -1.6890 | 0.104 |
| <30 – 30-49 | 0.0181 | 0.4423 | 0.0410 | 0.968 |
| Ethnicity: | | | | |
| Asian – White | -2.5190 | 0.7364 | -3.4205 | 0.002 |
| Black – White | -0.1872 | 0.2998 | -0.6243 | 0.538 |
| Hispanic – White | -0.1001 | 0.5734 | -0.1747 | 0.863 |
| Gender: | | | | |
| Male – Female | -0.1088 | 0.3544 | -0.3071 | 0.761 |
| PMI converted to hours | -0.0196 | 0.0147 | -1.3324 | 0.195 |
| pH | -0.2255 | 0.5083 | -0.4436 | 0.661 |

[a] Represents reference level

### Linear Regression

Model Fit Measures

| Model | R | $R^2$ |
|---|---|---|
| 1 | 0.710 | 0.504 |

Model Coefficients - NDUFS3

| Predictor | Estimate | SE | t | p |
|---|---|---|---|---|
| Intercept [a] | 29.9569 | 0.67089 | 44.6528 | < .001 |
| Arm: | | | | |
| alcohol use disorder – control | 0.0173 | 0.07059 | 0.2449 | 0.809 |
| opioid use disorder – control | -0.0357 | 0.06604 | -0.5402 | 0.594 |
| opioid+alcohol use disorder – control | -0.2495 | 0.07809 | -3.1949 | 0.004 |
| Age Cateogry: | | | | |
| 50-64 – 30-49 | -0.0524 | 0.07036 | -0.7452 | 0.463 |
| 50-65 – 30-49 | 0.1444 | 0.10809 | 1.3357 | 0.194 |
| 65+ – 30-49 | -0.0221 | 0.08152 | -0.2711 | 0.789 |
| <30 – 30-49 | 0.0144 | 0.08523 | 0.1686 | 0.868 |
| Ethnicity: | | | | |
| Asian – White | -0.1152 | 0.14191 | -0.8118 | 0.425 |
| Black – White | -0.0426 | 0.05777 | -0.7377 | 0.468 |
| Hispanic – White | 0.0594 | 0.11049 | 0.5378 | 0.596 |
| Gender: | | | | |
| Male – Female | -0.0819 | 0.06829 | -1.1995 | 0.242 |
| PMI converted to hours | 1.25e-4 | 0.00283 | 0.0440 | 0.965 |
| pH | 0.0526 | 0.09796 | 0.5369 | 0.596 |

[a] Represents reference level

### Linear Regression

Model Fit Measures

| Model | R | $R^2$ |
|---|---|---|
| 1 | 0.680 | 0.463 |

Model Coefficients - LGI1

| Predictor | Estimate | SE | t | p |
|---|---|---|---|---|
| Intercept [a] | 25.8941 | 3.5595 | 7.275 | < .001 |
| Arm: | | | | |
| alcohol use disorder – control | 0.2414 | 0.3745 | 0.644 | 0.525 |
| opioid use disorder – control | 0.1598 | 0.3504 | 0.456 | 0.652 |
| opioid+alcohol use disorder – control | -1.2371 | 0.4143 | -2.986 | 0.006 |
| Age Cateogry: | | | | |
| 50-64 – 30-49 | 0.3276 | 0.3733 | 0.877 | 0.389 |
| 50-65 – 30-49 | 0.2617 | 0.5735 | 0.456 | 0.652 |
| 65+ – 30-49 | 0.1885 | 0.4325 | 0.436 | 0.667 |
| <30 – 30-49 | 0.4528 | 0.4522 | 1.001 | 0.327 |
| Ethnicity: | | | | |
| Asian – White | 0.1627 | 0.7529 | 0.216 | 0.831 |
| Black – White | -0.1576 | 0.3065 | -0.514 | 0.612 |
| Hispanic – White | -0.3203 | 0.5862 | -0.546 | 0.590 |
| Gender: | | | | |
| Male – Female | -0.4414 | 0.3623 | -1.218 | 0.235 |
| PMI converted to hours | 0.0167 | 0.0150 | 1.112 | 0.277 |
| pH | 0.1739 | 0.5197 | 0.335 | 0.741 |

[a] Represents reference level

### Linear Regression

Model Fit Measures

| Model | R | $R^2$ |
|---|---|---|
| 1 | 0.719 | 0.518 |

Model Coefficients - C1QB

| Predictor | Estimate | SE | t | p |
|---|---|---|---|---|
| Intercept [a] | 23.52397 | 2.6885 | 8.74988 | < .001 |
| Arm: | | | | |
| alcohol use disorder – control | -0.16396 | 0.2829 | -0.57957 | 0.568 |
| opioid use disorder – control | 0.20552 | 0.2646 | 0.77659 | 0.445 |
| opioid+alcohol use disorder – control | -1.01829 | 0.3130 | -3.25383 | 0.003 |
| Age Cateogry: | | | | |
| 50-64 – 30-49 | 0.04932 | 0.2820 | 0.17492 | 0.863 |
| 50-65 – 30-49 | 0.38909 | 0.4332 | 0.89825 | 0.378 |
| 65+ – 30-49 | 0.41246 | 0.3267 | 1.26266 | 0.219 |
| <30 – 30-49 | -0.02027 | 0.3415 | -0.05936 | 0.953 |
| Ethnicity: | | | | |
| Asian – White | 0.00462 | 0.5687 | 0.00812 | 0.994 |
| Black – White | -0.08507 | 0.2315 | -0.36743 | 0.717 |
| Hispanic – White | -0.23762 | 0.4428 | -0.53665 | 0.596 |
| Gender: | | | | |
| Male – Female | -0.08682 | 0.2737 | -0.31726 | 0.754 |
| PMI converted to hours | -0.00582 | 0.0114 | -0.51266 | 0.613 |
| pH | 0.03925 | 0.3926 | 0.09999 | 0.921 |

[a] Represents reference level

### Linear Regression

Model Fit Measures

| Model | R | $R^2$ |
|---|---|---|
| 1 | 0.837 | 0.700 |

Model Coefficients - EIF3F

| Predictor | Estimate | SE | t | p |
|---|---|---|---|---|
| Intercept [a] | 19.0588 | 2.4706 | 7.714 | < .001 |
| Arm: | | | | |
| alcohol use disorder – control | -0.9415 | 0.2600 | -3.622 | 0.001 |
| opioid use disorder – control | -0.2307 | 0.2432 | -0.949 | 0.352 |
| opioid+alcohol use disorder – control | 0.8936 | 0.2876 | 3.107 | 0.005 |
| Age Cateogry: | | | | |
| 50-64 – 30-49 | -0.6724 | 0.2591 | -2.595 | 0.016 |
| 50-65 – 30-49 | -0.4332 | 0.3981 | -1.088 | 0.287 |
| 65+ – 30-49 | -0.6274 | 0.3002 | -2.090 | 0.047 |
| <30 – 30-49 | -0.7656 | 0.3139 | -2.439 | 0.022 |
| Ethnicity: | | | | |
| Asian – White | -0.0785 | 0.5226 | -0.150 | 0.882 |
| Black – White | 0.1350 | 0.2128 | 0.635 | 0.532 |
| Hispanic – White | -0.7913 | 0.4069 | -1.945 | 0.064 |
| Gender: | | | | |
| Male – Female | 0.2676 | 0.2515 | 1.064 | 0.298 |
| PMI converted to hours | 0.0355 | 0.0104 | 3.406 | 0.002 |
| pH | 0.6879 | 0.3607 | 1.907 | 0.069 |

[a] Represents reference level

### Linear Regression

Model Fit Measures

| Model | R | $R^2$ |
|---|---|---|
| 1 | 0.628 | 0.395 |

Model Coefficients - SSC5D

| Predictor | Estimate | SE | t | p |
|---|---|---|---|---|
| Intercept [a] | 22.54687 | 3.6921 | 6.107 | < .001 |
| Arm: | | | | |
| alcohol use disorder – control | 0.23293 | 0.3885 | 0.600 | 0.554 |
| opioid use disorder – control | 0.54261 | 0.3634 | 1.493 | 0.148 |
| opioid+alcohol use disorder – control | -0.59452 | 0.4298 | -1.383 | 0.179 |
| Age Cateogry: | | | | |
| 50-64 – 30-49 | 0.22761 | 0.3872 | 0.588 | 0.562 |
| 50-65 – 30-49 | 0.22023 | 0.5949 | 0.370 | 0.714 |
| 65+ – 30-49 | 0.93446 | 0.4486 | 2.083 | 0.048 |
| <30 – 30-49 | 0.09470 | 0.4690 | 0.202 | 0.842 |
| Ethnicity: | | | | |
| Asian – White | 0.45432 | 0.7810 | 0.582 | 0.566 |
| Black – White | 0.37119 | 0.3179 | 1.167 | 0.254 |
| Hispanic – White | 0.23442 | 0.6081 | 0.386 | 0.703 |
| Gender: | | | | |
| Male – Female | 0.18452 | 0.3758 | 0.491 | 0.628 |
| PMI converted to hours | -0.00433 | 0.0156 | -0.277 | 0.784 |
| pH | 0.07968 | 0.5391 | 0.148 | 0.884 |

[a] Represents reference level

### Linear Regression

Model Fit Measures

| Model | R | $R^2$ |
|---|---|---|
| 1 | 0.668 | 0.447 |

Model Coefficients - MROH6

| Predictor | Estimate | SE | t | p |
|---|---|---|---|---|
| Intercept [a] | 23.92039 | 2.8415 | 8.4181 | < .001 |
| Arm: | | | | |
| alcohol use disorder – control | -0.17015 | 0.2990 | -0.5690 | 0.575 |
| opioid use disorder – control | 0.01747 | 0.2797 | 0.0624 | 0.951 |
| opioid+alcohol use disorder – control | -1.06391 | 0.3308 | -3.2165 | 0.004 |
| Age Cateogry: | | | | |
| 50-64 – 30-49 | 0.19112 | 0.2980 | 0.6413 | 0.527 |
| 50-65 – 30-49 | 0.12295 | 0.4578 | 0.2685 | 0.791 |
| 65+ – 30-49 | 0.25537 | 0.3453 | 0.7396 | 0.467 |
| <30 – 30-49 | 0.29438 | 0.3610 | 0.8155 | 0.423 |
| Ethnicity: | | | | |
| Asian – White | -0.58946 | 0.6011 | -0.9807 | 0.337 |
| Black – White | -0.07905 | 0.2447 | -0.3231 | 0.749 |
| Hispanic – White | -0.17571 | 0.4680 | -0.3755 | 0.711 |
| Gender: | | | | |
| Male – Female | -0.36746 | 0.2892 | -1.2704 | 0.216 |
| PMI converted to hours | 0.00714 | 0.0120 | 0.5945 | 0.558 |
| pH | -0.04653 | 0.4149 | -0.1122 | 0.912 |

[a] Represents reference level

### Linear Regression

Model Fit Measures

| Model | R | $R^2$ |
|---|---|---|
| 1 | 0.553 | 0.306 |

Model Coefficients - ARPC2

| Predictor | Estimate | SE | t | p |
|---|---|---|---|---|
| Intercept [a] | 30.7587 | 6.0169 | 5.1120 | < .001 |
| Arm: | | | | |
| alcohol use disorder – control | 0.7438 | 0.6331 | 1.1748 | 0.252 |
| opioid use disorder – control | 0.3501 | 0.5923 | 0.5911 | 0.560 |
| opioid+alcohol use disorder – control | 1.5585 | 0.7004 | 2.2252 | 0.036 |
| Age Cateogry: | | | | |
| 50-64 – 30-49 | -0.0421 | 0.6310 | -0.0666 | 0.947 |
| 50-65 – 30-49 | 0.7399 | 0.9694 | 0.7632 | 0.453 |
| 65+ – 30-49 | -0.0931 | 0.7311 | -0.1273 | 0.900 |
| <30 – 30-49 | 0.4179 | 0.7644 | 0.5467 | 0.590 |
| Ethnicity: | | | | |
| Asian – White | 1.3760 | 1.2727 | 1.0811 | 0.290 |
| Black – White | 0.7834 | 0.5181 | 1.5119 | 0.144 |
| Hispanic – White | 0.7706 | 0.9910 | 0.7776 | 0.444 |
| Gender: | | | | |
| Male – Female | -0.0937 | 0.6125 | -0.1531 | 0.880 |
| PMI converted to hours | -0.0158 | 0.0254 | -0.6210 | 0.540 |
| pH | -0.4358 | 0.8785 | -0.4960 | 0.624 |

[a] Represents reference level

### Linear Regression

Model Fit Measures

| Model | R | $R^2$ |
|---|---|---|
| 1 | 0.693 | 0.480 |

Model Coefficients - MPDU1

| Predictor | Estimate | SE | t | p |
|---|---|---|---|---|
| Intercept [a] | 24.82621 | 2.5751 | 9.6409 | < .001 |
| Arm: | | | | |
| alcohol use disorder – control | -0.14928 | 0.2710 | -0.5509 | 0.587 |
| opioid use disorder – control | -0.25660 | 0.2535 | -1.0123 | 0.321 |
| opioid+alcohol use disorder – control | -1.03714 | 0.2998 | -3.4600 | 0.002 |
| Age Cateogry: | | | | |
| 50-64 – 30-49 | -0.11931 | 0.2701 | -0.4418 | 0.663 |
| 50-65 – 30-49 | -0.87145 | 0.4149 | -2.1004 | 0.046 |
| 65+ – 30-49 | 0.01894 | 0.3129 | 0.0605 | 0.952 |
| <30 – 30-49 | -0.20675 | 0.3271 | -0.6320 | 0.533 |
| Ethnicity: | | | | |
| Asian – White | -0.60709 | 0.5447 | -1.1145 | 0.276 |
| Black – White | -0.17577 | 0.2218 | -0.7926 | 0.436 |
| Hispanic – White | 0.03752 | 0.4241 | 0.0885 | 0.930 |
| Gender: | | | | |
| Male – Female | 0.07693 | 0.2621 | 0.2935 | 0.772 |
| PMI converted to hours | -0.00262 | 0.0109 | -0.2407 | 0.812 |
| pH | -0.10930 | 0.3760 | -0.2907 | 0.774 |

[a] Represents reference level

### Results

#### Linear Regression

Model Fit Measures

| Model | R | $R^2$ |
|---|---|---|
| 1 | 0.649 | 0.421 |

Model Coefficients - ITGAM

| Predictor | Estimate | SE | t | p |
|---|---|---|---|---|
| Intercept [a] | 23.3984 | 2.7414 | 8.5352 | < .001 |
| Arm: | | | | |
| alcohol use disorder – control | -0.3628 | 0.2885 | -1.2578 | 0.221 |
| opioid use disorder – control | -0.0706 | 0.2698 | -0.2616 | 0.796 |
| opioid+alcohol use disorder – control | -0.9287 | 0.3191 | -2.9101 | 0.008 |
| Age Cateogry: | | | | |
| 50-64 – 30-49 | 0.0696 | 0.2875 | 0.2421 | 0.811 |
| 50-65 – 30-49 | -0.4084 | 0.4417 | -0.9246 | 0.364 |
| 65+ – 30-49 | 0.2363 | 0.3331 | 0.7095 | 0.485 |
| <30 – 30-49 | -0.2040 | 0.3483 | -0.5859 | 0.563 |
| Ethnicity: | | | | |
| Asian – White | 0.0350 | 0.5799 | 0.0604 | 0.952 |
| Black – White | -0.1895 | 0.2361 | -0.8028 | 0.430 |
| Hispanic – White | -0.4515 | 0.4515 | -1.0001 | 0.327 |
| Gender: | | | | |
| Male – Female | -0.0832 | 0.2790 | -0.2982 | 0.768 |
| PMI converted to hours | 0.0202 | 0.0116 | 1.7485 | 0.093 |
| pH | -0.0390 | 0.4003 | -0.0974 | 0.923 |

[a] Represents reference level

#### Linear Regression

Model Fit Measures

| Model | R | $R^2$ |
|---|---|---|
| 1 | 0.631 | 0.398 |

Model Coefficients - SNRPA

| Predictor | Estimate | SE | t | p |
|---|---|---|---|---|
| Intercept [a] | 19.9303 | 3.1443 | 6.3385 | < .001 |
| Arm: | | | | |
| alcohol use disorder – control | -0.0600 | 0.3309 | -0.1813 | 0.858 |
| opioid use disorder – control | -0.2372 | 0.3095 | -0.7662 | 0.451 |
| opioid+alcohol use disorder – control | -0.8828 | 0.3660 | -2.4120 | 0.024 |
| Age Cateogry: | | | | |
| 50-64 – 30-49 | -0.3017 | 0.3298 | -0.9150 | 0.369 |
| 50-65 – 30-49 | 0.5442 | 0.5066 | 1.0742 | 0.293 |
| 65+ – 30-49 | -0.0406 | 0.3820 | -0.1062 | 0.916 |
| <30 – 30-49 | 0.1140 | 0.3994 | 0.2853 | 0.778 |
| Ethnicity: | | | | |
| Asian – White | -0.2929 | 0.6651 | -0.4403 | 0.664 |
| Black – White | -0.1697 | 0.2708 | -0.6268 | 0.537 |
| Hispanic – White | -0.0576 | 0.5179 | -0.1113 | 0.912 |
| Gender: | | | | |
| Male – Female | -0.1393 | 0.3201 | -0.4352 | 0.667 |
| PMI converted to hours | 6.52e-4 | 0.0133 | 0.0491 | 0.961 |
| pH | 0.6051 | 0.4591 | 1.3179 | 0.200 |

[a] Represents reference level

### Linear Regression

Model Fit Measures

| Model | R | $R^2$ |
|---|---|---|
| 1 | 0.695 | 0.483 |

Model Coefficients - UQCRB (2)

| Predictor | Estimate | SE | t | p |
|---|---|---|---|---|
| Intercept [a] | 30.93873 | 1.27223 | 24.3185 | < .001 |
| Arm: | | | | |
| alcohol use disorder – control | 0.05525 | 0.13387 | 0.4127 | 0.684 |
| opioid use disorder – control | 0.19738 | 0.12523 | 1.5762 | 0.128 |
| opioid+alcohol use disorder – control | -0.34357 | 0.14809 | -2.3199 | 0.029 |
| Age Cateogry: | | | | |
| 50-64 – 30-49 | 0.05203 | 0.13343 | 0.3900 | 0.700 |
| 50-65 – 30-49 | 0.16856 | 0.20498 | 0.8223 | 0.419 |
| 65+ – 30-49 | 0.00412 | 0.15458 | 0.0267 | 0.979 |
| <30 – 30-49 | 0.03187 | 0.16162 | 0.1972 | 0.845 |
| Ethnicity: | | | | |
| Asian – White | -0.14057 | 0.26911 | -0.5223 | 0.606 |
| Black – White | -0.03577 | 0.10956 | -0.3265 | 0.747 |
| Hispanic – White | -0.29533 | 0.20954 | -1.4095 | 0.172 |
| Gender: | | | | |
| Male – Female | -0.12760 | 0.12950 | -0.9853 | 0.334 |
| PMI converted to hours | 0.00129 | 0.00537 | 0.2394 | 0.813 |
| pH | -0.07427 | 0.18576 | -0.3998 | 0.693 |

[a] Represents reference level

### Linear Regression

Model Fit Measures

| Model | R | $R^2$ |
|---|---|---|
| 1 | 0.855 | 0.731 |

Model Coefficients - RPL3 (2)

| Predictor | Estimate | SE | t | p |
|---|---|---|---|---|
| Intercept [a] | 28.66087 | 0.88750 | 32.2938 | < .001 |
| Arm: | | | | |
| alcohol use disorder – control | 0.10251 | 0.09339 | 1.0976 | 0.283 |
| opioid use disorder – control | 0.07871 | 0.08736 | 0.9010 | 0.377 |
| opioid+alcohol use disorder – control | -0.46002 | 0.10331 | -4.4528 | < .001 |
| Age Cateogry: | | | | |
| 50-64 – 30-49 | -0.11793 | 0.09308 | -1.2670 | 0.217 |
| 50-65 – 30-49 | -0.18514 | 0.14299 | -1.2947 | 0.208 |
| 65+ – 30-49 | -0.00552 | 0.10784 | -0.0512 | 0.960 |
| <30 – 30-49 | 0.07549 | 0.11274 | 0.6696 | 0.510 |
| Ethnicity: | | | | |
| Asian – White | -0.15188 | 0.18773 | -0.8090 | 0.426 |
| Black – White | -0.04286 | 0.07643 | -0.5608 | 0.580 |
| Hispanic – White | -0.41842 | 0.14617 | -2.8625 | 0.009 |
| Gender: | | | | |
| Male – Female | -0.26125 | 0.09034 | -2.8919 | 0.008 |
| PMI converted to hours | 0.00613 | 0.00375 | 1.6361 | 0.115 |
| pH | -0.17734 | 0.12959 | -1.3685 | 0.184 |

[a] Represents reference level

### Linear Regression

Model Fit Measures

| Model | R | $R^2$ |
|---|---|---|
| 1 | 0.662 | 0.438 |

Model Coefficients - PCP4

| Predictor | Estimate | SE | t | p |
| --- | --- | --- | --- | --- |
| Intercept [a] | 21.3349 | 5.2383 | 4.073 | < .001 |
| Arm: | | | | |
| alcohol use disorder – control | -0.2375 | 0.5512 | -0.431 | 0.670 |
| opioid use disorder – control | -0.6526 | 0.5156 | -1.266 | 0.218 |
| opioid+alcohol use disorder – control | -1.1678 | 0.6098 | -1.915 | 0.067 |
| Age Cateogry: | | | | |
| 50-64 – 30-49 | -0.7231 | 0.5494 | -1.316 | 0.201 |
| 50-65 – 30-49 | -0.1003 | 0.8440 | -0.119 | 0.906 |
| 65+ – 30-49 | -0.3467 | 0.6365 | -0.545 | 0.591 |
| <30 – 30-49 | -0.1360 | 0.6654 | -0.204 | 0.840 |
| Ethnicity: | | | | |
| Asian – White | -1.7645 | 1.1080 | -1.592 | 0.124 |
| Black – White | 0.1352 | 0.4511 | 0.300 | 0.767 |
| Hispanic – White | 1.6678 | 0.8627 | 1.933 | 0.065 |
| Gender: | | | | |
| Male – Female | 0.2313 | 0.5332 | 0.434 | 0.668 |
| PMI converted to hours | -0.0243 | 0.0221 | -1.098 | 0.283 |
| pH | 0.5620 | 0.7648 | 0.735 | 0.470 |

[a] Represents reference level

### Linear Regression

Model Fit Measures

| Model | R | $R^2$ |
| --- | --- | --- |
| 1 | 0.718 | 0.515 |

Model Coefficients - HSPA9

| Predictor | Estimate | SE | t | p |
|---|---|---|---|---|
| Intercept [a] | 31.18176 | 0.94599 | 32.9619 | < .001 |
| Arm: | | | | |
| alcohol use disorder – control | 0.01309 | 0.09954 | 0.1315 | 0.896 |
| opioid use disorder – control | 0.14188 | 0.09312 | 1.5237 | 0.141 |
| opioid+alcohol use disorder – control | -0.27699 | 0.11012 | -2.5154 | 0.019 |
| Age Cateogry: | | | | |
| 50-64 – 30-49 | -0.04830 | 0.09921 | -0.4868 | 0.631 |
| 50-65 – 30-49 | -0.04691 | 0.15242 | -0.3078 | 0.761 |
| 65+ – 30-49 | 0.04629 | 0.11494 | 0.4027 | 0.691 |
| <30 – 30-49 | -0.09000 | 0.12017 | -0.7489 | 0.461 |
| Ethnicity: | | | | |
| Asian – White | -0.10450 | 0.20010 | -0.5222 | 0.606 |
| Black – White | -0.13694 | 0.08146 | -1.6810 | 0.106 |
| Hispanic – White | -0.10385 | 0.15580 | -0.6666 | 0.511 |
| Gender: | | | | |
| Male – Female | -0.00916 | 0.09629 | -0.0952 | 0.925 |
| PMI converted to hours | 0.00435 | 0.00400 | 1.0889 | 0.287 |
| pH | 0.08582 | 0.13813 | 0.6213 | 0.540 |

[a] Represents reference level

### Linear Regression

Model Fit Measures

| Model | R | $R^2$ |
|---|---|---|
| 1 | 0.797 | 0.635 |

Model Coefficients - EIF1

| Predictor | Estimate | SE | t | p |
|---|---|---|---|---|
| Intercept [a] | 15.47867 | 3.3724 | 4.5898 | < .001 |
| Arm: | | | | |
| alcohol use disorder – control | -0.40422 | 0.3549 | -1.1391 | 0.266 |
| opioid use disorder – control | -0.92992 | 0.3320 | -2.8013 | 0.010 |
| opioid+alcohol use disorder – control | -1.37152 | 0.3926 | -3.4937 | 0.002 |
| Age Cateogry: | | | | |
| 50-64 – 30-49 | -0.65985 | 0.3537 | -1.8656 | 0.074 |
| 50-65 – 30-49 | 0.06076 | 0.5434 | 0.1118 | 0.912 |
| 65+ – 30-49 | 0.16691 | 0.4098 | 0.4073 | 0.687 |
| <30 – 30-49 | 0.01365 | 0.4284 | 0.0319 | 0.975 |
| Ethnicity: | | | | |
| Asian – White | -0.55169 | 0.7134 | -0.7734 | 0.447 |
| Black – White | 0.27738 | 0.2904 | 0.9551 | 0.349 |
| Hispanic – White | 0.40747 | 0.5554 | 0.7336 | 0.470 |
| Gender: | | | | |
| Male – Female | -0.29892 | 0.3433 | -0.8708 | 0.393 |
| PMI converted to hours | -0.00876 | 0.0142 | -0.6149 | 0.544 |
| pH | 1.42411 | 0.4924 | 2.8921 | 0.008 |

[a] Represents reference level

### Linear Regression

Model Fit Measures

| Model | R | $R^2$ |
|---|---|---|
| 1 | 0.734 | 0.539 |

Model Coefficients - SHMT2 (2)

| Predictor | Estimate | SE | t | p |
|---|---|---|---|---|
| Intercept [a] | 24.7655 | 4.6613 | 5.313 | < .001 |
| Arm: | | | | |
| alcohol use disorder – control | -0.4564 | 0.4905 | -0.931 | 0.361 |
| opioid use disorder – control | -1.3707 | 0.4588 | -2.987 | 0.006 |
| opioid+alcohol use disorder – control | -1.0567 | 0.5426 | -1.948 | 0.063 |
| Age Cateogry: | | | | |
| 50-64 – 30-49 | 0.2770 | 0.4889 | 0.567 | 0.576 |
| 50-65 – 30-49 | -1.0108 | 0.7510 | -1.346 | 0.191 |
| 65+ – 30-49 | 0.6002 | 0.5664 | 1.060 | 0.300 |
| <30 – 30-49 | 0.1934 | 0.5921 | 0.327 | 0.747 |
| Ethnicity: | | | | |
| Asian – White | 0.9478 | 0.9860 | 0.961 | 0.346 |
| Black – White | 0.3825 | 0.4014 | 0.953 | 0.350 |
| Hispanic – White | 1.2322 | 0.7677 | 1.605 | 0.122 |
| Gender: | | | | |
| Male – Female | 0.0555 | 0.4745 | 0.117 | 0.908 |
| PMI converted to hours | -0.0229 | 0.0197 | -1.164 | 0.256 |
| pH | 0.1018 | 0.6806 | 0.150 | 0.882 |

[a] Represents reference level

### Linear Regression

Model Fit Measures

| Model | R | $R^2$ |
|---|---|---|
| 1 | 0.619 | 0.383 |

Model Coefficients - TXLNA

| Predictor | Estimate | SE | t | p |
|---|---|---|---|---|
| Intercept [a] | 21.55210 | 2.8827 | 7.4763 | < .001 |
| Arm: | | | | |
| alcohol use disorder – control | -0.39401 | 0.3033 | -1.2989 | 0.206 |
| opioid use disorder – control | 0.04510 | 0.2838 | 0.1589 | 0.875 |
| opioid+alcohol use disorder – control | -0.84654 | 0.3356 | -2.5228 | 0.019 |
| Age Cateogry: | | | | |
| 50-64 – 30-49 | -0.04258 | 0.3023 | -0.1408 | 0.889 |
| 50-65 – 30-49 | 0.13697 | 0.4645 | 0.2949 | 0.771 |
| 65+ – 30-49 | -0.02557 | 0.3503 | -0.0730 | 0.942 |
| <30 – 30-49 | -0.17373 | 0.3662 | -0.4744 | 0.639 |
| Ethnicity: | | | | |
| Asian – White | -0.29099 | 0.6098 | -0.4772 | 0.638 |
| Black – White | -0.35682 | 0.2482 | -1.4374 | 0.164 |
| Hispanic – White | -0.11855 | 0.4748 | -0.2497 | 0.805 |
| Gender: | | | | |
| Male – Female | -0.07801 | 0.2934 | -0.2659 | 0.793 |
| PMI converted to hours | 0.00134 | 0.0122 | 0.1103 | 0.913 |
| pH | 0.34745 | 0.4209 | 0.8255 | 0.417 |

[a] Represents reference level

### Linear Regression

Model Fit Measures

| Model | R | $R^2$ |
|---|---|---|
| 1 | 0.659 | 0.434 |

Model Coefficients - PRPH

| Predictor | Estimate | SE | t | p |
|---|---|---|---|---|
| Intercept [a] | 24.16838 | 2.7752 | 8.7088 | < .001 |
| Arm: | | | | |
| alcohol use disorder – control | -0.26823 | 0.2920 | -0.9186 | 0.367 |
| opioid use disorder – control | 0.03918 | 0.2732 | 0.1434 | 0.887 |
| opioid+alcohol use disorder – control | -1.02114 | 0.3230 | -3.1610 | 0.004 |
| Age Cateogry: | | | | |
| 50-64 – 30-49 | 0.28666 | 0.2911 | 0.9849 | 0.334 |
| 50-65 – 30-49 | 0.04569 | 0.4471 | 0.1022 | 0.919 |
| 65+ – 30-49 | 0.31158 | 0.3372 | 0.9240 | 0.365 |
| <30 – 30-49 | 0.39323 | 0.3525 | 1.1154 | 0.276 |
| Ethnicity: | | | | |
| Asian – White | 0.05764 | 0.5870 | 0.0982 | 0.923 |
| Black – White | -0.18617 | 0.2390 | -0.7790 | 0.444 |
| Hispanic – White | 0.02462 | 0.4571 | 0.0539 | 0.957 |
| Gender: | | | | |
| Male – Female | -0.19355 | 0.2825 | -0.6852 | 0.500 |
| PMI converted to hours | 0.00804 | 0.0117 | 0.6860 | 0.499 |
| pH | -0.10275 | 0.4052 | -0.2536 | 0.802 |

[a] Represents reference level

### Linear Regression

Model Fit Measures

| Model | R | $R^2$ |
|---|---|---|
| 1 | 0.609 | 0.370 |

Model Coefficients - CORO1A

| Predictor | Estimate | SE | t | p |
|---|---|---|---|---|
| Intercept [a] | 28.99932 | 1.16013 | 24.9967 | < .001 |
| Arm: | | | | |
| alcohol use disorder – control | -0.11810 | 0.12207 | -0.9675 | 0.343 |
| opioid use disorder – control | 0.05104 | 0.11419 | 0.4469 | 0.659 |
| opioid+alcohol use disorder – control | 0.31791 | 0.13504 | 2.3541 | 0.027 |
| Age Cateogry: | | | | |
| 50-64 – 30-49 | -0.01020 | 0.12167 | -0.0839 | 0.934 |
| 50-65 – 30-49 | 0.13724 | 0.18692 | 0.7342 | 0.470 |
| 65+ – 30-49 | 0.11459 | 0.14096 | 0.8129 | 0.424 |
| <30 – 30-49 | 0.01597 | 0.14738 | 0.1083 | 0.915 |
| Ethnicity: | | | | |
| Asian – White | 0.25295 | 0.24540 | 1.0308 | 0.313 |
| Black – White | 0.10266 | 0.09990 | 1.0276 | 0.314 |
| Hispanic – White | 0.16340 | 0.19107 | 0.8552 | 0.401 |
| Gender: | | | | |
| Male – Female | -0.00237 | 0.11809 | -0.0201 | 0.984 |
| PMI converted to hours | -0.00139 | 0.00490 | -0.2843 | 0.779 |
| pH | 0.14033 | 0.16939 | 0.8284 | 0.416 |

[a] Represents reference level

### Linear Regression

Model Fit Measures

| Model | R | $R^2$ |
|---|---|---|
| 1 | 0.691 | 0.477 |

Model Coefficients - GABRB3

| Predictor | Estimate | SE | t | p |
|---|---|---|---|---|
| Intercept [a] | 28.4028 | 3.3409 | 8.502 | < .001 |
| Arm: | | | | |
| alcohol use disorder – control | -0.1645 | 0.3515 | -0.468 | 0.644 |
| opioid use disorder – control | -0.3159 | 0.3289 | -0.961 | 0.346 |
| opioid+alcohol use disorder – control | -1.5382 | 0.3889 | -3.955 | < .001 |
| Age Cateogry: | | | | |
| 50-64 – 30-49 | 0.1092 | 0.3504 | 0.312 | 0.758 |
| 50-65 – 30-49 | -0.4078 | 0.5383 | -0.758 | 0.456 |
| 65+ – 30-49 | 0.0902 | 0.4059 | 0.222 | 0.826 |
| <30 – 30-49 | 0.6082 | 0.4244 | 1.433 | 0.165 |
| Ethnicity: | | | | |
| Asian – White | -0.7269 | 0.7067 | -1.029 | 0.314 |
| Black – White | -0.5665 | 0.2877 | -1.969 | 0.061 |
| Hispanic – White | 0.2918 | 0.5502 | 0.530 | 0.601 |
| Gender: | | | | |
| Male – Female | -0.3503 | 0.3401 | -1.030 | 0.313 |
| PMI converted to hours | -0.0108 | 0.0141 | -0.762 | 0.453 |
| pH | -0.5954 | 0.4878 | -1.221 | 0.234 |

[a] Represents reference level

### Linear Regression

Model Fit Measures

| Model | R | $R^2$ |
|---|---|---|
| 1 | 0.637 | 0.405 |

Model Coefficients - DGKQ

| Predictor | Estimate | SE | t | p |
|---|---|---|---|---|
| Intercept [a] | 27.73282 | 3.0165 | 9.19370 | < .001 |
| Arm: | | | | |
| alcohol use disorder – control | -0.00456 | 0.3174 | -0.01435 | 0.989 |
| opioid use disorder – control | 0.03085 | 0.2969 | 0.10390 | 0.918 |
| opioid+alcohol use disorder – control | -1.00854 | 0.3511 | -2.87223 | 0.008 |
| Age Cateogry: | | | | |
| 50-64 – 30-49 | 0.13454 | 0.3164 | 0.42527 | 0.674 |
| 50-65 – 30-49 | -0.12547 | 0.4860 | -0.25815 | 0.798 |
| 65+ – 30-49 | 0.32257 | 0.3665 | 0.88010 | 0.388 |
| <30 – 30-49 | -0.08437 | 0.3832 | -0.22018 | 0.828 |
| Ethnicity: | | | | |
| Asian – White | 0.19167 | 0.6381 | 0.30040 | 0.766 |
| Black – White | 0.07154 | 0.2598 | 0.27539 | 0.785 |
| Hispanic – White | -0.40550 | 0.4968 | -0.81619 | 0.422 |
| Gender: | | | | |
| Male – Female | -0.11896 | 0.3071 | -0.38743 | 0.702 |
| PMI converted to hours | -2.93e–5 | 0.0127 | -0.00230 | 0.998 |
| pH | -0.63849 | 0.4404 | -1.44964 | 0.160 |

[a] Represents reference level

### Linear Regression

Model Fit Measures

| Model | R | $R^2$ |
|---|---|---|
| 1 | 0.735 | 0.541 |

Model Coefficients - CAPZA1

| Predictor | Estimate | SE | t | p |
|---|---|---|---|---|
| Intercept [a] | 27.52902 | 1.02035 | 26.980 | < .001 |
| Arm: | | | | |
| alcohol use disorder – control | -0.05002 | 0.10737 | -0.466 | 0.646 |
| opioid use disorder – control | -0.19745 | 0.10044 | -1.966 | 0.061 |
| opioid+alcohol use disorder – control | -0.48006 | 0.11877 | -4.042 | < .001 |
| Age Cateogry: | | | | |
| 50-64 – 30-49 | 0.10108 | 0.10701 | 0.945 | 0.354 |
| 50-65 – 30-49 | -0.04503 | 0.16440 | -0.274 | 0.787 |
| 65+ – 30-49 | -0.12361 | 0.12398 | -0.997 | 0.329 |
| <30 – 30-49 | 0.27077 | 0.12962 | 2.089 | 0.047 |
| Ethnicity: | | | | |
| Asian – White | 0.07666 | 0.21583 | 0.355 | 0.726 |
| Black – White | -0.11552 | 0.08787 | -1.315 | 0.201 |
| Hispanic – White | 0.02660 | 0.16805 | 0.158 | 0.876 |
| Gender: | | | | |
| Male – Female | -0.19924 | 0.10386 | -1.918 | 0.067 |
| PMI converted to hours | -0.00208 | 0.00431 | -0.482 | 0.634 |
| pH | 0.12929 | 0.14898 | 0.868 | 0.394 |

[a] Represents reference level

### Linear Regression

Model Fit Measures

| Model | R | $R^2$ |
|---|---|---|
| 1 | 0.715 | 0.511 |

Model Coefficients - DBN1

| Predictor | Estimate | SE | t | p |
|---|---|---|---|---|
| Intercept [a] | 26.31745 | 1.99248 | 13.2084 | < .001 |
| Arm: | | | | |
| alcohol use disorder – control | -0.34340 | 0.20966 | -1.6379 | 0.114 |
| opioid use disorder – control | -0.20899 | 0.19613 | -1.0656 | 0.297 |
| opioid+alcohol use disorder – control | 0.53424 | 0.23193 | 2.3034 | 0.030 |
| Age Cateogry: | | | | |
| 50-64 – 30-49 | 0.03271 | 0.20897 | 0.1565 | 0.877 |
| 50-65 – 30-49 | 0.13860 | 0.32103 | 0.4317 | 0.670 |
| 65+ – 30-49 | 0.29842 | 0.24209 | 1.2326 | 0.230 |
| <30 – 30-49 | 0.07726 | 0.25311 | 0.3052 | 0.763 |
| Ethnicity: | | | | |
| Asian – White | 0.49409 | 0.42147 | 1.1723 | 0.253 |
| Black – White | 0.00739 | 0.17158 | 0.0431 | 0.966 |
| Hispanic – White | 0.49453 | 0.32816 | 1.5070 | 0.145 |
| Gender: | | | | |
| Male – Female | 0.14540 | 0.20282 | 0.7169 | 0.480 |
| PMI converted to hours | -0.00713 | 0.00842 | -0.8467 | 0.406 |
| pH | 0.40060 | 0.29093 | 1.3770 | 0.181 |

[a] Represents reference level

### Linear Regression

Model Fit Measures

| Model | R | $R^2$ |
|---|---|---|
| 1 | 0.664 | 0.441 |

Model Coefficients - USP49

| Predictor | Estimate | SE | t | p |
|---|---|---|---|---|
| Intercept [a] | 23.28922 | 2.6467 | 8.799 | < .001 |
| Arm: | | | | |
| alcohol use disorder – control | -0.36439 | 0.2785 | -1.308 | 0.203 |
| opioid use disorder – control | -0.02889 | 0.2605 | -0.111 | 0.913 |
| opioid+alcohol use disorder – control | -0.93008 | 0.3081 | -3.019 | 0.006 |
| Age Cateogry: | | | | |
| 50-64 – 30-49 | -0.18883 | 0.2776 | -0.680 | 0.503 |
| 50-65 – 30-49 | -0.05264 | 0.4264 | -0.123 | 0.903 |
| 65+ – 30-49 | 0.09929 | 0.3216 | 0.309 | 0.760 |
| <30 – 30-49 | 0.17540 | 0.3362 | 0.522 | 0.607 |
| Ethnicity: | | | | |
| Asian – White | -0.49706 | 0.5598 | -0.888 | 0.383 |
| Black – White | -0.11128 | 0.2279 | -0.488 | 0.630 |
| Hispanic – White | -0.07802 | 0.4359 | -0.179 | 0.859 |
| Gender: | | | | |
| Male – Female | -0.16830 | 0.2694 | -0.625 | 0.538 |
| PMI converted to hours | 0.00867 | 0.0112 | 0.776 | 0.445 |
| pH | 0.04610 | 0.3864 | 0.119 | 0.906 |

[a] Represents reference level

### Linear Regression

Model Fit Measures

| Model | R | $R^2$ |
|---|---|---|
| 1 | 0.682 | 0.465 |

Model Coefficients - DAK

| Predictor | Estimate | SE | t | p |
|---|---|---|---|---|
| Intercept [a] | 25.64259 | 4.2147 | 6.0841 | < .001 |
| Arm: | | | | |
| alcohol use disorder – control | 0.31033 | 0.4435 | 0.6998 | 0.491 |
| opioid use disorder – control | 0.10727 | 0.4149 | 0.2586 | 0.798 |
| opioid+alcohol use disorder – control | -1.33734 | 0.4906 | -2.7259 | 0.012 |
| Age Cateogry: | | | | |
| 50-64 – 30-49 | 0.38046 | 0.4420 | 0.8607 | 0.398 |
| 50-65 – 30-49 | 0.02674 | 0.6791 | 0.0394 | 0.969 |
| 65+ – 30-49 | -0.20274 | 0.5121 | -0.3959 | 0.696 |
| <30 – 30-49 | -0.46229 | 0.5354 | -0.8634 | 0.396 |
| Ethnicity: | | | | |
| Asian – White | 0.09356 | 0.8915 | 0.1049 | 0.917 |
| Black – White | -0.38610 | 0.3629 | -1.0638 | 0.298 |
| Hispanic – White | -0.12067 | 0.6942 | -0.1738 | 0.863 |
| Gender: | | | | |
| Male – Female | -0.17294 | 0.4290 | -0.4031 | 0.690 |
| PMI converted to hours | -0.00592 | 0.0178 | -0.3324 | 0.742 |
| pH | 0.08529 | 0.6154 | 0.1386 | 0.891 |

[a] Represents reference level

### Linear Regression

Model Fit Measures

| Model | R | $R^2$ |
|---|---|---|
| 1 | 0.607 | 0.368 |

Model Coefficients - PRKCD

| Predictor | Estimate | SE | t | p |
| --- | --- | --- | --- | --- |
| Intercept [a] | 22.21862 | 3.0234 | 7.3488 | < .001 |
| Arm: | | | | |
| alcohol use disorder – control | -0.00417 | 0.3181 | -0.0131 | 0.990 |
| opioid use disorder – control | 0.29571 | 0.2976 | 0.9936 | 0.330 |
| opioid+alcohol use disorder – control | -0.63482 | 0.3519 | -1.8038 | 0.084 |
| Age Cateogry: | | | | |
| 50-64 – 30-49 | -0.10654 | 0.3171 | -0.3360 | 0.740 |
| 50-65 – 30-49 | 0.52362 | 0.4871 | 1.0749 | 0.293 |
| 65+ – 30-49 | 0.14107 | 0.3674 | 0.3840 | 0.704 |
| <30 – 30-49 | -0.28052 | 0.3841 | -0.7304 | 0.472 |
| Ethnicity: | | | | |
| Asian – White | 0.00755 | 0.6395 | 0.0118 | 0.991 |
| Black – White | -0.07351 | 0.2604 | -0.2823 | 0.780 |
| Hispanic – White | 0.19884 | 0.4980 | 0.3993 | 0.693 |
| Gender: | | | | |
| Male – Female | 0.09018 | 0.3078 | 0.2930 | 0.772 |
| PMI converted to hours | 0.00557 | 0.0128 | 0.4364 | 0.666 |
| pH | 0.15277 | 0.4415 | 0.3461 | 0.732 |

[a] Represents reference level

### Linear Regression

Model Fit Measures

| Model | R | $R^2$ |
| --- | --- | --- |
| 1 | 0.735 | 0.540 |

Model Coefficients - GRIN2A

| Predictor | Estimate | SE | t | p |
|---|---|---|---|---|
| Intercept [a] | 27.1219 | 2.8509 | 9.513 | < .001 |
| Arm: | | | | |
| alcohol use disorder – control | -0.3390 | 0.3000 | -1.130 | 0.270 |
| opioid use disorder – control | 0.1430 | 0.2806 | 0.510 | 0.615 |
| opioid+alcohol use disorder – control | -1.1367 | 0.3319 | -3.425 | 0.002 |
| Age Cateogry: | | | | |
| 50-64 – 30-49 | 0.5025 | 0.2990 | 1.681 | 0.106 |
| 50-65 – 30-49 | 0.3333 | 0.4593 | 0.726 | 0.475 |
| 65+ – 30-49 | 0.6613 | 0.3464 | 1.909 | 0.068 |
| <30 – 30-49 | 0.1121 | 0.3622 | 0.310 | 0.760 |
| Ethnicity: | | | | |
| Asian – White | 0.1119 | 0.6031 | 0.186 | 0.854 |
| Black – White | 0.0399 | 0.2455 | 0.163 | 0.872 |
| Hispanic – White | -0.7739 | 0.4695 | -1.648 | 0.112 |
| Gender: | | | | |
| Male – Female | -0.2674 | 0.2902 | -0.921 | 0.366 |
| PMI converted to hours | 0.0109 | 0.0120 | 0.901 | 0.376 |
| pH | -0.5714 | 0.4163 | -1.373 | 0.183 |

[a] Represents reference level

### Linear Regression

Model Fit Measures

| Model | R | $R^2$ |
|---|---|---|
| 1 | 0.660 | 0.436 |

Model Coefficients - RGS7BP

| Predictor | Estimate | SE | t | p |
|---|---|---|---|---|
| Intercept [a] | 24.6864 | 3.5835 | 6.8889 | < .001 |
| Arm: | | | | |
| alcohol use disorder – control | -0.2288 | 0.3771 | -0.6069 | 0.550 |
| opioid use disorder – control | -0.2468 | 0.3527 | -0.6996 | 0.491 |
| opioid+alcohol use disorder – control | -1.0850 | 0.4171 | -2.6011 | 0.016 |
| Age Cateogry: | | | | |
| 50-64 – 30-49 | -0.4289 | 0.3758 | -1.1413 | 0.265 |
| 50-65 – 30-49 | -0.1647 | 0.5774 | -0.2852 | 0.778 |
| 65+ – 30-49 | -0.2204 | 0.4354 | -0.5062 | 0.617 |
| <30 – 30-49 | -0.0731 | 0.4552 | -0.1606 | 0.874 |
| Ethnicity: | | | | |
| Asian – White | -1.1250 | 0.7580 | -1.4841 | 0.151 |
| Black – White | -0.1031 | 0.3086 | -0.3342 | 0.741 |
| Hispanic – White | 0.7722 | 0.5902 | 1.3083 | 0.203 |
| Gender: | | | | |
| Male – Female | -0.1181 | 0.3648 | -0.3238 | 0.749 |
| PMI converted to hours | -0.0211 | 0.0151 | -1.3940 | 0.176 |
| pH | 0.0418 | 0.5232 | 0.0800 | 0.937 |

[a] Represents reference level

### Linear Regression

Model Fit Measures

| Model | R | $R^2$ |
|---|---|---|
| 1 | 0.567 | 0.321 |

Model Coefficients - UBE4A

| Predictor | Estimate | SE | t | p |
| --- | --- | --- | --- | --- |
| Intercept [a] | 21.90743 | 3.5251 | 6.2147 | < .001 |
| Arm: | | | | |
| alcohol use disorder – control | -0.27107 | 0.3709 | -0.7308 | 0.472 |
| opioid use disorder – control | 0.27062 | 0.3470 | 0.7799 | 0.443 |
| opioid+alcohol use disorder – control | -0.65003 | 0.4103 | -1.5842 | 0.126 |
| Age Cateogry: | | | | |
| 50-64 – 30-49 | 0.12011 | 0.3697 | 0.3249 | 0.748 |
| 50-65 – 30-49 | 0.59744 | 0.5680 | 1.0519 | 0.303 |
| 65+ – 30-49 | 0.02869 | 0.4283 | 0.0670 | 0.947 |
| <30 – 30-49 | 0.22345 | 0.4478 | 0.4990 | 0.622 |
| Ethnicity: | | | | |
| Asian – White | 0.21688 | 0.7457 | 0.2909 | 0.774 |
| Black – White | -0.04624 | 0.3036 | -0.1523 | 0.880 |
| Hispanic – White | -0.00978 | 0.5806 | -0.0168 | 0.987 |
| Gender: | | | | |
| Male – Female | -0.16861 | 0.3588 | -0.4699 | 0.643 |
| PMI converted to hours | 0.01418 | 0.0149 | 0.9527 | 0.350 |
| pH | 0.18356 | 0.5147 | 0.3566 | 0.724 |

[a] Represents reference level

### Linear Regression

Model Fit Measures

| Model | R | $R^2$ |
| --- | --- | --- |
| 1 | 0.738 | 0.545 |

Model Coefficients - CCBL2 (2)

| Predictor | Estimate | SE | t | p |
| --- | --- | --- | --- | --- |
| Intercept [a] | 20.7383 | 4.7822 | 4.3366 | < .001 |
| Arm: | | | | |
| alcohol use disorder – control | -0.2624 | 0.5032 | -0.5214 | 0.607 |
| opioid use disorder – control | -1.2648 | 0.4707 | -2.6869 | 0.013 |
| opioid+alcohol use disorder – control | -1.4566 | 0.5567 | -2.6167 | 0.015 |
| Age Cateogry: | | | | |
| 50-64 – 30-49 | -0.1991 | 0.5015 | -0.3969 | 0.695 |
| 50-65 – 30-49 | 0.1550 | 0.7705 | 0.2011 | 0.842 |
| 65+ – 30-49 | -0.1481 | 0.5811 | -0.2548 | 0.801 |
| <30 – 30-49 | 0.2484 | 0.6075 | 0.4089 | 0.686 |
| Ethnicity: | | | | |
| Asian – White | -0.0203 | 1.0116 | -0.0201 | 0.984 |
| Black – White | -0.5336 | 0.4118 | -1.2956 | 0.207 |
| Hispanic – White | -0.8877 | 0.7876 | -1.1270 | 0.271 |
| Gender: | | | | |
| Male – Female | -0.6024 | 0.4868 | -1.2375 | 0.228 |
| PMI converted to hours | 0.0602 | 0.0202 | 2.9806 | 0.006 |
| pH | 0.8574 | 0.6983 | 1.2279 | 0.231 |

[a] Represents reference level

### Linear Regression

Model Fit Measures

| Model | R | $R^2$ |
| --- | --- | --- |
| 1 | 0.631 | 0.398 |

Model Coefficients - UBE2G1

| Predictor | Estimate | SE | t | p |
|---|---|---|---|---|
| Intercept [a] | 25.07371 | 3.1912 | 7.85721 | < .001 |
| Arm: | | | | |
| alcohol use disorder – control | -0.00219 | 0.3358 | -0.00651 | 0.995 |
| opioid use disorder – control | 0.32157 | 0.3141 | 1.02372 | 0.316 |
| opioid+alcohol use disorder – control | -0.80123 | 0.3715 | -2.15693 | 0.041 |
| Age Cateogry: | | | | |
| 50-64 – 30-49 | 0.50404 | 0.3347 | 1.50602 | 0.145 |
| 50-65 – 30-49 | 0.37590 | 0.5142 | 0.73109 | 0.472 |
| 65+ – 30-49 | 0.14804 | 0.3877 | 0.38179 | 0.706 |
| <30 – 30-49 | 0.43180 | 0.4054 | 1.06515 | 0.297 |
| Ethnicity: | | | | |
| Asian – White | 0.53088 | 0.6750 | 0.78646 | 0.439 |
| Black – White | 0.07077 | 0.2748 | 0.25752 | 0.799 |
| Hispanic – White | -0.26465 | 0.5256 | -0.50353 | 0.619 |
| Gender: | | | | |
| Male – Female | -0.36123 | 0.3248 | -1.11207 | 0.277 |
| PMI converted to hours | 0.00856 | 0.0135 | 0.63482 | 0.532 |
| pH | -0.27532 | 0.4659 | -0.59088 | 0.560 |

[a] Represents reference level

### Linear Regression

Model Fit Measures

| Model | R | $R^2$ |
|---|---|---|
| 1 | 0.759 | 0.576 |

Model Coefficients - IMMT

| Predictor | Estimate | SE | t | p |
|---|---|---|---|---|
| Intercept [a] | 32.0028 | 0.71046 | 45.0451 | < .001 |
| Arm: | | | | |
| alcohol use disorder – control | 0.0981 | 0.07476 | 1.3129 | 0.202 |
| opioid use disorder – control | 0.0377 | 0.06993 | 0.5387 | 0.595 |
| opioid+alcohol use disorder – control | -0.2679 | 0.08270 | -3.2389 | 0.003 |
| Age Cateogry: | | | | |
| 50-64 – 30-49 | -0.0142 | 0.07451 | -0.1909 | 0.850 |
| 50-65 – 30-49 | 0.0993 | 0.11447 | 0.8676 | 0.394 |
| 65+ – 30-49 | -0.1354 | 0.08632 | -1.5690 | 0.130 |
| <30 – 30-49 | 0.0702 | 0.09025 | 0.7779 | 0.444 |
| Ethnicity: | | | | |
| Asian – White | -0.1313 | 0.15028 | -0.8736 | 0.391 |
| Black – White | -0.1323 | 0.06118 | -2.1628 | 0.041 |
| Hispanic – White | -0.1861 | 0.11701 | -1.5902 | 0.125 |
| Gender: | | | | |
| Male – Female | -0.1139 | 0.07232 | -1.5755 | 0.128 |
| PMI converted to hours | -4.63e−5 | 0.00300 | -0.0154 | 0.988 |
| pH | -0.0525 | 0.10374 | -0.5065 | 0.617 |

[a] Represents reference level

### Linear Regression

Model Fit Measures

| Model | R | $R^2$ |
|---|---|---|
| 1 | 0.635 | 0.403 |

Model Coefficients - OTUD7B

| Predictor | Estimate | SE | t | p |
|---|---|---|---|---|
| Intercept [a] | 25.73783 | 3.0660 | 8.395 | < .001 |
| Arm: | | | | |
| alcohol use disorder – control | -0.13509 | 0.3226 | -0.419 | 0.679 |
| opioid use disorder – control | -0.12033 | 0.3018 | -0.399 | 0.694 |
| opioid+alcohol use disorder – control | -1.21772 | 0.3569 | -3.412 | 0.002 |
| Age Cateogry: | | | | |
| 50-64 – 30-49 | 0.21478 | 0.3216 | 0.668 | 0.511 |
| 50-65 – 30-49 | -0.17298 | 0.4940 | -0.350 | 0.729 |
| 65+ – 30-49 | 0.21979 | 0.3725 | 0.590 | 0.561 |
| <30 – 30-49 | 0.10419 | 0.3895 | 0.268 | 0.791 |
| Ethnicity: | | | | |
| Asian – White | -0.45698 | 0.6485 | -0.705 | 0.488 |
| Black – White | -0.11382 | 0.2640 | -0.431 | 0.670 |
| Hispanic – White | -0.43148 | 0.5050 | -0.854 | 0.401 |
| Gender: | | | | |
| Male – Female | -0.32863 | 0.3121 | -1.053 | 0.303 |
| PMI converted to hours | -0.00594 | 0.0129 | -0.459 | 0.651 |
| pH | -0.23995 | 0.4477 | -0.536 | 0.597 |

[a] Represents reference level

### Linear Regression

Model Fit Measures

| Model | R | $R^2$ |
|---|---|---|
| 1 | 0.667 | 0.445 |

Model Coefficients - GOLGA7B

| Predictor | Estimate | SE | t | p |
|---|---|---|---|---|
| Intercept [a] | 28.3587 | 3.2225 | 8.8001 | < .001 |
| Arm: | | | | |
| alcohol use disorder – control | 0.0326 | 0.3391 | 0.0962 | 0.924 |
| opioid use disorder – control | 0.3473 | 0.3172 | 1.0949 | 0.284 |
| opioid+alcohol use disorder – control | -0.9954 | 0.3751 | -2.6536 | 0.014 |
| Age Cateogry: | | | | |
| 50-64 – 30-49 | 0.5916 | 0.3380 | 1.7504 | 0.093 |
| 50-65 – 30-49 | -0.2521 | 0.5192 | -0.4856 | 0.632 |
| 65+ – 30-49 | 0.3523 | 0.3916 | 0.8998 | 0.377 |
| <30 – 30-49 | 0.1273 | 0.4094 | 0.3109 | 0.759 |
| Ethnicity: | | | | |
| Asian – White | 0.0460 | 0.6817 | 0.0674 | 0.947 |
| Black – White | -0.0272 | 0.2775 | -0.0979 | 0.923 |
| Hispanic – White | -0.5761 | 0.5308 | -1.0855 | 0.288 |
| Gender: | | | | |
| Male – Female | -0.2654 | 0.3280 | -0.8091 | 0.426 |
| PMI converted to hours | 0.0183 | 0.0136 | 1.3423 | 0.192 |
| pH | -0.8182 | 0.4705 | -1.7390 | 0.095 |

[a] Represents reference level

### Linear Regression

Model Fit Measures

| Model | R | $R^2$ |
|---|---|---|
| 1 | 0.687 | 0.472 |

Model Coefficients - KANK2

| Predictor | Estimate | SE | t | p |
|---|---|---|---|---|
| Intercept [a] | 26.20358 | 2.8789 | 9.1021 | < .001 |
| Arm: | | | | |
| alcohol use disorder – control | -0.16435 | 0.3029 | -0.5425 | 0.592 |
| opioid use disorder – control | 0.32356 | 0.2834 | 1.1418 | 0.265 |
| opioid+alcohol use disorder – control | -0.88521 | 0.3351 | -2.6415 | 0.014 |
| Age Cateogry: | | | | |
| 50-64 – 30-49 | 0.49304 | 0.3019 | 1.6330 | 0.116 |
| 50-65 – 30-49 | 0.35204 | 0.4638 | 0.7590 | 0.455 |
| 65+ – 30-49 | 0.76539 | 0.3498 | 2.1881 | 0.039 |
| <30 – 30-49 | 0.12098 | 0.3657 | 0.3308 | 0.744 |
| Ethnicity: | | | | |
| Asian – White | -0.23789 | 0.6090 | -0.3906 | 0.700 |
| Black – White | 0.00539 | 0.2479 | 0.0218 | 0.983 |
| Hispanic – White | -0.42154 | 0.4741 | -0.8891 | 0.383 |
| Gender: | | | | |
| Male – Female | 0.04892 | 0.2930 | 0.1669 | 0.869 |
| PMI converted to hours | 0.01189 | 0.0122 | 0.9782 | 0.338 |
| pH | -0.51556 | 0.4203 | -1.2265 | 0.232 |

[a] Represents reference level

### Linear Regression

Model Fit Measures

| Model | R | $R^2$ |
|---|---|---|
| 1 | 0.691 | 0.477 |

Model Coefficients - PDE12

| Predictor | Estimate | SE | t | p |
|---|---|---|---|---|
| Intercept [a] | 21.01378 | 4.4211 | 4.753 | < .001 |
| Arm: | | | | |
| alcohol use disorder – control | -0.58557 | 0.4652 | -1.259 | 0.220 |
| opioid use disorder – control | -0.63376 | 0.4352 | -1.456 | 0.158 |
| opioid+alcohol use disorder – control | -1.44433 | 0.5146 | -2.807 | 0.010 |
| Age Cateogry: | | | | |
| 50-64 – 30-49 | -0.47876 | 0.4637 | -1.033 | 0.312 |
| 50-65 – 30-49 | -0.39935 | 0.7123 | -0.561 | 0.580 |
| 65+ – 30-49 | -0.48444 | 0.5372 | -0.902 | 0.376 |
| <30 – 30-49 | -0.25581 | 0.5616 | -0.455 | 0.653 |
| Ethnicity: | | | | |
| Asian – White | 0.36188 | 0.9352 | 0.387 | 0.702 |
| Black – White | 0.57322 | 0.3807 | 1.506 | 0.145 |
| Hispanic – White | 0.22960 | 0.7281 | 0.315 | 0.755 |
| Gender: | | | | |
| Male – Female | -0.52076 | 0.4500 | -1.157 | 0.259 |
| PMI converted to hours | 0.00829 | 0.0187 | 0.444 | 0.661 |
| pH | 0.62089 | 0.6455 | 0.962 | 0.346 |

[a] Represents reference level

### Linear Regression

Model Fit Measures

| Model | R | $R^2$ |
|---|---|---|
| 1 | 0.691 | 0.478 |

Model Coefficients - PDK3

| Predictor | Estimate | SE | t | p |
|---|---|---|---|---|
| Intercept [a] | 28.61468 | 3.6805 | 7.7746 | < .001 |
| Arm: | | | | |
| alcohol use disorder – control | 0.42223 | 0.3873 | 1.0902 | 0.286 |
| opioid use disorder – control | -0.03418 | 0.3623 | -0.0944 | 0.926 |
| opioid+alcohol use disorder – control | -1.44240 | 0.4284 | -3.3667 | 0.003 |
| Age Cateogry: | | | | |
| 50-64 – 30-49 | 0.46878 | 0.3860 | 1.2144 | 0.236 |
| 50-65 – 30-49 | -0.20669 | 0.5930 | -0.3486 | 0.730 |
| 65+ – 30-49 | -0.60156 | 0.4472 | -1.3452 | 0.191 |
| <30 – 30-49 | 0.37315 | 0.4676 | 0.7981 | 0.433 |
| Ethnicity: | | | | |
| Asian – White | -0.00941 | 0.7785 | -0.0121 | 0.990 |
| Black – White | -0.10940 | 0.3169 | -0.3452 | 0.733 |
| Hispanic – White | -0.44653 | 0.6062 | -0.7366 | 0.468 |
| Gender: | | | | |
| Male – Female | -0.39714 | 0.3746 | -1.0601 | 0.300 |
| PMI converted to hours | -0.00394 | 0.0155 | -0.2533 | 0.802 |
| pH | -0.27881 | 0.5374 | -0.5188 | 0.609 |

[a] Represents reference level

### Linear Regression

Model Fit Measures

| Model | R | $R^2$ |
|---|---|---|
| 1 | 0.687 | 0.472 |

Model Coefficients - EFR3A

| Predictor | Estimate | SE | t | p |
|---|---|---|---|---|
| Intercept [a] | 27.31890 | 3.6613 | 7.4615 | < .001 |
| Arm: | | | | |
| alcohol use disorder – control | -0.04706 | 0.3853 | -0.1221 | 0.904 |
| opioid use disorder – control | 0.18309 | 0.3604 | 0.5080 | 0.616 |
| opioid+alcohol use disorder – control | -1.41178 | 0.4262 | -3.3125 | 0.003 |
| Age Cateogry: | | | | |
| 50-64 – 30-49 | 0.17125 | 0.3840 | 0.4460 | 0.660 |
| 50-65 – 30-49 | -0.05602 | 0.5899 | -0.0950 | 0.925 |
| 65+ – 30-49 | 0.25206 | 0.4449 | 0.5666 | 0.576 |
| <30 – 30-49 | -0.06290 | 0.4651 | -0.1352 | 0.894 |
| Ethnicity: | | | | |
| Asian – White | -0.00913 | 0.7745 | -0.0118 | 0.991 |
| Black – White | -0.14317 | 0.3153 | -0.4541 | 0.654 |
| Hispanic – White | -0.82108 | 0.6030 | -1.3616 | 0.186 |
| Gender: | | | | |
| Male – Female | -0.41917 | 0.3727 | -1.1247 | 0.272 |
| PMI converted to hours | 0.02056 | 0.0155 | 1.3298 | 0.196 |
| pH | -0.62051 | 0.5346 | -1.1607 | 0.257 |

[a] Represents reference level

### Linear Regression

Model Fit Measures

| Model | R | $R^2$ |
|---|---|---|
| 1 | 0.679 | 0.462 |

Model Coefficients - YWHAZ

| Predictor | Estimate | SE | t | p |
|---|---|---|---|---|
| Intercept [a] | 33.55904 | 0.99771 | 33.6360 | < .001 |
| Arm: | | | | |
| alcohol use disorder – control | -0.07677 | 0.10498 | -0.7313 | 0.472 |
| opioid use disorder – control | -0.02148 | 0.09821 | -0.2187 | 0.829 |
| opioid+alcohol use disorder – control | 0.27553 | 0.11614 | 2.3724 | 0.026 |
| Age Cateogry: | | | | |
| 50-64 – 30-49 | 0.02857 | 0.10464 | 0.2730 | 0.787 |
| 50-65 – 30-49 | 0.07358 | 0.16075 | 0.4577 | 0.651 |
| 65+ – 30-49 | 0.13309 | 0.12123 | 1.0979 | 0.283 |
| <30 – 30-49 | 0.15578 | 0.12674 | 1.2291 | 0.231 |
| Ethnicity: | | | | |
| Asian – White | 0.21348 | 0.21104 | 1.0115 | 0.322 |
| Black – White | 0.04430 | 0.08592 | 0.5156 | 0.611 |
| Hispanic – White | 0.26805 | 0.16432 | 1.6313 | 0.116 |
| Gender: | | | | |
| Male – Female | 0.08683 | 0.10156 | 0.8550 | 0.401 |
| PMI converted to hours | -0.00841 | 0.00421 | -1.9964 | 0.057 |
| pH | 0.01257 | 0.14568 | 0.0863 | 0.932 |

[a] Represents reference level

### Linear Regression

Model Fit Measures

| Model | R | $R^2$ |
|---|---|---|
| 1 | 0.667 | 0.445 |

Model Coefficients - SLC4A10

| Predictor | Estimate | SE | t | p |
|---|---|---|---|---|
| Intercept [a] | 26.28281 | 1.51644 | 17.332 | < .001 |
| Arm: | | | | |
| alcohol use disorder – control | -0.09405 | 0.15957 | -0.589 | 0.561 |
| opioid use disorder – control | -0.05162 | 0.14927 | -0.346 | 0.733 |
| opioid+alcohol use disorder – control | 0.40829 | 0.17652 | 2.313 | 0.030 |
| Age Cateogry: | | | | |
| 50-64 – 30-49 | -0.17507 | 0.15904 | -1.101 | 0.282 |
| 50-65 – 30-49 | 0.10432 | 0.24433 | 0.427 | 0.673 |
| 65+ – 30-49 | -0.20796 | 0.18425 | -1.129 | 0.270 |
| <30 – 30-49 | -0.16669 | 0.19264 | -0.865 | 0.395 |
| Ethnicity: | | | | |
| Asian – White | 0.51699 | 0.32077 | 1.612 | 0.120 |
| Black – White | 0.12191 | 0.13059 | 0.934 | 0.360 |
| Hispanic – White | -1.10e−4 | 0.24976 | -4.39e−4 | 1.000 |
| Gender: | | | | |
| Male – Female | -0.17298 | 0.15436 | -1.121 | 0.274 |
| PMI converted to hours | 0.00674 | 0.00640 | 1.052 | 0.303 |
| pH | 0.21205 | 0.22142 | 0.958 | 0.348 |

[a] Represents reference level

### Linear Regression

Model Fit Measures

| Model | R | $R^2$ |
|---|---|---|
| 1 | 0.633 | 0.401 |

Model Coefficients - CLPP

| Predictor | Estimate | SE | t | p |
| --- | --- | --- | --- | --- |
| Intercept [a] | 24.10929 | 2.8473 | 8.4675 | < .001 |
| Arm: | | | | |
| alcohol use disorder – control | -0.30373 | 0.2996 | -1.0138 | 0.321 |
| opioid use disorder – control | 0.06803 | 0.2803 | 0.2427 | 0.810 |
| opioid+alcohol use disorder – control | -0.92671 | 0.3314 | -2.7961 | 0.010 |
| Age Cateogry: | | | | |
| 50-64 – 30-49 | 0.07123 | 0.2986 | 0.2385 | 0.813 |
| 50-65 – 30-49 | 0.00999 | 0.4588 | 0.0218 | 0.983 |
| 65+ – 30-49 | 0.10485 | 0.3460 | 0.3031 | 0.764 |
| <30 – 30-49 | -0.35619 | 0.3617 | -0.9847 | 0.335 |
| Ethnicity: | | | | |
| Asian – White | -0.11924 | 0.6023 | -0.1980 | 0.845 |
| Black – White | -0.19399 | 0.2452 | -0.7912 | 0.437 |
| Hispanic – White | -0.26670 | 0.4689 | -0.5687 | 0.575 |
| Gender: | | | | |
| Male – Female | 0.02125 | 0.2898 | 0.0733 | 0.942 |
| PMI converted to hours | 0.01252 | 0.0120 | 1.0413 | 0.308 |
| pH | -0.11414 | 0.4157 | -0.2746 | 0.786 |

[a] Represents reference level

### Linear Regression

Model Fit Measures

| Model | R | $R^2$ |
| --- | --- | --- |
| 1 | 0.630 | 0.397 |

Model Coefficients - SYPL1

| Predictor | Estimate | SE | t | p |
|---|---|---|---|---|
| Intercept [a] | 27.12631 | 3.1884 | 8.5078 | < .001 |
| Arm: | | | | |
| alcohol use disorder – control | 0.03903 | 0.3355 | 0.1163 | 0.908 |
| opioid use disorder – control | -0.15460 | 0.3138 | -0.4926 | 0.627 |
| opioid+alcohol use disorder – control | -1.15684 | 0.3711 | -3.1170 | 0.005 |
| Age Cateogry: | | | | |
| 50-64 – 30-49 | 0.34299 | 0.3344 | 1.0257 | 0.315 |
| 50-65 – 30-49 | -0.31507 | 0.5137 | -0.6133 | 0.545 |
| 65+ – 30-49 | 0.29021 | 0.3874 | 0.7491 | 0.461 |
| <30 – 30-49 | 0.02214 | 0.4050 | 0.0547 | 0.957 |
| Ethnicity: | | | | |
| Asian – White | -0.63334 | 0.6744 | -0.9391 | 0.357 |
| Black – White | -0.16179 | 0.2746 | -0.5892 | 0.561 |
| Hispanic – White | -1.05499 | 0.5251 | -2.0090 | 0.056 |
| Gender: | | | | |
| Male – Female | -0.18470 | 0.3245 | -0.5691 | 0.575 |
| PMI converted to hours | -0.00128 | 0.0135 | -0.0949 | 0.925 |
| pH | -0.52187 | 0.4655 | -1.1210 | 0.273 |

[a] Represents reference level

### Linear Regression

Model Fit Measures

| Model | R | $R^2$ |
|---|---|---|
| 1 | 0.718 | 0.516 |

Model Coefficients - SEC61B

| Predictor | Estimate | SE | t | p |
|---|---|---|---|---|
| Intercept [a] | 25.01676 | 2.4560 | 10.186 | < .001 |
| Arm: | | | | |
| alcohol use disorder – control | -0.24362 | 0.2584 | -0.943 | 0.355 |
| opioid use disorder – control | 0.44172 | 0.2417 | 1.827 | 0.080 |
| opioid+alcohol use disorder – control | -0.61937 | 0.2859 | -2.166 | 0.040 |
| Age Cateogry: | | | | |
| 50-64 – 30-49 | 0.14385 | 0.2576 | 0.558 | 0.582 |
| 50-65 – 30-49 | 0.38578 | 0.3957 | 0.975 | 0.339 |
| 65+ – 30-49 | 0.41365 | 0.2984 | 1.386 | 0.178 |
| <30 – 30-49 | -0.15375 | 0.3120 | -0.493 | 0.627 |
| Ethnicity: | | | | |
| Asian – White | -0.06370 | 0.5195 | -0.123 | 0.903 |
| Black – White | 0.19035 | 0.2115 | 0.900 | 0.377 |
| Hispanic – White | 0.23151 | 0.4045 | 0.572 | 0.572 |
| Gender: | | | | |
| Male – Female | -0.05985 | 0.2500 | -0.239 | 0.813 |
| PMI converted to hours | 0.01000 | 0.0104 | 0.964 | 0.345 |
| pH | -0.28432 | 0.3586 | -0.793 | 0.436 |

[a] Represents reference level

### Linear Regression

Model Fit Measures

| Model | R | $R^2$ |
|---|---|---|
| 1 | 0.827 | 0.684 |

Model Coefficients - RPL32

| Predictor | Estimate | SE | t | p |
|---|---|---|---|---|
| Intercept [a] | 20.95320 | 3.2343 | 6.478 | < .001 |
| Arm: | | | | |
| alcohol use disorder – control | 0.49336 | 0.3403 | 1.450 | 0.160 |
| opioid use disorder – control | 0.92408 | 0.3184 | 2.903 | 0.008 |
| opioid+alcohol use disorder – control | -0.43496 | 0.3765 | -1.155 | 0.259 |
| Age Cateogry: | | | | |
| 50-64 – 30-49 | -0.42492 | 0.3392 | -1.253 | 0.222 |
| 50-65 – 30-49 | -0.46071 | 0.5211 | -0.884 | 0.385 |
| 65+ – 30-49 | -0.26429 | 0.3930 | -0.673 | 0.508 |
| <30 – 30-49 | -1.25363 | 0.4109 | -3.051 | 0.005 |
| Ethnicity: | | | | |
| Asian – White | -0.36388 | 0.6841 | -0.532 | 0.600 |
| Black – White | 0.52476 | 0.2785 | 1.884 | 0.072 |
| Hispanic – White | 0.14103 | 0.5327 | 0.265 | 0.793 |
| Gender: | | | | |
| Male – Female | 0.44103 | 0.3292 | 1.340 | 0.193 |
| PMI converted to hours | 0.00833 | 0.0137 | 0.610 | 0.548 |
| pH | 0.35161 | 0.4722 | 0.745 | 0.464 |

[a] Represents reference level

### Linear Regression

Model Fit Measures

| Model | R | $R^2$ |
|---|---|---|
| 1 | 0.802 | 0.644 |

Model Coefficients - COA6

| Predictor | Estimate | SE | t | p |
|---|---|---|---|---|
| Intercept [a] | 25.00818 | 1.34637 | 18.5746 | < .001 |
| Arm: | | | | |
| alcohol use disorder – control | 0.00592 | 0.14167 | 0.0418 | 0.967 |
| opioid use disorder – control | 0.30917 | 0.13253 | 2.3329 | 0.028 |
| opioid+alcohol use disorder – control | -0.35457 | 0.15672 | -2.2624 | 0.033 |
| Age Cateogry: | | | | |
| 50-64 – 30-49 | -0.23789 | 0.14120 | -1.6847 | 0.105 |
| 50-65 – 30-49 | 0.09959 | 0.21693 | 0.4591 | 0.650 |
| 65+ – 30-49 | 0.04475 | 0.16359 | 0.2736 | 0.787 |
| <30 – 30-49 | -0.29930 | 0.17104 | -1.7499 | 0.093 |
| Ethnicity: | | | | |
| Asian – White | -0.22708 | 0.28479 | -0.7973 | 0.433 |
| Black – White | 0.04231 | 0.11594 | 0.3650 | 0.718 |
| Hispanic – White | -0.19058 | 0.22175 | -0.8594 | 0.399 |
| Gender: | | | | |
| Male – Female | 0.11928 | 0.13705 | 0.8703 | 0.393 |
| PMI converted to hours | 0.01082 | 0.00569 | 1.9034 | 0.069 |
| pH | 0.22032 | 0.19659 | 1.1207 | 0.273 |

[a] Represents reference level

### Linear Regression

Model Fit Measures

| Model | R | $R^2$ |
|---|---|---|
| 1 | 0.667 | 0.445 |

Model Coefficients - TWF2

| Predictor | Estimate | SE | t | p |
| --- | --- | --- | --- | --- |
| Intercept [a] | 27.30876 | 2.3706 | 11.520 | < .001 |
| Arm: | | | | |
| alcohol use disorder – control | -0.34662 | 0.2494 | -1.390 | 0.177 |
| opioid use disorder – control | -0.39314 | 0.2333 | -1.685 | 0.105 |
| opioid+alcohol use disorder – control | 0.40737 | 0.2759 | 1.476 | 0.153 |
| Age Cateogry: | | | | |
| 50-64 – 30-49 | 0.03888 | 0.2486 | 0.156 | 0.877 |
| 50-65 – 30-49 | 0.21245 | 0.3819 | 0.556 | 0.583 |
| 65+ – 30-49 | 0.05283 | 0.2880 | 0.183 | 0.856 |
| <30 – 30-49 | 0.11812 | 0.3011 | 0.392 | 0.698 |
| Ethnicity: | | | | |
| Asian – White | 0.41968 | 0.5014 | 0.837 | 0.411 |
| Black – White | 0.14374 | 0.2041 | 0.704 | 0.488 |
| Hispanic – White | 0.42562 | 0.3904 | 1.090 | 0.286 |
| Gender: | | | | |
| Male – Female | -0.20112 | 0.2413 | -0.833 | 0.413 |
| PMI converted to hours | -0.00652 | 0.0100 | -0.651 | 0.521 |
| pH | 0.16325 | 0.3461 | 0.472 | 0.641 |

[a] Represents reference level

### Linear Regression

Model Fit Measures

| Model | R | $R^2$ |
| --- | --- | --- |
| 1 | 0.676 | 0.457 |

Model Coefficients - SLC25A42

| Predictor | Estimate | SE | t | p |
|---|---|---|---|---|
| Intercept [a] | 27.63854 | 3.1571 | 8.7544 | < .001 |
| Arm: | | | | |
| alcohol use disorder – control | -0.35323 | 0.3322 | -1.0633 | 0.298 |
| opioid use disorder – control | 0.17730 | 0.3108 | 0.5705 | 0.574 |
| opioid+alcohol use disorder – control | -0.94420 | 0.3675 | -2.5693 | 0.017 |
| Age Cateogry: | | | | |
| 50-64 – 30-49 | -0.00640 | 0.3311 | -0.0193 | 0.985 |
| 50-65 – 30-49 | -0.45276 | 0.5087 | -0.8901 | 0.382 |
| 65+ – 30-49 | 0.06014 | 0.3836 | 0.1568 | 0.877 |
| <30 – 30-49 | -0.02355 | 0.4011 | -0.0587 | 0.954 |
| Ethnicity: | | | | |
| Asian – White | -0.75040 | 0.6678 | -1.1237 | 0.272 |
| Black – White | 0.23181 | 0.2719 | 0.8526 | 0.402 |
| Hispanic – White | -0.35703 | 0.5200 | -0.6866 | 0.499 |
| Gender: | | | | |
| Male – Female | 0.03281 | 0.3214 | 0.1021 | 0.920 |
| PMI converted to hours | 0.00263 | 0.0133 | 0.1971 | 0.845 |
| pH | -0.54764 | 0.4610 | -1.1880 | 0.246 |

[a] Represents reference level

### Linear Regression

Model Fit Measures

| Model | R | $R^2$ |
|---|---|---|
| 1 | 0.654 | 0.427 |

Model Coefficients - CYB5R4

| Predictor | Estimate | SE | t | p |
|---|---|---|---|---|
| Intercept [a] | 22.54020 | 2.9383 | 7.6713 | < .001 |
| Arm: | | | | |
| alcohol use disorder – control | -0.12955 | 0.3092 | -0.4190 | 0.679 |
| opioid use disorder – control | 0.35007 | 0.2892 | 1.2104 | 0.238 |
| opioid+alcohol use disorder – control | -0.73502 | 0.3420 | -2.1490 | 0.042 |
| Age Cateogry: | | | | |
| 50-64 – 30-49 | 0.06677 | 0.3082 | 0.2167 | 0.830 |
| 50-65 – 30-49 | 0.42943 | 0.4734 | 0.9071 | 0.373 |
| 65+ – 30-49 | 0.33863 | 0.3570 | 0.9485 | 0.352 |
| <30 – 30-49 | -0.04580 | 0.3733 | -0.1227 | 0.903 |
| Ethnicity: | | | | |
| Asian – White | 0.01398 | 0.6215 | 0.0225 | 0.982 |
| Black – White | 0.18163 | 0.2530 | 0.7178 | 0.480 |
| Hispanic – White | 0.35192 | 0.4839 | 0.7272 | 0.474 |
| Gender: | | | | |
| Male – Female | -0.04344 | 0.2991 | -0.1452 | 0.886 |
| PMI converted to hours | 0.00763 | 0.0124 | 0.6147 | 0.545 |
| pH | 0.08434 | 0.4290 | 0.1966 | 0.846 |

[a] Represents reference level

### Linear Regression

Model Fit Measures

| Model | R | $R^2$ |
|---|---|---|
| 1 | 0.759 | 0.577 |

Model Coefficients - WDR48

| Predictor | Estimate | SE | t | p |
|---|---|---|---|---|
| Intercept [a] | 28.3715 | 4.9105 | 5.7778 | < .001 |
| Arm: | | | | |
| alcohol use disorder – control | 0.2245 | 0.5167 | 0.4346 | 0.668 |
| opioid use disorder – control | -0.0954 | 0.4834 | -0.1973 | 0.845 |
| opioid+alcohol use disorder – control | -1.8318 | 0.5716 | -3.2048 | 0.004 |
| Age Cateogry: | | | | |
| 50-64 – 30-49 | -0.6726 | 0.5150 | -1.3060 | 0.204 |
| 50-65 – 30-49 | -1.4898 | 0.7912 | -1.8830 | 0.072 |
| 65+ – 30-49 | -1.0690 | 0.5966 | -1.7917 | 0.086 |
| <30 – 30-49 | 0.3482 | 0.6238 | 0.5581 | 0.582 |
| Ethnicity: | | | | |
| Asian – White | -1.4917 | 1.0387 | -1.4361 | 0.164 |
| Black – White | -0.2907 | 0.4229 | -0.6875 | 0.498 |
| Hispanic – White | -0.0582 | 0.8087 | -0.0720 | 0.943 |
| Gender: | | | | |
| Male – Female | -0.9848 | 0.4998 | -1.9702 | 0.060 |
| PMI converted to hours | 0.0352 | 0.0207 | 1.6971 | 0.103 |
| pH | -0.5865 | 0.7170 | -0.8180 | 0.421 |

[a] Represents reference level

### Linear Regression

Model Fit Measures

| Model | R | $R^2$ |
|---|---|---|
| 1 | 0.666 | 0.443 |

Model Coefficients - PGAM5

| Predictor | Estimate | SE | t | p |
|---|---|---|---|---|
| Intercept [a] | 23.32557 | 4.1472 | 5.6244 | < .001 |
| Arm: | | | | |
| alcohol use disorder – control | -0.76519 | 0.4364 | -1.7535 | 0.092 |
| opioid use disorder – control | -0.39921 | 0.4082 | -0.9779 | 0.338 |
| opioid+alcohol use disorder – control | -1.48937 | 0.4828 | -3.0852 | 0.005 |
| Age Cateogry: | | | | |
| 50-64 – 30-49 | -0.29973 | 0.4349 | -0.6891 | 0.497 |
| 50-65 – 30-49 | -0.88203 | 0.6682 | -1.3200 | 0.199 |
| 65+ – 30-49 | 0.30306 | 0.5039 | 0.6014 | 0.553 |
| <30 – 30-49 | 0.21871 | 0.5268 | 0.4151 | 0.682 |
| Ethnicity: | | | | |
| Asian – White | -0.63022 | 0.8772 | -0.7184 | 0.479 |
| Black – White | -0.04044 | 0.3571 | -0.1132 | 0.911 |
| Hispanic – White | -0.10611 | 0.6830 | -0.1554 | 0.878 |
| Gender: | | | | |
| Male – Female | 0.03847 | 0.4221 | 0.0911 | 0.928 |
| PMI converted to hours | 0.00818 | 0.0175 | 0.4667 | 0.645 |
| pH | 0.15609 | 0.6055 | 0.2578 | 0.799 |

[a] Represents reference level

### Linear Regression

Model Fit Measures

| Model | R | $R^2$ |
|---|---|---|
| 1 | 0.758 | 0.575 |

Model Coefficients - SYNE1

| Predictor | Estimate | SE | t | p |
|---|---|---|---|---|
| Intercept [a] | 10.0969 | 5.5730 | 1.812 | 0.083 |
| Arm: | | | | |
| alcohol use disorder – control | -1.0348 | 0.5864 | -1.765 | 0.090 |
| opioid use disorder – control | -1.4002 | 0.5486 | -2.552 | 0.017 |
| opioid+alcohol use disorder – control | -1.2700 | 0.6487 | -1.958 | 0.062 |
| Age Cateogry: | | | | |
| 50-64 – 30-49 | -0.9584 | 0.5845 | -1.640 | 0.114 |
| 50-65 – 30-49 | -0.3705 | 0.8979 | -0.413 | 0.684 |
| 65+ – 30-49 | -0.7275 | 0.6771 | -1.074 | 0.293 |
| <30 – 30-49 | -1.0980 | 0.7080 | -1.551 | 0.134 |
| Ethnicity: | | | | |
| Asian – White | -1.5851 | 1.1789 | -1.345 | 0.191 |
| Black – White | 1.0455 | 0.4799 | 2.179 | 0.039 |
| Hispanic – White | -0.2933 | 0.9179 | -0.320 | 0.752 |
| Gender: | | | | |
| Male – Female | 0.7910 | 0.5673 | 1.394 | 0.176 |
| PMI converted to hours | 0.0298 | 0.0235 | 1.266 | 0.218 |
| pH | 2.0816 | 0.8137 | 2.558 | 0.017 |

[a] Represents reference level

### Linear Regression

Model Fit Measures

| Model | R | $R^2$ |
|---|---|---|
| 1 | 0.636 | 0.404 |

Model Coefficients - SCAMP4

| Predictor | Estimate | SE | t | p |
|---|---|---|---|---|
| Intercept [a] | 22.33637 | 4.6759 | 4.77696 | < .001 |
| Arm: | | | | |
| alcohol use disorder – control | -0.20182 | 0.4920 | -0.41019 | 0.685 |
| opioid use disorder – control | 0.22027 | 0.4603 | 0.47858 | 0.637 |
| opioid+alcohol use disorder – control | -1.18018 | 0.5443 | -2.16829 | 0.040 |
| Age Cateogry: | | | | |
| 50-64 – 30-49 | -9.76e−4 | 0.4904 | -0.00199 | 0.998 |
| 50-65 – 30-49 | -0.06460 | 0.7534 | -0.08575 | 0.932 |
| 65+ – 30-49 | 0.00717 | 0.5681 | 0.01261 | 0.990 |
| <30 – 30-49 | 0.34620 | 0.5940 | 0.58283 | 0.565 |
| Ethnicity: | | | | |
| Asian – White | -0.51396 | 0.9891 | -0.51964 | 0.608 |
| Black – White | -0.14305 | 0.4027 | -0.35526 | 0.725 |
| Hispanic – White | -0.60471 | 0.7701 | -0.78523 | 0.440 |
| Gender: | | | | |
| Male – Female | -0.57303 | 0.4760 | -1.20395 | 0.240 |
| PMI converted to hours | 0.03367 | 0.0197 | 1.70490 | 0.101 |
| pH | 0.14654 | 0.6827 | 0.21464 | 0.832 |

[a] Represents reference level

### Linear Regression

Model Fit Measures

| Model | R | $R^2$ |
|---|---|---|
| 1 | 0.661 | 0.437 |

Model Coefficients - RAB3C

| Predictor | Estimate | SE | t | p |
|---|---|---|---|---|
| Intercept [a] | 30.11065 | 1.34350 | 22.4122 | < .001 |
| Arm: | | | | |
| alcohol use disorder – control | -0.04026 | 0.14137 | -0.2848 | 0.778 |
| opioid use disorder – control | 0.00233 | 0.13224 | 0.0176 | 0.986 |
| opioid+alcohol use disorder – control | 0.40927 | 0.15639 | 2.6170 | 0.015 |
| Age Cateogry: | | | | |
| 50-64 – 30-49 | -0.10802 | 0.14090 | -0.7666 | 0.451 |
| 50-65 – 30-49 | -0.30491 | 0.21646 | -1.4086 | 0.172 |
| 65+ – 30-49 | -0.17524 | 0.16324 | -1.0735 | 0.294 |
| <30 – 30-49 | -0.13581 | 0.17067 | -0.7957 | 0.434 |
| Ethnicity: | | | | |
| Asian – White | 0.17834 | 0.28419 | 0.6275 | 0.536 |
| Black – White | -0.02791 | 0.11570 | -0.2412 | 0.811 |
| Hispanic – White | -0.09901 | 0.22127 | -0.4474 | 0.659 |
| Gender: | | | | |
| Male – Female | 0.10365 | 0.13676 | 0.7579 | 0.456 |
| PMI converted to hours | 0.00181 | 0.00567 | 0.3194 | 0.752 |
| pH | -0.13643 | 0.19617 | -0.6955 | 0.493 |

[a] Represents reference level

### Linear Regression

Model Fit Measures

| Model | R | $R^2$ |
|---|---|---|
| 1 | 0.620 | 0.385 |

Model Coefficients - SAT2

| Predictor | Estimate | SE | t | p |
|---|---|---|---|---|
| Intercept [a] | 23.66302 | 3.1059 | 7.6187 | < .001 |
| Arm: | | | | |
| alcohol use disorder – control | -0.26482 | 0.3268 | -0.8103 | 0.426 |
| opioid use disorder – control | -0.00365 | 0.3057 | -0.0119 | 0.991 |
| opioid+alcohol use disorder – control | -1.07792 | 0.3615 | -2.9814 | 0.006 |
| Age Cateogry: | | | | |
| 50-64 – 30-49 | -0.02369 | 0.3257 | -0.0727 | 0.943 |
| 50-65 – 30-49 | -0.02300 | 0.5004 | -0.0460 | 0.964 |
| 65+ – 30-49 | 0.07699 | 0.3774 | 0.2040 | 0.840 |
| <30 – 30-49 | -0.12395 | 0.3946 | -0.3141 | 0.756 |
| Ethnicity: | | | | |
| Asian – White | -0.09554 | 0.6570 | -0.1454 | 0.886 |
| Black – White | -0.17870 | 0.2675 | -0.6681 | 0.510 |
| Hispanic – White | -0.15988 | 0.5115 | -0.3125 | 0.757 |
| Gender: | | | | |
| Male – Female | -0.12278 | 0.3162 | -0.3884 | 0.701 |
| PMI converted to hours | 0.01309 | 0.0131 | 0.9977 | 0.328 |
| pH | -0.04007 | 0.4535 | -0.0884 | 0.930 |

[a] Represents reference level

### Linear Regression

Model Fit Measures

| Model | R | $R^2$ |
|---|---|---|
| 1 | 0.649 | 0.422 |

Model Coefficients - AHNAK2

| Predictor | Estimate | SE | t | p |
|---|---|---|---|---|
| Intercept [a] | 21.3391 | 7.6459 | 2.791 | 0.010 |
| Arm: | | | | |
| alcohol use disorder – control | -0.3709 | 0.8045 | -0.461 | 0.649 |
| opioid use disorder – control | -0.2286 | 0.7526 | -0.304 | 0.764 |
| opioid+alcohol use disorder – control | -1.8360 | 0.8900 | -2.063 | 0.050 |
| Age Cateogry: | | | | |
| 50-64 – 30-49 | -0.2322 | 0.8019 | -0.290 | 0.775 |
| 50-65 – 30-49 | 0.3340 | 1.2319 | 0.271 | 0.789 |
| 65+ – 30-49 | 0.6322 | 0.9290 | 0.680 | 0.503 |
| <30 – 30-49 | 0.7687 | 0.9713 | 0.791 | 0.436 |
| Ethnicity: | | | | |
| Asian – White | 1.4271 | 1.6173 | 0.882 | 0.386 |
| Black – White | -0.4340 | 0.6584 | -0.659 | 0.516 |
| Hispanic – White | 2.8077 | 1.2593 | 2.230 | 0.035 |
| Gender: | | | | |
| Male – Female | -0.1239 | 0.7783 | -0.159 | 0.875 |
| PMI converted to hours | -0.0526 | 0.0323 | -1.630 | 0.116 |
| pH | 0.7948 | 1.1164 | 0.712 | 0.483 |

[a] Represents reference level

### Linear Regression

Model Fit Measures

| Model | R | $R^2$ |
|---|---|---|
| 1 | 0.616 | 0.379 |

Model Coefficients - TAB3

| Predictor | Estimate | SE | t | p |
| --- | --- | --- | --- | --- |
| Intercept [a] | 23.05308 | 3.0712 | 7.5062 | < .001 |
| Arm: | | | | |
| alcohol use disorder – control | -0.21565 | 0.3232 | -0.6673 | 0.511 |
| opioid use disorder – control | -0.13855 | 0.3023 | -0.4583 | 0.651 |
| opioid+alcohol use disorder – control | -0.93012 | 0.3575 | -2.6017 | 0.016 |
| Age Cateogry: | | | | |
| 50-64 – 30-49 | 0.27521 | 0.3221 | 0.8544 | 0.401 |
| 50-65 – 30-49 | 0.73348 | 0.4948 | 1.4823 | 0.151 |
| 65+ – 30-49 | 0.29231 | 0.3732 | 0.7833 | 0.441 |
| <30 – 30-49 | 0.42346 | 0.3902 | 1.0854 | 0.289 |
| Ethnicity: | | | | |
| Asian – White | -0.03730 | 0.6496 | -0.0574 | 0.955 |
| Black – White | -0.06456 | 0.2645 | -0.2441 | 0.809 |
| Hispanic – White | -0.24303 | 0.5058 | -0.4805 | 0.635 |
| Gender: | | | | |
| Male – Female | -0.40290 | 0.3126 | -1.2888 | 0.210 |
| PMI converted to hours | 0.00203 | 0.0130 | 0.1567 | 0.877 |
| pH | 0.08964 | 0.4484 | 0.1999 | 0.843 |

[a] Represents reference level

### Linear Regression

Model Fit Measures

| Model | R | $R^2$ |
| --- | --- | --- |
| 1 | 0.565 | 0.319 |

Model Coefficients - SYN2

| Predictor | Estimate | SE | t | p |
|---|---|---|---|---|
| Intercept [a] | 34.10857 | 1.63528 | 20.8579 | < .001 |
| Arm: | | | | |
| alcohol use disorder – control | -0.02102 | 0.17207 | -0.1221 | 0.904 |
| opioid use disorder – control | -0.09326 | 0.16097 | -0.5794 | 0.568 |
| opioid+alcohol use disorder – control | 0.32617 | 0.19035 | 1.7135 | 0.100 |
| Age Cateogry: | | | | |
| 50-64 – 30-49 | -0.13475 | 0.17150 | -0.7857 | 0.440 |
| 50-65 – 30-49 | -0.19880 | 0.26348 | -0.7545 | 0.458 |
| 65+ – 30-49 | -0.10739 | 0.19869 | -0.5405 | 0.594 |
| <30 – 30-49 | 0.04274 | 0.20774 | 0.2057 | 0.839 |
| Ethnicity: | | | | |
| Asian – White | 0.06886 | 0.34591 | 0.1991 | 0.844 |
| Black – White | -0.02782 | 0.14082 | -0.1976 | 0.845 |
| Hispanic – White | 0.01932 | 0.26933 | 0.0717 | 0.943 |
| Gender: | | | | |
| Male – Female | -0.01725 | 0.16646 | -0.1036 | 0.918 |
| PMI converted to hours | -0.00323 | 0.00691 | -0.4682 | 0.644 |
| pH | -0.20421 | 0.23877 | -0.8553 | 0.401 |

[a] Represents reference level

### Linear Regression

Model Fit Measures

| Model | R | $R^2$ |
|---|---|---|
| 1 | 0.722 | 0.522 |

Model Coefficients - CMC1

| Predictor | Estimate | SE | t | p |
|---|---|---|---|---|
| Intercept [a] | 17.60900 | 3.1415 | 5.6052 | < .001 |
| Arm: | | | | |
| alcohol use disorder – control | 0.04124 | 0.3306 | 0.1247 | 0.902 |
| opioid use disorder – control | -0.10180 | 0.3092 | -0.3292 | 0.745 |
| opioid+alcohol use disorder – control | -1.01195 | 0.3657 | -2.7672 | 0.011 |
| Age Cateogry: | | | | |
| 50-64 – 30-49 | -0.22856 | 0.3295 | -0.6937 | 0.495 |
| 50-65 – 30-49 | 0.50512 | 0.5062 | 0.9979 | 0.328 |
| 65+ – 30-49 | -0.20431 | 0.3817 | -0.5352 | 0.597 |
| <30 – 30-49 | -0.30843 | 0.3991 | -0.7728 | 0.447 |
| Ethnicity: | | | | |
| Asian – White | -0.13614 | 0.6645 | -0.2049 | 0.839 |
| Black – White | -0.19928 | 0.2705 | -0.7366 | 0.468 |
| Hispanic – White | -0.35452 | 0.5174 | -0.6852 | 0.500 |
| Gender: | | | | |
| Male – Female | 0.01159 | 0.3198 | 0.0363 | 0.971 |
| PMI converted to hours | 0.00700 | 0.0133 | 0.5275 | 0.603 |
| pH | 0.93128 | 0.4587 | 2.0302 | 0.054 |

[a] Represents reference level

### Linear Regression

Model Fit Measures

| Model | R | $R^2$ |
|---|---|---|
| 1 | 0.672 | 0.452 |

Model Coefficients - CRELD1

| Predictor | Estimate | SE | t | p |
|---|---|---|---|---|
| Intercept [a] | 27.3566 | 3.1720 | 8.6243 | < .001 |
| Arm: | | | | |
| alcohol use disorder – control | 0.0491 | 0.3338 | 0.1471 | 0.884 |
| opioid use disorder – control | -0.0686 | 0.3122 | -0.2197 | 0.828 |
| opioid+alcohol use disorder – control | -0.9541 | 0.3692 | -2.5840 | 0.016 |
| Age Cateogry: | | | | |
| 50-64 – 30-49 | 0.0854 | 0.3327 | 0.2566 | 0.800 |
| 50-65 – 30-49 | 0.2626 | 0.5111 | 0.5138 | 0.612 |
| 65+ – 30-49 | 0.1014 | 0.3854 | 0.2632 | 0.795 |
| <30 – 30-49 | 0.0936 | 0.4030 | 0.2324 | 0.818 |
| Ethnicity: | | | | |
| Asian – White | -0.1273 | 0.6710 | -0.1898 | 0.851 |
| Black – White | 0.3212 | 0.2732 | 1.1758 | 0.251 |
| Hispanic – White | 0.6771 | 0.5224 | 1.2960 | 0.207 |
| Gender: | | | | |
| Male – Female | -0.5160 | 0.3229 | -1.5980 | 0.123 |
| PMI converted to hours | -4.92e−4 | 0.0134 | -0.0367 | 0.971 |
| pH | -0.5213 | 0.4632 | -1.1255 | 0.272 |

[a] Represents reference level

### Linear Regression

Model Fit Measures

| Model | R | $R^2$ |
|---|---|---|
| 1 | 0.619 | 0.383 |

Model Coefficients - TTC5

| Predictor | Estimate | SE | t | p |
|---|---|---|---|---|
| Intercept [a] | 25.30689 | 3.8076 | 6.646 | < .001 |
| Arm: | | | | |
| alcohol use disorder – control | -0.43000 | 0.4007 | -1.073 | 0.294 |
| opioid use disorder – control | 0.04769 | 0.3748 | 0.127 | 0.900 |
| opioid+alcohol use disorder – control | -0.76016 | 0.4432 | -1.715 | 0.099 |
| Age Cateogry: | | | | |
| 50-64 – 30-49 | 0.50395 | 0.3993 | 1.262 | 0.219 |
| 50-65 – 30-49 | 0.41667 | 0.6135 | 0.679 | 0.504 |
| 65+ – 30-49 | 0.44023 | 0.4626 | 0.952 | 0.351 |
| <30 – 30-49 | 0.27743 | 0.4837 | 0.574 | 0.572 |
| Ethnicity: | | | | |
| Asian – White | -0.76207 | 0.8054 | -0.946 | 0.353 |
| Black – White | 0.42706 | 0.3279 | 1.302 | 0.205 |
| Hispanic – White | -0.28855 | 0.6271 | -0.460 | 0.650 |
| Gender: | | | | |
| Male – Female | 0.19814 | 0.3876 | 0.511 | 0.614 |
| PMI converted to hours | 0.00448 | 0.0161 | 0.278 | 0.783 |
| pH | -0.35794 | 0.5560 | -0.644 | 0.526 |

[a] Represents reference level

### Linear Regression

Model Fit Measures

| Model | R | $R^2$ |
|---|---|---|
| 1 | 0.738 | 0.545 |

Model Coefficients - LRRC4B

| Predictor | Estimate | SE | t | p |
|---|---|---|---|---|
| Intercept [a] | 27.37189 | 2.7558 | 9.9323 | < .001 |
| Arm: | | | | |
| alcohol use disorder – control | -0.27522 | 0.2900 | -0.9491 | 0.352 |
| opioid use disorder – control | 0.50522 | 0.2713 | 1.8624 | 0.075 |
| opioid+alcohol use disorder – control | -0.69321 | 0.3208 | -2.1609 | 0.041 |
| Age Cateogry: | | | | |
| 50-64 – 30-49 | -0.10034 | 0.2890 | -0.3471 | 0.732 |
| 50-65 – 30-49 | -0.21933 | 0.4440 | -0.4940 | 0.626 |
| 65+ – 30-49 | 0.00479 | 0.3348 | 0.0143 | 0.989 |
| <30 – 30-49 | -0.51770 | 0.3501 | -1.4788 | 0.152 |
| Ethnicity: | | | | |
| Asian – White | -0.34919 | 0.5829 | -0.5990 | 0.555 |
| Black – White | -0.03507 | 0.2373 | -0.1478 | 0.884 |
| Hispanic – White | -0.69774 | 0.4539 | -1.5373 | 0.137 |
| Gender: | | | | |
| Male – Female | 0.29554 | 0.2805 | 1.0536 | 0.303 |
| PMI converted to hours | 0.01255 | 0.0116 | 1.0780 | 0.292 |
| pH | -0.64164 | 0.4024 | -1.5946 | 0.124 |

[a] Represents reference level

### Linear Regression

Model Fit Measures

| Model | R | $R^2$ |
|---|---|---|
| 1 | 0.639 | 0.408 |

Model Coefficients - TCP11L1

| Predictor | Estimate | SE | t | p |
|---|---|---|---|---|
| Intercept [a] | 24.47977 | 4.4906 | 5.45140 | < .001 |
| Arm: | | | | |
| alcohol use disorder – control | -0.70447 | 0.4725 | -1.49090 | 0.149 |
| opioid use disorder – control | -0.19879 | 0.4420 | -0.44973 | 0.657 |
| opioid+alcohol use disorder – control | -1.62245 | 0.5227 | -3.10386 | 0.005 |
| Age Cateogry: | | | | |
| 50-64 – 30-49 | 0.30264 | 0.4710 | 0.64260 | 0.527 |
| 50-65 – 30-49 | -0.10296 | 0.7235 | -0.14230 | 0.888 |
| 65+ – 30-49 | -0.06060 | 0.5456 | -0.11106 | 0.912 |
| <30 – 30-49 | -0.04245 | 0.5705 | -0.07441 | 0.941 |
| Ethnicity: | | | | |
| Asian – White | -0.68166 | 0.9499 | -0.71763 | 0.480 |
| Black – White | -0.60714 | 0.3867 | -1.57003 | 0.130 |
| Hispanic – White | -0.54686 | 0.7396 | -0.73941 | 0.467 |
| Gender: | | | | |
| Male – Female | -0.47058 | 0.4571 | -1.02950 | 0.314 |
| PMI converted to hours | 0.00422 | 0.0190 | 0.22226 | 0.826 |
| pH | 0.00308 | 0.6557 | 0.00470 | 0.996 |

[a] Represents reference level

### Linear Regression

Model Fit Measures

| Model | R | $R^2$ |
|---|---|---|
| 1 | 0.694 | 0.482 |

Model Coefficients - FXYD6

| Predictor | Estimate | SE | t | p |
|---|---|---|---|---|
| Intercept [a] | 27.63883 | 1.49828 | 18.4470 | < .001 |
| Arm: | | | | |
| alcohol use disorder – control | -0.02041 | 0.15766 | -0.1295 | 0.898 |
| opioid use disorder – control | -0.00650 | 0.14748 | -0.0440 | 0.965 |
| opioid+alcohol use disorder – control | -0.50089 | 0.17441 | -2.8719 | 0.008 |
| Age Cateogry: | | | | |
| 50-64 – 30-49 | -0.22433 | 0.15714 | -1.4276 | 0.166 |
| 50-65 – 30-49 | 0.00360 | 0.24140 | 0.0149 | 0.988 |
| 65+ – 30-49 | -0.15161 | 0.18205 | -0.8328 | 0.413 |
| <30 – 30-49 | -0.00989 | 0.19033 | -0.0520 | 0.959 |
| Ethnicity: | | | | |
| Asian – White | -0.23653 | 0.31693 | -0.7463 | 0.463 |
| Black – White | -0.17230 | 0.12902 | -1.3354 | 0.194 |
| Hispanic – White | 0.05997 | 0.24677 | 0.2430 | 0.810 |
| Gender: | | | | |
| Male – Female | -0.10895 | 0.15251 | -0.7144 | 0.482 |
| PMI converted to hours | 0.00758 | 0.00633 | 1.1972 | 0.243 |
| pH | 0.16049 | 0.21877 | 0.7336 | 0.470 |

[a] Represents reference level

### Linear Regression

Model Fit Measures

| Model | R | $R^2$ |
|---|---|---|
| 1 | 0.727 | 0.529 |

Model Coefficients - NDUFAF4

| Predictor | Estimate | SE | t | p |
|---|---|---|---|---|
| Intercept [a] | 21.1415 | 3.6080 | 5.860 | < .001 |
| Arm: | | | | |
| alcohol use disorder – control | -0.4740 | 0.3797 | -1.248 | 0.224 |
| opioid use disorder – control | -0.6286 | 0.3551 | -1.770 | 0.089 |
| opioid+alcohol use disorder – control | -1.0546 | 0.4200 | -2.511 | 0.019 |
| Age Cateogry: | | | | |
| 50-64 – 30-49 | -0.5972 | 0.3784 | -1.578 | 0.128 |
| 50-65 – 30-49 | -0.0707 | 0.5813 | -0.122 | 0.904 |
| 65+ – 30-49 | -0.6697 | 0.4384 | -1.528 | 0.140 |
| <30 – 30-49 | -0.2598 | 0.4583 | -0.567 | 0.576 |
| Ethnicity: | | | | |
| Asian – White | -1.1307 | 0.7632 | -1.482 | 0.151 |
| Black – White | -0.1345 | 0.3107 | -0.433 | 0.669 |
| Hispanic – White | 1.5191 | 0.5942 | 2.556 | 0.017 |
| Gender: | | | | |
| Male – Female | 0.1303 | 0.3673 | 0.355 | 0.726 |
| PMI converted to hours | -0.0234 | 0.0152 | -1.536 | 0.138 |
| pH | 0.6132 | 0.5268 | 1.164 | 0.256 |

[a] Represents reference level

### Linear Regression

Model Fit Measures

| Model | R | $R^2$ |
|---|---|---|
| 1 | 0.758 | 0.574 |

Model Coefficients - SDHC

| Predictor | Estimate | SE | t | p |
|---|---|---|---|---|
| Intercept [a] | 30.5328 | 6.0959 | 5.0088 | < .001 |
| Arm: | | | | |
| alcohol use disorder – control | 0.1088 | 0.6414 | 0.1697 | 0.867 |
| opioid use disorder – control | -0.0929 | 0.6000 | -0.1548 | 0.878 |
| opioid+alcohol use disorder – control | 2.4355 | 0.7096 | 3.4323 | 0.002 |
| Age Cateogry: | | | | |
| 50-64 – 30-49 | -0.4060 | 0.6393 | -0.6351 | 0.531 |
| 50-65 – 30-49 | -0.1622 | 0.9822 | -0.1652 | 0.870 |
| 65+ – 30-49 | -0.7983 | 0.7407 | -1.0779 | 0.292 |
| <30 – 30-49 | -0.6739 | 0.7744 | -0.8703 | 0.393 |
| Ethnicity: | | | | |
| Asian – White | -1.7493 | 1.2894 | -1.3566 | 0.188 |
| Black – White | -0.1674 | 0.5249 | -0.3189 | 0.753 |
| Hispanic – White | 0.0456 | 1.0040 | 0.0454 | 0.964 |
| Gender: | | | | |
| Male – Female | 0.1200 | 0.6205 | 0.1934 | 0.848 |
| PMI converted to hours | 0.0246 | 0.0257 | 0.9557 | 0.349 |
| pH | -1.0431 | 0.8901 | -1.1719 | 0.253 |

[a] Represents reference level

### Linear Regression

Model Fit Measures

| Model | R | $R^2$ |
|---|---|---|
| 1 | 0.727 | 0.529 |

Model Coefficients - GDE1

| Predictor | Estimate | SE | t | p |
|---|---|---|---|---|
| Intercept [a] | 19.92895 | 2.5264 | 7.8883 | < .001 |
| Arm: | | | | |
| alcohol use disorder – control | -0.44159 | 0.2658 | -1.6611 | 0.110 |
| opioid use disorder – control | -0.17195 | 0.2487 | -0.6914 | 0.496 |
| opioid+alcohol use disorder – control | -0.94960 | 0.2941 | -3.2290 | 0.004 |
| Age Cateogry: | | | | |
| 50-64 – 30-49 | -0.43639 | 0.2650 | -1.6470 | 0.113 |
| 50-65 – 30-49 | -0.06729 | 0.4071 | -0.1653 | 0.870 |
| 65+ – 30-49 | -0.29959 | 0.3070 | -0.9760 | 0.339 |
| <30 – 30-49 | -0.17579 | 0.3209 | -0.5477 | 0.589 |
| Ethnicity: | | | | |
| Asian – White | -0.64482 | 0.5344 | -1.2066 | 0.239 |
| Black – White | -0.00808 | 0.2176 | -0.0371 | 0.971 |
| Hispanic – White | 0.00587 | 0.4161 | 0.0141 | 0.989 |
| Gender: | | | | |
| Male – Female | -0.17911 | 0.2572 | -0.6965 | 0.493 |
| PMI converted to hours | 0.01199 | 0.0107 | 1.1237 | 0.272 |
| pH | 0.58582 | 0.3689 | 1.5881 | 0.125 |

[a] Represents reference level

### Linear Regression

Model Fit Measures

| Model | R | $R^2$ |
|---|---|---|
| 1 | 0.677 | 0.458 |

Model Coefficients - OPA3

| Predictor | Estimate | SE | t | p |
|---|---|---|---|---|
| Intercept [a] | 22.17657 | 3.0082 | 7.372 | < .001 |
| Arm: | | | | |
| alcohol use disorder – control | 0.31970 | 0.3165 | 1.010 | 0.323 |
| opioid use disorder – control | -0.25692 | 0.2961 | -0.868 | 0.394 |
| opioid+alcohol use disorder – control | -0.88974 | 0.3502 | -2.541 | 0.018 |
| Age Cateogry: | | | | |
| 50-64 – 30-49 | -0.04556 | 0.3155 | -0.144 | 0.886 |
| 50-65 – 30-49 | -0.28280 | 0.4847 | -0.583 | 0.565 |
| 65+ – 30-49 | -0.49525 | 0.3655 | -1.355 | 0.188 |
| <30 – 30-49 | -0.04404 | 0.3821 | -0.115 | 0.909 |
| Ethnicity: | | | | |
| Asian – White | -0.57644 | 0.6363 | -0.906 | 0.374 |
| Black – White | -0.18024 | 0.2590 | -0.696 | 0.493 |
| Hispanic – White | 0.54372 | 0.4954 | 1.097 | 0.283 |
| Gender: | | | | |
| Male – Female | -0.25363 | 0.3062 | -0.828 | 0.416 |
| PMI converted to hours | -0.00690 | 0.0127 | -0.543 | 0.592 |
| pH | 0.29616 | 0.4392 | 0.674 | 0.507 |

[a] Represents reference level

### Linear Regression

Model Fit Measures

| Model | R | $R^2$ |
|---|---|---|
| 1 | 0.672 | 0.452 |

Model Coefficients - TRNT1

| Predictor | Estimate | SE | t | p |
|---|---|---|---|---|
| Intercept [a] | 25.35140 | 2.8546 | 8.8808 | < .001 |
| Arm: | | | | |
| alcohol use disorder – control | -0.09199 | 0.3004 | -0.3063 | 0.762 |
| opioid use disorder – control | 0.37679 | 0.2810 | 1.3409 | 0.192 |
| opioid+alcohol use disorder – control | -0.69121 | 0.3323 | -2.0801 | 0.048 |
| Age Cateogry: | | | | |
| 50-64 – 30-49 | 0.26106 | 0.2994 | 0.8720 | 0.392 |
| 50-65 – 30-49 | -0.07657 | 0.4599 | -0.1665 | 0.869 |
| 65+ – 30-49 | 0.13385 | 0.3468 | 0.3859 | 0.703 |
| <30 – 30-49 | -0.00667 | 0.3626 | -0.0184 | 0.985 |
| Ethnicity: | | | | |
| Asian – White | -0.20347 | 0.6038 | -0.3370 | 0.739 |
| Black – White | -0.13562 | 0.2458 | -0.5517 | 0.586 |
| Hispanic – White | -1.21309 | 0.4702 | -2.5802 | 0.016 |
| Gender: | | | | |
| Male – Female | 0.09868 | 0.2906 | 0.3396 | 0.737 |
| PMI converted to hours | 0.02093 | 0.0121 | 1.7360 | 0.095 |
| pH | -0.39900 | 0.4168 | -0.9573 | 0.348 |

[a] Represents reference level

### Linear Regression

Model Fit Measures

| Model | R | $R^2$ |
|---|---|---|
| 1 | 0.626 | 0.392 |

Model Coefficients - MRPL13

| Predictor | Estimate | SE | t | p |
|---|---|---|---|---|
| Intercept [a] | 23.81960 | 3.2239 | 7.3885 | < .001 |
| Arm: | | | | |
| alcohol use disorder – control | -0.46305 | 0.3392 | -1.3650 | 0.185 |
| opioid use disorder – control | -0.24833 | 0.3173 | -0.7826 | 0.442 |
| opioid+alcohol use disorder – control | -0.94453 | 0.3753 | -2.5169 | 0.019 |
| Age Cateogry: | | | | |
| 50-64 – 30-49 | 0.20866 | 0.3381 | 0.6171 | 0.543 |
| 50-65 – 30-49 | -0.03593 | 0.5194 | -0.0692 | 0.945 |
| 65+ – 30-49 | 0.58022 | 0.3917 | 1.4812 | 0.152 |
| <30 – 30-49 | 0.22790 | 0.4095 | 0.5565 | 0.583 |
| Ethnicity: | | | | |
| Asian – White | 0.31583 | 0.6819 | 0.4631 | 0.647 |
| Black – White | 0.29802 | 0.2776 | 1.0735 | 0.294 |
| Hispanic – White | 0.07507 | 0.5310 | 0.1414 | 0.889 |
| Gender: | | | | |
| Male – Female | -0.06047 | 0.3282 | -0.1843 | 0.855 |
| PMI converted to hours | -0.00746 | 0.0136 | -0.5481 | 0.589 |
| pH | -5.08e−5 | 0.4707 | -1.08e−4 | 1.000 |

[a] Represents reference level

### Linear Regression

Model Fit Measures

| Model | R | $R^2$ |
|---|---|---|
| 1 | 0.676 | 0.457 |

Model Coefficients - MRPL4

| Predictor | Estimate | SE | t | p |
| --- | --- | --- | --- | --- |
| Intercept [a] | 20.43235 | 2.8442 | 7.18387 | < .001 |
| Arm: | | | | |
| alcohol use disorder – control | -0.53488 | 0.2993 | -1.78724 | 0.087 |
| opioid use disorder – control | -0.04179 | 0.2800 | -0.14925 | 0.883 |
| opioid+alcohol use disorder – control | -0.78963 | 0.3311 | -2.38503 | 0.025 |
| Age Cateogry: | | | | |
| 50-64 – 30-49 | 0.19513 | 0.2983 | 0.65416 | 0.519 |
| 50-65 – 30-49 | 0.81045 | 0.4583 | 1.76854 | 0.090 |
| 65+ – 30-49 | 0.12249 | 0.3456 | 0.35444 | 0.726 |
| <30 – 30-49 | -0.00220 | 0.3613 | -0.00608 | 0.995 |
| Ethnicity: | | | | |
| Asian – White | -0.55938 | 0.6016 | -0.92978 | 0.362 |
| Black – White | -0.00404 | 0.2449 | -0.01651 | 0.987 |
| Hispanic – White | 0.05537 | 0.4684 | 0.11820 | 0.907 |
| Gender: | | | | |
| Male – Female | 0.19274 | 0.2895 | 0.66573 | 0.512 |
| PMI converted to hours | -0.00115 | 0.0120 | -0.09603 | 0.924 |
| pH | 0.46608 | 0.4153 | 1.12230 | 0.273 |

[a] Represents reference level

### Linear Regression

Model Fit Measures

| Model | R | $R^2$ |
| --- | --- | --- |
| 1 | 0.680 | 0.463 |

Model Coefficients - LMTK3

| Predictor | Estimate | SE | t | p |
|---|---|---|---|---|
| Intercept [a] | 20.7620 | 2.5417 | 8.1687 | < .001 |
| Arm: | | | | |
| alcohol use disorder – control | -0.2705 | 0.2674 | -1.0114 | 0.322 |
| opioid use disorder – control | 0.1531 | 0.2502 | 0.6118 | 0.546 |
| opioid+alcohol use disorder – control | -0.6834 | 0.2959 | -2.3098 | 0.030 |
| Age Cateogry: | | | | |
| 50-64 – 30-49 | 0.2724 | 0.2666 | 1.0218 | 0.317 |
| 50-65 – 30-49 | 0.0584 | 0.4095 | 0.1426 | 0.888 |
| 65+ – 30-49 | 0.3663 | 0.3088 | 1.1861 | 0.247 |
| <30 – 30-49 | -0.0144 | 0.3229 | -0.0445 | 0.965 |
| Ethnicity: | | | | |
| Asian – White | 0.0975 | 0.5376 | 0.1814 | 0.858 |
| Black – White | -0.0474 | 0.2189 | -0.2166 | 0.830 |
| Hispanic – White | 0.0120 | 0.4186 | 0.0286 | 0.977 |
| Gender: | | | | |
| Male – Female | -0.0907 | 0.2587 | -0.3504 | 0.729 |
| PMI converted to hours | 0.0163 | 0.0107 | 1.5175 | 0.142 |
| pH | 0.3500 | 0.3711 | 0.9431 | 0.355 |

[a] Represents reference level

### Linear Regression

Model Fit Measures

| Model | R | $R^2$ |
|---|---|---|
| 1 | 0.550 | 0.303 |

Model Coefficients - CPNE1

| Predictor | Estimate | SE | t | p |
|---|---|---|---|---|
| Intercept [a] | 26.3878 | 4.6919 | 5.6242 | < .001 |
| Arm: | | | | |
| alcohol use disorder – control | -0.3147 | 0.4937 | -0.6375 | 0.530 |
| opioid use disorder – control | -0.4780 | 0.4618 | -1.0351 | 0.311 |
| opioid+alcohol use disorder – control | -0.8834 | 0.5462 | -1.6174 | 0.119 |
| Age Cateogry: | | | | |
| 50-64 – 30-49 | 0.1347 | 0.4921 | 0.2738 | 0.787 |
| 50-65 – 30-49 | -0.0140 | 0.7560 | -0.0186 | 0.985 |
| 65+ – 30-49 | -1.1209 | 0.5701 | -1.9662 | 0.061 |
| <30 – 30-49 | 0.4301 | 0.5960 | 0.7216 | 0.478 |
| Ethnicity: | | | | |
| Asian – White | -0.0350 | 0.9925 | -0.0353 | 0.972 |
| Black – White | -0.2109 | 0.4040 | -0.5219 | 0.606 |
| Hispanic – White | -0.3809 | 0.7727 | -0.4929 | 0.627 |
| Gender: | | | | |
| Male – Female | -0.3283 | 0.4776 | -0.6874 | 0.498 |
| PMI converted to hours | 0.0161 | 0.0198 | 0.8129 | 0.424 |
| pH | -0.1126 | 0.6851 | -0.1644 | 0.871 |

[a] Represents reference level

### Linear Regression

Model Fit Measures

| Model | R | $R^2$ |
|---|---|---|
| 1 | 0.594 | 0.353 |

Model Coefficients - DERL1

| Predictor | Estimate | SE | t | p |
|---|---|---|---|---|
| Intercept [a] | 22.6555 | 3.0124 | 7.521 | < .001 |
| Arm: | | | | |
| alcohol use disorder – control | -0.3791 | 0.3170 | -1.196 | 0.243 |
| opioid use disorder – control | 0.1144 | 0.2965 | 0.386 | 0.703 |
| opioid+alcohol use disorder – control | -0.6883 | 0.3507 | -1.963 | 0.061 |
| Age Cateogry: | | | | |
| 50-64 – 30-49 | -0.1094 | 0.3159 | -0.346 | 0.732 |
| 50-65 – 30-49 | 0.1694 | 0.4854 | 0.349 | 0.730 |
| 65+ – 30-49 | 0.1854 | 0.3660 | 0.507 | 0.617 |
| <30 – 30-49 | 0.0801 | 0.3827 | 0.209 | 0.836 |
| Ethnicity: | | | | |
| Asian – White | -0.4244 | 0.6372 | -0.666 | 0.512 |
| Black – White | -0.1188 | 0.2594 | -0.458 | 0.651 |
| Hispanic – White | -0.4909 | 0.4961 | -0.989 | 0.332 |
| Gender: | | | | |
| Male – Female | -0.0376 | 0.3066 | -0.123 | 0.903 |
| PMI converted to hours | 0.0185 | 0.0127 | 1.451 | 0.160 |
| pH | 0.0635 | 0.4398 | 0.144 | 0.886 |

[a] Represents reference level

### Linear Regression

Model Fit Measures

| Model | R | $R^2$ |
|---|---|---|
| 1 | 0.555 | 0.308 |

Model Coefficients - CSTF2T

| Predictor | Estimate | SE | t | p |
|---|---|---|---|---|
| Intercept [a] | 25.39043 | 3.5269 | 7.1991 | < .001 |
| Arm: | | | | |
| alcohol use disorder – control | -0.54195 | 0.3711 | -1.4603 | 0.157 |
| opioid use disorder – control | -0.32457 | 0.3472 | -0.9349 | 0.359 |
| opioid+alcohol use disorder – control | -1.10741 | 0.4105 | -2.6974 | 0.013 |
| Age Cateogry: | | | | |
| 50-64 – 30-49 | 0.05488 | 0.3699 | 0.1484 | 0.883 |
| 50-65 – 30-49 | -0.10917 | 0.5682 | -0.1921 | 0.849 |
| 65+ – 30-49 | 0.40101 | 0.4285 | 0.9358 | 0.359 |
| <30 – 30-49 | 0.16072 | 0.4480 | 0.3587 | 0.723 |
| Ethnicity: | | | | |
| Asian – White | -0.56213 | 0.7460 | -0.7535 | 0.458 |
| Black – White | -0.26289 | 0.3037 | -0.8656 | 0.395 |
| Hispanic – White | 0.08433 | 0.5809 | 0.1452 | 0.886 |
| Gender: | | | | |
| Male – Female | -0.26883 | 0.3590 | -0.7488 | 0.461 |
| PMI converted to hours | -0.00135 | 0.0149 | -0.0905 | 0.929 |
| pH | -0.19973 | 0.5150 | -0.3879 | 0.702 |

[a] Represents reference level

### Linear Regression

Model Fit Measures

| Model | R | $R^2$ |
|---|---|---|
| 1 | 0.783 | 0.613 |

Model Coefficients - FBXO44

| Predictor | Estimate | SE | t | p |
|---|---|---|---|---|
| Intercept [a] | 30.1833 | 3.6010 | 8.382 | < .001 |
| Arm: | | | | |
| alcohol use disorder – control | 0.3918 | 0.3789 | 1.034 | 0.311 |
| opioid use disorder – control | 0.0705 | 0.3545 | 0.199 | 0.844 |
| opioid+alcohol use disorder – control | 1.0768 | 0.4192 | 2.569 | 0.017 |
| Age Cateogry: | | | | |
| 50-64 – 30-49 | 0.2530 | 0.3777 | 0.670 | 0.509 |
| 50-65 – 30-49 | 0.4867 | 0.5802 | 0.839 | 0.410 |
| 65+ – 30-49 | 0.2856 | 0.4375 | 0.653 | 0.520 |
| <30 – 30-49 | -0.3173 | 0.4575 | -0.694 | 0.495 |
| Ethnicity: | | | | |
| Asian – White | -0.2449 | 0.7617 | -0.321 | 0.751 |
| Black – White | 0.7480 | 0.3101 | 2.412 | 0.024 |
| Hispanic – White | -0.9750 | 0.5931 | -1.644 | 0.113 |
| Gender: | | | | |
| Male – Female | -0.6009 | 0.3666 | -1.639 | 0.114 |
| PMI converted to hours | 0.0237 | 0.0152 | 1.556 | 0.133 |
| pH | -1.0778 | 0.5258 | -2.050 | 0.051 |

[a] Represents reference level

### Linear Regression

Model Fit Measures

| Model | R | $R^2$ |
|---|---|---|
| 1 | 0.683 | 0.466 |

Model Coefficients - NAPG

| Predictor | Estimate | SE | t | p |
|---|---|---|---|---|
| Intercept [a] | 28.3667 | 1.02856 | 27.57898 | < .001 |
| Arm: | | | | |
| alcohol use disorder – control | -0.1267 | 0.10823 | -1.17023 | 0.253 |
| opioid use disorder – control | -3.77e−4 | 0.10124 | -0.00372 | 0.997 |
| opioid+alcohol use disorder – control | -0.2804 | 0.11973 | -2.34166 | 0.028 |
| Age Cateogry: | | | | |
| 50-64 – 30-49 | -0.1241 | 0.10787 | -1.14997 | 0.261 |
| 50-65 – 30-49 | -0.0872 | 0.16572 | -0.52607 | 0.604 |
| 65+ – 30-49 | -0.0340 | 0.12497 | -0.27196 | 0.788 |
| <30 – 30-49 | -0.0835 | 0.13066 | -0.63896 | 0.529 |
| Ethnicity: | | | | |
| Asian – White | -0.2017 | 0.21757 | -0.92704 | 0.363 |
| Black – White | -0.1444 | 0.08857 | -1.63065 | 0.116 |
| Hispanic – White | -0.1859 | 0.16940 | -1.09755 | 0.283 |
| Gender: | | | | |
| Male – Female | 0.0224 | 0.10470 | 0.21393 | 0.832 |
| PMI converted to hours | 0.0119 | 0.00434 | 2.74495 | 0.011 |
| pH | 0.1612 | 0.15018 | 1.07353 | 0.294 |

[a] Represents reference level

### Linear Regression

Model Fit Measures

| Model | R | $R^2$ |
|---|---|---|
| 1 | 0.741 | 0.549 |

Model Coefficients - LSM5

| Predictor | Estimate | SE | t | p |
|---|---|---|---|---|
| Intercept [a] | 23.52504 | 2.08564 | 11.2795 | < .001 |
| Arm: | | | | |
| alcohol use disorder – control | -0.24537 | 0.21946 | -1.1180 | 0.275 |
| opioid use disorder – control | 0.25654 | 0.20530 | 1.2496 | 0.223 |
| opioid+alcohol use disorder – control | -0.76746 | 0.24278 | -3.1612 | 0.004 |
| Age Cateogry: | | | | |
| 50-64 – 30-49 | 0.02534 | 0.21874 | 0.1159 | 0.909 |
| 50-65 – 30-49 | 0.29623 | 0.33604 | 0.8815 | 0.387 |
| 65+ – 30-49 | 0.18828 | 0.25341 | 0.7430 | 0.465 |
| <30 – 30-49 | -0.13180 | 0.26495 | -0.4975 | 0.623 |
| Ethnicity: | | | | |
| Asian – White | -0.27503 | 0.44117 | -0.6234 | 0.539 |
| Black – White | 0.00683 | 0.17961 | 0.0380 | 0.970 |
| Hispanic – White | -0.34869 | 0.34350 | -1.0151 | 0.320 |
| Gender: | | | | |
| Male – Female | 0.02947 | 0.21230 | 0.1388 | 0.891 |
| PMI converted to hours | 0.00421 | 0.00881 | 0.4784 | 0.637 |
| pH | -0.02179 | 0.30453 | -0.0716 | 0.944 |

[a] Represents reference level

### Linear Regression

Model Fit Measures

| Model | R | $R^2$ |
|---|---|---|
| 1 | 0.508 | 0.258 |

Model Coefficients - CYB5R1

| Predictor | Estimate | SE | t | p |
|---|---|---|---|---|
| Intercept [a] | 30.8986 | 5.4568 | 5.6624 | < .001 |
| Arm: | | | | |
| alcohol use disorder – control | 0.4515 | 0.5742 | 0.7864 | 0.439 |
| opioid use disorder – control | -0.1816 | 0.5371 | -0.3381 | 0.738 |
| opioid+alcohol use disorder – control | 0.9635 | 0.6352 | 1.5168 | 0.142 |
| Age Cateogry: | | | | |
| 50-64 – 30-49 | -0.0810 | 0.5723 | -0.1415 | 0.889 |
| 50-65 – 30-49 | -0.2471 | 0.8792 | -0.2811 | 0.781 |
| 65+ – 30-49 | -0.6278 | 0.6630 | -0.9469 | 0.353 |
| <30 – 30-49 | 0.0916 | 0.6932 | 0.1321 | 0.896 |
| Ethnicity: | | | | |
| Asian – White | 0.2807 | 1.1543 | 0.2432 | 0.810 |
| Black – White | 0.4624 | 0.4699 | 0.9841 | 0.335 |
| Hispanic – White | 0.5173 | 0.8987 | 0.5756 | 0.570 |
| Gender: | | | | |
| Male – Female | 0.0327 | 0.5554 | 0.0589 | 0.954 |
| PMI converted to hours | -0.0191 | 0.0230 | -0.8297 | 0.415 |
| pH | -0.5709 | 0.7968 | -0.7165 | 0.481 |

[a] Represents reference level

### Linear Regression

Model Fit Measures

| Model | R | $R^2$ |
|---|---|---|
| 1 | 0.618 | 0.381 |

Model Coefficients - KIAA1107

| Predictor | Estimate | SE | t | p |
|---|---|---|---|---|
| Intercept [a] | 24.16025 | 3.0921 | 7.814 | < .001 |
| Arm: | | | | |
| alcohol use disorder – control | -0.35026 | 0.3254 | -1.077 | 0.292 |
| opioid use disorder – control | 0.11832 | 0.3044 | 0.389 | 0.701 |
| opioid+alcohol use disorder – control | 0.57529 | 0.3599 | 1.598 | 0.123 |
| Age Cateogry: | | | | |
| 50-64 – 30-49 | 0.36610 | 0.3243 | 1.129 | 0.270 |
| 50-65 – 30-49 | 0.73842 | 0.4982 | 1.482 | 0.151 |
| 65+ – 30-49 | 0.30255 | 0.3757 | 0.805 | 0.429 |
| <30 – 30-49 | 0.04638 | 0.3928 | 0.118 | 0.907 |
| Ethnicity: | | | | |
| Asian – White | 0.46201 | 0.6541 | 0.706 | 0.487 |
| Black – White | 0.05724 | 0.2663 | 0.215 | 0.832 |
| Hispanic – White | 0.21603 | 0.5093 | 0.424 | 0.675 |
| Gender: | | | | |
| Male – Female | -0.08072 | 0.3147 | -0.256 | 0.800 |
| PMI converted to hours | -0.00501 | 0.0131 | -0.384 | 0.704 |
| pH | 0.17753 | 0.4515 | 0.393 | 0.698 |

[a] Represents reference level

### Linear Regression

Model Fit Measures

| Model | R | $R^2$ |
|---|---|---|
| 1 | 0.762 | 0.581 |

Model Coefficients - WDR37

| Predictor | Estimate | SE | t | p |
|---|---|---|---|---|
| Intercept [a] | 28.5977 | 3.9214 | 7.2927 | < .001 |
| Arm: | | | | |
| alcohol use disorder – control | 0.0226 | 0.4126 | 0.0547 | 0.957 |
| opioid use disorder – control | -0.1820 | 0.3860 | -0.4715 | 0.642 |
| opioid+alcohol use disorder – control | 0.7963 | 0.4565 | 1.7444 | 0.094 |
| Age Cateogry: | | | | |
| 50-64 – 30-49 | 0.7347 | 0.4113 | 1.7863 | 0.087 |
| 50-65 – 30-49 | 0.3908 | 0.6318 | 0.6186 | 0.542 |
| 65+ – 30-49 | 0.3791 | 0.4765 | 0.7956 | 0.434 |
| <30 – 30-49 | -0.0247 | 0.4982 | -0.0495 | 0.961 |
| Ethnicity: | | | | |
| Asian – White | 0.3694 | 0.8295 | 0.4453 | 0.660 |
| Black – White | -0.5465 | 0.3377 | -1.6182 | 0.119 |
| Hispanic – White | 0.8033 | 0.6459 | 1.2438 | 0.226 |
| Gender: | | | | |
| Male – Female | -0.3032 | 0.3992 | -0.7596 | 0.455 |
| PMI converted to hours | -0.0233 | 0.0166 | -1.4041 | 0.173 |
| pH | -0.5340 | 0.5726 | -0.9327 | 0.360 |

[a] Represents reference level

### Linear Regression

Model Fit Measures

| Model | R | $R^2$ |
|---|---|---|
| 1 | 0.656 | 0.430 |

Model Coefficients - HSPB8

| Predictor | Estimate | SE | t | p |
|---|---|---|---|---|
| Intercept [a] | 24.3709 | 2.9704 | 8.205 | < .001 |
| Arm: | | | | |
| alcohol use disorder – control | -0.1269 | 0.3126 | -0.406 | 0.688 |
| opioid use disorder – control | 0.3155 | 0.2924 | 1.079 | 0.291 |
| opioid+alcohol use disorder – control | -0.8097 | 0.3458 | -2.342 | 0.028 |
| Age Cateogry: | | | | |
| 50-64 – 30-49 | 0.0726 | 0.3115 | 0.233 | 0.818 |
| 50-65 – 30-49 | 0.3743 | 0.4786 | 0.782 | 0.442 |
| 65+ – 30-49 | 0.6277 | 0.3609 | 1.739 | 0.095 |
| <30 – 30-49 | -0.0406 | 0.3773 | -0.108 | 0.915 |
| Ethnicity: | | | | |
| Asian – White | 0.5892 | 0.6283 | 0.938 | 0.358 |
| Black – White | -0.0990 | 0.2558 | -0.387 | 0.702 |
| Hispanic – White | -0.2885 | 0.4892 | -0.590 | 0.561 |
| Gender: | | | | |
| Male – Female | 0.0436 | 0.3024 | 0.144 | 0.887 |
| PMI converted to hours | 0.0101 | 0.0125 | 0.803 | 0.430 |
| pH | -0.1903 | 0.4337 | -0.439 | 0.665 |

[a] Represents reference level

### Linear Regression

Model Fit Measures

| Model | R | $R^2$ |
|---|---|---|
| 1 | 0.747 | 0.559 |

Model Coefficients - DNAJB11

| Predictor | Estimate | SE | t | p |
|---|---|---|---|---|
| Intercept [a] | 22.3861 | 3.0526 | 7.333 | < .001 |
| Arm: | | | | |
| alcohol use disorder – control | -0.1458 | 0.3212 | -0.454 | 0.654 |
| opioid use disorder – control | -0.1577 | 0.3005 | -0.525 | 0.605 |
| opioid+alcohol use disorder – control | -0.6032 | 0.3553 | -1.697 | 0.103 |
| Age Cateogry: | | | | |
| 50-64 – 30-49 | -0.5584 | 0.3202 | -1.744 | 0.094 |
| 50-65 – 30-49 | -0.3808 | 0.4918 | -0.774 | 0.446 |
| 65+ – 30-49 | -0.4427 | 0.3709 | -1.193 | 0.244 |
| <30 – 30-49 | -0.1704 | 0.3878 | -0.439 | 0.664 |
| Ethnicity: | | | | |
| Asian – White | -0.3811 | 0.6457 | -0.590 | 0.561 |
| Black – White | -0.0740 | 0.2629 | -0.282 | 0.781 |
| Hispanic – White | 1.4053 | 0.5028 | 2.795 | 0.010 |
| Gender: | | | | |
| Male – Female | 0.1936 | 0.3107 | 0.623 | 0.539 |
| PMI converted to hours | -0.0348 | 0.0129 | -2.697 | 0.013 |
| pH | 0.3725 | 0.4457 | 0.836 | 0.412 |

[a] Represents reference level

### Linear Regression

Model Fit Measures

| Model | R | $R^2$ |
|---|---|---|
| 1 | 0.576 | 0.331 |

Model Coefficients - DGKB

| Predictor | Estimate | SE | t | p |
|---|---|---|---|---|
| Intercept [a] | 24.82436 | 4.0919 | 6.06668 | < .001 |
| Arm: | | | | |
| alcohol use disorder – control | 0.19515 | 0.4306 | 0.45324 | 0.654 |
| opioid use disorder – control | 0.18579 | 0.4028 | 0.46127 | 0.649 |
| opioid+alcohol use disorder – control | 1.11239 | 0.4763 | 2.33541 | 0.028 |
| Age Cateogry: | | | | |
| 50-64 – 30-49 | 0.29675 | 0.4292 | 0.69147 | 0.496 |
| 50-65 – 30-49 | -0.46131 | 0.6593 | -0.69970 | 0.491 |
| 65+ – 30-49 | 0.07969 | 0.4972 | 0.16028 | 0.874 |
| <30 – 30-49 | 0.15538 | 0.5198 | 0.29891 | 0.768 |
| Ethnicity: | | | | |
| Asian – White | 0.22764 | 0.8656 | 0.26300 | 0.795 |
| Black – White | 0.02594 | 0.3524 | 0.07362 | 0.942 |
| Hispanic – White | -0.05174 | 0.6739 | -0.07677 | 0.939 |
| Gender: | | | | |
| Male – Female | 0.00368 | 0.4165 | 0.00883 | 0.993 |
| PMI converted to hours | -0.01007 | 0.0173 | -0.58281 | 0.565 |
| pH | -0.17762 | 0.5975 | -0.29728 | 0.769 |

[a] Represents reference level

### Linear Regression

Model Fit Measures

| Model | R | $R^2$ |
|---|---|---|
| 1 | 0.591 | 0.350 |

Model Coefficients - EXOC6B

| Predictor | Estimate | SE | t | p |
|---|---|---|---|---|
| Intercept [a] | 25.5314 | 3.2440 | 7.8704 | < .001 |
| Arm: | | | | |
| alcohol use disorder – control | 0.3681 | 0.3413 | 1.0782 | 0.292 |
| opioid use disorder – control | 0.3820 | 0.3193 | 1.1962 | 0.243 |
| opioid+alcohol use disorder – control | 0.8005 | 0.3776 | 2.1198 | 0.045 |
| Age Cateogry: | | | | |
| 50-64 – 30-49 | 0.3755 | 0.3402 | 1.1036 | 0.281 |
| 50-65 – 30-49 | -0.3467 | 0.5227 | -0.6634 | 0.513 |
| 65+ – 30-49 | 0.2770 | 0.3942 | 0.7026 | 0.489 |
| <30 – 30-49 | 0.0546 | 0.4121 | 0.1324 | 0.896 |
| Ethnicity: | | | | |
| Asian – White | 0.2849 | 0.6862 | 0.4152 | 0.682 |
| Black – White | 0.0525 | 0.2794 | 0.1880 | 0.852 |
| Hispanic – White | -0.6079 | 0.5343 | -1.1379 | 0.266 |
| Gender: | | | | |
| Male – Female | -0.0301 | 0.3302 | -0.0911 | 0.928 |
| PMI converted to hours | 0.0120 | 0.0137 | 0.8738 | 0.391 |
| pH | -0.4017 | 0.4737 | -0.8481 | 0.405 |

[a] Represents reference level

### Linear Regression

Model Fit Measures

| Model | R | $R^2$ |
|---|---|---|
| 1 | 0.679 | 0.461 |

Model Coefficients - PPP1R1B

| Predictor | Estimate | SE | t | p |
| --- | --- | --- | --- | --- |
| Intercept [a] | 25.2679 | 6.6706 | 3.7880 | < .001 |
| Arm: | | | | |
| alcohol use disorder – control | 1.5062 | 0.7019 | 2.1459 | 0.042 |
| opioid use disorder – control | 0.6167 | 0.6566 | 0.9392 | 0.357 |
| opioid+alcohol use disorder – control | 1.9352 | 0.7765 | 2.4923 | 0.020 |
| Age Cateogry: | | | | |
| 50-64 – 30-49 | 1.7102 | 0.6996 | 2.4445 | 0.022 |
| 50-65 – 30-49 | 1.1257 | 1.0748 | 1.0474 | 0.305 |
| 65+ – 30-49 | 0.3168 | 0.8105 | 0.3909 | 0.699 |
| <30 – 30-49 | 0.7447 | 0.8474 | 0.8788 | 0.388 |
| Ethnicity: | | | | |
| Asian – White | 1.0966 | 1.4110 | 0.7772 | 0.445 |
| Black – White | 0.5758 | 0.5744 | 1.0023 | 0.326 |
| Hispanic – White | -0.0725 | 1.0986 | -0.0660 | 0.948 |
| Gender: | | | | |
| Male – Female | 0.1040 | 0.6790 | 0.1532 | 0.880 |
| PMI converted to hours | 0.0145 | 0.0282 | 0.5158 | 0.611 |
| pH | -0.1484 | 0.9740 | -0.1524 | 0.880 |

[a] Represents reference level
